# Hypoxia-Induced Cardiopulmonary Remodeling and Recovery: Critical Roles of the Proximal Pulmonary Artery, Macrophages, and Exercise

**DOI:** 10.1101/2025.02.15.638455

**Authors:** Abhay B. Ramachandra, Prapti Sharma, Ruben De Man, Fadi Nikola, Nicole Guerrera, Pramath Doddaballapur, Cristina Cavinato, Rira Choi, Micha Sam Brickman Raredon, Jason M. Szafron, Zhen W. Zhuang, Thomas Barnthaler, Aurelien Justet, Ngozi D. Akingbesote, Nebal S. Abu Hussein, Lonnette Diggs, Rachel J. Perry, Taylor S. Adams, Inderjit Singh, Naftali Kaminski, Xiting Yan, George Tellides, Jay D. Humphrey, Edward P. Manning

## Abstract

Hypoxemia impairs cardiopulmonary function. We investigated pulmonary artery remodeling in mice exposed to chronic hypoxia for up to five weeks and quantified associated changes in cardiac and lung function, without or with subsequent normoxic recovery in the absence or presence of exercise or pharmacological intervention. Hypoxia-induced stiffening of the proximal pulmonary artery stemmed primarily from remodeling of the adventitial collagen, which resulted in part from altered inter-cellular signaling associated with phenotypic changes in the mural smooth muscle cells and macrophages. Such stiffening appeared to precede and associate with both right ventricular and lung dysfunction, with changes emerging to similar degrees regardless of the age of onset of hypoxia during postnatal development. Key homeostatic target values of the wall mechanics were recovered by the pulmonary arteries with normoxic recovery while other values recovered only partially. Overall cardiopulmonary dysfunction due to hypoxia was similarly only partially reversible. Remodeling of the cardiopulmonary system due to hypoxia is a complex, multi-scale process that involves maladaptations of the proximal pulmonary artery.

## Introduction

Congenital heart disease (CHD) comprises myriad structural and functional cardiovascular defects, the most serious of which are life-threatening without early surgical intervention. In most cases, the primary clinical goal is to restore normal blood oxygenation by improving flow through the pulmonary vasculature to the lungs via a series of surgical procedures, including the Glenn at 4-6 months of age and the Fontan at 18-36 months of age. Although life-extending, these palliative procedures leave the patient with a non-physiologic circulation that is characterized by reduced cardiac output and elevated central venous pressure.^1–3^ It is commonly thought that these hemodynamic perturbations predispose to diverse sequelae that contribute to morbidity and pre-mature mortality in this patient population.^1–4^ Yet, another poorly understood contributor to these devastating sequelae is the extended period of hypoxemia prior to Fontan completion – systemic blood oxygen saturation levels are as low as ∼70% prior to the Glenn procedure, then 75-85% following the Glenn, only to be returned to 90-95% following the Fontan procedure.^3,5^ Normal levels are 95-100% while less than 90% is considered low and less than 88% often demands treatment. Toward this end, there is a pressing need to understand better the effects of pre-Fontan hypoxemia on the developing cardiopulmonary system, namely, function of the right heart, pulmonary vasculature, and lungs. Notwithstanding tremendous understanding gleaned from clinical findings, animal models continue to provide significant insight.

There is now an extensive literature on effects of hypoxia on cardiopulmonary remodeling in animal models and much is known,^6–13^ though with diverse findings likely reflecting differences in experimental design: different species, ages at hypoxic onset, durations of hypoxic exposure and recovery, regions of the pulmonary arterial tree, and so forth. Importantly, it remains difficult to separate direct effects of hypoxia from indirect effects, including increased right ventricular and pulmonary artery pressures during hypoxic exposure. For example, there have been reports of no adverse tissue-level remodeling of the proximal pulmonary arteries (PPAs) when nifedipine preserved blood pressure during hypoxic exposure,^14^ suggesting that the vascular remodeling was in response to the pressure rise, not the hypoxia per se. Conversely, it is increasingly thought that hypoxia stimulates early monocyte / macrophage activity,^15^ which has been suggested to drive pulmonary artery remodeling and the subsequent development of pulmonary hypertension.^16–18^ Clearly there is a need to consider these and other factors in any study of hypoxia-induced remodeling and possible recovery thereafter.

There has been considerable emphasis on distal vessels within the lung and the smooth muscle cells (SMCs) within them. In health, SMCs in these muscular pulmonary arteries optimize ventilation-perfusion (V/Q) matching by contracting and limiting blood flow to poorly ventilated alveoli.^19,20^ In ambient hypoxia, SMC contractility occurs diffusely throughout the lung, as if the body senses that all of the alveoli are poorly ventilated, thus pulmonary vascular resistance (PVR) increases acutely.^20^ Less attention has been directed to the proximal conduit arteries, however. PPAs yet play two critical roles: via their elasticity, they endow the pulmonary vasculature with both distensibility (compliance) and resilience (recoil) and via their contractility, they regulate lumen caliber and modulate local blood flow. The normally high distensibility of the PPAs decreases pulse wave velocity (PWV), which reduces the penetration of pulse pressures into the resistance vessels, whereas the normally robust SMC-driven vasoactivity can reduce PPA wall stress during periods of increased pressure and cardiac output, thus reducing the extent of mural remodeling. Together, these two functions help to optimize blood flow to the lungs.

Many studies using animal models report results in either juveniles or adults but, motivated by conditions in CHD, we sought first to contrast effects of early versus later onset hypoxic exposure on cardiopulmonary remodeling and its possible recovery upon a return to normoxic conditions as this sequence mimics the restoration of normoxia after staged surgical interventions ending with the Fontan procedure. Toward this end, we employed a functional mechano-genomics workflow^21^ to quantify effects of hypoxia on right heart function, PPA remodeling, and lung function in neonatal, juvenile, and young adult mice. It has long been known that hypoxia induces adverse remodeling of pulmonary arteries, both proximal and distal.^22^ Characteristic features of such remodeling include adventitial thickening of proximal vessels and so-called hyper-muscularization of distal vessels, thus implicating significant remodeling of adventitial and medial layers, respectively. Our comparison of time-courses of cardiac, vascular, and pulmonary changes suggested that PPA remodeling preceded changes in heart and lung function. Hence, we focused largely on the PPAs, motivated further by the growing awareness of roles of large artery stiffening on cardiac afterload and end-organ damage in both the systemic and pulmonary circulations.^23–25^ Fibroblasts are known to play critical roles in the observed adventitial remodeling in proximal arteries^26^ and interactions between fibroblasts and macrophages are increasingly recognized as important in both homeostasis and disease progression.^27,28^ We thus used single cell RNA-sequencing (scRNA-seq) to investigate possible intercellular interactions between the vascular fibroblasts and macrophages. Finally, we also evaluated possible recovery following restoration of normoxia and potential efficacy of four different treatments administered concurrent with the hypoxic exposure. It is highly concerning that neither normoxic recovery nor targeted treatments were able to preserve or restore normal cardiopulmonary structure and function, suggesting that chronic exposure to hypoxia results in persistent adverse effects.

## Methods

### Research design

The overall study design is summarized in Supplemental Figures S1A and S1B: over 20 groups of neonatal (0-3 weeks old), juvenile (3-8 weeks), and young adult (8-13 weeks) mice include, as appropriate, age-matched normoxic controls, hypoxic, hypoxic with subsequent normoxic recovery without or with voluntary exercise, and hypoxic with concurrent treatment with either different medications or forced exercise. Periods of hypoxia were 2.5 to 5 weeks. At the scheduled endpoint, cardiac and pulmonary function were measured under anesthesia and a battery of tests were performed on the PPAs following euthanasia. Specifically, these vessels were subjected to *ex vivo* biomechanical phenotyping followed by multiphoton microscopy and histology or used to isolate cells for scRNA-seq to identify transcriptional changes.

### Mice

All live animal procedures were approved by the Institutional Animal Care and Use Committee of Yale University and the experimental design followed ARRIVE guidelines. Preliminary studies revealed qualitative similarities between adult female and male mice for multiple metrics of interest. Notable differences were largely due to differences in body mass rather than sex as a biological variable.^29^ Hence, to complement better the extensive data available on effects of hypoxia on male mice, we focused thereafter on female C57BL/6J wild-type (WT) mice (Jackson Laboratory, Bar Harbor, ME and inbred litters). Mice were housed in an antigen-free and virus-free animal care facility under a 12-hour light and dark cycle and fed a standard rodent chow with free access to water. Some groups of hypoxic mice received concurrent treatment with rapamycin (modified LabDiet® 5LG6 with 420 ppm encapsulated rapamycin, irradiated, 9-12% purity, purchased from TestDiet and W.F. Fisher) or metformin (in chow Bio-Serv F10463 Rodent Diet, AIN-93G, Metformin (0.1%), 1/2” Pellets; or 0.5 mg/mL (∼100 mg/kg/day) in the drinking water Sigma Aldrich 317240). Finally, two additional groups of mice used adult female *Ccr2^-/-^* mice (Jackson Laboratory, Bar Harbor, ME and inbred litters). At study endpoints, mice were euthanized via an overdose of urethane by intraperitoneal injection, followed by exsanguination achieved via harvest of the heart, lungs, and PPAs.

### Hypoxic exposure

Mice were exposed to hypoxic conditions by placing them in a BioSpherix A-Chamber connected to a ProOx 360 High Infusion Rate Controller (BioSpherix, Ltd., RRID: SCR 021130, Syracuse, NY, USA). This gas control delivery system regulated the flow of N_2_ into the chamber to maintain a prescribed fraction of inspired oxygen (FiO_2_). Persistent hypoxia defined by a 10% FiO_2_ was maintained 24 hours/day for 2.5 to 5 weeks. When studying neonatal mice, the mother and litter were placed in the chamber together at postnatal day P0 to P1. Age- and sex-matched control mice were kept in similar conditions but exposed to room air (∼20% FiO_2_).

### Voluntary exercise

We monitored voluntary running of select mice with in-cage running wheels (Actimetrics Wireless Low-profile running Wheel Model ACT-557-WLP). Data were collected wirelessly, and daily running was measured using ClockLab Data Collection Software from Lafayette Instrument. Mice naturally run,^30^ and in-cage running wheels have the advantage of providing accessibility to exercise at the convenience of the mice, day and night, without interfering with normal housing conditions for this experiment. We monitored the mice daily for non-participation.

### Exercise capacity (fastest speed)

Maximal running speed was assessed for select mice using the Exer3/6 treadmill (Columbus Instruments, Columbus, OH) with a light shock provided when the animals failed to run and came to the end of the treadmill belt. After 3 days of acclimation to the treadmills, all with a gentle incentivizing shock (0.46 mA) when the mice failed to continue to run (day 1: 10 minutes without the treadmill moving; days 2 and 3: 10 minutes running at 10 m/min), mice were subjected to the maximal speed test. The test began at treadmill speed 10 m/min for 10 min, after which the speed was increased by 1 m/min every minute. The mice were monitored continuously, and maximal running speed was determined to be that speed at which the mice remained on the shock pad without responding to 3 gentle hand brushes.

### Systemic blood analyses

Blood gases, arterial oxygenation saturations, and hemoglobin/hematocrit were measured inside the hypoxia chamber from blood drawn from the descending thoracic aorta for hypoxic mice and in ambient air for normoxic controls. Specially fabricated ports were constructed to pass investigator hands and forearms through the front face of the hypoxia A-chambers without elevating internal ambient oxygen levels above 10 ± 1%. Samples were analyzed using the VetScan i-STAT 1 handheld analyzer (Abbott, Melville, NY). In addition, we used CHEM8+ and CG4+ cartridges (Zoetis, Parsippany, NJ) to obtain the desired combination of parameters for the arterial blood analysis.

### Right heart function

Noninvasive investigation of cardiac function was performed using transthoracic echocardiograms under light anesthesia (1.5% isoflurane) while maintaining physiological temperatures.^31^ Standardized cardiac views were obtained with a high-resolution ultrasound system (Vevo 2100, VisualSonics, Toronto, ON, Canada) equipped with an ultrahigh frequency (40 MHz) linear array transducer. B-mode two-dimensional (2D) images of the right ventricle and right atrium were obtained from an apical four-chamber view, and the PPA was imaged from a parasternal short-axis view at the level of the aortic valve. In addition, M-mode and tissue Doppler imaging (TDI) of the lateral tricuspid annulus were obtained from the apical four-chamber view. The pulmonic valve was visualized at the level of the leaflet tips with pulsed wave Doppler. Right ventricle outflow tract (RVOT) diameter, tricuspid annular plane systolic excursion (TAPSE), right ventricular systolic myocardial velocity (s’), pulmonary artery acceleration time (PAT), pulmonary ejection time (PET), right ventricular free wall thickness, and right atrial area were measured offline using Vevo Lab software (version 3.2.6, VisualSonics) by an experienced sonographer.^32–35^ Cardiac output was calculated using the RVOT and velocity time integral of doppler flow.

### Lung function and histology

Mice were anesthetized via an intraperitoneal injection of urethane (1 g/kg administered in 10% solution with sterile water) and tracheostomized, then connected to the Flexivent system (FlexiVent®, SCIREQ©, Montreal, QC, Canada). Succinylcholine (1 mg/kg) was administered via intraperitoneal injection to eliminate spontaneous breathing. FlexiVent perturbations and oscillations were performed and analyzed using the FlexiWare Version 7.6 software, Service Pack 6 to obtain lung pressure-volume loops, static compliance (Cst), and inspiratory capacity (IC). Maneuvers and perturbations were continued until acquiring three suitable measurements. A coefficient of determination of 0.95 was the lower limit for suitable measurements. An average of three measurements for each parameter was calculated per mouse. Diffusion capacity was measured on select mice using the Inficon MicroFusion Gas Chromatographer (Inficon, Syracuse, NY, USA) with helium carrier gas. 600 µL 0.5% carbon monoxide (CO) 0.5% neon (Ne) nitrogen balanced gas was given to anesthetized and tracheostomized mice that were spontaneously breathing for 9 seconds. After 9 seconds the gas was diluted and analyzed by the gas chromatographer to determine the diffusion fraction of carbon monoxide (DFCO) as follows:

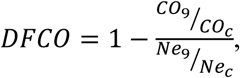

where CO_9_ and Ne_9_ are the concentrations of CO and Ne, respectively, present in the gas mixture after 9 seconds in the mouse lungs and CO_c_ and Ne_c_ are the initial concentrations of the CO and Ne in the mixture (0.5% for each).^36,37^

Immediately following pulmonary function testing, the lungs were removed with tracheal cannulation intact and distended with warm (37°C), 1% weight:volume low melting weight agarose in PBS x 1 at a pressure of 20 cmH_2_O. The agarose solution was allowed to cool, the tracheal cannula was removed, the trachea was ligated, and the stiffened agarose-lung preparation was placed in 10% formalin for 24 hours, then transferred to 70% ethanol. Preparations were embedded in paraffin, sectioned at 5 microns, and stained with hematoxylin and eosin (H&E) for imaging and analysis. Two slides from each lung were digitally imaged at 20x magnification and analyzed. The chord length ImageJ plug-in was used to quantify lung morphometrics such as mean linear intercept or alveolar chord length.^38^ At least five samples of 20x magnification images were analyzed per slide with emphasis on analyzing lung parenchyma to avoid larger airways or vessels.

### Micro-CT of the lung vasculature

A subset of mice (n=5, 2 female 13-week hypoxic and 3 female 13-week normoxic controls) were anesthetized with 2% isoflurane via an oral intubation tube (120 strokes per min). The chest was surgically opened to expose the heart and lung with a midline incision. The pulmonary artery was catheterized with a flared PE 160 tube and secured with a 6-0 silk suture, then 10% Bismuth nanoparticles in 5% gelatin (a custom CT contrast agent) were injected similar to previous descriptions.^39,40^ Briefly, the contrast was injected into the lung at 30 cmH_2_O through the needle after saline perfusion for 8-10 min and 2% PFA for 15-20 min. The lung was covered by ice-cold water to stop the contrast agent from proceeding into the pulmonary venous circulation as previously described. The whole body was further fixed in 2% PFA at 4°C overnight. The lungs were then excised and imaged using a high-resolution micro-CT imaging system (GE eXplore Locus SP), set to a 0.007-mm effective detector pixel size. Micro-CT acquisition parameters were 80-kVp X-ray tube voltage, 80-mA tube current, 3,000-ms-per-frame exposure time, 1 × 1 detector binning model, and 360 and 0.4 increments per view. This acquisition resulted in a set of contiguous VFF-formatted images through the entire lung. The VFF image data were calibrated in standardized Hounsfield unit (HU) with MicroView^TM^ software (GE Healthcare) for quantitative analysis. The calibrated images were further transferred to an AW workstation (GE Healthcare) for volume rendering and multiple format reconstruction. We qualitatively analyzed these images with NeuronStudio 2 to identify branching and overall vasculature geometry. For additional analysis, an open-source software package for vascular morphometric characterization, TubeMap, which is part of the ClearMap2 project, was used to segment the pulmonary arterial tree using a combination of thresholding and adaptive filtering.^41^ As there was some overflow of injected contrast into the airways of several of the right lung samples imaged, we focused this analysis solely on the left lung and manually removed any overfill from segmentations of the left lungs. During skeletonization of the resulting network, automated measurements of radius and length were stored locally for each vascular segment, which were then incorporated into a graph representation of the pulmonary artery tree. From the graph representation, a solver file was generated for a reduced order hemodynamic simulation available as part of the open-source SimVascular project^42^. We then estimated pulmonary vascular resistance (PVR) of the imaged network using standard hemodynamic assumptions (cardiac output of 16.9 mL/min, flow fraction of *FF* = 0.3 to the left lung, uniform outlet pressures of *P*_*o*_ = 4.3 mmHg, fluid density of 1.00 g/mL, and blood viscosity of 0.042 Poise^43–45^) and a Poiseuille flow simulation of pulmonary hemodynamics. Simulation parameters were chosen to be the same across all subjects to isolate effects of morphometry and geometry on PVR, which was calculated as

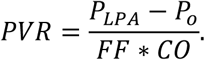

*P_LPA_* was the resulting pressure in the left pulmonary artery from each simulated subject. Additionally, average radius for each generation of vessels was calculated, where the generation number started at 1 for the LPA and incremented by 1 with each branching of the network.

### Biomechanical phenotyping of pulmonary arteries

Following euthanasia, arterial specimens were excised from the main pulmonary artery to the first branch of the right pulmonary artery as previously described.^46^ Perivascular tissue and fat were removed and the left pulmonary artery and small branch vessels were ligated with suture after flushing luminal blood with a Hank’s buffered physiologic solution (HBSS).^47^ The right pulmonary artery was cannulated on custom glass micropipettes and secured with ligatures beyond the main pulmonary artery on one end and the first branch at the other end. Each specimen was submerged in room temperature HBSS to ensure passive behavior, then tested using a custom computer-controlled testing device imposing a series of three cyclic pressure-distension and four axial force-extension protocols to quantify biaxial mechanics—mechanical quantities of interest such as circumferential and axial wall stretch, stress, and material stiffness as well as local distensibility were subsequently calculated from biaxial data and constitutive fits to the data.^48^

Since residual stresses tend to homogenize the stress field, we used a 2-D formulation to quantify the basic mechanical behaviors thereby rendering mean values as good estimates of overall wall stress.^49^ Mean wall stresses in circumferential (θ) and axial (z) directions were determined experimentally as

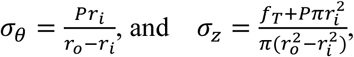

where *P* is the transmural pressure, *r*_*i*_ internal radius, *r*_*o*_ outer radius, and *f*_*r*_ the transducer measured axial force. Under the assumption of incompressibility, inner radius

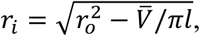

where *l* is the instantaneous length between the ligatures securing the vessel to the micropipettes and *V̄* is the volume of the vessel in the unloaded state; *V̄* = *πL*(*OD*^2^ − *ID*^2^)/4 where *L* is the unloaded length, *OD* the unloaded outer diameter, and *ID =OD-2H,* the unloaded inner diameter. Mean biaxial wall stretches are

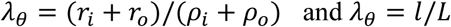

where *ρ*_*i*_ and *ρ*_*o*_ are inner and outer radii in the unloaded configuration. Finally, *D* is the distensibility coefficient (in kPa^-1^ or mmHg^-1^) determined by the end-diastolic inner diameter-normalized change in arterial diameter from end-diastole to end-systole divided by the change in end-diastolic and end-systolic pressures.^50^

### Material characterization of pulmonary arteries

Circumferential wall stress and material stiffness are highly mechano-regulated quantities, whereas distensibility and elastic energy storage are fundamental functional readouts.^51^ Full biomechanical phenotyping thus requires calculation and comparison of multiple metrics for each sample. Following prior work^46^, we further characterized the biaxial biomechanical data from the aforementioned seven different cyclic testing protocols using a single nonlinear (pseudo)elastic stored energy function *W* that has successfully described passive biaxial behaviors of PPAs, namely

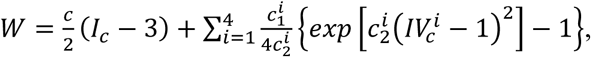

where *c* (kPa), 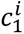 (kPa) and 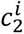 (-) are material parameters; *i*=1,2,3,4 represent four collagen-dominated families of fibers along axial, circumferential, and two symmetric diagonal directions, respectively. *I*_*c*_ is the first invariant of right Cauchy-Green tensor (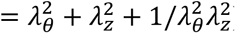) and 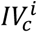 is the square of the stretch of the *i*^*th*^ fiber family (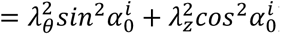) where 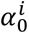 is the fiber angle relative to the axial direction in the reference configuration. Best-fit values of the material parameters and the fiber angle were determined via nonlinear regression of biaxial data from all seven passive protocols. More details on parameter estimation can be found elsewhere,^46^ noting that the associated biaxial wall stress and material stiffness can be computed from first and second derivatives of *W* with respect to an appropriate deformation metric.

Local values of pulse wave velocity (PWV) depend on both the geometry and mechanical properties of the artery and can be approximated based on material stiffness or distensibility, as, for example,

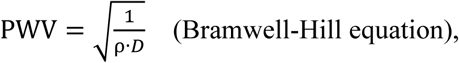

where *ρ* is the mass density of the contained fluid (approximately 1,050 kg·m^-3^) and *D* is the aforementioned distensibility coefficient (in Pa^-1^ or kg^-1^m·s^2^). PWV in the pulmonary artery can thus be estimated non-invasively.^52–54^

### Active mechanical testing of pulmonary arteries

For select PPAs, we measured smooth muscle contractility with an active protocol prior to the passive protocol used to measure biomechanical properties as described above. The specimen was submerged in a 37°C bath of Krebs-Ringer solution perfused with 95% O2 / 5% CO2 during testing with our custom computer-controlled testing device.^46^ We focused our active testing on SMC contractile responses near normal *in vivo* conditions, that is, a mean intraluminal pressure of 15 mmHg and the specimen-specific value of *in vivo* axial stretch as previously described,^21,55^ which more closely phenotypes *in situ* physiology of the pulmonary artery than “ring-tests” performed with uniaxial myographs.^56^ The SMCs were initially conditioned at 10 mmHg and an axial stretch of 1.1 with respect to the specimen-specific unloaded length by exposing the vessel to 100 mM potassium chloride (KCl) in the bath two times and returning the bath to normal Krebs-Ringer solution in between. Following conditioning, the axial stretch was increased slowly in steps of 0.01 and pressure was increased slowly by 1 mmHg increments to avoid rapid large deformations that might compromise viability of the SMCs. The vessels were then exposed to 100 mM KCl or 1 µM phenylephrine (PE) at 15 mmHg and subject-specific *in vivo* axial stretch. This protocol comprised 5 min equilibration, 15 min contraction, and 10 min relaxation after washout of the vasoactive substance with fresh, warm Krebs-Ringer solution.

### Multiphoton imaging of pulmonary arteries

A multiphoton microscope was used to image representative regions of PPAs at *in vivo* relevant loading conditions (*in vivo* axial stretches and distending pressures used during passive mechanical testing). Specifically, a Titanium-Sapphire Laser (Chameleon Vision II, Coherent) and LaVision Biotec TriMScope microscope equipped with a water immersion 20X objective lens (NA. 0.95) was tuned at 840 nm. The backward scattering second harmonic generation signal from fibrillar collagens was detected within the wavelength range 390-425 nm, the auto-fluorescent signal arising from elastin was detected at 500-550 nm, and the fluorescent signal of cell nuclei labelled with a Syto red stain was detected above 550 nm. Each in-plane field of view (axial-circumferential plane) was 500 µm x 500 µm with a volume of about 0.05 mm^3^; this provides a much greater volume of tissue for imaging than via standard histology and hence averaging over significantly greater numbers of cells and extracellular matrix. The in-plane resolution was 0.48 µm/pixel and the out-of-plane (radial direction) step size was 1 µm/pixel. 3D images acquired concurrently for the three signals (collagen, elastin, and cell nuclei) were post-processed using MATLAB R2019b and ImageJ 1.53a. The first processing step relied on the near cylindrical shape of the samples to fit a circle to the two-dimensional mid-thickness profile of the arterial wall and transform each circumferential-radial slice of the 3D images from Cartesian to polar coordinates (angle and radius). This allowed a layer-specific microstructural analysis to focus on collagen fiber alignment and cell volume density analyses, as described previously.^57,58^

### Histology of pulmonary arteries

Following biaxial testing and multiphoton imaging, the PPAs were fixed in 10% neutral buffered formalin and stored in 70% ethanol at 4°C for histology. After embedding in paraffin, they were sectioned (5 µm thickness) into radial-circumferential cross-sections (planes) and stained with Verhoeff Van Giesen (VVG) or Masson’s Trichrome (MTC). Details of image quantification can be found elsewhere.^59^ Briefly, each section was imaged with an Aperio AT2 High Capacity Digital Pathology Scanner under a 40x magnification objective. Following background subtraction and pixel-based thresholding, area fractions for elastin (from VVG) and cytoplasm (from MTC) were computed as the ratio of pixels corresponding to a stain divided by the total number of pixels in the image. Because MTC can overstain collagen, its area fraction was computed as 1 - area fraction of elastin + cytoplasm, with GAG content assumed negligible^60,61^ as confirmed with Movat Pentachrome staining (data not shown). Three sections were analyzed per vessel per stain.

### Single cell RNA sequencing (scRNA-seq) and analyses

We analyzed viable cells from non-biomechanically tested main pulmonary arteries of mice exposed to the various conditions (normoxic, hypoxic, hypoxic with normoxic recovery without or with exercise, as shown in Supplemental Figures S1 and S2). Following euthanasia, the excised pulmonary arteries were chopped mechanically and placed in 1.5 mg/ml collagenase (Gibco), 3 U/ml elastase (Worthington), 1 mg/ml DNASE I (Roche), 1.5 mg/ml dispase (Sigma-Aldrich), and 1 mg hyaluronidase (Sigma-Aldrich). Following cellular dissociation, we barcoded unique mRNA molecules of each cell using the 10x Genomics Chromium platform (3’ v3.1 kit), a droplet-based microfluidic system, and performed reverse transcription, cDNA amplification, fragmentation, adaptor ligation, and sample index PCR according to the manufacturer’s protocol. High sensitivity bioanalyzer traces of cDNA after barcoding and of the final cDNA library were evaluated for quality control. The final cDNA libraries were sequenced on a HiSeq 4000 Illumina platform in our core facility aiming for 150 million paired-end reads per library, at the manufacturer’s recommended read 1 and read 2 lengths. Raw sequencing reads were demultiplexed based on sample index adaptors, which were added during the last step of cDNA library preparation.

Read 2 5-prime anchored template-switch oligo and 3-prime poly(A) contamination were removed using cutadapt. Trimmed reads were mapped to the GRCm38 genome with GENCODE reference annotation release M22 (GRCm38.p6) with STARsolo. Data were analyzed and visualized with R using the Seurat package. Specifically, we clustered the cellular transcriptomes and visualized them in a uniform manifold approximation and projection (UMAP) space to delineate cell types (as shown in Supplemental Figure S3). To identify aberrant gene expression profiles in cellular subpopulations, we established differentially expressed genes between normoxic and hypoxic cells using the non-parametric Wilcoxon rank sum test with Bonferroni correction for multiple testing error. We then modeled changes in expression with age and hypoxic state as predictor variables using a generalized linear mixed effect regression model to identify differentially expressed genes (DEGs) with expression changes correlated with hypoxic exposure. Gene set enrichment analysis of these genes was conducted using PantherDB Gene Ontology (GO) search algorithm for biological processes of mouse genes. Statistically significant GO processes, their accession numbers, description and DEGs from our data associated with the enriched GO terms were identified.

To identify intercellular communications among SMCs, fibroblasts (FBs), and macrophages (MΦs), connectomic analysis in scRNA-seq data was performed using the NICHES package in R.^62^ NICHES computes cell-cell interactions by multiplying expression of ligand (in sender cell) with expression of receptor (in receiving cell). The scRNA-seq data were imput with the ALRA algorithm using genes expressed in at least 50 cells. Ligand-receptor lists from FANTOM5 were used to filter for biologically relevant signaling mechanisms. Cell-to-cell signaling was calculated using NICHES for each sample individually, and later merged across samples. The resulting dataset was filtered to include only cell pairs with non-zero connectivity for at least 100 mechanisms. Cell signaling data were scaled followed by principal component analysis (PCA) and UMAP calculation. Clustering was performed using KNN with a cluster detection sensitivity parameter of 0.3. Clusters were assessed for enrichment of conditions using a Fisher Exact Test. Finally, a hypoxia signaling score was calculated using the ‘AddModuleScore’ function in Seurat. Significant sender-receiver pairs for each condition (normoxia and hypoxia (Nox-Hox), hypoxia and normoxic recovery (Hox-NoxRec), and hypoxia and normoxic recovery including voluntary exercise (Hox-NoxRecExer)), were analyzed for gene enrichment similar to above. We compared the numbers of similar and differing sender and receiver genes found in the signaling archetypes across conditions and summarized these results as a Venn diagram. Based on our enrichment analysis of these genes relevant to SMC-FB-MΦ intercellular communication, and similar genes identified in the literature, we grouped ligand-receptor interactions into three categories to determine if they returned to pre-hypoxic levels after chronic hypoxic exposures: extracellular matrix (ECM) production (*Tgfb2, Bmpr2, Col1a1, Col18a1, Col15a1, Itgb1*);^63,64^ ECM degradation (*Serpine1, Serpine2, Mmp14, Tgm2, Mmp9, Lgals3bp, Mif*);^65^ and immune cell signaling and activity (*Hif1a, Il1rl2, Il6st, Il1a, Tnf, Tnfrsf1a, Il1rap, Il10, Il10rb, Il4, Alox5, Alox5ap, Icam1, Cxcl12, Ccl2*).^64^

### Statistics

Statistical tests were performed using GraphPad Prism version 7.01 for Windows, GraphPad Software, La Jolla California USA, www.graphpad.com. Differences between hypoxic and normoxic groups in morphological, functional, and mechanical properties were determined by a two-tailed, unpaired *t* test. For categorical variables with more than two categories, levels of stretch or pressure were compared using a two-factor analysis of variance (ANOVA). A global test across all levels was performed and then pair-wise comparisons were conducted with post-hoc tests using Bonferroni correction. A *p <* 0.05 level of significance was used, with data reported as mean ± standard error from the mean (SEM). Longitudinal data were fit with a four-parameter logistic regression model using cohort-specific best-fit parameters, which we used previously to model developmental characteristics and homeostasis of the cardiovascular system of this strain of mice.^29,66^ Sample sizes for experiments are shown in Supplemental Table S1.

## Results

### Hypoxia-induced cardiopulmonary dysfunction is independent of age of onset

Right ventricle (RV) function was attenuated following 5 weeks of continuous hypoxia (10% FiO_2_) in juvenile (hypoxic onset at 3 weeks of age) and adult (onset at 8 weeks of age) mice as revealed by significantly decreased values (*p*<0.01) of the ratio PAT:PET of pulmonary artery acceleration time (PAT) to pulmonary ejection time (PET) and systolic myocardial velocity (s’; albeit not in adults). Notwithstanding some increases in both of these metrics with normal postnatal development, ∼10% hypoxia-induced reductions of RV function were similar regardless of age of onset of hypoxic exposure (Figure 1A, Supplemental Figure S4). We also observed significantly increased RV free wall thickness (*p*=0.04) in the juvenile mice and RV cardiac output (*p*=0.01) in the adult mice despite no significant difference in right atrial area (Supplemental Figure S5) consistent with compensated RV failure.^67,68^

**Figure 1.**
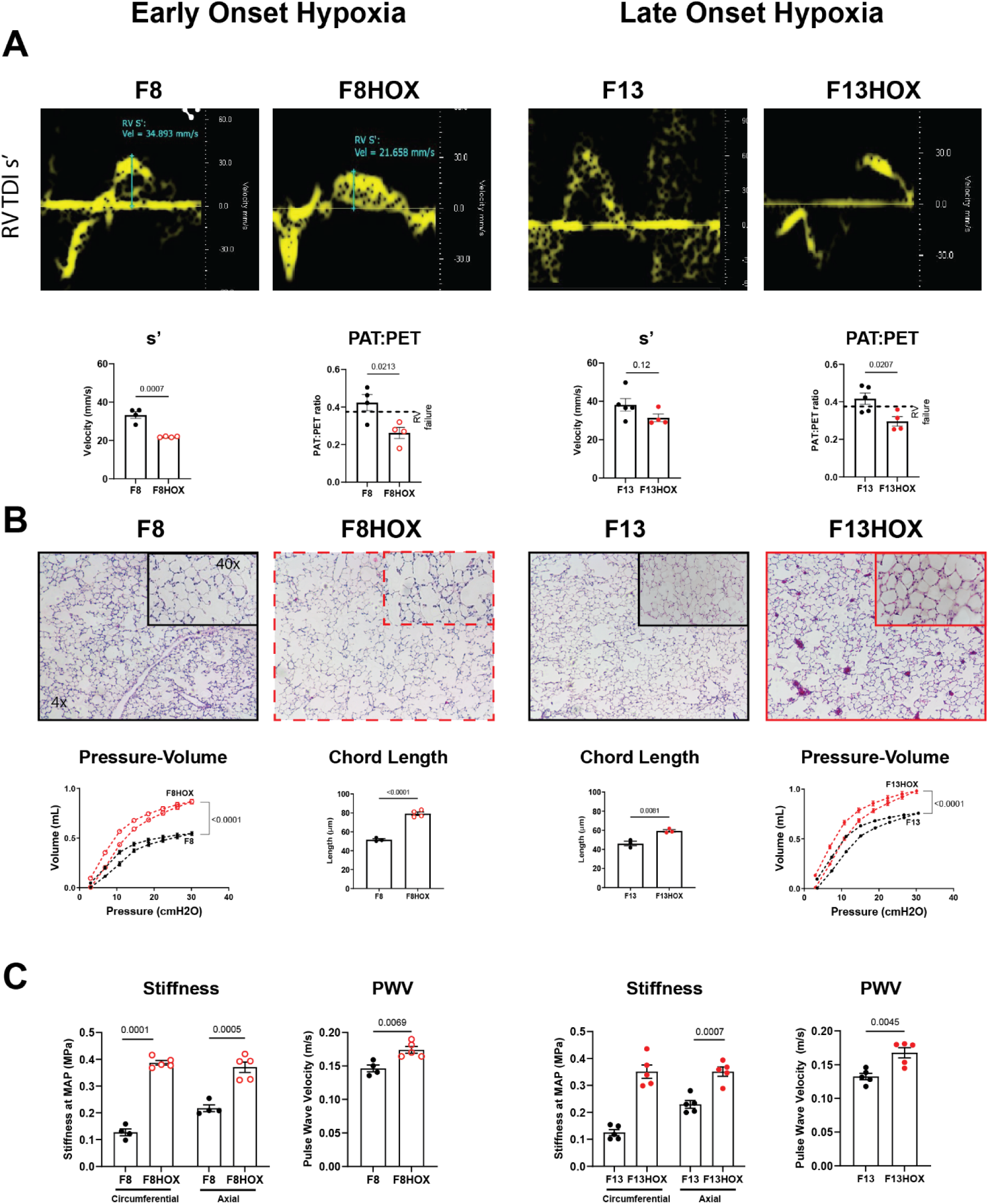
Adverse effects of 5 weeks of inhaled 10% FiO_2_ (hypoxia - HOX) in female WT mice on right ventricular (RV) (panel A), lung (panel B), and proximal pulmonary artery (panel C) function for both early onset (HOX from 3 to 8 weeks of age) and later onset (HOX from 8 to 13 weeks of age) hypoxia. Cardiac metrics are s’ (the peak rate of systolic contraction of the lateral aspect of the tricuspid annulus, a measure of RV function measured by tissue doppler echocardiography) and PAT:PET (the ratio of pulmonary artery acceleration time (PAT) to pulmonary ejection time (PET)), an echocardiographic measure of pulmonary arterial pressures where a lower ratio equates to a decreased ability of the RV to overcome higher PA pressures; PAT:PET < 39% associates with RV failure in mice. Lung metrics are pressure-volume relationships determined by pulmonary function testing and mean linear intercept calculations (chord length) of histologic slices of lungs, representative images of which are shown. Proximal pulmonary artery metrics include intrinsic circumferential and axial material stiffness and pulse wave velocity (PWV).

Similarly, lung structure and function were compromised in these two age groups of mice (Figure 1B). Histological analysis of alveolar chord length demonstrated significant hypoxia-induced increases independent of age of onset (*p*<0.01), which were reflected by increased lung compliance at the organ level as revealed by volume-pressure relationships (*p*<0.01). Similar differences were seen in inspiratory capacity (Supplemental Figure S6, *p*<0.01). Again, changes were similar in both age groups, though slightly more severe in the juvenile mice (Figure 1B). We found similar changes in lung function in neonatal mice exposed to hypoxia (F3HOX, Supplemental Figure S7).

Two key metrics of PPA health are the intrinsic circumferential material stiffness and PWV, the latter of which incorporates both material stiffness and geometry (luminal radius and wall thickness). Both metrics increased significantly (*p*<0.01) following 5 weeks of continuous hypoxia, again independent of age of onset (Figure 1C). It is important to note that both of these metrics are highly pressure dependent, thus they were evaluated at *in vivo* relevant mean arterial pressures of 15 mmHg (normoxia) and 25 mmHg (hypoxia) based on measurements of RV pressure (Supplemental Tables S2 through S4).^21^ PPA distensibility decreased significantly in both juvenile and adult mice after 5-week exposures to hypoxia (juvenile normoxia versus hypoxia: 336.66 ± 23.52 versus 198.21 ± 11.02 mmHg^-1^, *p*<0.01; adult: 387.44 ± 36.69 versus 217.57 ± 21.24 mmHg^-1^, *p*<0.01; Supplemental Table S3).

Given the potential importance of altered cardiopulmonary development prior to Fontan completion in children, we next assessed heart, pulmonary artery, and lung function in neonatal mice, finding that continuous hypoxic exposure from birth to weaning at 3 weeks of age similarly compromised cardiac, vascular, and pulmonary function (Supplemental Figures S4 and S7 and Supplemental Tables S5 and S6). Therefore, chronic hypoxia associated with age-independent functional impairment in the three primary components of the cardiopulmonary system: the right ventricle of the heart, the lungs, and the proximal pulmonary arteries that buffer their interactions. See Supplemental Tables S5 and S6 for further quantification of the biomechanical phenotype of the neonatal PPA under normoxic versus hypoxic conditions (restricted to the neonatal period) as well as from birth to 8 weeks of age (i.e., continuing through the juvenile period). Age-matched comparisons reveal compromised geometry and properties in both cases.

### Compromised pulmonary artery function precedes cardiopulmonary changes

To investigate the time-course of changes during the hypoxic period, we collected additional data at 2.5 weeks of hypoxia that was initiated at either 3 (in juveniles) or 8 (in adults) weeks of age. Both cardiac and pulmonary dysfunction emerged progressively, with differences from age-matched normoxic controls smaller after 2.5 weeks and greater after 5 weeks of hypoxia (Figure 2A). By contrast to this gradual worsening in function of the heart and lungs, with few exceptions, changes in PPA mechanics tended to differ at 2.5 weeks at the same levels seen at 5 weeks (Figure 2B; Supplemental Tables S2 through S4), suggesting a more rapid response by these pulmonary vessels. A similarly rapid response was seen in both increased RV blood pressure (Supplemental Figure S8) and increased hematocrit (Supplemental Figure S9). Taken together, these results suggest that, similar to findings in the systemic hypertension and aging^69,70^ and humans with pulmonary hypertension,^71^ changes in conduit artery function and properties can precede those at the organ level, here in the heart and lungs.^72^

**Figure 2.**
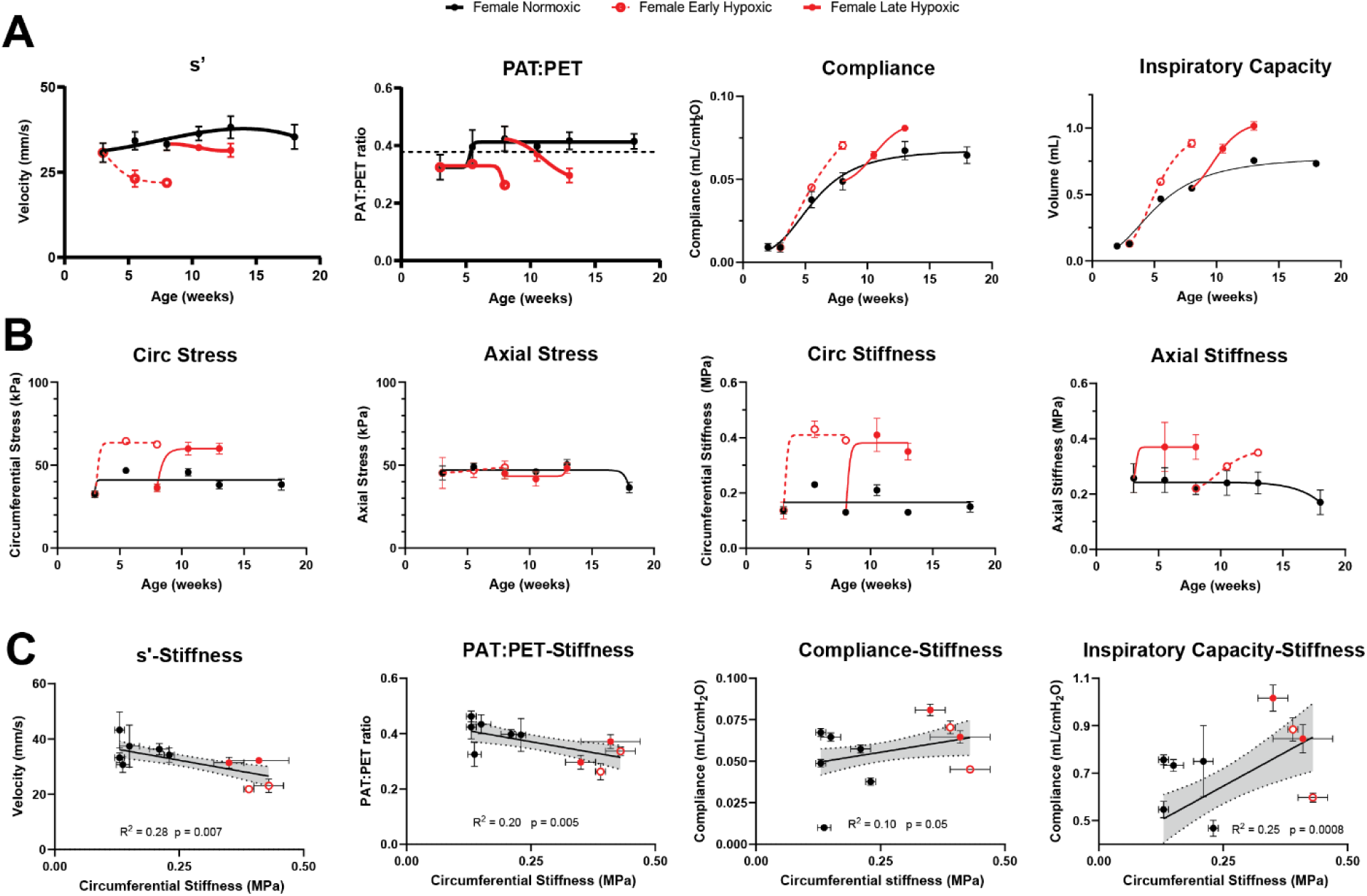
Similar to Figure 1 except time-course studies for early onset (HOX from 3 to 5.5 to 8 weeks) and later onset (HOX from 8 to 10.5 to 13 weeks of age) for RV and lung function (panel A), proximal pulmonary artery metrics (panel B), and correlations therein (panel C).

We thus evaluated possible correlations between both altered heart function (indicated by s’ and PAT:PET) and lung function (indicated by static compliance and inspiratory capacity) and the intrinsic circumferential stiffness of the PPA that increased rapidly following hypoxic exposure. As it can be seen (Figure 2C), decreases in both cardiac and pulmonary functional metrics correlated with increases in intrinsic PPA stiffness (*p*<0.05). We hypothesized that the association of increased PWV in the pulmonary circulation due to chronic hypoxia with worsening pulmonary metrics resulted from loss of functional gas exchange units at the termini of the pulmonary vasculature and airways. Select micro-CT imaging of the pulmonary vasculature supported this hypothesis. Chronic hypoxia causes a loss of microvasculature in the lung, also known as pruning, a phenomenon associated with increased PVR, RV failure, and increased mortality in humans diagnosed with pulmonary hypertension and emphysema.^73–77^ We found a 2-fold increase in PVR resulting from the reduced number of distal pulmonary vessels. We suspected that loss of microvasculature in the lung and increased chord length of the alveolar space indicated a loss of fundamental gas exchange units within the lungs of mice exposed to chronic hypoxia. Consistent with this hypothesis, the DFCO decreased significantly (F13 = 76%, F13HOX = 69%, *p*=0.01) following later onset chronic hypoxic exposure and the emphysematous changes of lung parenchyma^78,79^ (Supplemental Figure S10). Given that our findings are consistent with PPA changes preceding and driving organ-level changes, we focused thereafter primarily on more detailed assessments of PPA structure and function.

### Hypoxia-induced adventitial remodeling associates with adventitial collagen fiber re-orientation

To determine reasons underlying the observed age-independent stiffening of the PPAs in response to hypoxia, we contrasted multiple geometric, microstructural, and mechanical metrics as a function of time of onset (juvenile vs. adult) and duration (2.5 vs. 5 weeks) of continuous exposure to 10% FiO_2_. These vessels exhibited a significant loaded mural thickening in response to hypoxia, slightly more in the later onset group (*p*=0.05 in juvenile mice relative to age-matched controls, *p*<0.05 in adult mice; Figure 3A) though not sufficient to restore circumferential wall stress to normal values (Supplemental Table S3) given the attendant increase in blood pressure. These arteries also showed significant diffuse dilatation in both hypoxic groups (*p*<0.05); the percent increase of inner radius was slightly greater in the later onset group (<7% in juvenile mice, >10% in adult mice).

**Figure 3.**
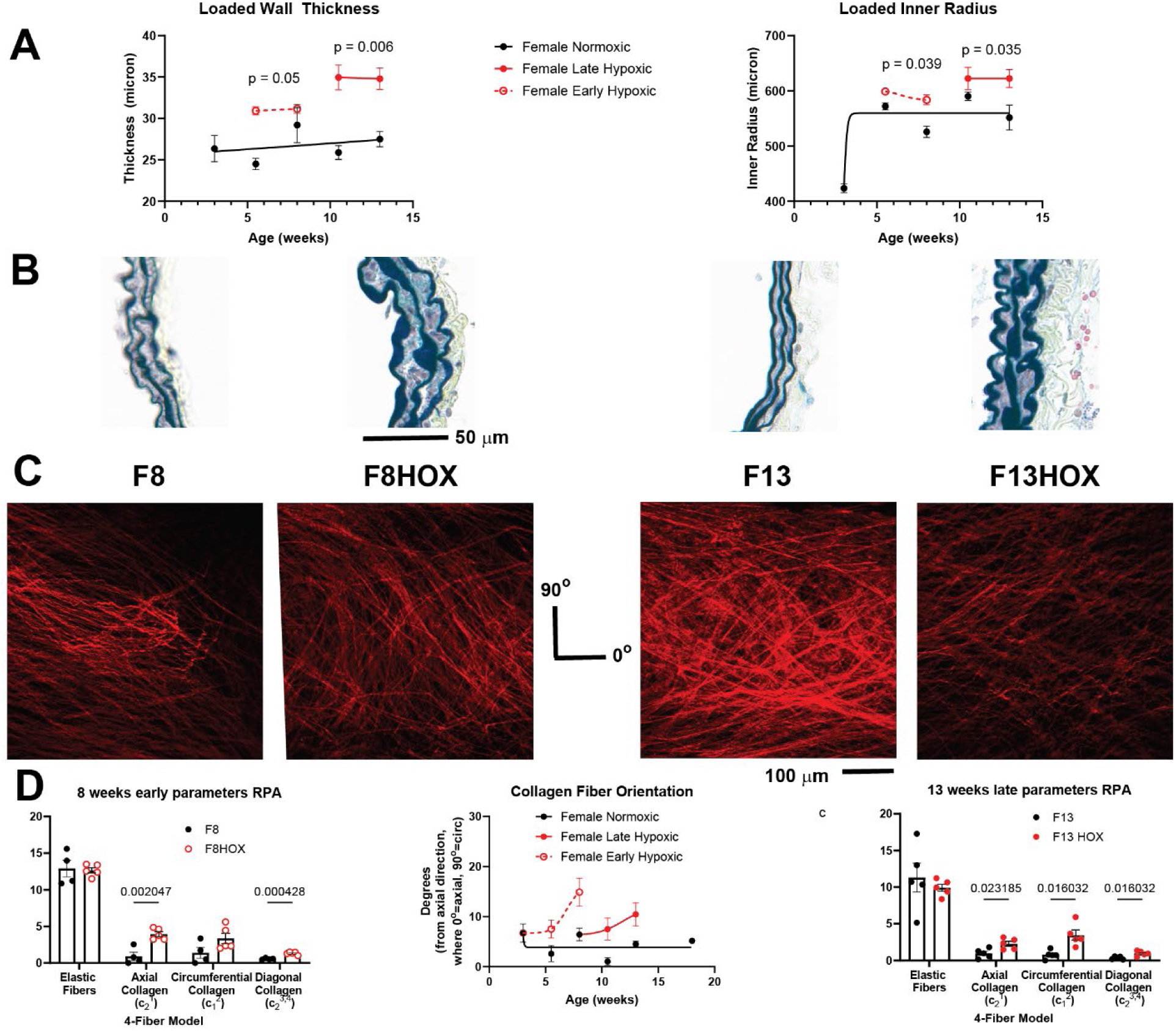
Hypoxia-induced changes in proximal pulmonary artery wall thickness and luminal radius (panel A), basic histological structure (panel B), collagen organization detected via second harmonic generation in multiphoton microscopy (panel C), and hypoxia-related changes in material parameters that describe the mechanical behavior of the proximal pulmonary artery (panel D).

Notwithstanding the marked increases in wall thickness due to hypoxia, standard histological examination revealed no major compositional changes to the pulmonary arterial wall, including no significant differences in elastin fiber content (Figure 3B). This finding was confirmed by detailed quantification of the biaxial mechanical properties, namely, no significant differences in the elastin-associated material parameter (Figure 3D, Supplemental Table S2). By contrast, second harmonic generation multiphoton images of adventitial collagen revealed a progressive hypoxia-associated reorientation by ∼10 degrees towards the circumferential direction (Figure 3C), which would serve to reduce distensibility. Such reorientation is also seen in cases of aortic dilatation, seemingly to help prevent increasing diameter.^80^ This measured reorientation was consistent with the independently quantified biaxial material properties, with material parameters associated with axial and symmetric diagonal (toward axial) collagen fibers greater in a hypoxic group (Figure 3D). This change in adventitial collagen orientation was similar regardless of age of onset of hypoxic exposure.

### Hypoxia-induced transcriptional changes in the pulmonary arteries affect metabolism, intercellular signaling, ECM re-organization, and cellular migration

Given similar tissue-level changes of the PPAs independent of age of onset of hypoxia, we hypothesized the existence of common molecular and cellular responses to hypoxia independent of underlying postnatal development.^81,82^ To test this hypothesis, we performed scRNA-seq at different stages of normal development (3, 5.5, 8, 10.5, and 13 weeks) and hypoxic exposure (for 2.5 versus 5 weeks when initiated at 3 or 8 weeks of age) to identify common transcriptomic signatures for cells that typically reside within the PPA. In particular, we sought insight into molecular-cellular processes involved in the observed adventitial remodeling; thus, we focused primarily on FBs and MΦs though SMCs as well. We identified genes for each cell type whose expressions were common amongst the hypoxic samples when compared with their age-matched normoxic controls (Figure 4A, Supplemental Tables S7 through S9). Biological processes significantly enriched in these genes (Figure 4B; Supplemental Table S10) reflected changes in metabolism / glycolysis (GO:0061621), extracellular matrix reorganization (GO:0030198), collagen fibril organization (GO:0030199), cell migration and adhesion (GO:0007155), regulation of intercellular signaling cell-cell communication (GO:0010646, GO:0023051), cell proliferation (GO:0042127), and vasculature development (GO:0001944). General changes in the organization of ECM, including collagen specifically, clearly reflected the multiphoton and standard histological findings.

**Figure 4.**
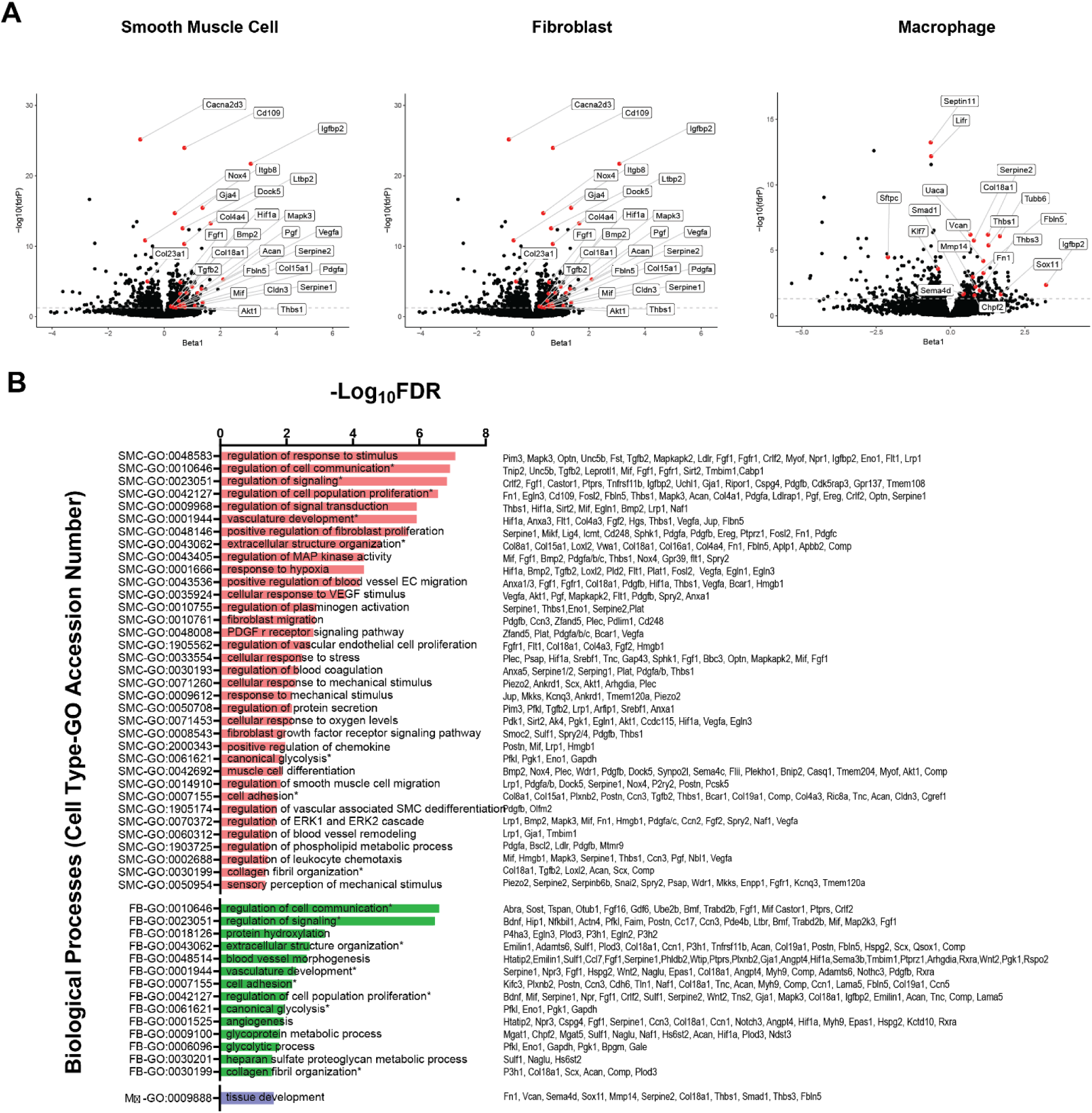
Differentially expressed genes by cell type that resulted from scRNA-seq analysis visualized as volcano plots (panel A, y-axis = significance (-log_10_(false discovery rate)), x-axis = gene expression association with hypoxia (β)). Gene enrichment analysis with biological processes for each cell type shown and key differentially expressed genes from our analysis that associated with each process (panel B). * = biological process present in more than one cell type though expressed gene associated with this process may vary from cell to cell.

We also found common molecular pathways through gene enrichment analysis, including transforming growth factor-beta (WP113 including genes *Smad1, Thbs1, Mapk3, Fst, and Serpine1*), endothelial growth factor receptor signaling (WP572, including *Hip1, Ralb, Tnk2, Inppl1, Ap2a1, Pld2, Gja1, Akt1, Spry2, Git1, Bcar1, Plec, Mapk3, Map2k3, Plec, Mapk3, Grb14*), interleukin 17 signaling (WP5242, with genes *Upf1, Gap43, Arpc2, Pdgfb, Mcm6, Mmp14, Fn1, Prkacb, Gap43, Akt1, Snai2, Ccn2, Pak3*), and a range of nutrient-sensing and biosynthesis pathways such as PI3k-Akt-mTOR (WP2841 and WP3855, including genes *Sreb1, Lama5, Angpt4, Epas1, Pdgfa, Pdgfb, Tnc, Fgf1, Hif1a, Efna4, Comp, Fgf16, Itga7, Mapk3, Csf1r, Fn1, Thbs1, Thbs3, Tnc, Fgf1, Vegfa, Col4a4, Akt1, Fgfr1, and Flot2*), insulin signaling (WP65, including genes *Map2k3, Pfkl, Vamp2, Mapk3, Grb13, Flot2, Rhoq, Inpp4a, Pfkl, Kif3a, Inppl1, Akt1, Enpp1, Map3k8, Mapk6, Vamp2, Mapk3, Grb14*), and cholesterol metabolism (WP4346 and WP2084: including genes *Acat1, Srebf1, Sqle, Acsl1, Pmvk, and Ldlr*). Some cell types shared increased expression of genes associated with particular biological processes; for example, processes common to FBs, MΦs, and SMCs included increased expression of genes that associate with tissue development. Common genes for these processes in these resident cell types included *P3h1, Hif1a, Comp, Fbln5, Thbs1,* and *Sost*, with other genes including *Thbs3, Bmp2, Fosl2, Smad1, Sema4d, Fbln5, Mmp14*, and *Mapk3*.

Both FBs and SMCs increased expression of genes associated with glycolytic metabolism (GO:0061621, *p*< 0.05), including *Pfkl*, *Eno1*, *Gapdh*, and *Pgk1*. FBs and SMCs also shared processes associated with collagen proteostasis and extracellular matrix remodeling, including extracellular structure organization (GO:0043062) and collagen fibril organization (GO:0030199). Many of these genes have been described previously in matrix remodeling in the vasculature and tissue types other than the PPA, including: *Plod3*, *Col18a1, Loxl2, Tgf 2, and Emilin.*^83–89^ SMC gene expression during chronic hypoxia also associated with processes involving responses to low oxygen environment and increased luminal pressures that increase transmural stress and intercellular communication. These processes include responses to hypoxia (GO:0001666, including genes *Egln3, Bmp2, Flt1, Hif1a, Tgfb2, Pgf, Loxl2, Vegfa, Fosl2, Akt1, Engl1, Pgk1, Plat, Pdk1,* and *Sirt2*), glycolysis (GO:0061621 with genes including *Pfkl, Eno1, Gapdh, and Pgk1*), mechano-transduction and cell response to mechanical stimuli (GO:0009612, GO:0033554, GO:0050954 and GO:0071260, with genes including *Piezo2, Mapk3, Mif, Hif1a,* and *Ankrd1*), altered signal transduction (GO:0009968), PDGF receptor signaling (GO:0048008), chemokine CXCL2 production (GO:2000343), endothelial and fibroblast migration (GO:0043536, GO:0010761 with genes including *Col18a1, Fgf2, Fgfr1, Thbs1, Pdgfb, Ccn3, Plec, Akt1, Zfand5, Anxa1, and Pdlim1, and Hmgb1*), and blood coagulation (regulation of plasminogen activation GO:0010755, involving genes *Serpine1, Thbs2, Eno1, Serpine2,* and *Plat* and regulation of blood coagulation GO:0030193).

As a result of chronic hypoxia, FBs typically associated with collagen deposition and expressed genes associated with extracellular organization through both degradation and collagen arrangement (GO:0030198 with genes *Adamts6, Col18a1, Plod3, Can* and GO:0018126 including genes *P4ha3, Egln3, Plod3, P3h1,* and *P3h2*) as well as proteoglycan metabolism and protein hydroxylation (GO:0030201 and GO:0018126 including genes *Can, Plod3, and Egln2*). We found no significant difference in *Col1a1* or *Col3a1* in FBs associated with chronic hypoxia, suggesting either that collagen deposition remained relatively constant between normoxic and hypoxic conditions or other cells were responsible for possible deposition. FBs also expressed genes in response to the low oxygen environment (GO:0006096 with genes including *Pfkl, Eno1, Gapdh,* and *Pgk1*), cellular adhesion (GO:0007255 with genes *Cdh6, Postn, Ccn3, Fbln5, and Col19a1*), and proliferation (GO:0042127 with genes *Bdnf, Mif, Wnt2, Mapk3, Fgf1*). We also found that FBs exposed to chronic hypoxia expressed genes associated with angiogenesis (GO:0001525), including *Serpine1, Ccn1, Col18a1*, and *Notch3, Angpt4*.^90^

MΦs often orchestrate remodeling in response to stress and insults, including increased mechanical loading, injury, or infection.^91–98^ Genes with increased expression in hypoxia by MΦs were enriched in differential tissue development (GO:0009888). We found increased expression of genes related to protease activity, including: *Mmp14*, which interacts with *Mmp2* and *Timp2* and associates with degradation of extracellular matrix and changes in cell adhesion that affect cellular motility;^99–101^ *Serpine2*, which associates with matrix regulation and cardiac fibrosis;^102–104^ *Thbs3*, which associates with altered matrix in cardiomyopathies and aortic aneurysm;^105–109^ *Smad1*, critical in the extracellular control of TGFβ signaling for vascular maintenance;^110^ and *Sema4d*, which is a mediator of cellular motility and angiogenesis in the context of malignancy but not previously noted in vascular remodeling.^111,112^ We also found a common increase in expression of *Fbln5* in FBs, MΦs, and SMCs in hypoxia. Although *Fbln5* is commonly expressed during development for elastogenesis, it also associates with increased expression after tissue injury to regulate matrix remodeling.^113–115^ We also found that *Chil3+* MΦs, whose gene markers resemble resident, yolk-sac derived, nerve-associated macrophages found in the peri-bronchovascular bundles of lungs that self-renew independent of CCR2+ monocytes,^116,117^ are absent from hypoxic samples, though our study design was insufficiently powered to test for significance of this cell population.

### Altered cell-cell communication in hypoxia: macrophages play a critical role in adventitial ECM remodeling

We also observed increased expression of genes in FBs and SMCs that associated with migration of cell types other than themselves, suggesting intercellular communication amongst resident cells within the wall, a phenomenon that we previously observed.^21^ For example, SMC genes associated with endothelial cell migration (GO:0043536), SMC migration (GO:0014910), and FB migration (GO:0010761). Based on the common changes observed in gene expression and processes associated with signal transduction, we hypothesized that altered intercellular interactions may regulate FB activity and play a key role in an attempt to promote homeostatic remodeling of the adventitia of hypoxic PPAs. To investigate alterations in intercellular communication within the adventitial environment, we performed intercellular communication analyses using NICHES. We assembled a list of genes that included increased mesenchymal and myeloid cells, noting that MΦs are found in the adventitia, and sought to identify signaling mechanisms associated with hypoxia (362 ligand-receptor pairs). This list was used to calculate a “hypoxia signaling score” for each sending-receiving cell pair in NICHES. As expected, hypoxia signaling increased in the hypoxic condition, with the strongest increase occurring SMCs and FBs (*p*<0.05, Figure 5A).

**Figure 5.**
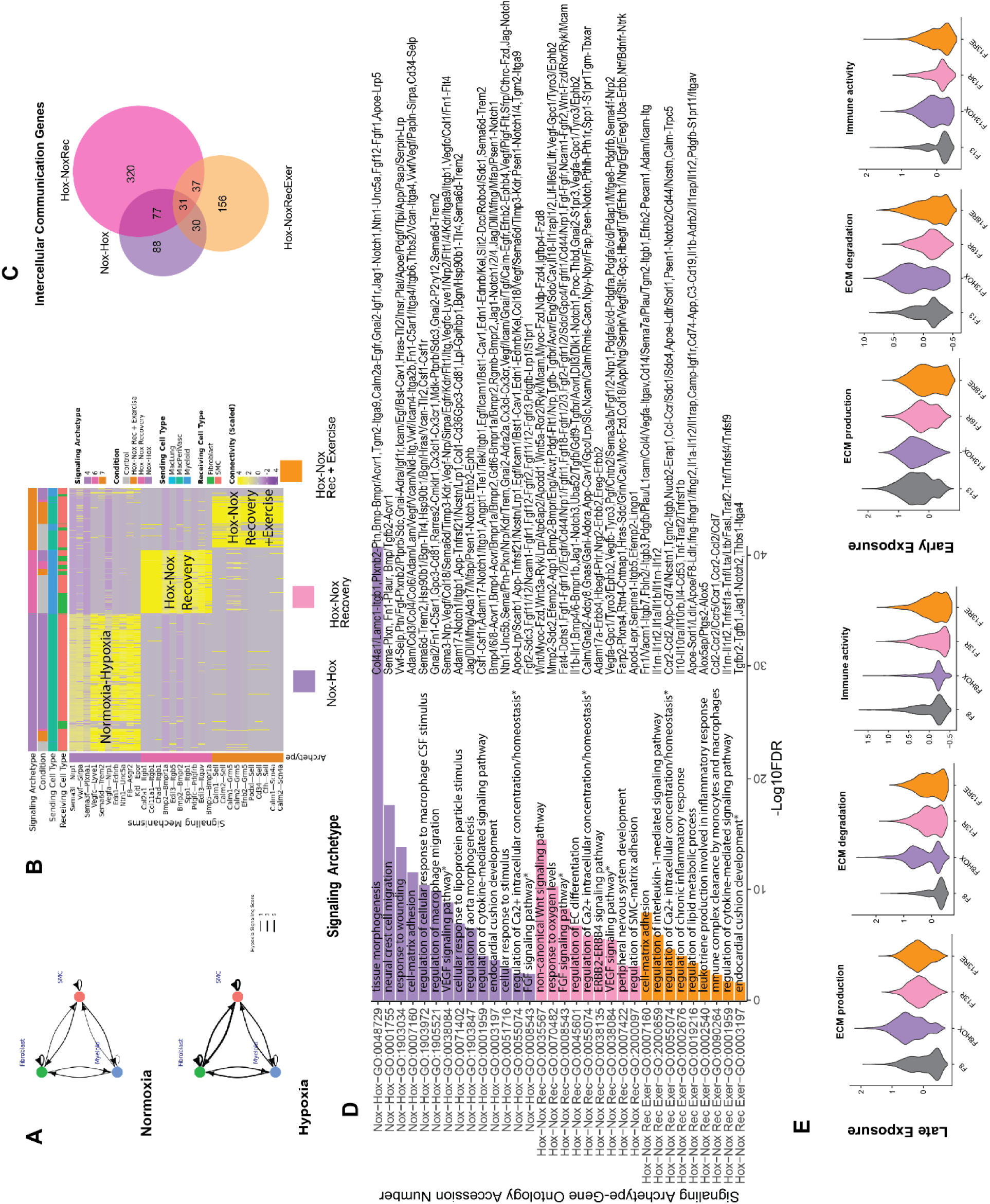
Visualization of increased signaling between smooth muscle cells (SMCs), fibroblasts (FBs), and myeloid cells in hypoxia based on NICHES using scRNA-seq data (panel A). Line thickness reflects the NICHES signaling score between sender-receiver pairs. A heatmap of the top ten genes for sender-receiver pairs for each comparison of conditions (panel B, normoxic compared with hypoxic signaling, normoxic recovery compared with hypoxia (HOX-NOX Recovery), and normoxic recovery with voluntary exercise compared with hypoxia (HOX-NOX Recovery + Exercise). Gene enrichment analysis of signaling genes from NICHES are shown for the different comparisons (panel C). Biological processes for normoxic signaling compared with hypoxic signaling is similar to our differential gene expression for normoxia compared with hypoxia in Figure 4. Comparison of the number of sender-receiver genes for each condition visualized with a Venn diagram demonstrate similarities and differences in the quantities of signaling genes in each condition (panel D). In late exposure, differentially expressed genes associated with ECM production and degradation appeared to increase in chronic hypoxia, a period with which we found re-orientation but not excessive deposition of adventitia collagen fibers and a return to pre-hypoxic baselines (panel E). A similar though less distinct trend was observed in early exposure data. There was a decrease in immune-related gene expression in the late exposure and an increased in early exposure (panel E).

To further characterize changes in cell signaling interactions, we adopted an unbiased clustering approach (see Methods). The top ten enriched sending-receiving pairs of genes for each signaling archetype in these conditions are shown in the heatmap of Figure 5B, with a complete list of significantly enriched sending-receiving pairs of genes provided in Supplemental Table S11. We found signs of enriched signaling among perivascular MΦs, FBs, and SMCs in hypoxia (cluster 4, Nox-Hox, *p*<0.001; Figure 5B; connectivity embedding in Supplemental Figure S11). Signaling interactions characterizing this signaling archetype included *Vwf-Sirpa*, *Vegfc-Lyve1*, and *Nnt1-Unc5a* amongst other sender-receiving pairs. The genes in this group tended to reflect biologic processes that associate with vascular remodeling and angiogenesis similar to results found using a generalized linear mixed effects model to identify gene expression that correlates with hypoxia.

We also found independent signaling archetypes between perivascular MΦs and lung FBs or SMCs enriched during the subsequent recovery from hypoxia (cluster 6, Hox-NoxRec, *p*<0.001) as well as recovery that included voluntary exercise (cluster 7, Hox-NoxRecExer, *p*<0.001). Unique signaling genes enriched in the normoxic recovery from hypoxia (Hox-NoxRec) include: *Bmp2-Bmpr1a, Bmp2-Bmpr2, Chad-Itgb1, Pdgfc-Pdgfrb,* and *Spp1-Itgb1*, which associate with vascular and perivascular remodeling.^118–120^ Unique genes enriched in the normoxic recovery with voluntary exercise (Hox-NoxRecExer) include: *Calm1-Sell*, *Calm1-Grm5*, *Podxl-Sell*, *Cd34-Sell* and *Calm1-Scn4a*. These pairs associate with immune signaling regulation,^121^ hematopoietic cell involvement of vascular development,^122^ and the pleotropic effects of bone morphogenic protein on vascular wall and ECM remodeling.^123^

The majority of genes involved in intercellular communications among MΦ-FB-SMC differed by condition (Figure 5C, Supplementary Table S12). Clearly, cellular-molecular responses during re-exposure to normal levels of oxygen following chronic hypoxia are not simply the reverse of those when mice are initially exposed to hypoxia from normoxic conditions. When comparing trends in biological processes involving the deposition and degradation of ECM and immune system activity that may be involved in regulating it, we found:

- Late exposure trends: during normoxic recovery without exercise, there was an increase in genes driving ECM deposition and degradation that approached pre-hypoxic levels. Immune activity was decreased during late exposures to hypoxia. See Figure 5E.
- Early exposure differed from late exposure in that baseline ECM production appeared to be greater during the development period and immune activity appeared to increase rather than decrease due to chronic hypoxia and during normoxic recovery when mice were allowed to exercise voluntarily. See Figure 5E.

### Hypoxic remodeling is only partially reversible after exposure to chronic hypoxia

We next addressed the clinically relevant question: Is maladaptive remodeling of the PPA and associated cardiac and pulmonary dysfunction reversible? We thus returned additional cohorts of juvenile (hypoxic from 3-8 weeks of age) and adult (hypoxic from 8-13 weeks of age) mice to normoxic conditions for 5 weeks either without or with concurrent voluntary exercise using an in-cage running wheel (recall Supplemental Figure S1). We first assessed overall physiologic consequences of hypoxia and possible normoxic recovery on cardiopulmonary function by measuring exercise tolerance (Figure 6A). There were no significant differences in the ability of the mice to handle forced exercise when exposed to hypoxia early (onset at 3 weeks) or later (onset at 8 weeks) compared to their normoxic, age-matched controls in terms of maximal running speed. Yet, when assessed in terms of daily distances run voluntarily during the post-hypoxic recovery period, still-developing juvenile mice had a significantly greater ability or desire to exercise (*p*<0.01), whereas fully developed adult mice exhibited reduced exercise (*p*<0.01). This finding suggests greater resiliency of the juvenile mice.

**Figure 6.**
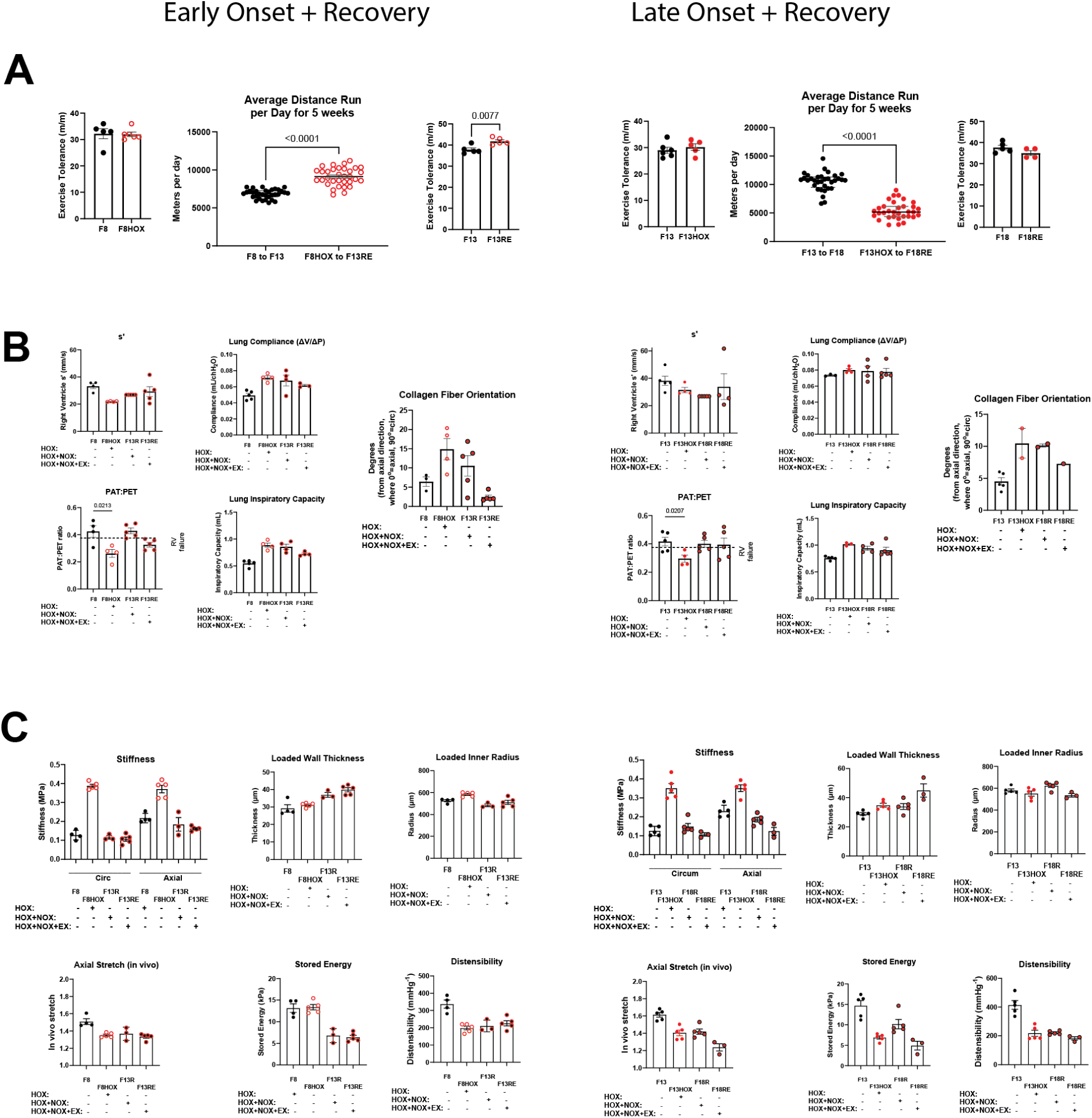
Overall physiological metrics (exercise) for possible normoxic recovery following 5 weeks of hypoxia in adult mice (panel A) as well as measures of cardiac and lung (panel B) and proximal pulmonary artery (panel C) function. Notice a partial recovery of some pre-hypoxic values, but persistent maladaptive changes in many others.

Nevertheless, there was only a partial recovery of cardiac function, measured in terms of s’ and PAT:PET, in both the early and later onset cohorts, with normoxic exercise helping to improve s’ more in the later onset group while blunting recovery of PAT:PET in the early onset group (Figure 6B). There was less recovery in lung function, measured in terms of lung compliance and inspiratory capacity, again independent of age of onset (Figure 6B). The inner radius of the PPA as well as its biaxial wall stress and intrinsic material stiffness (circumferential and axial) returned toward normal values following normoxic recovery (Figures 6C and Supplemental Table S13), augmented slightly by exercise in the later onset group. It is important to note that circumferential stress and material stiffness are typically highly mechano-regulated quantities, at least in adults.^124^ Nevertheless, many geometric and mechanical metrics did not return to normal within the 5-week normoxic recovery periods. The *in vivo* value of axial stretch, the elastic energy storage capability, and overall distensibility remained significantly lower in both the early and later onset cohorts, with voluntary exercise not improving the potential recovery. Importantly, energy storage and distensibility represent the primary functional characteristics (resilience and compliance) of conduit arteries, thus their lack of recovery indicates persistent maladaptive or dysfunctional remodeling. Additional mechanical properties and material parameters are included in Supplemental Tables S13 and S14.

### Interventions (exercise, mTOR inhibition, and metformin) to prevent maladaptive effects during hypoxia have limited effects

Chronic hypoxia associated with ECM reorganization and loss of contractility of SMCs involving Akt and mTOR signaling (WP2841 and WP3855), pathways that others have found to associate with glycolytic metabolism.^125,126^ Related to this, inhibition of Akt/mTOR protects against maladaptive remodeling of the distal pulmonary vasculature and RV in preclinical models of hypoxia-induced pulmonary hypertension by preserving SMC contractility, reducing SMC de-differentiation, and reducing FB proliferation.^127–131^ Based on the common nutrient-sensing molecular pathways associated with genes expressed by the SMCs, FBs and MΦs, particularly insulin signaling (WP65), we hypothesized that this may be a potential target to reduce or prevent maladaptive remodeling in the PPA during chronic hypoxia – we thus treated mice with rapamycin, a mTORC1 inhibitor.^130–132^ In addition, recalling possible changes in metabolism, we also treated mice with metformin, a common drug used in humans with pre-diabetes to regulate insulin signaling. Metformin targets cellular metabolism via AMPK-dependent and -independent pathways and has been shown to protect against distal pulmonary artery remodeling during chronic hypoxia.^130,133,134^ Therefore, we hypothesized that treatment with metformin, or similarly mTOR-targeting rapamycin, could provide some protection against maladaptive remodeling of the PPA in chronic hypoxia by preserving the contractile SMC phenotype and reducing the extracellular matrix remodeling that leads to stiffening. We found that rapamycin partially preserved contractility of SMCs during late onset hypoxia (*p*=0.03, 21.2 ± 3.57 % versus 14.0 ± 4.32 % vasoconstriction in response to high KCl for F13HOXrapa versus F13HOX) although overall SMC contractility remained significantly impaired compared to normoxic controls (*p*=0.01, 31.2 ± 5.04 versus 14.0 ± 4.32 % vasoconstriction for F13 versus F13HOXrapa; Figure 7A, Supplementary Tables S15 and S16). Conversely, SMC contractility in response to phenylephrine was significantly worse (*p*=0.0007, 15.4 ± 2.78 % versus 28 ± 3.65 % vasoconstriction for F13HOXrapa versus F13HOX; Figure 7A). The PPA of mice treated with rapamycin during chronic hypoxia had significantly lower arterial wall circumferential stress and stiffness (usually highly mechano-regulated) than mice exposed to chronic hypoxia without rapamycin treatment (*p*=0.02; Figure 7B, Supplementary Tables S15 and S16), but these values remained significantly higher than normoxic controls (*p*=0.002). Importantly, rapamycin similarly did not preserve the two primary passive functional metrics; the difference between elastic energy storage and distensibility of the PPA was lower (*p*=0.08 and *p*=0.01, respectively) in late onset hypoxic mice whether treated with rapamycin (F13HOXrapa) or not (F13HOX) – Figure 7C,D. Results were generally similar for metformin, whether delivered via the chow (F13HOXmetC) or drinking water (F13HOXmetW); stiffness was significantly less than F13HOX but remained significantly higher than F13 normoxic controls (*p*<0.01) in most cases. Again, see Supplementary Tables S15 and S16.

**Figure 7.**
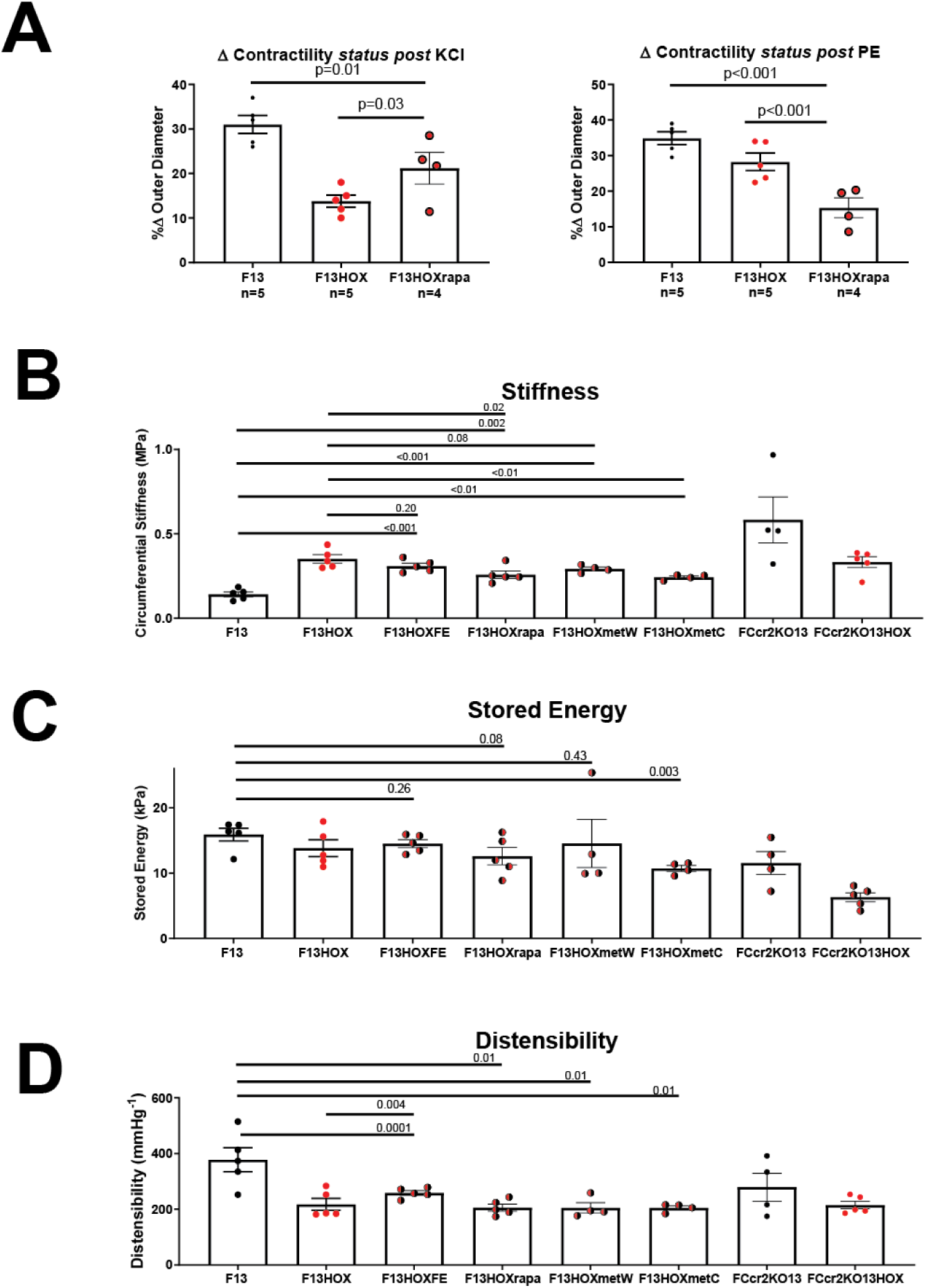
Comparisons of proximal pulmonary artery function across multiple possible interventions focusing on SMC contractility (panel A) as well as circumferential material stiffness (panel B), elastic energy storage (panel C), and distensibility (panel D).

We hypothesized further that the critical roles played by MΦs in hypoxic PPA remodeling resulted in part via communication with FBs and SMCs, similar to findings by others in distal vessels within the lungs.^135^ It was unclear what proportions of the MΦs were resident within the arterial wall or recruited during hypoxic exposure. CCR2+ macrophages have been associated with normoxic homeostasis in adult mice whereas CCR2 deficiency has been associated with increased pulmonary arterial pressures in normoxic conditions.^136,137^ To investigate possible roles of CCR+ recruited MΦs, we quantified mechanical properties of the PPAs in 13-week-old adult *Ccr2^-/-^* mice without or with 5 weeks of prior hypoxic exposure. The PPAs of normoxic *Ccr2^-/-^*mice were significantly stiffer than their age-matched normoxic WT counterparts (*p*< 0.01), resulting in increased PWV (*p*<0.01, Figure 7B and Supplementary Tables S15 and S16). Indeed, the material stiffness of the normoxic *Ccr2^-/-^* arteries more closely resembled those of the hypoxic arteries from WT mice, consistent with absence of CCR2+ macrophages resulting in pressure-induced PPA remodeling independent of hypoxia (Figure 7; Supplementary Table S15 and S16). Exposing adult *Ccr2^-/-^* mice to chronic hypoxic conditions had little effect on further maladaptive remodeling. Recruited macrophages thus appear to be necessary to establish and maintain biomechanical homeostasis in the PPA during normoxic conditions. Lack of these recruited MΦs may reset the homeostatic setpoint of the PPA, which is then preserved during chronic hypoxia.

Finally, based on the alterations of immunological signaling with adjuvant voluntary exercise during normoxic recovery, we hypothesized that exercise may attenuate maladaptive effects of chronic hypoxia. Based on our anecdotal observations that mice tend to huddle during chronic hypoxia and be less mobile, we imposed forced exercise. We found that forced exercise of 10 m/min (∼75% VO max for mice in our study at this age) for three sessions per week, one hour for each session during chronic hypoxia significantly improved distensibility (*p*=0.004, Distensibility = 258.43 ± 8.16 mmHg^-1^ versus 217.57 ± 21.24 mmHg^-1^ for F13HOXFE versus F13HOX, respectively), which nevertheless remained significantly lower than normoxic controls (*p*=0.0001, Distensibility = 387.44 ± 36.69 versus 258.43 ± 8.16 mmHg^-1^ for F13 versus F13HOXFE, respectively). Forced exercise preserved elastic energy storage (*p*=0.26, F13 versus F13HOXFE) but did not significantly reduce wall stiffness of the PPA (*p*=0.20, F13HOX versus F13HOXFE). See, too, Figure 7 and Supplementary Tables S15 and S16.

## Discussion

Numerous conditions and diseases lead to hypoxemia in neonatal and early postnatal periods as well as in adulthood. We quantified hypoxia-induced cardiopulmonary remodeling across these three age groups by exposing female WT mice to 2.5 to 5 weeks of 10% FiO_2_ and measuring functional, structural, and transcriptional changes. Adverse remodeling of the proximal pulmonary arteries (PPAs) appeared to precede impairment of RV and lung function. RV pressure and highly mechano-regulated metrics of PPA homeostasis, such as circumferential wall stress and material stiffness, were largely restored following normoxic recovery, but measures of PPA function such as vasoactive capability, passive elastic energy storage, and distensibility yet remained compromised. This persistent maladaptation is important despite the partial homeostatic recovery, reminding us that single metrics alone need not provide a complete picture of adaptive versus maladaptive changes that can occur simultaneously.

Although data from mice allowed to exercise during normoxic recovery suggested greater resiliency in juveniles compared to adults, the maladaptive changes in PPA function (especially reduced distensibility and reduced ability to store elastic energy during systolic distension to be used during diastole to augment flow) largely emerged and persisted independent of age. Indeed, neither concurrent treatment with Akt/mTOR inhibitors in adult mice nor germ-line depletion of CCR2+ macrophages could preserve these metrics of PPA function at pre-hypoxic levels. Given the critical roles played by SMC contraction and wall elasticity in healthy PPAs, these persistent losses in function represent a critical concern and unmet clinical need.

Collecting data following 2.5 and 5 weeks of hypoxia revealed early changes in RV pressure, hematocrit, and importantly PPA distensibility, which increased PWV. It is thought that increased PWV causes both earlier reflections of the pulse wave (retrograde), which increase afterload on the heart, and a deeper penetration of pulse waves into the distal vasculature (antegrade), which can damage these vessels and cause end organ dysfunction.^138^ These phenomena have been associated with changes in cardiac and pulmonary function,^139–143^ including our associations of PPA stiffening with impaired RV and lung function in natural aging^144^ as well as in chronic obstructive pulmonary disease and pulmonary hypertension.^145,146^ Herein, echocardiography and pulmonary function testing confirmed a hypoxia-induced reduction in multiple metrics of both RV (s’, TAPSE) and lung (diffusion capacity, compliance) function, the latter consistent with the observed microstructural damage.^145,147^ Our direct correlations between PPA stiffening and structural and functional impairment of the lungs appeared to be consistent with correlations between decreased extensibility of the main pulmonary artery and severity of pulmonary arterial hypertension and impaired RV function in patients with increased RV afterload.^24,138,142,148–152^

Notwithstanding an extensive literature on effects of chronic hypoxia on the cardiopulmonary system, differences in animal models and methods of study preclude many direct comparisons. Nevertheless, Liu and colleagues reported detailed temporal data on pulmonary artery remodeling across seven regions in 3-month-old male rats exposed to 10% FiO_2_ for 10 or 30 days.^14,22,153^ They found a rapid rise of pulmonary blood pressure within hours that reached a near steady state by 2 days that increased only slightly thereafter through 30 days of hypoxia (i.e., *P* = *P*_*o*_(1 − *be*^−*kt*^), where *P*_*o*_ is the original pressure and *b* and *k* are parameters), with no significant change in systemic pressure but an initial decrease in body mass over the first 10 days that was followed by a slight recovery through 30 days. Focusing on the PPA, they also provided time-courses of adventitial and medial thickening, which similarly increased rapidly over the first 10 days (i.e., *h* = *h*_*o*_(1 − *be*^−*kt*^) where *h*_*o*_ is the original thickness and *b* and *k* are parameters), with adventitial thickening outpacing medial thickening. Inner radius changed modestly, and mean circumferential wall stress (*σ*_*θ*_ = *Pa*/*h*, where *P* is pressure, *h* is thickness, and *a* is inner radius) increased ∼1.35 fold over the first 12 hours, then decreased monotonically back toward the original value (∼22 kPa) within ∼ 4-5 days and remained so through 10 days, thus suggesting a stress-based homeostatic response despite the continued hypoxia. This concept was supported by a companion study comparing normoxic recovery of PPA geometry and wall stress after 12 hours vs. 10 days of hypoxia, with wall stress returning toward normal in both cases though via different time-courses. Our recent data in 8-week-old male mice yet moderates the suggestion of a mechanical homeostatic response.^154^ Although circumferential wall stress first increased and then returned toward normal values during a 3-week exposure to 10% FiO_2_, circumferential wall stiffness exhibited only a partial restoration while axial stiffness remained higher, both elastic energy storage and vasoactive capacity remained lower, thus suggesting an overall compromised biomechanical functionality. Our present biaxial data are consistent with, but extend, prior findings of time-course changes to neonatal, juvenile, and adult female mice.

Also decades ago,^155^ it was shown that normoxic recovery following 10 days of 10% FiO_2_ in 6-week-old male rats showed monotonic returns of pulmonary pressure to normal by 14 days with a similar return of medial thickness of the PPA toward normal, which associated with a comparable return of mural protein content. Importantly, gelatinase activity increased and collagenase activity decreased slightly during the 10 days of hypoxia, but both increased dramatically (∼5-fold) early in normoxic recovery (day 3) before decreasing toward baseline values over the 14 days of recovery. The increase in collagenase activity correlated with the partial reversal of the collagen accumulation during the hypoxic period, which had manifested via marked (up to 8-fold) increases in synthesis rates.^156^ A consistent finding in their 12-week-old mice that were exposed to 10% FiO_2_ for 10 days and then normoxic recovery for 32 days was an increased circumferential stiffness that associated with increased collagen accumulation, which resolved in WT mice upon restoration of 20% FiO_2_ but not in mice having a mutation that slows collagen degradation.^157^ A companion study using β-aminopropionitrile (BAPN) to block cross-linking of newly synthesized collagen emphasized that increased stiffening requires appropriate cross-linking.^158^ Finally, this group also suggested that the normoxic recovery following hypoxic remodeling is partial, not full, because of persistent alterations to the collagen fibers,^159^ which is consistent with recent findings in mice on recovery following systemic hypertension.^160^ Although we did not measure turnover rates of collagen, our results are again consistent with these prior studies while emphasizing further that remodeling/reorientation of adventitial collagen persists despite normoxic recovery, suggesting some permanent alterations to the condition of the collagen even if not to the quantity of collagen. A computational modeling study of long-term (up to 6 months) aortic recovery from a brief period of angiotensin II-induced hypertension similarly suggested that persistent changes in collagen can prevent full recovery of pre-hypertensive properties^161^.

Clearly, focusing on cell-mediated changes in PPA structure and function may hold clues to targeted early interventions, which must build on current understanding. Consistent with the observed persistent remodeling of adventitial collagen, key biological processes emerging from our scRNA-seq data for the PPAs supported the centrality of altered collagen fibril organization, ECM organization, and cell-matrix adhesion. The transcriptomics suggested further that complex intercellular communication amongst SMCs, FBs, and MΦs likely plays a critical role in this hypoxia-induced remodeling. Many prior reports of acute transcriptomic responses to hypoxia focused on the important role of the gene encoding hypoxia-inducible factor-1 (*HIF1A* in humans, *Hif1a* in mice, which encode the α-subunit of the heterodimeric transcription factor HIF-1), which plays an important role in cellular responses to hypoxia,^8,10,162^ adaptations in cellular metabolism,^162^ regulation of gene expression,^163^ and cardiovascular remodeling.^164^ HIF-1 is ubiquinated (through actions including EGLN and von-Hipplel-Lindau oxygen-dependent enzymes) in normoxic physiologic conditions; in hypoxia, it avoids degradation and translocates to the nucleus.^165^ *Hif1a* is also known to increase the expression of genes that encode VEGF and VEGF receptors KDR/Flk and Flt.^166^ VEGF and its receptors have been associated with SMC migration^167^ and angiogenesis,^168^ similar to findings from our study. Prolonged hypoxia causes SMC hyperplasia and hypertrophy in the thickened walls of distal pulmonary vessels, effects mediated by *Hif1a*.^163,164,169^ There is also evidence that HIF-1 plays a role in regulating intercellular signaling between SMCs and other resident cells within the pulmonary artery wall through HIF-1 dependent production of signaling molecules such as endothelin-1 (ET-1).^164,170^ Meta-analysis of transcriptomic responses to early hypoxic responses (< 48 hours) of cells in vitro revealed that acute hypoxia associates with increased differential expression of genes (e.g., *EGLN1, LOXL2, TGFBx, DCN, IL6*) rather than decreased (repression).^165^ In comparison to many of these studies by others, we focused on changes after longer periods of hypoxia, 2.5 to 5 weeks at 10% FiO_2_, and found that *Hif1a* expression associated with SMC response to VEGF stimulation. In FBs *Hif1a* also associated with angiogenesis as well as remodeling of the extracellular matrix of the PPA, specifically collagen and glycoprotein remodeling.

Distal pulmonary arterial responses in mice exposed to chronic hypoxia (4 to 28 days) include mononuclear cell (inflammatory) infiltration, primarily during early phases of hypoxia, and mural remodeling in later phases, with decreases in IL-10, increases in IL-12, as well as initial spikes with eventual declines in IL-1β and IL-33.^18^ Also noted were increases in the number of CD68+ macrophages, with an acute inflammatory response on day 4 of hypoxic exposure that progressively impacts the distal pulmonary arteries due to increased expression of iNOS (M1 macrophage marker) and decrease of Arg-1 (M2 macrophage marker, though initially increased during days 4-14).^18^ Chronic hypoxia is also known to alter the microenvironment of cells within the walls of pulmonary arteries, affecting cellular adhesion and differentiation, cytokines (TGFβ), growth factors (VEGF-A), fibrosis (TIMP-1, MMP-2), and chemokines for monocyte/macrophage recruitment (CCL2/MCP-1, CCR2, IL-6, IL-6R, and TLR-4).^64^ Furthermore, others have speculated the “elimination or inactivation” of subsets of macrophages may attenuate maladaptive changes to pulmonary vessels in the setting of hypoxia or disease.^16^ We found that hypoxia associated with the loss of YM1+ (*Chil3+)* macrophages, though our study was underpowered to determine statistical significance of this loss. We also did not identify a significant association of M1 or M2 macrophage polarization markers with hypoxic exposure; rather, we found significant differences in overall immune signaling activity. Clearly, the roles of MΦs in signaling SMC and FBs within PPAs is complex.

CCL2/MCP-1 (ligand) and CCR2 (receptor) are key markers of MΦ involvement in hypoxic remodeling of the pulmonary vasculature in mice.^136,137^ Moreover, it has long been suspected that early and continued leukocyte recruitment, including CCR2+ monocytes, to the pulmonary vasculature is a key step to adventitial remodeling of pulmonary vessels.^171,172^ Our study in *Ccr2^-/-^*mice extends prior findings to include the PPAs, which under normoxic conditions were similar to WT PPAs following hypoxic exposure. Interestingly, exposing the adult *Ccr2^-/-^* mice to chronic hypoxic conditions had little to no effect on further remodeling. These results suggest that CCR2*+* macrophages are necessary for normoxic development and homeostatic remodeling of PPAs, but their absence prevents further maladaptive remodeling in response to chronic hypoxia. Moreover, that a PPA stiffening phenotype associates with the absence of CCR2+ macrophages in the *Ccr2^-/-^* mice in normoxic conditions and loss of YM1+ expressing macrophages, which are yolk-sac derived and capable of self-renewal,^116^ in our hypoxic WT mice suggests that MΦs play a critical role in regulating resident cell activity within the PPA, a finding similar to that which is described in the distal arteries.^16,18^ It yet remains unclear if this shift of macrophage phenotype is due to a lack of recruitment of bone marrow derived myeloid cells, a transient in resident myeloid cells in response to inflammatory cytokines, or both. A larger sample size focusing on MΦ populations in the PPA is required to better address these questions. Regardless, MΦs appear to play a critical role in promoting homeostasis of the PPA in part by orchestrating intercellular communications amongst the resident cells.

That hypoxia-induced changes in PPA structure and function were independent of age of hypoxic onset was surprising since it is often thought that developing blood vessels often display “heightened or unique responses to injurious stimuli”.^173^ It may be that any transition from normoxia to chronic hypoxia initiates common changes in intercellular communication amongst resident arterial cells, regardless of their current phenotypic status, that result in maladaptive adventitial collagen remodeling at any age.^91^ Intercellular communications amongst resident cells of the hypoxic PPA may thus play a more critical role in remodeling of the adventitia than external directors of development.^174^

On restoration of normoxic conditions, we found a return of many hypoxia-related genes to near normal pre-hypoxic levels of expression, similar to transcriptomic changes described by others including *Cxcl12*, *Ccl2, Tgfbx,* and *Icam1*.^64^ It may be that resolution of the complex, inflammatory microenvironment associated with macrophage-mediated maladaptive remodeling facilitated the return of some geometric features, such as wall thickness and inner radius, and biomechanical metrics, such as wall stress and material stiffness, to pre-hypoxic homeostatic values. Overall recovery was incomplete, however, as functional indices including distensibility and elastic stored energy of the arterial wall—biomechanical properties having meaningful physiologic associations, noting that decreased distensibility associates with impaired hemodynamics and impaired cardiopulmonary function^147^—failed to return to pre-hypoxic values. The persistent impairment of pulmonary arterial function associated with an incomplete return of collagen fibers to their original axial orientation and compromised lung and cardiac mechanics. Although young mice displayed supernormal compensatory cardiopulmonary function in the form of increased exercise tolerance following hypoxia, the sustained maladaptive functional remodeling of the cardiopulmonary system appeared to decrease cardiopulmonary ability in adulthood and may place adults at greater risk of future cardiopulmonary failure if perturbed again by hypoxia, illness, or other stresses. The persistence of maladaptive remodeling due to chronic exposure to hypoxia may also aid in understanding why distensibility of the pulmonary vasculature has been shown to decrease in humans following chronic exposures to hypoxia but not acute exposures.^175^

Unfortunately, neither concurrent treatment with a candidate drug (rapamycin or metformin) nor forced exercise were able to prevent the observed persistent hypoxia-induced losses of function. In particular, given the general belief that exercise provides beneficial effects to the cardiopulmonary system as well as an extensive literature regarding its beneficial remodeling of the aorta in the context of systemic hypertension,^176–184^ we were surprised that adjuvant exercise during recovery from chronic hypoxia in juvenile mice (“early exposure” group) had no effect and only partially beneficial effects in maintaining PPA wall thickness and radius of adult mice (“later exposure” group). Further, adjuvant exercise had mixed effects on recovery of functional indices of the pulmonary vasculature as well as lung and RV functions. Differences in our results and beneficial results in other studies may reflect the context with which exercise was performed. In systemic hypertension, aging, and other pro-inflammatory states, exercise may reduce cytokines such as IL1β, IL6, TNFα as well as reactive oxygen species, which associate with inflammation. In the normoxic, post-hypoxic recovery states of our studies, we found that the expression of these genes returned to pre-hypoxic levels without the need for adjuvant exercise. The small improvements with exercise that we observed in biomechanical properties of adult mice may thus be related more to benefits observed from moderate exercise in aging mice,^185–188^ though the adult mice in our study were too young to be considered aged and the overall functional indices of their pulmonary vasculature suffered sustained impairment. Perhaps the increased cardiac output and increased pulmonary circulatory pressure that occur during exercise contribute to the lack of benefit from exercise during normoxic recovery because they may re-create stress-induced conditions that associate with the hypoxic exposure, thus hindering rather than aiding normoxic recovery. Thus, interventions to reduce intraluminal pressure within the proximal pulmonary arteries may be a potential target for prophylaxis of maladaptive remodeling of large arteries.^189^

Genomic adaptations to prolonged exercise training have suggested links between exercise and altered regulation of immune, metabolic, stress response and mitochondrial biogenesis, which translates to roles of exercise in cardiovascular health, tissue injury, and recovery.^190^ Weeks of exercise in female mice associated with changes in gene networks related to heart disease, including genes *Mmp14* and *Lox*, and enrichment of enzymes in the glycolysis-gluconeogenesis pathway. Further, increased expression of genes associated with the MAPK and TGFβ signaling pathways, which were higher in our hypoxic cohorts of mice, were also found in mice exposed to prolonged exercise.^190^ While exercise for sedentary individuals may be beneficial to cardiopulmonary health, signals increased during exercise may have deleterious effects on cardiopulmonary systems that are under stress. There also appears to be a role of development in this process. C57BL/6J mice do not complete development until 13 weeks of age.^29^ Mice in the early exposure group of hypoxia were thus continuing to develop during normoxic recovery ± exercise; mice in the late hypoxia exposure group had completed development at the same time at which their hypoxic exposure ended. The younger mice showed no beneficial remodeling in response to exercise, whereas the older mice showed some benefit in maintaining geometric values. It is possible that elements of development enable the cardiopulmonary system to remodel completely, whereas exercise in addition to normoxia is necessary in the older group. Regardless, PPA stored energy and distensibility did not return to pre-hypoxic baselines which suggests that exercise is not able to improve recovery of the cardiopulmonary system immediately following hypoxia. Even forced exercise during hypoxic exposure had only modest beneficial effects.

Although we considered many groups across multiple ages, our studies are nevertheless limited. We focused on female mice based on our data that show that mechanical changes of the lungs in female and male mice are largely due to differences in body mass rather than sex^29^ and in part to balance the prior preponderance of studies on male mice; effects of sex should yet be considered further given known transcriptomic effects that are sex-dependent.^190^ Ancillary studies such as our micro-CT imaging would benefit from a dedicated study comprising a larger sample size to evaluate effects of remodeling of the distal arteries and microvasculature of the lung as well as comparing the contributions of vascular resistance of the right and left lungs separately. We focused on a severe (10% FiO_2_) hypoxic insult, but there is a need to consider graded insults (from 10 to 20% FiO_2_). We considered multiple potential interventions, but we did not consider different doses of the drugs, other drug classes, or different exercise regimes. For example, we did not investigate effects of chronic hypoxia on vasodilatation and the potential protective roles that vasodilators such as PDE-5 inhibitors have on pulmonary vascular remodeling during hypoxia.^191–193^ That our interventions – rapamycin, metformin, forced exercise, or depletion of CCR2+ cells – did not prevent adverse hypoxia-induced maladaptations in adult mice may suggest a need for combination therapies, particularly given the multi-scale (from resistance to conduit pulmonary arteries) and multi-organ effects of chronic hypoxia. Given the persistence of many of these changes, it appears however that there is a need to focus on prevention, not treatment, of symptoms. Notwithstanding these and other limitations, much was learned.

## Conclusions

In summary, our working hypothesis to explain changes in the PPA resulting from chronic hypoxic exposure includes a sequence of events that alter the microenvironment of the arterial wall. First, acute hypoxia likely causes distal pulmonary arteries to constrict, an evolved mechanism that reduces blood flow to regions of the lung with decreased oxygen tension to allow a higher proportion of blood to flow through regions of lung that are better ventilated.^13^ This optimizes the ventilation:perfusion ratio and ensures optimal blood oxygenation under normal conditions. In global hypoxia, however, all distal, small pulmonary arteries constrict resulting in increased PVR, which increases proximal blood pressures that initiate remodeling of the PPAs. The structural stiffening of the PPAs, evidenced by decreased distensibility and thus increased PWV, can then propagate the insult to the right heart (increased afterload) and lung (increased pulse pressures in the distal vessels), the latter including capillary damage, small vessel loss, and resulting interstitial damage. Prolonged hypoxia causes SMC dysfunction in the PPA and hyperplasia and hypertrophy in distal pulmonary vessels through *Hif1a* mediated processes and thickens the distal arterial walls.^163,164,169^ Pulmonary blood pressure either increases further or is maintained at the elevated level, with aspects of PPA remodeling seemingly entrenched. This remodeling likely results from changes in increased mechanical stress-driven mechanotransduction and increased HIF-1α signaling within the low-oxygen, pro-inflammatory, elevated wall stress microenvironment. Altered cell-specific responses include SMC shifts from a contractive to synthetic phenotype and intercellular signaling that facilitates recruitment of monocytes. This new environment may evolve into an equilibrium state in which degradation of extant collagen allows deposition of new collagen oriented toward the circumferential direction. A positive feedback loop entails – small vessel damage in the lungs and increased stress on the RV lead to greater pressures within the PPAs that continue to operate in a higher pressure, lower-oxygen, pro-inflammatory environment, and so on. Suggested changes in the pulmonary vasculature due to chronic hypoxia are summarized in Figure 8. Upon the return to normoxia, there is return toward a normal oxygen saturation, lower inflammation, and a decreased wall stress microenvironment in the walls of the PPAs. There may be some return of adventitial collagen toward the axial orientation, though incomplete. Although some geometric and biomechanical metrics such as wall thickness and material stiffness appear to be restored to pre-hypoxic homeostatic values, multiple functional indices such as distensibility and stored energy do not return to pre-hypoxic values, thus, representing a persistent functional impairment. This likely poses prolonged risk to the RV and lungs whereby additional insults (e.g., acute hypoxemic events, increases in pulmonary arterial pressure) are likely to hasten further RV and lung failure.

**Figure 8.**
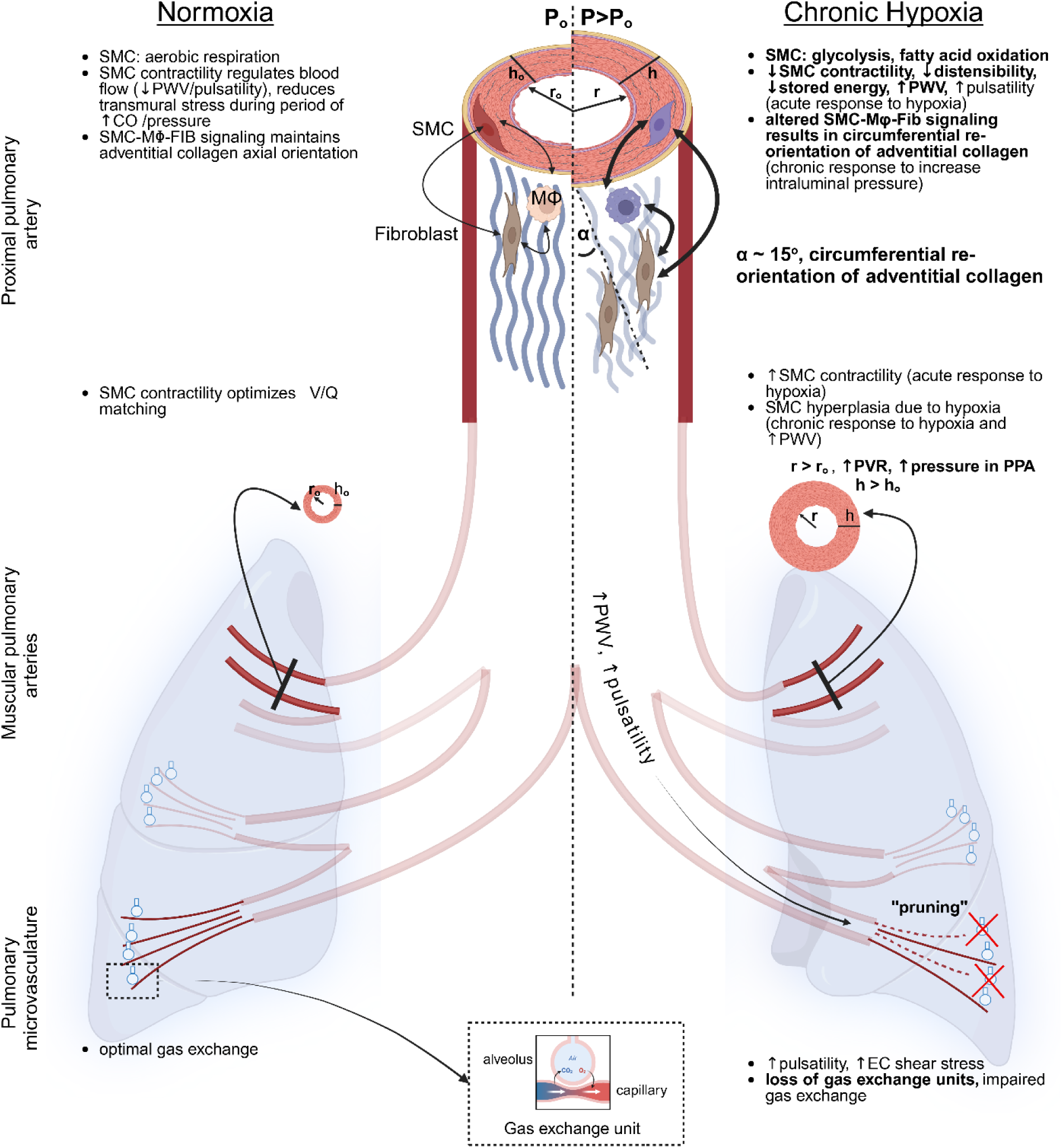
Schema of the proposed multi-scale, multi-organ dysfunction in chronic hypoxia. Highlights of the structure and function of pulmonary arteries in normoxia are on the left; changes in the structure and function of pulmonary arteries due to chronic hypoxia are shown on the right. Changes observed in this study are in bold font.

Remodeling of the pulmonary vasculature due to hypoxia is thus a complex, multi-scale process that involves many of the hallmarks of pulmonary hypertension, including hypoxia signaling, altered metabolism, cellular proliferation, altered immune responses, angiogenesis, ECM remodeling, transcriptional changes, and associated RV failure and pulmonary dysfunction.^194^ Whereas the majority of studies have focused on remodeling of the distal pulmonary vasculature within the lung parenchyma, we demonstrate here that a complete understanding of hypoxia-induced cardiopulmonary dysfunction requires inclusion of the proximal pulmonary artery independent of whether it is causative or a consequence to pathophysiologic remodeling of the rest of the cardiopulmonary system.^195^

## Acknowledgments

This work was supported by grants from Additional Ventures (SVRF – JDH and EPM, AVCC – JDH) as well as the Veteran’s Affairs (EPM) and Pepper Center at Yale University (EPM). NK is supported by R01HL127349, R01HL141852, U01HL145567, UH3TR002445, R21HL161723. XY is supported by R01LM014087 from NLM. We also thank the George M. O’Brien Kidney Center at Yale (P30 DK079310) for blood test measurements.

## Contributions of Authors

AB – biomechanical testing of pulmonary arteries, edited manuscript and figures

PS – single cell RNA sequencing analysis, edited manuscripts and figures

RDM - NICHES intercellular signaling analysis, edited manuscript and figures

FN – single cell RNA sequencing analysis

NG – echocardiography of mice and image analysis

PD – two-photon microscopy imaging analysis

CC – performed two-photon microscopy imaging, performed and supervised two-photon microscopy analysis and edited manuscript

RC – biomechanical measurements of pulmonary arteries

MSR – NICHES intercellular signaling analysis, edited manuscript

JMS – microCT imaging and analysis

ZZ – pulmonary vascular contrast injection, microCT imaging

TB – pulmonary airway mechanical testing of mice

AJ – single cell RNA sequencing preparation NA – exercise testing of mice

NAH – performed single cell RNA sequencing

LD – blood testing of mice

RP – exercise testing of mice, editing of manuscript

TSA, XY – single cell RNA sequencing analysis, edited manuscript

IS – edited manuscript

NK –supervised experiments, edited manuscript

XY – single cell RNA sequencing analysis, edited manuscript

GT-supervised experiments, edited manuscript

JDH – conceived project, supervised experiments, analyzed data, revised figures, wrote manuscript, edited manuscript

EPM – conceived project, imposed hypoxic exposures and exercises, performed biomechanical experiments of pulmonary arteries and lungs, histology, two-photon imaging, single cell RNA sequencing experiments and analysis, supervised other experiments, prepared figures and supplementary materials wrote manuscript, edited manuscript,

Additional Figures and Tables are in Supplementary Materials.

