## Supplemental Materials for "Hypoxia-Induced Cardiopulmonary Remodeling and Recovery: Critical Roles of the Proximal Pulmonary Artery, Macrophages, and Exercise"

1 – Dept of Mechanical Engineering, Iowa State University,
2 - Section of Pulmonary, Critical Care and Sleep Medicine, Dept of Internal Medicine, Yale School of Medicine
3 – Translational Research Imaging Center, Yale School of Medicine
4 – University of California, Santa Barbara
5 –LMGC, Univ. Montpellier, CNRS, Montpellier, France
6 – Dept Biomedical Engineering, Yale University
7 – Department of Anesthesiology, Vascular Biology & Therapeutics, Program in Translational Biomedicine, Yale School of Medicine.
8 – Department of Biomedical Engineering, Carnegie Mellon University
9 – Yale Cardiovascular Research Center, Department of Internal Medicine, Yale School of Medicine, New Haven, CT, USA
10 – Otto Loewi Research Center, Division of Pharmacology, Medical University of Graz, Austria
11 – Service de Pneumologie-Centre de Competence Maladies Pulmonaires Rares – UMR 6030 CNRS- ISTCT, Université de Normandie Caen France, France, Section of Pulmonary, Critical Care and Sleep Medicine, Dept of Internal Medicine, Yale School of Medicine
12 – Department of Cellular and Molecular Physiology, Yale School of Medicine
13– O'Brien Kidney Center Renal Physiology Core, Yale School of Medicine
14 –School of Public Health, Yale University
15 – Dept of Surgery (Cardiothoracic), Yale School of Medicine
16 – Vascular Biology & Therapeutics, Yale School of Medicine
17 – VA Connecticut Healthcare System

**Supplemental Figures**

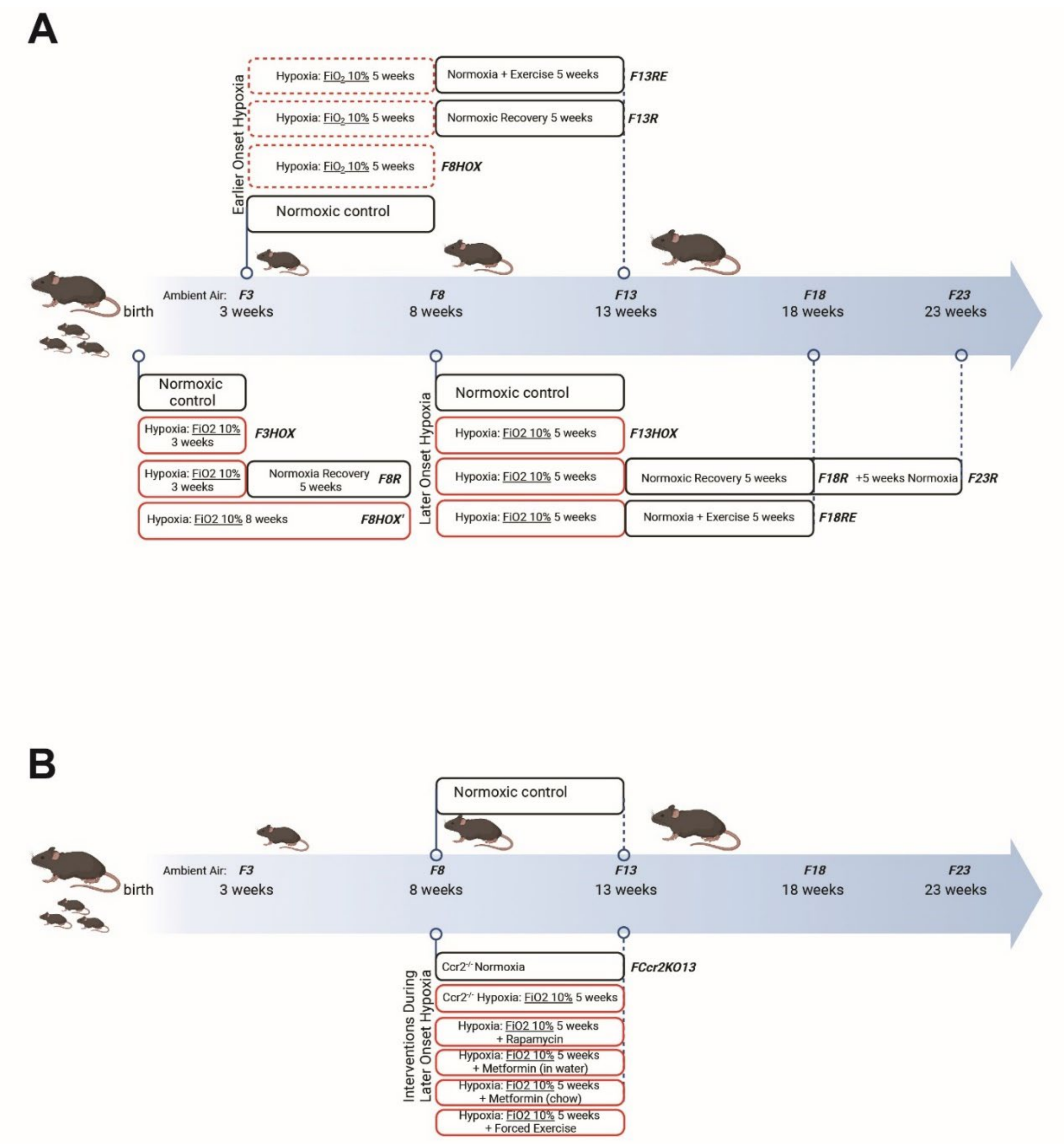


**Figure S1. Experimental Design.** The research design comprised groups of female mice including age-matched normoxic controls, hypoxic, hypoxic with subsequent normoxic recovery without and with voluntary exercise, and hypoxic with concurrent treatment with either different medications or forced exercise. Periods of hypoxia began from birth, 3 weeks or 8 weeks of age and ranged from 2.5 to 5 weeks. At the scheduled endpoints, cardiac and pulmonary function were measured and a battery of tests were performed. F3 = 3-week-old normoxic controls; F8 = 8-week-old normoxic controls; F13 = 13-week-old normoxic controls; etc. F3HOX = 3-week-old mice exposed to hypoxia within 24 hours of birth; F8HOX = 8-week-old mice exposed to 5 weeks of hypoxia starting at 3 weeks of age; F8HOX'= 8-week-old mice exposed to 8 weeks of hypoxia after birth; F13HOX = 13-week-old mice exposed to 5 weeks of hypoxia starting at 8 weeks of age. F8R = 8-week-old mice exposed to hypoxia for 3 weeks after birth then 5 weeks of normoxia; F13R = 13-week-old mice exposed to 5 weeks of hypoxia starting at 3 weeks of age then 5 weeks of normoxia; F18R = 18-week-old mice exposed to 5 weeks of hypoxia starting at 8 weeks of age then 5 weeks of normoxia. F13RE = 13-week-old mice exposed to 5 weeks of hypoxia starting at 3 weeks of age then 5 weeks of normoxia including voluntary exercise; F18RE = 18-week-old mice exposed to 5 weeks of hypoxia starting at 8 weeks of age then 5 weeks of normoxia including voluntary exercise.

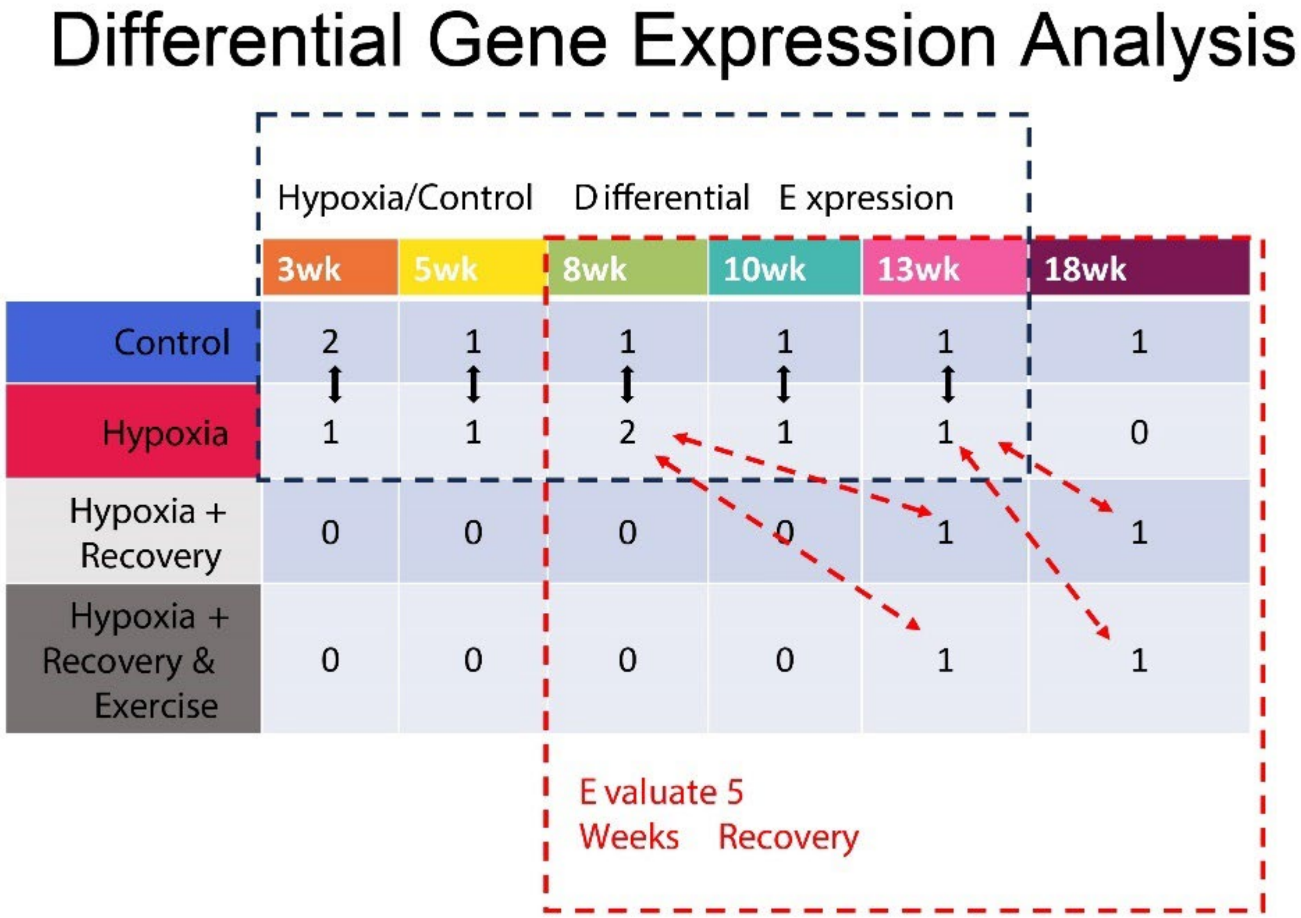


**Figure S2. Single Cell RNA Sequencing Experimental Design**. We performed single cell RNA sequencing on viable cells from non-biomechanically tested main pulmonary arteries of mice at various ages exposed to the various conditions (normoxic, hypoxic, hypoxic with normoxic recovery without or with exercise). We used a generalized linear mixed effects statistical model to find differentially expressed genes (DEGs) due to hypoxia independent of age. This enabled us to isolate effects of hypoxia from developmental changes in mice. X-axis is age; Y-axis is condition. The numbers in the cells indicate number of replicates. Each replicate comprises combined arteries from three mice of that age and condition. The solid black arrows indicate comparisons for hypoxia compared with normoxia, whereas the dashed red arrows indicate comparisons for recovery versus hypoxic condition. The wk = weeks of age at the time of RNA isolation.

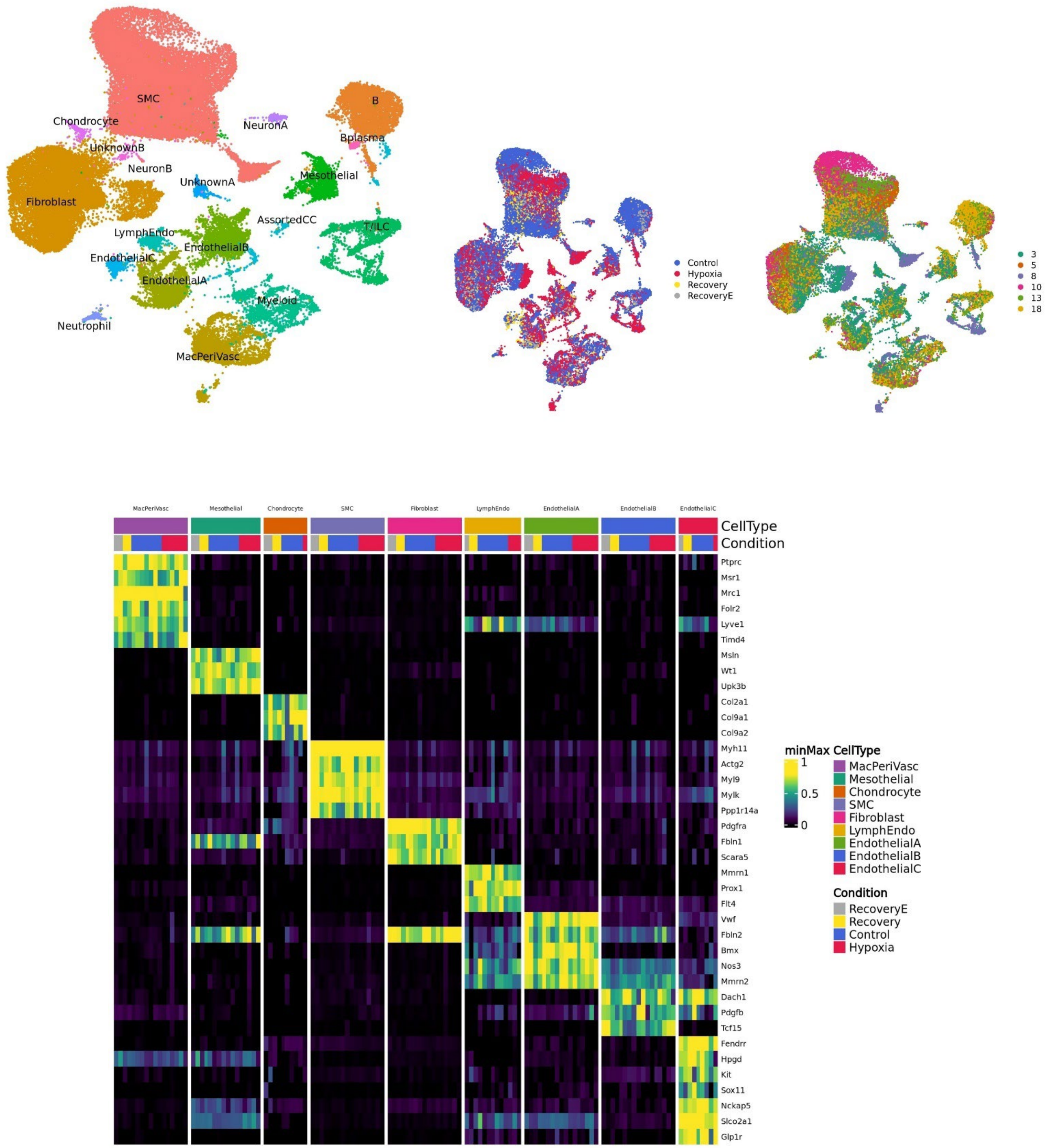


**Figure S3. Single Cell RNA Sequencing Gene Marker List embedding for Cell Type Identification.** Cells were identified using uniform manifold approximation and projection (UMAP) representing various cell types (top left), conditions (top middle), and age in weeks (top right), suggesting that clustering based on cell type rather than cell state. Cell markers to define the cell type annotations are shown in the heatmap (bottom) with emphasis on smooth muscle cells (SMC), fibroblasts, and myeloid cells.

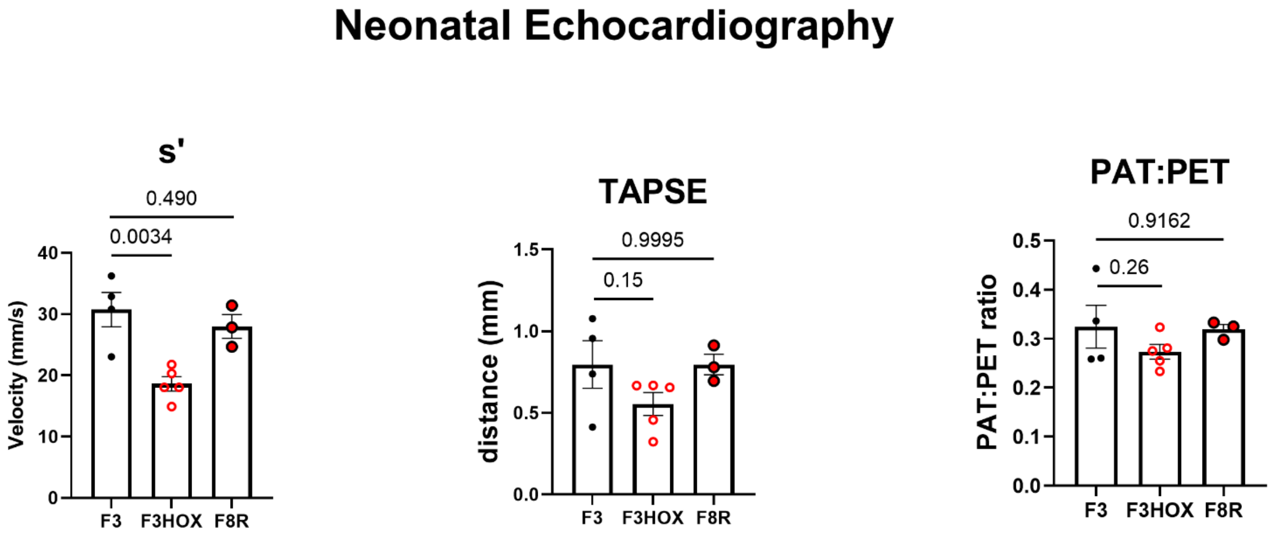


**Figure S4. Neonatal echocardiographic data of right ventricular (RV) function.** Degrees of RV impairment (altered rate of contractility measured by tissue doppler (s'), distance of tricuscpid annular plan systolic excursion (TAPSE), and PAT:PET ratio) of three-week-old mice exposed to hypoxia immediately after birth (F3HOX) were similar to those seen in hypoxic exposures of juvenile (F8HOX) and adolescent mice (F13HOX) compared to age-matched normoxic mice. However, we found a greater degree of recovery in cardiac function in neonatal mice exposed to three weeks of hypoxia followed by five weeks of normoxia (F8R). Younger, developing mice appear to be more capable of greater recovery and thus are more resilient to hypoxic injuries.

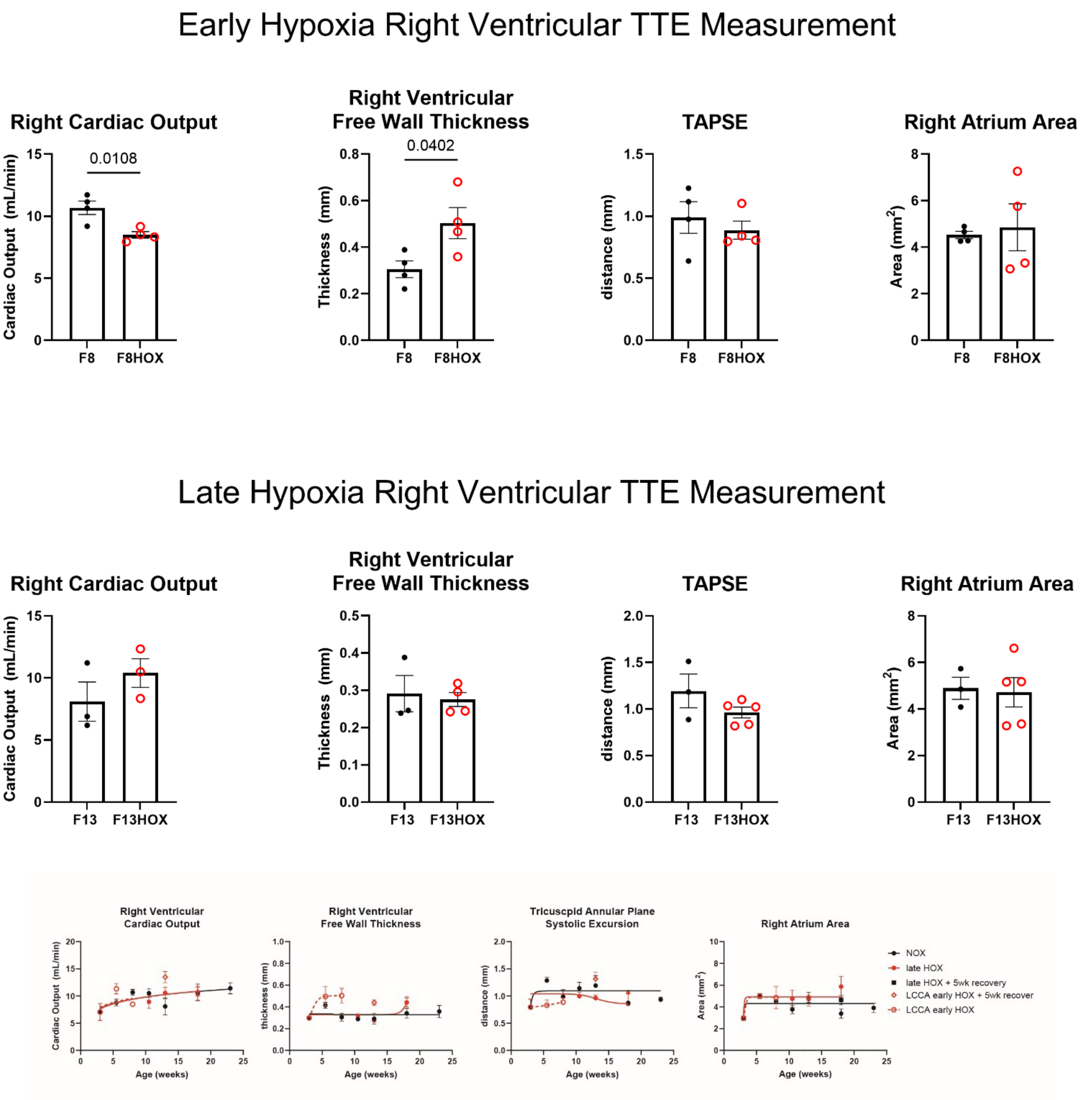


**Figure S5. Echocardiogram Supplemental Data.** Additional comparative and time-course echocardiographic studies for early onset (HOX from 3 to 5.5 to 8 weeks) and later onset (HOX from 8 to 10.5 to 13 weeks of age) for RV function.

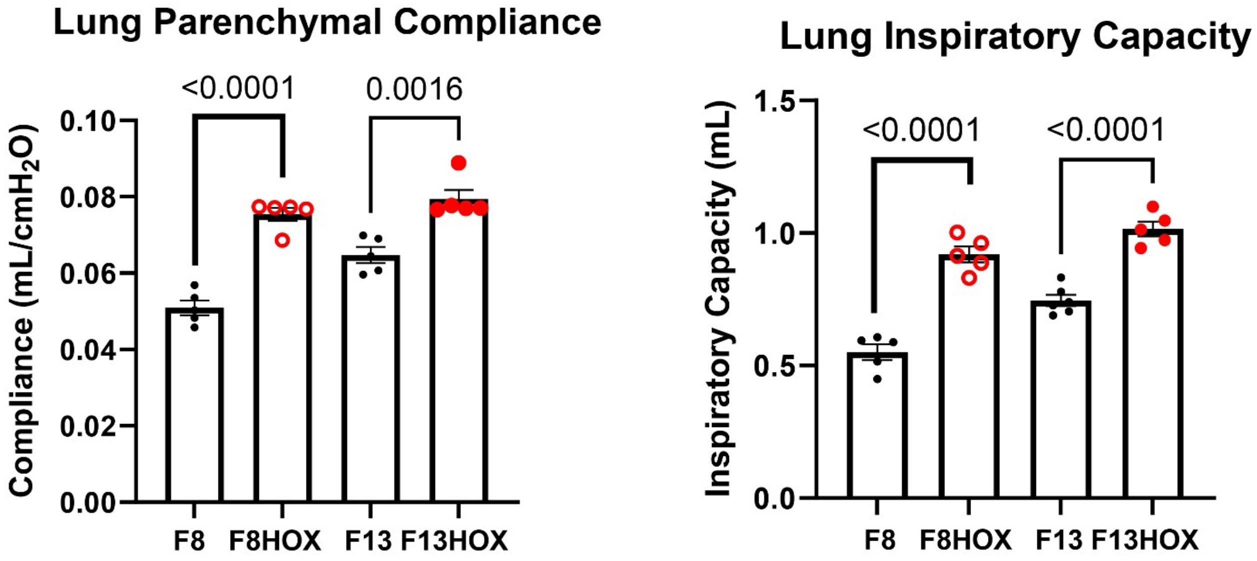


**Figure S6. Supplemental Pulmonary Mechanics Data.** Additional comparative studies for early onset (HOX from 3 to 8 weeks) and later onset (HOX from 8 to 13 weeks of age) for lung function.

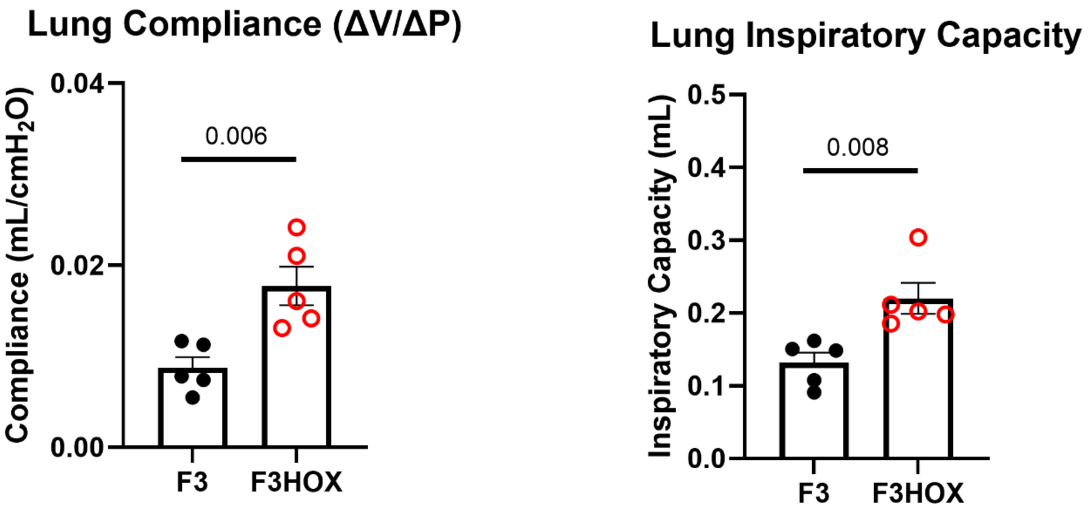


**Figure S7. Neonatal Pulmonary Mechanics Data.** Neonatal mice exposed to 3 weeks of hypoxia from birth (F3HOX) display similar changes in pulmonary mechanics (significantly increased compliance and inspiratory capacity) as we found in juvenile (F8 vs. F8HOX, early exposure) and adolescent adult mice (F13 vs. F13HOX, late exposure).

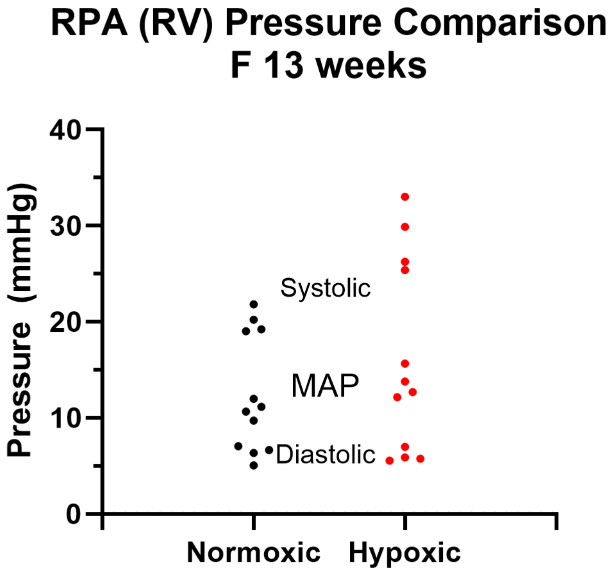


**Figure S8. RV Pressures measured by Millar catheter.** Inset graph: comparison 13-week normoxic control versus 13-week previously exposed to five weeks of hypoxia. Black line = trend of normoxic controls. Gray line = trend of continuous hypoxic exposure.

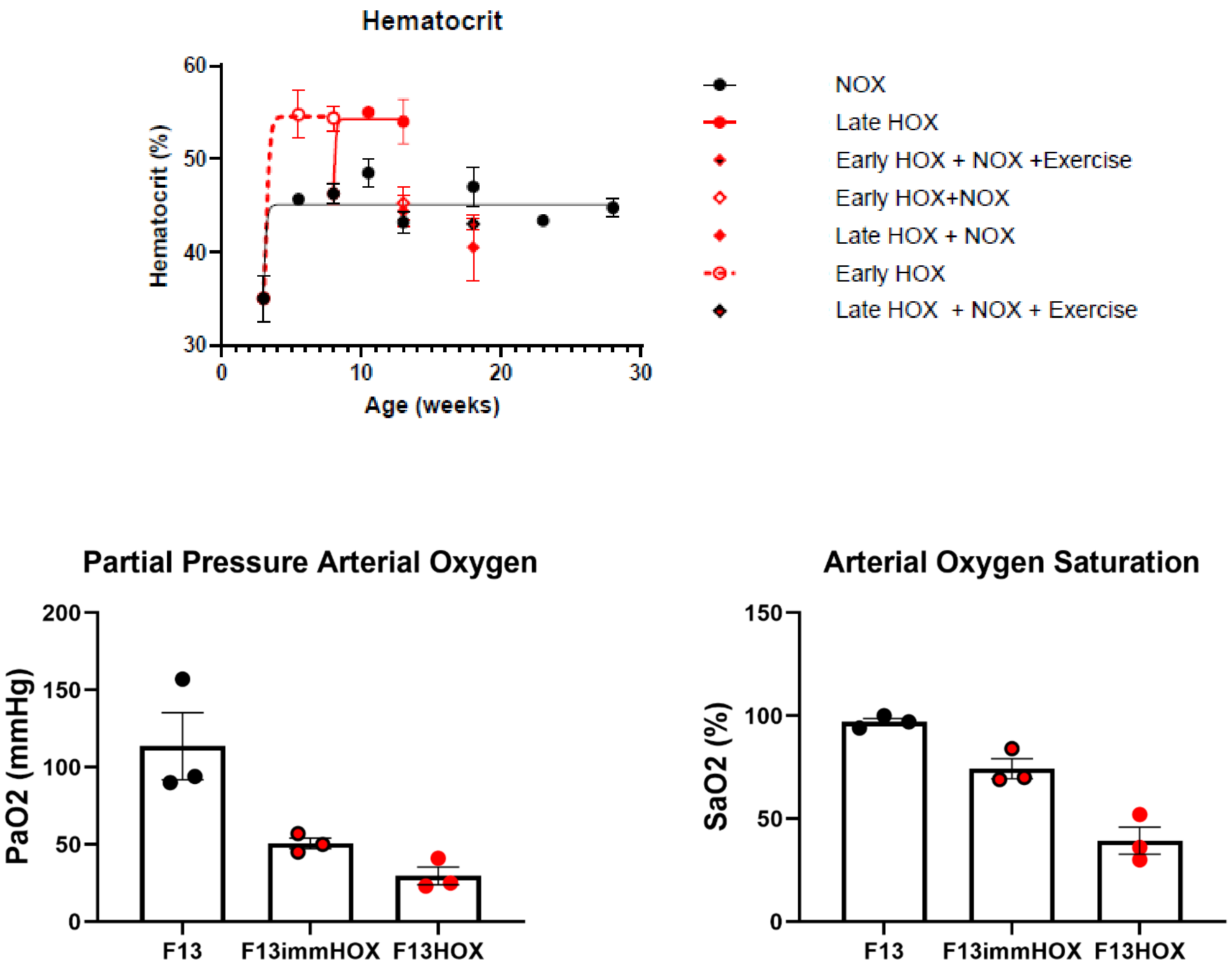


**Figure S9. Hematocrit and Oxygen Saturation of Arterial Blood Samples.** F13 = normoxic control; F13_immediate_HOX = arterial blood sampling of 13-week normoxic control immediately upon exposure to ambient hypoxia (FiO2 10%); F13HOX = 13 week old previously exposed to 5 weeks of chronic hypoxia (FiO2 10%).

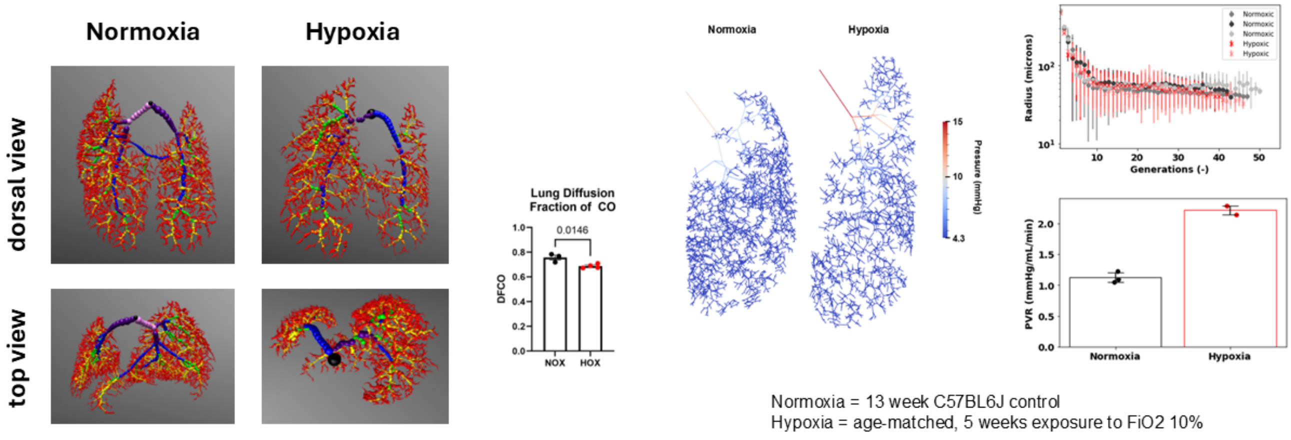


**Figure S10. Hypoxia-induced "pruning" associates with decreases in gas exchange units in mice.** Computed tomography of pulmonary vasculature of mouse lung enabled qualitative visualization of "pruning" (left images). Pruning associated with decreased diffusion of CO from alveoli to the pulmonary vasculature. Quantitative analysis also found pruning of the pulmonary vasculature. Mice exposed to chronic hypoxia beginning at 8 weeks of age (FiO2 10% for five weeks, n=2) had fewer orders of lung branches than normoxic controls (n=3) resulting in increased pulmonary vascular resistance.

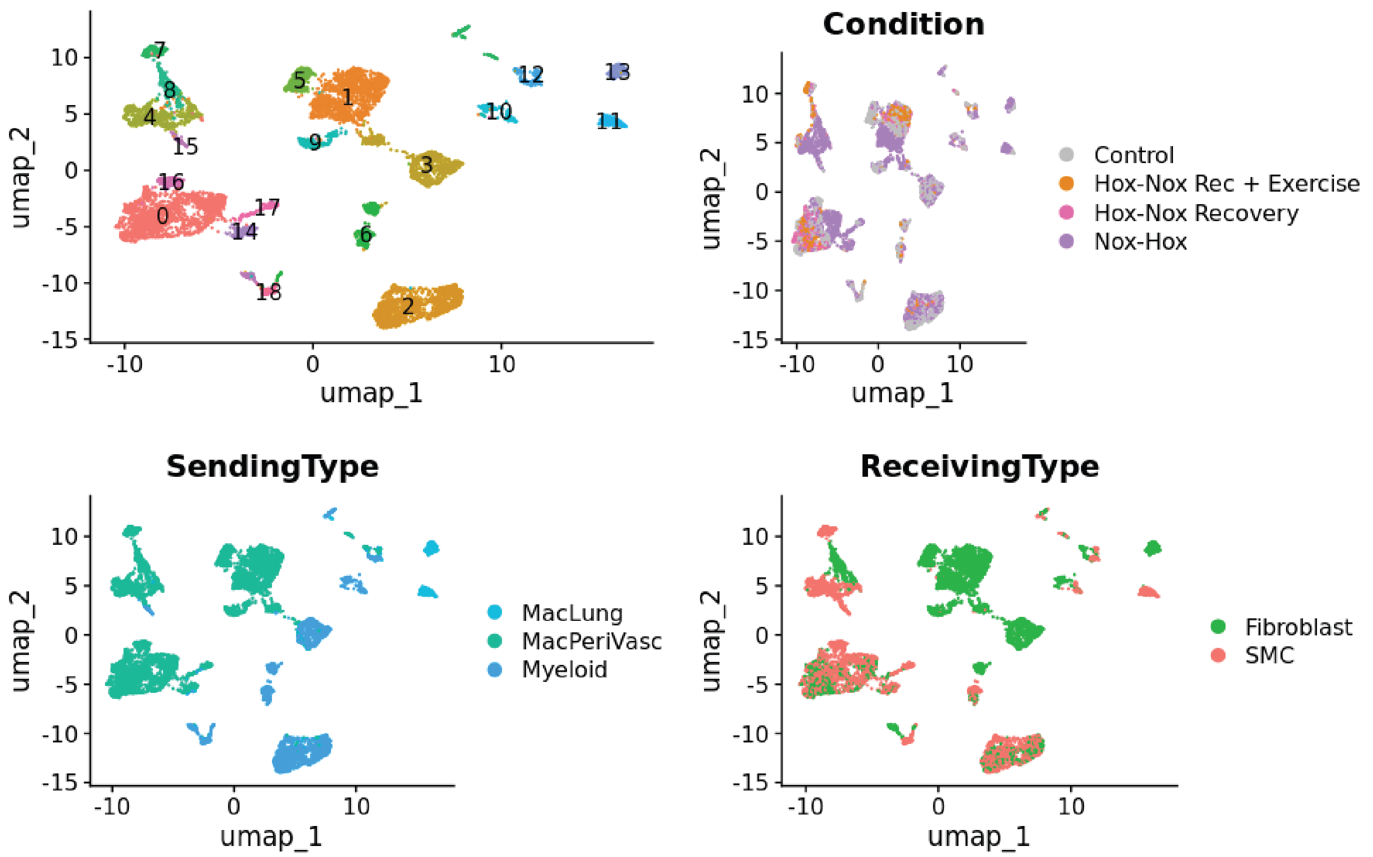


**Figure S11. NICHES archetype embedding.** Independent signaling clusters are identified in the top left. Top right panel separates cells within the signaling clusters based on condition. Sending cell types are identified in the bottom left panel, and receiving cell types are identified by cell type in the bottom right panel.

#### Supplemental Tables

#### Table S1. Sample Size Summary Table

| | | Single Cell RNA Sequencing | Biomechanics | Pulmonary Function Mechanics | Lung Histology | Echocardiography | Voluntary Exercise Monitoring | Cariopulmonary Exercise Testing | Multiphoton Imaging | Hematocrit |
|---|---|---|---|---|---|---|---|---|---|---|
| Female 3 week old normoxia | **F3** | 6 | 5 | 5 | 0 | 4 | 0 | 3 | 4 | 4 |
| | **F3HOX** | 3 | | 5 | | 5 | | | | |
| Female 5.5 week old normoxia | **F5.5** | 3 | 4 | 5 | 0 | 5 | 0 | 0 | 4 | 3 |
| Female 5.5 week old (2.5 weeks hypoxia) | **F5.5HOX** | 3 | 5 | 3 | 0 | 3 | 0 | 0 | 3 | 4 |
| Female 8 week old normoxia | **F8** | 3 | 4 | 5 | 3 | 4 | 0 | 5 | 3 | 4 |
| Female 8 week old (5 weeks hypoxia) | **F8HOX** | 6 | 5 | 5 | 4 | 4 | 0 | 5 | 4 | 5 |
| | **F8R** | 3 | | | | 3 | | | 0 | |
| Female 10.5 week old normoxia | **F10.5** | 3 | 4 | 5 | 0 | 5 | 0 | 0 | 3 | 2 |
| Female 10.5 week old (2.5 week hypoxia) | **F10.5HOX** | 3 | 5 | 5 | 0 | 4 | 0 | 0 | 3 | 2 |
| Female 13 week old normoxia | **F13** | 3 | 5 | 5 | 3 | 5 | 0 | 5 | 5 | 5 |
| Female 13 week old (5 weeks hypoxia) | **F13HOX** | 3 | 5 | 5 | 3 | 5 | 0 | 0 | 3 | 5 |
| Female 13 week old (5 weeks hypoxia followed by 5 weeks normoxia) | **F13R** | 3 | 3 | 4 | 0 | 5 | 0 | 0 | 5 | 4 |
| Female 13 week old (5 weeks hypoxia followed by 5 weeks normoxia with exercise) | **F13RE** | 3 | 0 | 3 | 0 | 5 | 5 | 5 | 5 | 5 |
| Female 13 week old (5 weeks hypoxia + forced exercise) | **F13HOXFE** | | 5 | | | | | | | |
| Female 13 week old (5 weeks hypoxia + rapamycin chow) | **F13HOXrapa** | | 4 | | | | | | | |
| Female 13 week old (5 weeks hypoxia + metformin water) | **F13HOXmetW** | | 4 | | | | | | | |
| Female 13 week old (5 weeks hypoxia + metformin chow) | **F13HOXmetC** | | 4 | | | | | | | |
| Female 13 week old Ccr2$^{-/-}$ normoxia | **FCcr2KO13** | | 4 | | | | | | | |
| Female 13 week old Ccr2$^{-/-}$ (5 weeks hypoxia) | **FCcr2KO13HOX** | | 5 | | | | | | | |
| Female 18 week old normoxia | **F18** | 3 | 5 | 4 | 0 | 5 | 0 | 0 | 2 | 3 |
| Female 18 week old (5 weeks hypoxia followed by 5 weeks normoxia) | **F18R** | 3 | 5 | 4 | 0 | 5 | 0 | 0 | 1 | 4 |
| Female 18 week old (5 weeks hypoxia followed by 5 weeks normoxia with exercise) | **F18RE** | 3 | 3 | 5 | 0 | 5 | 5 | 5 | 2 | 5 |

**Table S2.** Best-fit values of the material parameters in the (pseudo)strain-energy function describing the passive mechanical behavior of the proximal pulmonary artery in juvenile (3 to 8 weeks of age) and adult (8 to 13 weeks of age) mice without and with chronic hypoxic exposure.

Normoxic $FiO_2$ 20%

5wk $FiO_2$ 10%

5wk $FiO_2$ 10%

| | F3 | F5.5 NOX | F5.5 HOX | F8 NOX | F8 HOX | F10.5 NOX | F10.5 HOX | F13 NOX | F13 HOX | F18 NOX |
|---|---|---|---|---|---|---|---|---|---|---|
| | n = 5 | n = 4 | n = 5 | n = 4 | n = 5 | n = 4 | n = 5 | n = 6 | n = 5 | n = 4 |
| **Elastic Fibers** | | | | | | | | | | |
| $c$ (kPa) | 6.064 | 9.545 | 17.191 | 12.913 | 12.659 | 11.486 | 8.349 | 11.335 | 9.919 | 11.986 |
| **Axial Collagen** | | | | | | | | | | |
| $c_1^1$ (kPa) | 4.260 | 1.207 | 0.179 | 1.093 | 0.131 | 0.926 | 0.537 | 0.288 | 0.576 | 0.063 |
| $c_2^1$ | 0.616 | 1.661 | 6.575 | 0.919 | 3.945 | 1.580 | 3.621 | 0.906 | 2.274 | 2.781 |
| **Circumferential CollagenCollagen** | | | | | | | | | | |
| $c_1^2$ (kPa) | 2.515 | 2.283 | 2.183 | 1.404 | 3.356 | 2.266 | 3.892 | 0.648 | 3.449 | 0.607 |
| $c_2^2$ | 0.000 | 1.674 | 0.374 | 0.200 | 0.231 | 1.725 | 1.023 | 0.468 | 0.073 | 0.502 |
| **Diagonal Collagen** | | | | | | | | | | |
| $c_1^{3,4}$ (kPa) | 4.574 | 3.237 | 2.766 | 1.932 | 3.447 | 2.129 | 3.537 | 3.306 | 3.559 | 2.431 |
| $c_2^{3,4}$ | 0.438 | 0.542 | 2.296 | 0.565 | 1.329 | 0.719 | 1.266 | 0.363 | 0.960 | 0.645 |
| $\alpha_0$ (deg) | 37.725 | 47.622 | 40.705 | 41.500 | 41.587 | 44.490 | 42.814 | 42.522 | 42.643 | 41.637 |
| **Error (RMSE)** | 0.12 | 0.13 | 0.07 | 0.10 | 0.07 | 0.10 | 0.09 | 0.09 | 0.07 | 0.08 |

**Table S3**. Key geometric and passive biomechanical metrics for the proximal pulmonary artery in juvenile (3 to 8 weeks of age) and adult (8 to 13 weeks of age) mice without and with two different durations (2.5 or 5.0 weeks) of hypoxic exposure calculated at the mean *in vivo* arterial pressure (15 mmHg for normoxia, 25 mmHg for hypoxia).

Normoxic $FiO_2$ 20%

5wk $FiO_2$ 10%

5wk $FiO_2$ 10%

| | F3 | F5.5 NOX | F5.5 HOX | F8 NOX | F8 HOX | F10.5 NOX | F10.5 HOX | F13 NOX | F13 HOX | F18 NOX |
|---|---|---|---|---|---|---|---|---|---|---|
| | n = 5 | n = 4 | n = 5 | n = 4 | n = 5 | n = 4 | n = 5 | n = 6 | n = 5 | n = 4 |
| **Unloaded Dimensions** | | | | | | | | | | |
| Wall Thickness (µm) | 56.14 ± 1.16 | 56.48 ± 1.80 | 58.23 ± 1.11 | 63.72 ± 4.91 | 62.81 ± 0.69 | 58.74 ± 0.66 | 72.82 ± 1.68 | 66.50 ± 3.18 | 74.94 ± 1.52 | 67.71 ± 1.62 |
| Outer Diameter (µm) | 647.02 ± 28.88 | 851.43 ± 15.73 | 887.15 ± 7.49 | 810.83 ± 5.59 | 865.23 ± 6.73 | 862.89 ± 9.44 | 894.25 ± 27.32 | 813.07 ± 28.59 | 906.61 ± 17.42 | 831.52 ± 9.32 |
| Axial Length (mm) | 2.07 ± 0.06 | 2.70 ± 0.22 | 2.75 ± 0.15 | 2.31 ± 0.13 | 3.06 ± 0.20 | 2.61 ± 0.11 | 2.32 ± 0.37 | 2.39 ± 0.14 | 3.01 ± 0.29 | 2.54 ± 0.05 |
| **Loaded Dimensions** | **P = 15.0** | **P = 15.0** | **P = 25.0** | **P = 15.0** | **P = 25.0** | **P = 15.0** | **P = 25.0** | **P = 15.0** | **P = 25.0** | **P = 15.0** |
| Wall Thickness (µm) | 26.35 ± 1.59 | 24.50 ± 0.69 | 30.93 ± 0.49 | 29.20 ± 2.12 | 31.16 ± 0.54 | 25.89 ± 0.83 | 34.95 ± 1.51 | 28.18 ± 1.01 | 34.80 ± 1.30 | 30.31 ± 1.20 |
| Outer Diameter (µm) | 899.88 ± 17.11 | 1193.41 ± 14.42 | 1259.42 ± 9.60 | 1110.37 ± 22.31 | 1229.56 ± 17.65 | 1232.63 ± 14.15 | 1314.74 ± 39.04 | 1122.96 ± 51.97 | 1314.30 ± 31.25 | 1211.64 ± 60.03 |
| Inner Radius (µm) | 423.59 8.08 | 572.20 6.58 | 598.7786 ± 4.66 | 525.99 10.32 | 583.62 ± 8.99 | 590.43 7.87 | 622.42 ± 20.27 | 533.30 26.06 | 622.35 ± 16.27 | 575.51 30.83 |
| *in vivo* Axial Stretch ($\lambda_z^{iv}$) | 1.45 ± 0.02 | 1.57 ± 0.02 | 1.27 ± 0.01 | 1.51 ± 0.03 | 1.35 ± 0.01 | 1.52 ± 0.03 | 1.37 ± 0.06 | 1.60 ± 0.03 | 1.40 ± 0.03 | 1.42 ± 0.04 |
| *in vivo* Circumferential Stress ($\lambda_\vartheta^{iv}$) | 1.49 ± 0.07 | 1.47 ± 0.02 | 1.48 ± 0.01 | 1.45 ± 0.03 | 1.49 ± 0.02 | 1.50 ± 0.03 | 1.56 ± 0.04 | 1.48 ± 0.04 | 1.54 ± 0.04 | 1.52 ± 0.08 |
| **Cacucy Stresses (kPa)** | | | | | | | | | | |
| Circumferential, $\sigma_\vartheta$ | 32.59 ± 2.01 | 46.78 ± 0.91 | 64.58 ± 1.02 | 36.49 ± 2.30 | 62.54 ± 1.72 | 45.81 ± 2.15 | 59.95 ± 3.82 | 38.12 ± 2.37 | 60.06 ± 3.31 | 38.34 ± 3.42 |
| Axial, $\sigma_z$ | 45.42 ± 4.18 | 48.93 ± 2.18 | 46.93 ± 1.95 | 45.01 ± 3.23 | 48.63 ± 1.76 | 46.13 ± 1.27 | 41.73 ± 4.28 | 50.49 ± 3.02 | 48.28 ± 3.21 | 36.51 ± 3.23 |
| **Linearized Stiffness (MPa)** | | | | | | | | | | |
| Circumferential, $C_{\theta\theta\theta\theta}$ | 0.14 ± 0.01 | 0.23 ± 0.01 | 0.43 ± 0.03 | 0.13 ± 0.01 | 0.39 ± 0.01 | 0.21 ± 0.02 | 0.41 ± 0.06 | 0.13 ± 0.01 | 0.35 ± 0.03 | 0.15 ± 0.02 |
| Axial, $C_{zzzz}$ | 0.26 ± 0.02 | 0.25 ± 0.02 | 0.37 ± 0.04 | 0.22 ± 0.01 | 0.37 ± 0.02 | 0.24 ± 0.02 | 0.30 ± 0.04 | 0.24 ± 0.02 | 0.35 ± 0.02 | 0.17 ± 0.02 |
| **Distensibility ($mmHg^{-1}$)** | 259.11 ± 18.50 | 195.32 ± 19.97 | 195.18 ± 3.04 | 336.66 ± 23.52 | 198.21 ± 11.02 | 235.11 ± 29.77 | 194.21 ± 13.43 | 387.44 ± 36.69 | 217.57 ± 21.24 | 300.62 ± 33.77 |
| **Disetnsibility (1/MPa)** | 0.0345 ± 0.0025 | 0.0260 ± 0.0027 | 0.0260 ± 0.0004 | 0.0449 ± 0.0031 | 0.0264 ± 0.0015 | 0.0313 ± 0.0040 | 0.0259 ± 0.0018 | 0.0517 ± 0.0049 | 0.0290 ± 0.0028 | 0.0401 ± 0.0045 |
| **PWV (m/s)** | 0.17 ± 0.01 | 0.19 ± 0.01 | 0.17 ± 0.00 | 0.15 ± 0.01 | 0.17 ± 0.01 | 0.18 ± 0.01 | 0.18 ± 0.01 | 0.14 ± 0.01 | 0.17 ± 0.01 | 0.16 ± 0.01 |
| **Stored Energy (kPa)** | 10.52 ± 0.86 | 14.38 ± 1.46 | 12.93 ± 0.32 | 13.15 ± 0.98 | 13.36 ± 0.61 | 14.22 ± 1.22 | 12.45 ± 1.36 | 15.12 ± 1.12 | 13.84 ± 1.28 | 11.80 ± 1.04 |

**Table S4**. Key geometric and passive biomechanical metrics for the proximal pulmonary artery in juvenile (3 to 8 weeks of age) and adult (8 to 13 weeks of age) mice without and with two different durations (2.5 or 5.0 weeks) of hypoxic exposure calculated at a common *in vivo* arterial pressure (15 mmHg).

Normoxic $FiO_2$ 20%

5wk $FiO_2$ 10%

5wk $FiO_2$ 10%

| | F3 | F5.5 NOX | F5.5 HOX | F8 NOX | F8 HOX | F10.5 NOX | F10.5 HOX | F13 NOX | F13 HOX | F18 NOX |
|---|---|---|---|---|---|---|---|---|---|---|
| | n = 5 | n = 4 | n = 5 | n = 4 | n = 5 | n = 4 | n = 5 | n = 6 | n = 5 | n = 4 |
| **Unloaded Dimensions** | | | | | | | | | | |
| Wall Thickness (µm) | 56.14 ± 1.16 | 56.48 ± 1.80 | 58.23 ± 1.11 | 63.72 ± 4.91 | 62.81 ± 0.69 | 58.74 ± 0.66 | 72.82 ± 1.68 | 66.50 ± 3.18 | 74.94 ± 1.52 | 67.71 ± 1.62 |
| Outer Diameter (µm) | 647.02 ± 28.88 | 851.43 ± 15.73 | 887.15 ± 7.49 | 810.83 ± 5.59 | 865.23 ± 6.73 | 862.89 ± 9.44 | 894.25 ± 27.32 | 813.07 ± 28.59 | 906.61 ± 17.42 | 831.52 ± 9.32 |
| Axial Length (mm) | 2.07 ± 0.06 | 2.70 ± 0.22 | 2.75 ± 0.15 | 2.31 ± 0.13 | 3.06 ± 0.20 | 2.61 ± 0.11 | 2.32 ± 0.37 | 2.39 ± 0.14 | 3.01 ± 0.29 | 2.54 ± 0.05 |
| **Loaded Dimensions** | P = 15.0 | P = 15.0 | P = 15.0 | P = 15.0 | P = 15.0 | P = 15.0 | P = 15.0 | P = 15.0 | P = 15.0 | P = 15.0 |
| Wall Thickness (µm) | 26.35 ± 1.59 | 24.50 ± 0.69 | 36.31 ± 0.51 | 29.20 ± 2.12 | 36.55 ± 0.56 | 25.89 ± 0.83 | 40.47 ± 1.67 | 28.18 ± 1.01 | 41.41 ± 1.17 | 30.31 ± 1.20 |
| Outer Diameter (µm) | 899.88 ± 17.11 | 1193.41 ± 14.42 | 1083.06 ± 14.61 | 1110.37 ± 22.31 | 1057.77 ± 10.83 | 1232.63 ± 14.15 | 1146.32 ± 40.82 | 1122.96 ± 51.97 | 1115.06 ± 23.92 | 1211.64 ± 60.03 |
| Inner Radius (µm) | 423.59 8.08 | 572.20 6.58 | 505.22 7.32 | 525.99 10.32 | 492.34 5.38 | 590.43 7.87 | 532.69 21.25 | 533.30 26.06 | 516.12 12.17 | 575.51 30.83 |
| *in vivo* Axial Stretch ($\lambda_z^{iv}$) | 1.45 ± 0.02 | 1.57 ± 0.02 | 1.27 ± 0.01 | 1.51 ± 0.03 | 1.35 ± 0.01 | 1.52 ± 0.03 | 1.37 ± 0.06 | 1.60 ± 0.03 | 1.40 ± 0.03 | 1.42 ± 0.04 |
| *in vivo* Circumferential Stress ($\lambda_\theta^{iv}$) | 1.49 ± 0.07 | 1.47 ± 0.02 | 1.26 ± 0.01 | 1.45 ± 0.03 | 1.27 ± 0.01 | 1.50 ± 0.03 | 1.35 ± 0.03 | 1.48 ± 0.04 | 1.29 ± 0.03 | 1.52 ± 0.08 |
| **Cacucy Stresses (kPa)** | | | | | | | | | | |
| Circumferential, $\sigma_\theta$ | 32.59 ± 2.01 | 46.78 ± 0.91 | 27.85 ± 0.58 | 36.49 ± 2.30 | 26.97 ± 0.53 | 45.81 ± 2.15 | 26.59 ± 1.85 | 38.12 ± 2.37 | 25.02 ± 1.02 | 38.34 ± 3.42 |
| Axial, $\sigma_z$ | 45.42 ± 4.18 | 48.93 ± 2.18 | 29.47 ± 1.30 | 45.01 ± 3.23 | 31.87 ± 1.14 | 46.13 ± 1.27 | 26.10 ± 3.27 | 50.49 ± 3.02 | 31.77 ± 2.04 | 36.51 ± 3.23 |
| **Linearized Stiffness (MPa)** | | | | | | | | | | |
| Circumferential, $C_{\theta\theta\theta\theta}$ | 0.14 ± 0.01 | 0.23 ± 0.01 | 0.11 ± 0.00 | 0.13 ± 0.01 | 0.12 ± 0.00 | 0.21 ± 0.02 | 0.12 ± 0.01 | 0.13 ± 0.01 | 0.11 ± 0.01 | 0.15 ± 0.02 |
| Axial, $C_{zzzz}$ | 0.26 ± 0.02 | 0.25 ± 0.02 | 0.18 ± 0.02 | 0.22 ± 0.01 | 0.22 ± 0.01 | 0.24 ± 0.02 | 0.16 ± 0.02 | 0.24 ± 0.02 | 0.22 ± 0.02 | 0.17 ± 0.02 |
| **Distensibility ($mmHg^{-1}$)** | 259.11 ± 18.50 | 195.32 ± 19.97 | 195.18 ± 3.04 | 336.66 ± 23.52 | 198.21 ± 11.02 | 235.11 ± 29.77 | 194.21 ± 13.43 | 387.44 ± 36.69 | 217.57 ± 21.24 | 300.62 ± 33.77 |
| **Disetnsibility (1/MPa)** | 0.0345 ± 0.0025 | 0.0260 ± 0.0027 | 0.0260 ± 0.0004 | 0.0449 ± 0.0031 | 0.0264 ± 0.0015 | 0.0313 ± 0.0040 | 0.0259 ± 0.0018 | 0.0517 ± 0.0049 | 0.0290 ± 0.0028 | 0.0401 ± 0.0045 |
| **PWV (m/s)** | 0.17 ± 0.01 | 0.19 ± 0.01 | 0.17 ± 0.00 | 0.15 ± 0.01 | 0.17 ± 0.01 | 0.18 ± 0.01 | 0.18 ± 0.01 | 0.14 ± 0.01 | 0.17 ± 0.01 | 0.16 ± 0.01 |
| **Stored Energy (kPa)** | 10.52 ± 0.86 | 14.38 ± 1.46 | 6.13 ± 0.21 | 13.15 ± 0.98 | 6.73 ± 0.29 | 14.22 ± 1.22 | 6.58 ± 0.97 | 15.12 ± 1.12 | 6.89 ± 0.39 | 11.80 ± 1.04 |

**Table S5.** Best-fit values of the material parameters in the (pseudo)strain-energy function describing the passive mechanical behavior of the proximal pulmonary artery in the neonatal mice without and with chronic hypoxic exposure (10% FiO2).

| | | | | |
|---|---|---|---|---|
| | Normoxic $FiO_2$ 20% | | | 8wk $FiO_2$ 10% |
| | 10% | | | |
| birth | F3 | F3 HOX | F8 NOX | F8 HOX' |
| | n = 5 | n = 5 | n = 4 | n = 4 |
| **Elastic Fibers** | | | | |
| $c$ (kPa) | 6.064 | 5.019 | 12.913 | 2.4228 |
| **Axial Collagen** | | | | |
| $c_1^1$ (kPa) | 4.260 | 3.655 | 1.093 | 13.311 |
| $c_2^1$ | 0.616 | 5.039 | 0.919 | 1.2768 |
| **Circumferential CollagenCollagen** | | | | |
| $c_1^2$ (kPa) | 2.515 | 7.323 | 1.404 | 4.2837 |
| $c_2^2$ | 0.000 | 2.835 | 0.200 | 0.0185 |
| **Diagonal Collagen** | | | | |
| $c_1^{3,4}$ (kPa) | 4.574 | 12.10646 | 1.932 | 11.824 |
| $c_2^{3,4}$ | 0.438 | 2.180345 | 0.565 | 0.4255 |
| $\alpha_0$ (deg) | 37.725 | 38.00453 | 41.500 | 36.153 |
| **Error (RMSE)** | 0.12 | 0.115361 | 0.10 | 0.0828 |

**Table S6.** Key geometric and passive biomechanical metrics for the proximal pulmonary artery in neonatal mice without and with chronic (birth to 3 weeks of age, birth to 8 weeks of age) exposure to hypoxia.

Normoxic $FiO_2$ 20%

8wk $FiO_2$ 10%

3wk FiO2 10%

| birth | F3 | F3 HOX | F8 NOX | F8 HOX' |
|---|---|---|---|---|
| | n = 5 | n = 5 | n = 4 | n = 5 |
| **Unloaded Dimensions** | | | | |
| Wall Thickness (μm) | 56.14 ± 1.16 | 61.52 ± 2.47 | 63.72 ± 4.91 | 67.70 ± 3.03 |
| Outer Diameter (μm) | 647.02 ± 28.88 | 682.37 ± 27.99 | 810.83 ± 5.59 | 704.26 ± 43.61 |
| Axial Length (mm) | 2.07 ± 0.06 | 2.59 ± 0.09 | 2.31 ± 0.13 | 3.16 ± 0.21 |
| **Loaded Dimensions** | P = 15.0 | P = 25.0 | P = 15.0 | P = 25.0 |
| Wall Thickness (μm) | 26.35 ± 1.59 | 36.26 ± 1.78 | 29.20 ± 2.12 | 32.50 ± 1.45 |
| Outer Diameter (μm) | 899.88 ± 17.11 | 861.73 ± 23.97 | 1110.37 ± 22.31 | 1041.40 ± 121.14 |
| Inner Radius (μm) | 423.59 8.08 | 394.60 12.93 | 525.99 10.32 | 488.20 59.70 |
| *in vivo* Axial Stretch ($\lambda_z^{iv}$) | 1.45 ± 0.02 | 1.28 ± 0.05 | 1.51 ± 0.03 | 1.33 ± 0.05 |
| *in vivo* Circumferential Stress ($\lambda_\theta^{iv}$) | 1.49 ± 0.07 | 1.34 ± 0.04 | 1.45 ± 0.03 | 1.57 ± 0.10 |
| **Cacucy Stresses (kPa)** | | | | |
| Circumferential, $\sigma_\theta$ | 32.59 ± 2.01 | 36.75 ± 2.59 | 36.49 ± 2.30 | 49.87 ± 4.79 |
| Axial, $\sigma_z$ | 45.42 ± 4.18 | 38.72 ± 5.15 | 45.01 ± 3.23 | 74.50 ± 6.66 |
| **Linearized Stiffness (MPa)** | | | | |
| Circumferential, $C_{\theta\theta\theta\theta}$ | 0.14 ± 0.01 | 0.21 ± 0.01 | 0.13 ± 0.01 | 0.26 ± 0.05 |
| Axial, $C_{zzzz}$ | 0.26 ± 0.02 | 0.37 ± 0.02 | 0.22 ± 0.01 | 0.59 ± 0.03 |
| **Distensibility (mmHg$^{-1}$)** | 259.11 ± 18.50 | 149.40 ± 10.24 | 336.66 ± 23.52 | 224.66 ± 36.42 |
| **Disetnsibility (1/MPa)** | 0.0345 ± 0.0025 | 0.02 ± 0.00 | 0.04 ± 0.00 | 0.03 ± 0.00 |
| **PWV (m/s)** | 0.17 ± 0.01 | 0.20 ± 0.01 | 0.15 ± 0.01 | 0.17 ± 0.01 |
| **Stored Energy (kPa)** | 10.52 ± 0.86 | 7.12 ± 0.94 | 13.15 ± 0.98 | 13.82 ± 0.90 |

### Table S7. Fibroblast Differentially Expressed Genes: Hypoxia versus Normoxia

**Fibroblasts: Genes that positively correlate with chronic hypoxia**

| Column1 | Beta0 | Beta1 | Disp | Sigma | PiC | PiI | P.Beta | P.Pi | Hessian | Convergence | Message | AIC | BIC | logLik | deviance | df.resid | fdrP |
|---|---|---|---|---|---|---|---|---|---|---|---|---|---|---|---|---|---|
| Map7d2 | -12.05446702 | 0.77684537 | 1.065187644 | 0.689458491 | 1.888315428 | 0.472351817 | 0.5 | 0.002257231 | FALSE | 0 | relative convergence (4) | 5045.35044369939 | 5095.85982833422 | -2515.67522184969 | 5031.35044369939 | 10046 | 0.043221201 |
| Eif2s3y | -17.32158997 | 1.650340826 | -9.874019651 | -12.54103645 | 3.906606397 | 3.074077022 | 0.5 | 3.78359E-06 | FALSE | 0 | relative convergence (4) | 970.568944352648 | 1021.07832898749 | -478.284472176324 | 956.568944352648 | 10046 | 0.000598114 |
| Ddx3y | -14.56111251 | 0.992391834 | -1.032370809 | -17.85288267 | 3.528549272 | 1.689541202 | 0.5 | 8.55713E-06 | FALSE | 0 | relative convergence (4) | 1527.96349824796 | 1578.4728828828 | -756.981749123981 | 1513.96349824796 | 10046 | 0.001118598 |
| mt-Ts1 | -13.33796056 | 1.00103188 | 0.531681063 | 2.439379081 | 0.167322097 | 0.861536252 | 0.5 | 0.000439848 | FALSE | 0 | relative convergence (4) | 2771.08657642705 | 2821.59596106189 | -1378.54328821352 | 2757.08657642705 | 10046 | 0.016031475 |
| Septin6 | -10.42666122 | 0.46111793 | 0.085182345 | 0.07858188 | 1.083956294 | 0.343555673 | 0.5 | 0.000150404 | FALSE | 0 | relative convergence (4) | 11284.5723975121 | 11335.0817821469 | -5635.28619875605 | 11270.5723975121 | 10046 | 0.00787839 |
| Fgf16 | -12.11722983 | 0.616086733 | 0.50082941 | 0.369436334 | 3.9239626 | 0.000101906 | 0.5 | 0.000164755 | FALSE | 0 | relative convergence (4) | 3815.64122150214 | 3866.15060613698 | -1900.82061075107 | 3801.64122150214 | 10046 | 0.008420461 |
| Pgk1 | -9.783452477 | 0.409483049 | -0.109267684 | 0.273738108 | 0.308433142 | 0.000111349 | 0.5 | 0.001027845 | FALSE | 0 | relative convergence (4) | 16285.467703706 | 16335.9770883409 | -8135.73385185302 | 16271.467703706 | 10046 | 0.026769267 |
| Tspan6 | -11.01459159 | 0.227642009 | -0.062822799 | 0.361473771 | 0.622370681 | 0.000264763 | 0.5 | 0.000707556 | FALSE | 0 | relative convergence (4) | NA | NA | NA | NA | 10046 | 0.021519528 |
| Tceal6 | -13.95106881 | 1.744841992 | 2.330261008 | 2.323412718 | 7.119906407 | 0.344805529 | 0.5 | 0.00039412 | FALSE | 0 | relative convergence (4) | 2322.6953739846 | 2373.20475861944 | -1154.3476869923 | 2308.6953739846 | 10046 | 0.01528425 |
| Ehbp1l1 | -9.839583538 | 0.483577076 | 0.290164924 | 0.431274591 | 0.348409752 | 0.091789081 | 0.5 | 0.000200584 | FALSE | 0 | relative convergence (4) | 16016.5426476246 | 16067.0520322595 | -8001.27132381231 | 16002.5426476246 | 10046 | 0.009615485 |
| Pola2 | -11.53302146 | 0.272188544 | -0.127982582 | -0.216137064 | 1.313609992 | 6.45968E-11 | 0.5 | 0.001857316 | FALSE | 1 | singular convergence (7) | NA | NA | NA | NA | 10046 | 0.038532416 |
| Gpr137 | -11.6129422 | 0.356540706 | -0.1304003 | 0.31532845 | 0.035294387 | 0.064564635 | 0.5 | 2.71511E-05 | FALSE | 0 | relative convergence (4) | 4938.30222146932 | 4988.81160610415 | -2462.15111073466 | 4924.30222146932 | 10046 | 0.002469284 |
| Otub1 | -10.48382761 | 0.170206138 | 0.156026841 | -0.352927132 | 5.96579E-09 | 0.146585796 | 0.5 | 0.001591957 | FALSE | 1 | false convergence (8) | 9363.25616185005 | 9413.76554648489 | -4674.62808092503 | 9349.25616185005 | 10046 | 0.035273898 |
| Ddb1 | -9.237208638 | 0.279184954 | -0.095248235 | 0.164819659 | 0.079037036 | 0.000352684 | 0.5 | 0.001557777 | FALSE | 0 | relative convergence (4) | 20254.7927311143 | 20305.3021157492 | -10120.3963655572 | 20240.7927311143 | 10046 | 0.034996369 |
| Tmem132a | -11.65126142 | 0.80666069 | 0.241990766 | 0.616853792 | 0.732469766 | 0.400864225 | 0.5 | 6.3557E-06 | FALSE | 0 | relative convergence (4) | 6450.1965032778 | 6500.70588791264 | -3218.0982516389 | 6436.1965032778 | 10046 | 0.00090006 |
| Pitpnm1 | -11.66743082 | 0.40214786 | 0.19221559 | 0.403445776 | 0.108388495 | 1.9636E-05 | 0.5 | 0.000788007 | FALSE | 0 | relative convergence (4) | 4903.45332125234 | 4953.96270588717 | -2444.72666062617 | 4889.45332125234 | 10046 | 0.022810305 |
| Sart1 | -10.7408824 | 0.240809631 | 0.257194248 | 0.303188902 | 0.114980118 | 0.164957898 | 0.5 | 1.96356E-05 | FALSE | 0 | relative convergence (4) | 8657.01724299637 | 8707.52662763121 | -4321.50862149819 | 8643.01724299637 | 10046 | 0.002022318 |
| Fibp | -10.56543949 | 0.222660856 | -0.015776796 | 0.2671049 | 3.25626E-07 | 0.101881436 | 0.5 | 1.95901E-05 | FALSE | 1 | false convergence (8) | NA | NA | NA | NA | 10046 | 0.002022318 |
| Hs6st2 | -12.62750699 | 1.029201617 | -0.153894614 | 0.116677628 | 11.83591962 | 0.250241022 | 0.5 | 8.65452E-05 | FALSE | 0 | relative convergence (4) | 2815.14396096403 | 2865.65334559887 | -1400.57198048201 | 2801.14396096403 | 10046 | 0.005323902 |
| Eno1 | -9.799815823 | 0.612274814 | 0.350677449 | -0.151164821 | 0.383197909 | 0.09393069 | 0.5 | 8.16385E-05 | FALSE | 0 | relative convergence (4) | 16545.8659092432 | 16596.3752938781 | -8265.93295462161 | 16531.8659092432 | 10046 | 0.005114632 |
| Ankrd1 | -12.56665457 | 0.960357007 | 0.055257883 | 0.192675155 | 7.900959041 | 0.390275551 | 0.5 | 0.001367803 | FALSE | 0 | relative convergence (4) | 3235.48460178526 | 3285.99398642009 | -1610.74230089263 | 3221.48460178526 | 10046 | 0.031950663 |
| Wtip | -10.32617783 | 0.66398002 | 0.490270947 | 0.582451626 | 0.350675463 | 0.000351527 | 0.5 | 0.001387207 | FALSE | 0 | relative convergence (4) | 13195.6732495712 | 13246.182634206 | -6590.8366247856 | 13181.6732495712 | 10046 | 0.032292938 |
| Uba2 | -9.703266699 | 0.196758532 | -0.024375367 | 0.18661242 | 0.112925287 | 0.066062106 | 0.5 | 0.000931669 | FALSE | 0 | relative convergence (4) | 15696.2082020741 | 15746.7175867089 | -7841.10410103706 | 15682.2082020741 | 10046 | 0.025332078 |
| Gpi1 | -9.135200727 | 0.39690142 | 0.123305447 | 0.377478591 | 0.142388458 | 0.171098674 | 0.5 | 1.04433E-05 | FALSE | 0 | relative convergence (4) | 22043.3838905236 | 22093.8932751585 | -11014.6919452618 | 22029.3838905236 | 10046 | 0.001326881 |
| Nudt19 | -11.12829714 | 0.321975351 | 0.045933475 | 0.475645304 | 0.104898188 | 0.000580178 | 0.5 | 0.002481105 | FALSE | 0 | relative convergence (4) | 7110.57752657477 | 7161.0869112096 | -3548.28876328738 | 7096.57752657477 | 10046 | 0.045717476 |
| Dpy19l3 | -11.84837728 | 0.394501335 | 0.269256429 | 0.013353848 | 1.236054262 | 1.85484E-19 | 0.5 | 5.28025E-05 | FALSE | 1 | singular convergence (7) | 4163.13462132059 | 4213.64400595543 | -2074.56731066029 | 4149.13462132059 | 10046 | 0.003966022 |
| 5430431A17Rik | -11.7899438 | 0.863822572 | 1.215318066 | 1.153848403 | 7.945284281 | 0.000178911 | 0.5 | 0.000173364 | FALSE | 0 | relative convergence (4) | 5450.74425829315 | 5501.25364292799 | -2718.37212914657 | 5436.74425829315 | 10046 | 0.008671193 |
| Tbc1d17 | -10.81533301 | 0.210949403 | 0.039551639 | 0.312758963 | 0.018550918 | 4.74966E-05 | 0.5 | 0.000239611 | FALSE | 0 | relative convergence (4) | 8323.18828761316 | 8373.69767224799 | -4154.59414380658 | 8309.18828761316 | 10046 | 0.01078645 |
| Ap2a1 | -10.27753035 | 0.353942039 | 0.211907345 | 0.153879757 | 0.126960621 | 0.079427924 | 0.5 | 0.001903755 | FALSE | 0 | relative convergence (4) | 11671.5967240134 | 11722.1061086483 | -5828.79836200672 | 11657.5967240134 | 10046 | 0.03874483 |
| Hrc | -12.49283794 | 1.042132978 | 0.768578719 | 0.281390865 | 2.553745094 | 0.000142086 | 0.5 | 0.001603392 | FALSE | 0 | relative convergence (4) | 3765.0978802682 | 3815.60726490303 | -1875.5489401341 | 3751.0978802682 | 10046 | 0.035386541 |
| Nucb1 | -9.067033703 | 0.238305212 | -0.127707804 | 0.185015295 | 0.062908806 | 0.110268713 | 0.5 | 0.000118033 | FALSE | 0 | relative convergence (4) | 21399.1698524036 | 21449.6792370385 | -10692.5849262018 | 21385.1698524036 | 10046 | 0.006547632 |
| Dbp | -11.45248203 | 1.667730336 | -0.716119983 | -1.188003426 | 2.172306468 | 0.000363223 | 0.5 | 0.000249657 | FALSE | 0 | relative convergence (4) | 8288.76219451822 | 8339.27157915306 | -4137.38109725911 | 8274.76219451822 | 10046 | 0.011164754 |
| Nomo1 | -10.42852281 | 0.299378054 | -0.077784395 | 0.214800608 | 0.108030099 | 0.040110207 | 0.5 | 0.000474116 | FALSE | 0 | relative convergence (4) | 10543.5799706678 | 10594.0893553027 | -5264.78998533392 | 10529.5799706678 | 10046 | 0.016945621 |
| Ldha | -9.14583293 | 0.472994774 | 0.083596947 | 0.214267976 | 0.280703138 | 0.12138094 | 0.5 | 0.000423552 | FALSE | 0 | relative convergence (4) | 22546.565039139 | 22597.0744237738 | -11266.2825195695 | 22532.565039139 | 10046 | 0.015748876 |
| Chpf2 | -11.04074968 | 0.466674894 | 0.038395822 | 0.695411408 | 0.02110594 | 0.166806566 | 0.5 | 1.42775E-13 | FALSE | 0 | relative convergence (4) | NA | NA | NA | NA | 10046 | 3.88205E-10 |
| Emilin1 | -9.737124196 | 0.3750021 | 0.263941085 | 0.503094011 | 0.285708576 | 0.289747402 | 0.5 | 0.000579806 | FALSE | 0 | relative convergence (4) | 17379.0464846394 | 17429.5558692743 | -8682.52324231972 | 17365.0464846394 | 10046 | 0.018812554 |
| Preb | -10.36784768 | 0.16762604 | 0.1422909 | 0.268513672 | 0.108624372 | 0.026744809 | 0.5 | 0.000490547 | FALSE | 0 | relative convergence (4) | NA | NA | NA | NA | 10046 | 0.017242321 |
| Marveld1 | -9.335291706 | 0.551512244 | 0.368631283 | 0.370328804 | 0.433636534 | 0.000434241 | 0.5 | 0.000523376 | FALSE | 0 | relative convergence (4) | 21743.4037479833 | 21793.9131326181 | -10864.7018739916 | 21729.4037479833 | 10046 | 0.017788246 |
| Actr1a | -9.844048892 | 0.158444798 | -0.189730466 | 0.1679659 | 0.120872055 | 0.071510388 | 0.5 | 5.67776E-05 | FALSE | 0 | relative convergence (4) | 14146.6410478702 | 14197.150432505 | -7066.32052393508 | 14132.6410478702 | 10046 | 0.004105806 |
| Akr1c14 | -12.05391564 | 0.369091276 | 0.140095971 | 0.353116194 | 2.013084709 | 0.038348578 | 0.5 | 0.001306139 | FALSE | 0 | relative convergence (4) | NA | NA | NA | NA | 10046 | 0.031067795 |
| Mylk4 | -9.955025558 | 0.964189803 | 1.346613185 | 1.091623302 | 4.122491875 | 0.41482576 | 0.5 | 0.000713317 | FALSE | 0 | relative convergence (4) | 18638.9371849249 | 18689.4465695597 | -9312.46859246243 | 18624.9371849249 | 10046 | 0.021602492 |
| Htatip2 | -11.6700932 | 0.426630407 | 0.030086624 | 0.355447388 | 0.427666543 | 0.179210177 | 0.5 | 0.001109806 | FALSE | 0 | relative convergence (4) | 5120.91051763136 | 5171.4199022662 | -2553.45525881568 | 5106.91051763136 | 10046 | 0.02825432 |
| Acan | -13.41068685 | 1.994445967 | -0.022520932 | 0.060589605 | 8.512223747 | 0.000163592 | 0.5 | 2.63161E-09 | FALSE | 0 | relative convergence (4) | 2573.90026684921 | 2624.40965148405 | -1279.95013342461 | 2559.90026684921 | 10046 | 1.78884E-06 |
| P4ha3 | -13.32518555 | 0.982726388 | -1.787910831 | 0.486671846 | 9.782134264 | 0.500169596 | 0.5 | 1.56905E-06 | FALSE | 0 | relative convergence (4) | NA | NA | NA | NA | 10046 | 0.000309149 |
| P2ry2 | -12.20475201 | 0.411788752 | -0.522586425 | -0.214072949 | 0.807399072 | 3.31744E-39 | 0.5 | 0.000436769 | FALSE | 0 | relative convergence (4) | 3104.60492430214 | 3155.11430893698 | -1545.30246215107 | 3090.60492430214 | 10046 | 0.016005038 |
| Trim47 | -10.02151129 | 0.401116645 | 0.126682981 | 0.200627656 | 0.373297378 | 0.112519164 | 0.5 | 0.001798145 | FALSE | 0 | relative convergence (4) | 14215.7482624824 | 14266.2576471172 | -7100.87413124118 | 14201.7482624824 | 10046 | 0.037841758 |
| Rhbdf2 | -11.84045084 | 0.509141549 | -0.101142176 | -0.401596505 | 1.207018607 | 0.277393773 | 0.5 | 0.002239488 | FALSE | 0 | relative convergence (4) | 4062.1915259372 | 4112.70091057204 | -2024.0957629686 | 4048.1915259372 | 10046 | 0.04316278 |
| Mgat5b | -13.91493309 | 1.318508214 | 0.979954451 | 0.333883653 | 5.172239212 | 0.00014192 | 0.5 | 9.59761E-05 | FALSE | 0 | relative convergence (4) | 1400.88288633067 | 1451.39227096551 | -693.441443165337 | 1386.88288633067 | 10046 | 0.00559998 |
| Septin9 | -10.22781643 | 0.51893157 | 0.432480087 | 0.135356638 | 0.870444329 | 0.134234439 | 0.5 | 0.001307155 | FALSE | 0 | relative convergence (4) | 13288.298945313 | 13338.8083299478 | -6637.1494726565 | 13274.298945313 | 10046 | 0.031067795 |
| Aatk | -12.48508284 | 0.468505235 | -0.088372409 | 0.540379506 | 0.966211965 | 0.000171568 | 0.5 | 0.000259104 | FALSE | 0 | relative convergence (4) | 2780.36137958197 | 2830.8707642168 | -1383.18068979098 | 2766.36137958197 | 10046 | 0.011326444 |
| Slc38a10 | -9.770058976 | 0.200335267 | -0.070014143 | 0.366815804 | 0.165255838 | 0.174156455 | 0.5 | 0.001476503 | FALSE | 0 | relative convergence (4) | 16041.0852492358 | 16091.5946338706 | -8013.5426246179 | 16027.0852492358 | 10046 | 0.033907202 |
| Arhgdia | -8.643614695 | 0.255802766 | -0.114679486 | 0.155941318 | 0.10304469 | 9.07615E-05 | 0.5 | 0.000162455 | FALSE | 0 | relative convergence (4) | 26972.0433251457 | 27022.5527097806 | -13479.0216625729 | 26958.0433251457 | 10046 | 0.008334249 |
| Pcyt2 | -11.17881345 | 0.287618794 | 0.259101682 | 0.421861197 | 0.440720077 | 0.090493584 | 0.5 | 6.17685E-05 | FALSE | 0 | relative convergence (4) | 6864.10751467718 | 6914.61689931202 | -3425.05375733859 | 6850.10751467718 | 10046 | 0.004309979 |
| Slc16a3 | -11.78370106 | 0.926258606 | -0.212638475 | -0.059010287 | 0.811261731 | 0.000128424 | 0.5 | 2.259E-07 | FALSE | 0 | relative convergence (4) | 5125.71522893154 | 5176.22461356638 | -2555.85761446577 | 5111.71522893154 | 10046 | 6.31428E-05 |
| 4930417O22Rik | -13.70731576 | 1.439410986 | 1.08351667 | 0.692764392 | 1.819060685 | 0.000110918 | 0.5 | 0.00079442 | FALSE | 0 | relative convergence (4) | 1774.15894439839 | 1824.66832903323 | -880.079472199197 | 1760.15894439839 | 10046 | 0.022881666 |
| Sost | -9.262876007 | 1.499334726 | 1.095755147 | 0.72137131 | 4.373949821 | 0.338427553 | 0.5 | 0.000307583 | FALSE | 0 | relative convergence (4) | 26467.1940356369 | 26517.7034202717 | -13226.5970178184 | 26453.1940356369 | 10046 | 0.012946112 |
| Dusp3 | -9.381965952 | 0.357121506 | 0.174557831 | -0.024703409 | 0.278923321 | 0.047838419 | 0.5 | 0.000397836 | FALSE | 0 | relative convergence (4) | 18797.2247326274 | 18847.7341172622 | -9391.61236631368 | 18783.2247326274 | 10046 | 0.01528425 |
| Cdc42ep4 | -10.8946052 | 0.199478335 | -0.08381593 | 0.407927657 | 0.312506415 | 0.11087085 | 0.5 | 0.002049362 | FALSE | 0 | relative convergence (4) | NA | NA | NA | NA | 10046 | 0.041153725 |
| Asap2 | -8.834284074 | 0.330829275 | 0.425502302 | 0.139742665 | 0.657744357 | 0.179252545 | 0.5 | 0.001876677 | FALSE | 0 | relative convergence (4) | 25998.8980251374 | 26049.4074097723 | -12992.4490125687 | 25984.8980251374 | 10046 | 0.038593779 |
| Sqle | -11.98442935 | 0.429268677 | -0.185172194 | 0.251554191 | 1.580111724 | 0.391205272 | 0.5 | 4.19303E-05 | FALSE | 0 | relative convergence (4) | 4061.7709598323 | 4112.28034446714 | -2023.88547991615 | 4047.7709598323 | 10046 | 0.003393111 |
| Gfus | -11.22699927 | 0.25859478 | 0.129970154 | 0.318813613 | 3.41144E-07 | 5.32985E-42 | 0.5 | 0.000200868 | FALSE | 1 | false convergence (8) | NA | NA | NA | NA | 10046 | 0.009615485 |
| Scrib | -11.39450088 | 0.289740276 | 0.075833828 | 0.275375104 | 0.132633597 | 2.74787E-18 | 0.5 | 0.000135503 | FALSE | 1 | singular convergence (7) | NA | NA | NA | NA | 10046 | 0.007281268 |
| Plec | -9.133041394 | 0.259593172 | 0.225446769 | 0.246137487 | 0.224192748 | 0.085219952 | 0.5 | 0.000924012 | FALSE | 0 | relative convergence (4) | 21570.1579895683 | 21620.6673742031 | -10778.0789947841 | 21556.1579895683 | 10046 | 0.025174243 |
| Bop1 | -11.13122084 | 0.325209941 | 0.011618692 | 0.04827498 | 0.202074709 | 0.000115764 | 0.5 | 7.29885E-05 | FALSE | 0 | relative convergence (4) | 6604.50539918784 | 6655.01478382267 | -3295.25269959392 | 6590.50539918784 | 10046 | 0.004840384 |
| Scx | -13.06660534 | 1.47036276 | 0.805249476 | 0.733658615 | 4.838582513 | 0.000178605 | 0.5 | 0.000253468 | FALSE | 0 | relative convergence (4) | 2925.77682065449 | 2976.28620528933 | -1455.88841032725 | 2911.77682065449 | 10046 | 0.011224403 |
| Fbxl6 | -11.57981011 | 0.29364455 | 0.111371287 | 0.609571664 | 0.000637672 | 0.158861952 | 0.5 | 0.000423986 | FALSE | 0 | relative convergence (4) | 5267.98584203051 | 5318.49522666535 | -2626.99292101525 | 5253.98584203051 | 10046 | 0.015748876 |
| Kifc2 | -13.16197569 | 0.540105514 | 0.58181627 | 0.449333399 | 1.419117589 | 1.24759E-85 | 0.5 | 0.002254183 | FALSE | 1 | singular convergence (7) | 1753.79895067185 | 1804.30833530669 | -869.899475335924 | 1739.79895067185 | 10046 | 0.043221201 |
| Myh9 | -8.866275046 | 0.175654198 | 0.0949717 | 0.00301224 | 0.423748553 | 0.072726572 | 0.5 | 0.000618509 | FALSE | 0 | relative convergence (4) | 24010.2921287637 | 24060.8015133985 | -11998.1460643818 | 23996.2921287637 | 10046 | 0.019646346 |
| Kctd17 | -10.12086854 | 0.156915168 | -0.20338002 | 0.134727546 | 0.139672368 | 0.139458205 | 0.5 | 0.000384276 | FALSE | 0 | relative convergence (4) | 12486.1782969882 | 12536.687681623 | -6236.0891484941 | 12472.1782969882 | 10046 | 0.015055414 |
| Ltbp2 | -11.03569071 | 1.287439287 | 0.252802703 | -0.144057177 | 3.920018112 | 0.000231403 | 0.5 | 3.24538E-05 | FALSE | 0 | relative convergence (4) | 10209.6603076478 | 10260.1696922827 | -5097.83015382391 | 10195.6603076478 | 10046 | 0.002810248 |
| Fbln5 | -7.918724745 | 0.341197119 | -0.308176956 | -0.038245814 | 0.497135193 | 0.051337452 | 0.5 | 2.60776E-05 | FALSE | 0 | relative convergence (4) | 38505.3631791459 | 38555.8725637807 | -19245.6815895729 | 38491.3631791459 | 10046 | 0.002411738 |
| Wwp2 | -9.35570534 | 0.612618695 | 0.539762105 | 0.30742342 | 0.930676865 | 0.000482099 | 0.5 | 0.001138608 | FALSE | 0 | relative convergence (4) | 22386.403699186 | 22436.9130838208 | -11186.201849593 | 22372.403699186 | 10046 | 0.028559728 |
| Hp | -13.68769493 | 1.224171614 | 0.248777031 | 2.27926973 | 32.18385233 | 0.635730674 | 0.5 | 4.81958E-06 | FALSE | 0 | relative convergence (4) | 1755.83876676558 | 1806.34815140041 | -870.919383382788 | 1741.83876676558 | 10046 | 0.000720024 |
| Bcar1 | -11.50335715 | 0.704954018 | 0.08232463 | 0.37699305 | 0.123458343 | 0.105723654 | 0.5 | 0.00242926 | FALSE | 0 | relative convergence (4) | 5999.85582503142 | 6050.36520966626 | -2992.92791251571 | 5985.85582503142 | 10046 | 0.045302867 |
| 2310022B05Rik | -9.713475113 | 0.645359735 | 0.266008662 | 0.18922458 | 0.386349216 | 0.000231049 | 0.5 | 0.001817995 | FALSE | 0 | relative convergence (4) | 18126.0275784918 | 18176.5369631266 | -9056.01378924588 | 18112.0275784918 | 10046 | 0.038004845 |
| Gdf6 | -13.47718549 | 1.375386375 | -0.622012853 | 0.512115335 | 22.5159407 | 7.64125E-05 | 0.5 | 8.66468E-07 | FALSE | 0 | relative convergence (4) | 1691.57497840415 | 1742.08436303899 | -838.787489202077 | 1677.57497840415 | 10046 | 0.000189994 |
| Egln3 | -12.77508671 | 1.761228428 | 1.381059536 | 1.261379467 | 1.9965397 | 0.234998941 | 0.5 | 9.96032E-05 | FALSE | 0 | relative convergence (4) | 4792.8223934284 | 4843.33177806324 | -2389.4111967142 | 4778.8223934284 | 10046 | 0.005737733 |
| Hif1a | -10.25773398 | 0.440134493 | 0.28579573 | 0.041966159 | 0.773412967 | 0.000128236 | 0.5 | 0.000306319 | FALSE | 0 | relative convergence (4) | 12021.2233386944 | 12071.7327233293 | -6003.61166934721 | 12007.2233386944 | 10046 | 0.012941456 |
| Nubp1 | -11.25852073 | 0.317029956 | 0.170096081 | 0.145637536 | 0.492595994 | 0.100226195 | 0.5 | 2.10298E-05 | FALSE | 0 | relative convergence (4) | 6249.30626009877 | 6299.8156447336 | -3117.65313004938 | 6235.30626009877 | 10046 | 0.002117774 |
| Scarf2 | -10.20009015 | 0.198634822 | -0.043659707 | 0.383357279 | 0.371564997 | 0.267548007 | 0.5 | 1.50499E-05 | FALSE | 0 | relative convergence (4) | 12554.6064994986 | 12605.1158841335 | -6270.30324974932 | 12540.6064994986 | 10046 | 0.001690941 |
| Mif | -10.07647825 | 0.946330146 | 0.558411354 | 0.480173434 | 0.419825766 | 0.000371833 | 0.5 | 0.001520522 | FALSE | 0 | relative convergence (4) | 16867.5299403987 | 16918.0393250336 | -8426.76497019937 | 16853.5299403987 | 10046 | 0.034562383 |
| Col18a1 | -9.013548996 | 1.147947915 | 0.812030645 | 0.647699794 | 2.277643412 | 0.343110154 | 0.5 | 6.52911E-05 | FALSE | 0 | relative convergence (4) | 29660.7011133569 | 29711.2104979917 | -14823.3505566784 | 29646.7011133569 | 10046 | 0.00443816 |
| Pfkl | -11.1968441 | 1.054985004 | 0.343346464 | 0.344105151 | 0.568931369 | 0.021311986 | 0.5 | 0.00056926 | FALSE | 0 | relative convergence (4) | 8817.47375552767 | 8867.98314016251 | -4401.73687776383 | 8803.47375552767 | 10046 | 0.018603568 |
| Pdgfb | -13.28365114 | 1.285030421 | -0.15376113 | 0.532469252 | 6.025040886 | 0.258790008 | 0.5 | 7.40528E-05 | FALSE | 0 | relative convergence (4) | 1990.22579177833 | 2040.73517641317 | -988.112895889164 | 1976.22579177833 | 10046 | 0.004863516 |
| Pim3 | -11.2830075 | 0.269036119 | -0.496840835 | 0.130825093 | 0.973169312 | 0.233264858 | 0.5 | 0.001013021 | FALSE | 0 | relative convergence (4) | 5687.29012650113 | 5737.79951113597 | -2836.64506325056 | 5673.29012650113 | 10046 | 0.026586911 |
| Plxnb2 | -9.955673325 | 0.263988639 | -0.27032294 | 0.341484861 | 0.087789919 | 0.128591477 | 0.5 | 4.97429E-12 | FALSE | 0 | relative convergence (4) | 14164.9210292983 | 14215.4304139331 | -7075.46051464915 | 14150.9210292983 | 10046 | 9.66078E-09 |
| Arsa | -11.10356649 | 0.330641351 | -0.254154405 | 0.25621182 | 0.527056313 | 8.04122E-05 | 0.5 | 0.002637143 | FALSE | 0 | relative convergence (4) | 6986.19279019576 | 7036.70217483059 | -3486.09639509788 | 6972.19279019576 | 10046 | 0.047597375 |
| Dnaja2 | -9.331728137 | 0.123396529 | -0.035809813 | 0.090006568 | 0.03296324 | 0.060583177 | 0.5 | 4.47905E-05 | FALSE | 0 | relative convergence (4) | 18437.923342077 | 18488.4327267118 | -9211.96167103851 | 18423.923342077 | 10046 | 0.003540269 |
| Kifc3 | -11.93172836 | 0.378681921 | -0.259896438 | -0.329959703 | 0.633197734 | 3.37634E-16 | 0.5 | 0.000138686 | FALSE | 0 | relative convergence (4) | 3755.14039156092 | 3805.64977619576 | -1870.57019578046 | 3741.14039156092 | 10046 | 0.007422951 |
| Ndrg4 | -10.96293366 | 0.360999269 | -0.514553721 | 0.220785366 | 0.470493691 | 5.47023E-05 | 0.5 | 0.00037481 | FALSE | 0 | relative convergence (4) | 7668.18718138599 | 7718.69656602083 | -3827.09359069299 | 7654.18718138599 | 10046 | 0.014769704 |
| Ripor1 | -10.58617802 | 0.364339784 | 0.093856643 | 0.356501567 | 0.113900669 | 0.091468656 | 0.5 | 0.002586953 | FALSE | 0 | relative convergence (4) | 9938.086218185 | 9988.59560281984 | -4962.0431090925 | 9924.086218185 | 10046 | 0.047048242 |
| Slc12a4 | -10.03557063 | 0.204067639 | -0.175880917 | 0.220944304 | 0.160879112 | 0.083533823 | 0.5 | 1.03882E-06 | FALSE | 0 | relative convergence (4) | 13182.0130008144 | 13232.5223854492 | -6584.0065004072 | 13168.0130008144 | 10046 | 0.000224172 |
| Ldlrap1 | -11.15420002 | 0.229560592 | 0.060221345 | -0.116361313 | 0.410983589 | 0.109889394 | 0.5 | 0.0023725 | FALSE | 0 | relative convergence (4) | 6372.29744046787 | 6422.80682510271 | -3179.14872023394 | 6358.29744046787 | 10046 | 0.044488461 |
| Maco1 | -9.666651882 | 0.256847869 | -0.147966009 | -0.014027634 | 0.44275813 | 0.100833477 | 0.5 | 0.000520286 | FALSE | 0 | relative convergence (4) | 15716.9064077192 | 15767.415792354 | -7851.45320385958 | 15702.9064077192 | 10046 | 0.017788246 |
| Gale | -13.45711305 | 0.750271895 | -0.438679622 | 0.659069045 | 1.758157283 | 5.08982E-05 | 0.5 | 0.000168851 | FALSE | 0 | relative convergence (4) | 1441.4214450813 | 1491.93082971614 | -713.710722540652 | 1427.4214450813 | 10046 | 0.008565431 |
| Hspg2 | -8.330600163 | 0.410254166 | -0.025776942 | -0.053066247 | 0.237050567 | 0.000434743 | 0.5 | 0.000202989 | FALSE | 0 | relative convergence (4) | 31791.4061520827 | 31841.9155367176 | -15888.7030760414 | 31777.4061520827 | 10046 | 0.009682926 |
| Txndc5 | -9.267065552 | 0.168087901 | -0.191830151 | 0.360834439 | 0.21504905 | 0.165504274 | 0.5 | 0.000742171 | FALSE | 0 | relative convergence (4) | 20125.5456043709 | 20176.0549890058 | -10055.7728021855 | 20111.5456043709 | 10046 | 0.022034183 |
| Diras2 | -12.13374601 | 0.482638734 | -0.431651153 | 0.09367896 | 4.138899214 | 3.60926E-05 | 0.5 | 0.000101895 | FALSE | 0 | relative convergence (4) | 3349.4731296934 | 3399.98251432823 | -1667.7365648467 | 3335.4731296934 | 10046 | 0.005844971 |
| Tspan17 | -11.17216256 | 0.448768686 | 0.061773681 | 0.481729511 | 0.297704163 | 5.62522E-05 | 0.5 | 0.000153156 | FALSE | 0 | relative convergence (4) | 7247.62431439564 | 7298.13369903048 | -3616.81215719782 | 7233.62431439564 | 10046 | 0.007920063 |
| Prelid1 | -9.853231184 | 0.486679903 | -0.124448697 | 0.413078765 | 0.094214112 | 0.000118105 | 0.5 | 0.002088616 | FALSE | 0 | relative convergence (4) | 15849.493891467 | 15900.0032761018 | -7917.74694573349 | 15835.493891467 | 10046 | 0.041424452 |
| Ankrd37 | -12.4148137 | 0.803125727 | -0.052235994 | -0.295765163 | 1.199564927 | 7.88795E-05 | 0.5 | 0.000571799 | FALSE | 0 | relative convergence (4) | 3257.52964512826 | 3308.0390297631 | -1621.76482256413 | 3243.52964512826 | 10046 | 0.018641759 |
| Acsl1 | -10.10941309 | 0.389646438 | 0.376016773 | 0.178716082 | 0.734071171 | 0.000119714 | 0.5 | 0.001869219 | FALSE | 0 | relative convergence (4) | 13398.3145494768 | 13448.8239341116 | -6692.15727473839 | 13384.3145494768 | 10046 | 0.038593779 |
| Cldn22 | -13.28883644 | 1.056580301 | 0.898301737 | 0.753162712 | 3.14719327 | 0.454696647 | 0.5 | 0.000356749 | FALSE | 0 | relative convergence (4) | 2272.65087765508 | 2323.16026228992 | -1129.32543882754 | 2258.65087765508 | 10046 | 0.014222863 |
| Tmem192 | -11.6066083 | 0.355667302 | -0.033443471 | 0.411612631 | 0.763126209 | 0.056882427 | 0.5 | 9.09721E-05 | FALSE | 0 | relative convergence (4) | 5118.35744208302 | 5168.86682671786 | -2552.17872104151 | 5104.35744208302 | 10046 | 0.005484378 |
| Naf1 | -10.94422888 | 0.552531181 | 0.382432713 | 0.450464028 | 0.408943875 | 0.197152303 | 0.5 | 0.000211813 | FALSE | 0 | relative convergence (4) | 8463.60628627927 | 8514.11567091411 | -4224.80314313963 | 8449.60628627927 | 10046 | 0.010033452 |
| Upf1 | -10.34908142 | 0.203499908 | 0.169256728 | 0.272727883 | 2.05728E-07 | 7.66475E-13 | 0.5 | 0.000398269 | FALSE | 0 | relative convergence (4) | NA | NA | NA | NA | 10046 | 0.01528425 |
| Comp | -12.49983278 | 1.365316698 | -0.214518937 | 0.789076637 | 24.7565435 | 0.060389015 | 0.5 | 0.002686641 | FALSE | 0 | relative convergence (4) | 3504.72521554461 | 3555.23460017945 | -1745.36260777231 | 3490.72521554461 | 10046 | 0.04824951 |
| Mapk3 | -9.893193963 | 0.253313934 | -0.00990963 | 0.304736326 | 0.113109869 | 0.104846596 | 0.5 | 1.1145E-05 | FALSE | 0 | relative convergence (4) | 14862.8092837577 | 14913.3186683925 | -7424.40464187884 | 14848.8092837577 | 10046 | 0.001377423 |
| Mvp | -10.2241067 | 0.217585658 | 0.029162311 | 0.517439395 | 0.181253137 | 0.099356959 | 0.5 | 2.49434E-06 | FALSE | 0 | relative convergence (4) | 12242.9355746939 | 12293.4449593287 | -6114.46778734693 | 12228.9355746939 | 10046 | 0.000429248 |
| Zfp688 | -12.10486528 | 0.490244191 | 0.342178514 | 0.318657041 | 0.292887655 | 8.34305E-37 | 0.5 | 0.00086244 | FALSE | 0 | relative convergence (4) | NA | NA | NA | NA | 10046 | 0.024075717 |
| Bckdk | -10.55695175 | 0.167394436 | 0.030988353 | 0.276884089 | 0.054265406 | 0.112376832 | 0.5 | 0.001305072 | FALSE | 0 | relative convergence (4) | 9731.16025429391 | 9781.66963892875 | -4858.58012714696 | 9717.16025429391 | 10046 | 0.031067795 |
| Armc5 | -11.43870841 | 0.402831315 | -0.150701182 | 0.389708316 | 0.216552149 | 0.075153339 | 0.5 | 4.04797E-07 | FALSE | 0 | relative convergence (4) | 5728.03783562498 | 5778.54722025981 | -2857.01891781249 | 5714.03783562498 | 10046 | 0.000101911 |
| Vasp | -10.05765098 | 0.593414404 | 0.400383594 | 0.48351861 | 0.330864149 | 0.256767469 | 0.5 | 0.001885211 | FALSE | 0 | relative convergence (4) | 15164.0738367435 | 15214.5832213784 | -7575.03691837177 | 15150.0738367435 | 10046 | 0.038593779 |
| Erf | -10.82069957 | 0.391554521 | 0.212380958 | 0.290784482 | 0.253867296 | 0.157360201 | 0.5 | 1.36848E-06 | FALSE | 0 | relative convergence (4) | 8337.94792692751 | 8388.45731156235 | -4161.97396346376 | 8323.94792692751 | 10046 | 0.000281886 |
| Egln2 | -10.38778537 | 0.186835393 | 0.103045523 | 0.284973268 | 3.9793E-07 | 0.029729118 | 0.5 | 9.1171E-05 | FALSE | 0 | relative convergence (4) | NA | NA | NA | NA | 10046 | 0.005484378 |
| Actn4 | -8.788502398 | 0.583994051 | 0.180712411 | 0.149366272 | 0.53146505 | 0.195418106 | 0.5 | 0.002408213 | FALSE | 0 | relative convergence (4) | 27641.9537350833 | 27692.4631197181 | -13813.9768675416 | 27627.9537350833 | 10046 | 0.045033366 |
| Paox | -12.27259728 | 0.392101719 | 0.07679287 | 0.437014839 | 0.5903538 | 0.180490799 | 0.5 | 0.001050867 | FALSE | 0 | relative convergence (4) | 3210.71484134868 | 3261.22422598351 | -1598.35742067434 | 3196.71484134868 | 10046 | 0.027264388 |
| Cracr2b | -12.09511211 | 0.508074918 | 0.491479805 | 0.359595047 | 0.796720844 | 0.285860637 | 0.5 | 5.92831E-07 | FALSE | 0 | relative convergence (4) | 3867.67838552618 | 3918.18777016102 | -1926.83919276309 | 3853.67838552618 | 10046 | 0.000144563 |
| Tspan4 | -10.29719825 | 0.249493521 | -0.257032108 | -0.038718309 | 0.436887438 | 0.214648245 | 0.5 | 0.000315585 | FALSE | 0 | relative convergence (4) | 11548.0188884016 | 11598.5282730364 | -5767.0094442008 | 11534.0188884016 | 10046 | 0.013201153 |
| Tssc4 | -10.93611086 | 0.385235326 | -0.092437922 | 0.385025424 | 0.078819273 | 0.100158363 | 0.5 | 2.67399E-10 | FALSE | 0 | relative convergence (4) | 7978.40796071398 | 8028.91734534882 | -3982.20398035699 | 7964.40796071398 | 10046 | 2.79638E-07 |
| Lamp1 | -7.838153089 | 0.146493305 | 0.016411357 | 0.233944397 | 0.063759597 | 0.071669575 | 0.5 | 0.001292147 | FALSE | 0 | relative convergence (4) | 35533.4896412214 | 35583.9990258563 | -17759.7448206107 | 35519.4896412214 | 10046 | 0.030955819 |
| Mir703 | -9.959946318 | 1.026275555 | -0.03082527 | 1.287152908 | 0.082525599 | 0.3949182 | 0.5 | 0.001239493 | FALSE | 0 | relative convergence (4) | 17596.8061504977 | 17647.3155351325 | -8791.40307524883 | 17582.8061504977 | 10046 | 0.030090896 |
| Crlf2 | -10.99841067 | 0.584495369 | -0.011635041 | 0.389908576 | 0.447748223 | 0.000290919 | 0.5 | 0.000669578 | FALSE | 0 | relative convergence (4) | 8356.88288217895 | 8407.39226681379 | -4171.44144108948 | 8342.88288217895 | 10046 | 0.020641536 |
| Chd4 | -8.636250866 | 0.124369145 | 0.098650721 | -0.093953769 | 0.19479611 | 0.08132205 | 0.5 | 6.79344E-07 | FALSE | 0 | relative convergence (4) | 25674.5314058473 | 25725.0407904821 | -12830.2657029236 | 25660.5314058473 | 10046 | 0.000153928 |

| | | | | | | | | | | | | | | | | | |
|---|---|---|---|---|---|---|---|---|---|---|---|---|---|---|---|---|---|
| Gapdh | -8.787478367 | 0.877255719 | 0.888997613 | 0.665703495 | 0.334179943 | 0.462090452 | 0.5 | 9.23002E-05 | FALSE | 0 | relative convergence (4) | 30109.5524502607 | 30160.0618348956 | -15047.7762251304 | 30095.5524502607 | 10046 | 0.0055036 |
| Ltbr | -10.58559766 | 0.246063882 | 0.010312044 | 0.216071275 | 0.023061404 | 0.150027083 | 0.5 | 0.000276419 | FALSE | 0 | relative convergence (4) | 9706.49095455306 | 9757.00033918789 | -4846.24547727653 | 9692.49095455306 | 10046 | 0.011976205 |
| Tnfrsf1a | -9.832355894 | 0.220169494 | -0.022697823 | 0.43106895 | 0.189335764 | 0.158891308 | 0.5 | 9.96699E-09 | FALSE | 0 | relative convergence (4) | 15253.3088856469 | 15303.8182702818 | -7619.65444282347 | 15239.3088856469 | 10046 | 5.27123E-06 |
| Uchl1 | -10.26355932 | 0.869282029 | 0.668762918 | 0.820633946 | 0.955704885 | 0.334212572 | 0.5 | 0.002276697 | FALSE | 0 | relative convergence (4) | 15261.6844542785 | 15312.1938389133 | -7623.84222713925 | 15247.6844542785 | 10046 | 0.043410518 |
| Commd8 | -10.37324512 | 0.188325114 | 0.024844933 | 0.191778539 | 0.040103282 | 0.014375051 | 0.5 | 6.25372E-05 | FALSE | 0 | relative convergence (4) | 10686.5591524295 | 10737.0685370644 | -5336.27957621476 | 10672.5591524295 | 10046 | 0.004309979 |
| Gm19610 | -13.5954561 | 1.046184128 | 0.608561655 | 0.262127571 | 3.362330896 | 0.137024946 | 0.5 | 0.000699893 | FALSE | 0 | relative convergence (4) | 1463.66447460745 | 1514.17385924229 | -724.832237303727 | 1449.66447460745 | 10046 | 0.021334195 |
| Parm1 | -11.2401807 | 0.753261296 | 0.738572659 | 0.455321783 | 1.771729666 | 0.000179461 | 0.5 | 0.00158363 | FALSE | 0 | relative convergence (4) | 7979.3517033145 | 8029.86108794934 | -3982.67585165725 | 7965.3517033145 | 10046 | 0.035273898 |
| Art3 | -12.43267382 | 0.468184466 | 0.606642534 | -0.00221172 | 8.518855056 | 0.090475796 | 0.5 | 0.002065632 | FALSE | 0 | relative convergence (4) | NA | NA | NA | NA | 10046 | 0.041297449 |
| Epas1 | -10.27294138 | 0.531259862 | 0.473163493 | 0.380470922 | 1.840227135 | 0.000164441 | 0.5 | 0.002138214 | FALSE | 0 | relative convergence (4) | 13358.6162351897 | 13409.1256198245 | -6672.30811759483 | 13344.6162351897 | 10046 | 0.04186121 |
| Ccl7 | -12.82860794 | 1.278086363 | 0.032366377 | -0.467301227 | 62.1026428 | 0.850947671 | 0.5 | 0.001792302 | FALSE | 0 | relative convergence (4) | 1728.28343812382 | 1778.79282275866 | -857.141719061909 | 1714.28343812382 | 10046 | 0.037841758 |
| Prr15 | -13.37023546 | 0.910732322 | 0.465305113 | 0.460055657 | 1.293056877 | 6.20031E-05 | 0.5 | 1.17014E-05 | FALSE | 0 | relative convergence (4) | 1636.91036762392 | 1687.41975225876 | -811.455183811962 | 1622.91036762392 | 10046 | 0.001407788 |
| Gars | -10.01335936 | 0.137974366 | 0.101483716 | 0.23984043 | 0.237277766 | 0.140387143 | 0.5 | 0.00179181 | FALSE | 0 | relative convergence (4) | 13191.9125975379 | 13242.4219821727 | -6588.95629876895 | 13177.9125975379 | 10046 | 0.037841758 |
| Eif3b | -9.711871816 | 0.131045362 | -0.075536666 | 0.009170573 | 0.048585407 | 0.039011185 | 0.5 | 0.000336237 | FALSE | 0 | relative convergence (4) | 15258.3042011276 | 15308.8135857624 | -7622.15210056378 | 15244.3042011276 | 10046 | 0.013645024 |
| Arpc1b | -8.183163561 | 0.225885897 | 0.076401306 | 0.411088039 | 0.284031406 | 0.121377548 | 0.5 | 1.67802E-05 | FALSE | 0 | relative convergence (4) | 33812.2708238075 | 33862.7802084423 | -16899.1354119037 | 33798.2708238075 | 10046 | 0.001796274 |
| 2410022M11Rik | -12.38420602 | 0.51534222 | 0.393359192 | 0.507769726 | 1.33299E-06 | 5.13558E-11 | 0.5 | 1.198E-05 | FALSE | 0 | relative convergence (4) | NA | NA | NA | NA | 10046 | 0.001416239 |
| Spata13 | -11.65996864 | 0.758213889 | 0.490934329 | 0.318449094 | 0.965018605 | 0.000538637 | 0.5 | 0.001195652 | FALSE | 0 | relative convergence (4) | 5642.52435030338 | 5693.03373493822 | -2814.26217515169 | 5628.52435030338 | 10046 | 0.029394014 |
| Ephx2 | -12.36563602 | 0.728039651 | 0.449243571 | 0.373734509 | 5.159548826 | 0.35668578 | 0.5 | 9.51268E-08 | FALSE | 0 | relative convergence (4) | 3220.6273115013 | 3271.13669613614 | -1603.31365575065 | 3206.6273115013 | 10046 | 3.00756E-05 |
| Polr3d | -10.95287216 | 0.226847201 | 0.073860587 | 0.096061684 | 0.354827383 | 0.070806285 | 0.5 | 0.000593827 | FALSE | 0 | relative convergence (4) | 7473.12316931135 | 7523.63255394619 | -3729.56158465567 | 7459.12316931135 | 10046 | 0.019130509 |
| Timm23 | -10.04696208 | 0.14404708 | -0.026866737 | 0.230196879 | 0.134298217 | 3.94685E-05 | 0.5 | 0.000449548 | FALSE | 0 | relative convergence (4) | 12875.7001148888 | 12926.2094995236 | -6430.85005744439 | 12861.7001148888 | 10046 | 0.016254263 |
| Lrp10 | -9.690358881 | 0.131875369 | 0.089702995 | 0.244116258 | 0.076756065 | 0.136716922 | 0.5 | 0.000221182 | FALSE | 0 | relative convergence (4) | 15827.1189079191 | 15877.628292554 | -7906.55945395957 | 15813.1189079191 | 10046 | 0.010240336 |
| H2aj | -9.541672449 | 0.321459562 | 0.127858181 | 0.26950263 | 0.252541527 | 8.53946E-05 | 0.5 | 0.001849674 | FALSE | 0 | relative convergence (4) | 18687.7719044844 | 18738.2812891193 | -9336.88595224221 | 18673.7719044844 | 10046 | 0.038508909 |
| Ppp1r12c | -9.779546591 | 0.474075089 | 0.231299777 | 0.245445674 | 0.317945366 | 0.000521501 | 0.5 | 0.000522051 | FALSE | 0 | relative convergence (4) | 16541.9909054036 | 16592.5002900384 | -8263.99545270179 | 16527.9909054036 | 10046 | 0.017788246 |
| Srebf1 | -10.95244545 | 0.300681434 | 0.202447248 | 0.573594029 | 0.106744524 | 8.05832E-05 | 0.5 | 0.00173648 | FALSE | 0 | relative convergence (4) | 8079.21587394765 | 8129.72525858249 | -4032.60793697383 | 8065.21587394765 | 10046 | 0.037353561 |
| Flii | -9.672265203 | 0.218441025 | 0.248578771 | 0.37174962 | 0.058088979 | 0.144265232 | 0.5 | 0.000426482 | FALSE | 0 | relative convergence (4) | 16072.3341032459 | 16122.8434878808 | -8029.16705162297 | 16058.3341032459 | 10046 | 0.015798445 |
| Map2k3 | -10.04149397 | 0.2212779 | -0.383278722 | -0.183068796 | 0.356525482 | 0.165858982 | 0.5 | 0.002470157 | FALSE | 0 | relative convergence (4) | 12336.1666215302 | 12386.676006165 | -6161.08331076509 | 12322.1666215302 | 10046 | 0.045689508 |
| Vamp2 | -10.08186862 | 0.38356034 | 0.151013228 | 0.32950131 | 0.082765491 | 6.6696E-05 | 0.5 | 3.92743E-05 | FALSE | 0 | relative convergence (4) | 13472.81961042 | 13523.3289950548 | -6729.40980521 | 13458.81961042 | 10046 | 0.003235961 |
| Crybb3 | -12.3168035 | 0.419692081 | -0.155568084 | 0.196786994 | 2.499248966 | 0.194229221 | 0.5 | 0.001409917 | FALSE | 0 | relative convergence (4) | 2913.16215760431 | 2963.67154223915 | -1449.58107880216 | 2899.16215760431 | 10046 | 0.032709582 |
| 2900026A02Rik | -9.848930669 | 0.511957221 | 0.112968129 | 0.193871169 | 0.443203812 | 0.058574028 | 0.5 | 3.44323E-07 | FALSE | 0 | relative convergence (4) | 15343.3498953198 | 15393.8592799547 | -7664.67494765992 | 15329.3498953198 | 10046 | 8.9689E-05 |
| Kctd10 | -9.623868417 | 0.435257738 | 0.27367965 | 0.379167562 | 0.277886155 | 0.120683217 | 0.5 | 0.002122084 | FALSE | 0 | relative convergence (4) | 17952.3238282918 | 18002.8332129267 | -8969.16191414592 | 17938.3238282918 | 10046 | 0.041690366 |
| Srsf9 | -10.52110744 | 0.609219483 | 0.135428399 | 0.74971003 | 1.24621E-05 | 1.24315E-38 | 0.5 | 0.001795072 | FALSE | 1 | singular convergence (7) | NA | NA | NA | NA | 10046 | 0.037841758 |
| Fbxo21 | -11.07733995 | 0.264107558 | -0.025958385 | 0.025614194 | 0.294801572 | 0.167755704 | 0.5 | 0.000228438 | FALSE | 0 | relative convergence (4) | 6806.83175259206 | 6857.3411372269 | -3396.41587629603 | 6792.83175259206 | 10046 | 0.010456617 |
| Ddx54 | -10.63840476 | 0.272497972 | 0.057277359 | 0.176101221 | 0.100057642 | 0.018309245 | 0.5 | 3.31935E-07 | FALSE | 0 | relative convergence (4) | 9154.04462617899 | 9204.55401081383 | -4570.02231308949 | 9140.04462617899 | 10046 | 8.84835E-05 |
| Abi3 | -13.52670322 | 1.029018466 | 0.808920852 | 0.707062651 | 2.711493009 | 0.269790751 | 0.5 | 0.000134295 | FALSE | 0 | relative convergence (4) | 1618.92985324395 | 1669.43923787879 | -802.464926621975 | 1604.92985324395 | 10046 | 0.007255479 |
| Cdk5rap3 | -10.31770956 | 0.260155787 | 0.145301807 | 0.434595996 | 0.097622052 | 0.183776942 | 0.5 | 1.08565E-08 | FALSE | 0 | relative convergence (4) | 11864.2544598271 | 11914.7638444619 | -5925.12722991354 | 11850.2544598271 | 10046 | 5.27123E-06 |
| Tcap | -10.54073245 | 1.359297541 | 1.898153794 | 1.043149051 | 3.983250391 | 0.829011646 | 0.5 | 0.00025618 | FALSE | 0 | relative convergence (4) | 15401.0150127608 | 15451.5243973957 | -7693.50750638042 | 15387.0150127608 | 10046 | 0.01129413 |
| Nr1d1 | -9.833144526 | 0.89227344 | -0.212142139 | -1.076758745 | 0.495051639 | 0.000169235 | 0.5 | 0.00196386 | FALSE | 0 | relative convergence (4) | 16601.3918447376 | 16651.9012293724 | -8293.69592236879 | 16587.3918447376 | 10046 | 0.039671131 |
| Naglu | -10.76978437 | 0.282660204 | 0.098848691 | 0.276715402 | 0.096128185 | 0.120697738 | 0.5 | 5.98828E-07 | FALSE | 0 | relative convergence (4) | 8773.64302802384 | 8824.15241265868 | -4379.82151401192 | 8759.64302802384 | 10046 | 0.000144563 |
| Ccdc62 | -11.86861413 | 0.3629754 | 0.368551847 | 0.532444032 | 1.966678092 | 0.075962255 | 0.5 | 0.00063345 | FALSE | 0 | relative convergence (4) | 4332.86813557548 | 4383.37752021031 | -2159.43406778774 | 4318.86813557548 | 10046 | 0.020027345 |
| Ran | -10.09929312 | 0.618225759 | 0.034640246 | 0.785436481 | 0.05348477 | 0.232573077 | 0.5 | 0.002144261 | FALSE | 0 | relative convergence (4) | 14773.0042259026 | 14823.5136105374 | -7379.50211295129 | 14759.0042259026 | 10046 | 0.041883943 |
| Stx1a | -11.10751774 | 0.561874055 | 0.725572895 | 0.610030301 | 0.75230811 | 0.000132823 | 0.5 | 0.001015492 | FALSE | 0 | relative convergence (4) | 8175.97096873486 | 8226.4803533697 | -4080.98548436743 | 8161.97096873486 | 10046 | 0.026600416 |
| Hip1 | -8.896363555 | 0.520380224 | 0.445728736 | 0.374324819 | 0.770833745 | 0.128934674 | 0.5 | 2.25651E-05 | FALSE | 0 | relative convergence (4) | 26911.2983876331 | 26961.8077722679 | -13448.6491938165 | 26897.2983876331 | 10046 | 0.002205153 |
| Plod3 | -10.31787061 | 0.174028242 | -0.131423138 | 0.329390391 | 0.055056191 | 4.05537E-05 | 0.5 | 0.001651172 | FALSE | 0 | relative convergence (4) | 11174.5367737654 | 11225.0461584003 | -5580.26838688272 | 11160.5367737654 | 10046 | 0.035925543 |
| Serpine1 | -10.82794314 | 1.290783474 | 0.489366436 | 0.445244481 | 3.397512103 | 0.000532375 | 0.5 | 0.002771251 | FALSE | 0 | relative convergence (4) | 11132.9173664974 | 11183.4267511322 | -5559.45868324869 | 11118.9173664974 | 10046 | 0.049377672 |
| P2rx1 | -13.09993373 | 1.147990461 | 0.38952785 | 0.913746605 | 7.768853589 | 0.227387712 | 0.5 | 6.24536E-05 | FALSE | 0 | relative convergence (4) | 2460.45823801992 | 2510.96762265475 | -1223.22911900996 | 2446.45823801992 | 10046 | 0.004309979 |
| Coro6 | -13.55849295 | 1.263597301 | 1.209259367 | 0.875865932 | 2.064465794 | 6.88448E-05 | 0.5 | 1.09163E-05 | FALSE | 0 | relative convergence (4) | 1929.62652371624 | 1980.13590835108 | -957.813261858121 | 1915.62652371624 | 10046 | 0.001374134 |
| Rpl23a | -8.620879617 | 0.828927397 | 0.074679122 | 0.893553737 | 0.104421114 | 0.000397729 | 0.5 | 0.002331671 | FALSE | 0 | relative convergence (4) | 30776.5393151928 | 30827.0486998276 | -15381.2696575964 | 30762.5393151928 | 10046 | 0.044087718 |
| Cdc16 | -10.10123136 | 0.146663095 | -0.027649213 | 0.146384722 | 0.159457492 | 0.110146118 | 0.5 | 0.000823471 | FALSE | 0 | relative convergence (4) | 12712.2582374476 | 12762.7676220825 | -6349.12911872382 | 12698.2582374476 | 10046 | 0.023420675 |
| Snord13 | -13.83418687 | 2.051972243 | 0.051805925 | 0.487732372 | 1.24031613 | 1.359911891 | 0.5 | 0.00010924 | FALSE | 0 | relative convergence (4) | 2010.63221760859 | 2061.14160224343 | -998.316108804294 | 1996.63221760859 | 10046 | 0.00616233 |
| Fat1 | -9.161256158 | 0.359142735 | 0.146198099 | 0.149491225 | 0.461754543 | 0.000141413 | 0.5 | 0.000149166 | FALSE | 0 | relative convergence (4) | 22254.3949452823 | 22304.9043299172 | -11120.1974726412 | 22240.3949452823 | 10046 | 0.00786014 |
| Stk19 | -10.96559492 | 0.344701805 | 0.232813751 | 0.304862888 | 0.134360283 | 5.51023E-07 | 0.5 | 1.55698E-08 | FALSE | 0 | relative convergence (4) | 7885.70371010155 | 7936.21309473639 | -3935.85185505077 | 7871.70371010155 | 10046 | 6.82813E-06 |
| Skiv2l | -10.91412767 | 0.283216353 | 0.149710866 | 0.318158455 | 0.017955036 | 0.084266315 | 0.5 | 0.000520593 | FALSE | 0 | relative convergence (4) | 7932.51318957026 | 7983.0225742051 | -3959.25659478513 | 7918.51318957026 | 10046 | 0.017788246 |
| Vars | -10.88921417 | 0.335036734 | -0.063174605 | 0.475513342 | 0.165580232 | 5.90387E-05 | 0.5 | 0.000390902 | FALSE | 0 | relative convergence (4) | 8277.97931779313 | 8328.48870242797 | -4131.98965889656 | 8263.97931779313 | 10046 | 0.015227258 |
| Nfkbil1 | -11.53971809 | 0.376946471 | -0.00902475 | 0.364956844 | 0.058208806 | 1.57204E-05 | 0.5 | 2.20532E-06 | FALSE | 0 | relative convergence (4) | 5246.38807843968 | 5296.89746307452 | -2616.19403921984 | 5232.38807843968 | 10046 | 0.000389367 |
| Mrps18b | -11.52627545 | 0.573336554 | 0.109757487 | 0.497452917 | 1.33176E-06 | 0.083689031 | 0.5 | 0.001209831 | FALSE | 0 | relative convergence (4) | 5726.33717321365 | 5776.84655784849 | -2856.16858660682 | 5712.33717321365 | 10046 | 0.029635421 |
| Gnl1 | -10.7122239 | 0.286673937 | -0.006430087 | 0.151380557 | 0.002372456 | 0.00044904 | 0.5 | 0.000498333 | FALSE | 0 | relative convergence (4) | 8787.03542659886 | 8837.54481123369 | -4386.51771329943 | 8773.03542659886 | 10046 | 0.017396216 |
| Klhdc3 | -10.85849923 | 0.327819817 | 0.083824245 | 0.140443656 | 0.117764847 | 0.151526142 | 0.5 | 0.001340923 | FALSE | 0 | relative convergence (4) | 8222.87592414248 | 8273.38530877732 | -4104.43796207124 | 8208.87592414248 | 10046 | 0.031466741 |
| Atoh8 | -11.57670609 | 0.705983527 | 0.896732513 | 0.049597262 | 1.34287996 | 0.000333422 | 0.5 | 0.002232857 | FALSE | 0 | relative convergence (4) | 6276.36563999965 | 6326.87502463449 | -3131.18281999983 | 6262.36563999965 | 10046 | 0.043118882 |
| Vamp8 | -9.506084513 | 0.243372385 | 0.215657941 | 0.140107407 | 0.157158386 | 7.41154E-05 | 0.5 | 0.00223019 | FALSE | 0 | relative convergence (4) | 17963.7552256061 | 18014.2646102409 | -8974.87761280303 | 17949.7552256061 | 10046 | 0.043118882 |
| Tmsb10 | -7.665070535 | 0.280163689 | 0.084798686 | 0.53892713 | 0.925947992 | 0.235784625 | 0.5 | 0.000129698 | FALSE | 0 | relative convergence (4) | 43287.6450933171 | 43338.154477952 | -21636.8225466586 | 43273.6450933171 | 10046 | 0.007052958 |
| Dok1 | -11.21954643 | 0.347229149 | -0.06115148 | 0.307541858 | 0.111476973 | 0.081228218 | 0.5 | 6.42897E-07 | FALSE | 0 | relative convergence (4) | 6551.4234210104 | 6601.93280564523 | -3268.7117105052 | 6537.4234210104 | 10046 | 0.000149359 |
| Pcgf1 | -12.27087337 | 0.397442499 | 0.301164483 | 0.544864815 | 5.01593E-07 | 0.168996101 | 0.5 | 0.000618339 | FALSE | 1 | false convergence (8) | 3269.1390231971 | 3319.64840783194 | -1627.56951159855 | 3255.1390231971 | 10046 | 0.019646346 |
| Dctn1 | -10.10223325 | 0.138603987 | 0.270375345 | 0.037144732 | 0.077921542 | 0.122725697 | 0.5 | 0.001287174 | FALSE | 0 | relative convergence (4) | 12148.2985465145 | 12198.8079311493 | -6067.14927325726 | 12134.2985465145 | 10046 | 0.030917201 |
| Smyd5 | -12.12156834 | 0.380472334 | -0.491379708 | 0.235830262 | 0.121291387 | 0.060343785 | 0.5 | 0.000179571 | FALSE | 0 | relative convergence (4) | NA | NA | NA | NA | 10046 | 0.00887169 |
| Lama5 | -11.47345739 | 1.458422466 | 0.997438376 | 0.725990119 | 1.153615219 | 0.00037675 | 0.5 | 0.000986686 | FALSE | 0 | relative convergence (4) | 9548.64559080027 | 9599.15497543511 | -4767.32279540014 | 9534.64559080027 | 10046 | 0.026250494 |
| Fry | -8.716249108 | 0.42064256 | 0.463854356 | 0.240332808 | 1.022072472 | 0.166450953 | 0.5 | 0.000491626 | FALSE | 0 | relative convergence (4) | 28964.7634074221 | 29015.272792057 | -14475.3817037111 | 28950.7634074221 | 10046 | 0.017242321 |
| Asns | -12.46424515 | 0.453330469 | 0.214924559 | 0.406237059 | 4.560038882 | 0.229811169 | 0.5 | 0.002767999 | FALSE | 0 | relative convergence (4) | 2799.12397080715 | 2849.63335544199 | -1392.56198540358 | 2785.12397080715 | 10046 | 0.049377672 |
| Ica1 | -12.20983156 | 0.534786172 | 0.370395299 | 0.237873449 | 4.215267222 | 0.25779682 | 0.5 | 2.94093E-05 | FALSE | 0 | relative convergence (4) | NA | NA | NA | NA | 10046 | 0.002613197 |
| Wnt2 | -10.96681907 | 0.40427437 | -0.17387736 | 0.807400602 | 8.468017123 | 0.349603541 | 0.5 | 8.55072E-05 | FALSE | 0 | relative convergence (4) | 7013.22033656304 | 7063.72972119788 | -3499.61016828152 | 6999.22033656304 | 10046 | 0.005308087 |
| Ptprz1 | -10.14265935 | 1.253473292 | 0.964848577 | 0.85793364 | 4.317301123 | 0.000249667 | 0.5 | 0.000306521 | FALSE | 0 | relative convergence (4) | 18056.8563682199 | 18107.3657528547 | -9021.42818410994 | 18042.8563682199 | 10046 | 0.012941456 |
| Bpgm | -11.35806574 | 0.28532391 | -0.013454472 | 0.153617718 | 0.450194648 | 0.000140613 | 0.5 | 0.000189717 | FALSE | 0 | relative convergence (4) | NA | NA | NA | NA | 10046 | 0.009244465 |
| Mgat1 | -10.24941702 | 0.352080167 | 0.095420496 | 0.375685723 | 0.005241141 | 0.098876616 | 0.5 | 2.93991E-05 | FALSE | 0 | relative convergence (4) | 12294.4121761845 | 12344.9215608193 | -6140.20608809223 | 12280.4121761845 | 10046 | 0.002613197 |
| Ube2b | -8.655050685 | 0.091284306 | 0.007374281 | -0.012910514 | 0.045393003 | 0.108006098 | 0.5 | 3.74968E-05 | FALSE | 0 | relative convergence (4) | NA | NA | NA | NA | 10046 | 0.003108345 |
| Trim3 | -10.65884788 | 0.181501715 | -0.034258838 | 0.146226166 | 0.155819932 | 0.048310032 | 0.5 | 0.001226254 | FALSE | 0 | relative convergence (4) | 8842.92646382144 | 8893.43584845628 | -4414.46323191072 | 8828.92646382144 | 10046 | 0.029876207 |
| Arfip2 | -10.76204549 | 0.219520892 | 0.149866991 | 0.47033994 | 0.032664236 | 0.173653601 | 0.5 | 8.87831E-05 | FALSE | 0 | relative convergence (4) | 8978.22508412737 | 9028.73446876221 | -4482.11254206369 | 8964.22508412737 | 10046 | 0.005436198 |
| Lmo1 | -11.5624562 | 0.267595478 | -0.082859218 | 0.411119779 | 1.298224973 | 0.000324136 | 0.5 | 0.002554964 | FALSE | 0 | relative convergence (4) | NA | NA | NA | NA | 10046 | 0.046686463 |
| Akip1 | -11.86431665 | 0.312243125 | 0.074074356 | 0.394125741 | 0.061292062 | 0.000265914 | 0.5 | 0.000998503 | FALSE | 0 | relative convergence (4) | 4236.88188332059 | 4287.39126795543 | -2111.44094166029 | 4222.88188332059 | 10046 | 0.026286506 |
| Notch3 | -11.28091731 | 0.776845878 | 0.850954658 | 0.097125781 | 3.030649502 | 0.145665994 | 0.5 | 0.000588203 | FALSE | 0 | relative convergence (4) | 7823.83843117447 | 7874.3478158093 | -3904.91921558723 | 7809.83843117447 | 10046 | 0.019039576 |
| Ctps | -10.81539532 | 0.363335167 | 0.091434609 | 0.372264432 | 0.864377242 | 0.000146309 | 0.5 | 9.40688E-05 | FALSE | 0 | relative convergence (4) | 8561.17983330803 | 8611.68921794287 | -4273.58991665402 | 8547.17983330803 | 10046 | 0.005564624 |
| Gja4 | -13.47663142 | 1.234112796 | 0.740736341 | 0.633686277 | 5.961680167 | 8.2268E-05 | 0.5 | 6.19286E-05 | FALSE | 0 | relative convergence (4) | 1819.79189379758 | 1870.30127843242 | -902.89594689879 | 1805.79189379758 | 10046 | 0.004309979 |
| Wwc1 | -12.749408 | 0.506463899 | 0.142645392 | 0.238695087 | 6.411127638 | 0.180226726 | 0.5 | 0.002140018 | FALSE | 0 | relative convergence (4) | 2149.11862718226 | 2199.6280118171 | -1067.55931359113 | 2135.11862718226 | 10046 | 0.04186121 |
| Adam33 | -10.29495535 | 0.270018572 | 0.404910247 | 0.412817726 | 0.871473036 | 0.10921572 | 0.5 | 0.001791205 | FALSE | 0 | relative convergence (4) | 12032.9549167753 | 12083.4643014101 | -6009.47745838764 | 12018.9549167753 | 10046 | 0.037841758 |
| Cds2 | -10.18826886 | 0.455034245 | 0.476543762 | 0.444085286 | 0.224006387 | 0.043515693 | 0.5 | 0.000192178 | FALSE | 0 | relative convergence (4) | 13400.6653824079 | 13451.1747670428 | -6693.33269120396 | 13386.6653824079 | 10046 | 0.009330923 |
| Pygb | -9.629973128 | 0.251384631 | 0.13352542 | 0.236786418 | 0.272050941 | 0.129248733 | 0.5 | 0.000231342 | FALSE | 0 | relative convergence (4) | 17072.4302314061 | 17122.939616041 | -8529.21511570306 | 17058.4302314061 | 10046 | 0.010531918 |
| Angpt4 | -12.49694209 | 0.73172692 | -0.239607329 | 1.257455172 | 15.54514104 | 0.418898398 | 0.5 | 0.001092639 | FALSE | 0 | relative convergence (4) | 3125.62846966174 | 3176.13785429657 | -1555.81423483087 | 3111.62846966174 | 10046 | 0.027974435 |
| H13 | -9.347786667 | 0.163539713 | -0.156032049 | 0.236358783 | 0.132275365 | 0.075493847 | 0.5 | 2.0722E-07 | FALSE | 0 | relative convergence (4) | NA | NA | NA | NA | 10046 | 6.06518E-05 |
| Cox4i2 | -11.60568978 | 1.259382311 | 1.019526314 | 0.561116237 | 2.050920987 | 0.306434306 | 0.5 | 0.001818882 | FALSE | 0 | relative convergence (4) | 8144.62362374773 | 8195.13300838257 | -4065.31181187386 | 8130.62362374773 | 10046 | 0.038004845 |
| Dlgap4 | -8.949223347 | 0.142802544 | 0.145717814 | 0.149820649 | 0.159710862 | 0.121976968 | 0.5 | 0.000622415 | FALSE | 0 | relative convergence (4) | 22615.5730376327 | 22666.0824222676 | -11300.7865188164 | 22601.5730376327 | 10046 | 0.019724306 |
| Serinc3 | -8.525137798 | 0.222319563 | -0.0774466 | 0.198761824 | 0.131544265 | 0.037907652 | 0.5 | 0.002139009 | FALSE | 0 | relative convergence (4) | 27966.2125830621 | 28016.7219676969 | -13976.1062915311 | 27952.2125830621 | 10046 | 0.04186121 |
| Pkig | -9.719155695 | 0.439359183 | 0.351582058 | 0.389485852 | 0.399285181 | 0.260003995 | 0.5 | 0.002096361 | FALSE | 0 | relative convergence (4) | 17659.4337898068 | 17709.9431744417 | -8822.71689490341 | 17645.4337898068 | 10046 | 0.041424452 |
| Ccn5 | -10.26354579 | 0.970803523 | 0.050635781 | 0.94813542 | 5.935735126 | 0.456018353 | 0.5 | 2.08331E-05 | FALSE | 0 | relative convergence (4) | 13939.6728797782 | 13990.182264413 | -6962.83643988908 | 13925.6728797782 | 10046 | 0.002113628 |
| Ctsa | -9.541211681 | 0.190667424 | -0.165218358 | 0.099569919 | 0.150945313 | 0.108334935 | 0.5 | 0.000999204 | FALSE | 0 | relative convergence (4) | 17323.5325931557 | 17374.0419777906 | -8654.76629657786 | 17309.5325931557 | 10046 | 0.026286506 |
| Ntpcr | -11.40470765 | 0.328114335 | -0.011774591 | 0.210290967 | 0.15692399 | 0.058870418 | 0.5 | 2.70588E-05 | FALSE | 0 | relative convergence (4) | 5666.35197297865 | 5716.86135761349 | -2826.17598648932 | 5652.35197297865 | 10046 | 0.002469284 |
| Gm10709 | -13.46237533 | 1.431883592 | -0.990649665 | -0.015403165 | 9.27899E-08 | 0.490005294 | 0.5 | 0.002196811 | FALSE | 1 | false convergence (8) | NA | NA | NA | NA | 10046 | 0.042707071 |
| Keap1 | -10.0702943 | 0.147313322 | 0.0063212 | 0.097502787 | 0.110012837 | 0.03895325 | 0.5 | 0.000446798 | FALSE | 0 | relative convergence (4) | 12593.3466539141 | 12643.856038549 | -6289.67332695706 | 12579.3466539141 | 10046 | 0.016214151 |
| Dock6 | -10.3456217 | 0.31909841 | 0.569097879 | 0.380296694 | 0.552448644 | 0.187329425 | 0.5 | 0.001653556 | FALSE | 0 | relative convergence (4) | 12005.2617773036 | 12055.7711619384 | -5995.63088865178 | 11991.2617773036 | 10046 | 0.035925543 |
| Prkcsh | -10.03985066 | 0.247036409 | 0.175100665 | 0.410064475 | 6.08214E-07 | 0.117223172 | 0.5 | 0.000839233 | FALSE | 1 | false convergence (8) | 13856.2909860401 | 13906.8003706749 | -6921.14549302005 | 13842.2909860401 | 10046 | 0.0236219 |
| Tmem203 | -11.40584352 | 0.321351575 | 0.110952882 | 0.18862884 | 3.31788E-07 | 0.167341701 | 0.5 | 2.28502E-05 | FALSE | 0 | relative convergence (4) | NA | NA | NA | NA | 10046 | 0.002205153 |
| Rxra | -10.54415549 | 0.220402781 | 0.167143182 | 0.031050473 | 5.688E-07 | 1.95037E-05 | 0.5 | 0.000858054 | FALSE | 1 | false convergence (8) | 9468.31071106657 | 9518.8200957014 | -4727.15535553328 | 9454.31071106657 | 10046 | 0.024002558 |
| Atp1a1 | -9.185180037 | 0.370244756 | 0.243019744 | 0.249909777 | 0.295936389 | 0.165543176 | 0.5 | 0.001567693 | FALSE | 0 | relative convergence (4) | 21751.8092026022 | 21802.3185872371 | -10868.9046013011 | 21737.8092026022 | 10046 | 0.034996369 |
| Kcnc4 | -12.4483923 | 0.903488258 | 1.00004323 | 0.540613065 | 2.81930987 | 0.372540622 | 0.5 | 0.00262535 | FALSE | 0 | relative convergence (4) | 3582.86567738101 | 3633.37506201585 | -1784.43283869051 | 3568.86567738101 | 10046 | 0.047525482 |
| Cyb561d1 | -13.11614263 | 0.585507932 | 0.008969052 | 0.629142942 | 0.283980009 | 0.088285855 | 0.5 | 0.00085048 | FALSE | 0 | relative convergence (4) | 1815.1320772485 | 1865.64146188333 | -900.566038624248 | 1801.1320772485 | 10046 | 0.023839755 |
| Sulf1 | -7.846357892 | 0.225809669 | -0.129568412 | 0.129436646 | 0.68186405 | 0.076708098 | 0.5 | 9.86866E-06 | FALSE | 0 | relative convergence (4) | 39154.4052691208 | 39204.9146537556 | -19570.2026345604 | 39140.4052691208 | 10046 | 0.001277757 |
| Sbspon | -13.9191218 | 0.986450933 | 0.800209482 | 1.195512144 | 1.97026877 | 7.34083E-05 | 0.5 | 0.001678606 | FALSE | 0 | relative convergence (4) | 1265.64763187926 | 1316.1570165141 | -625.82381593963 | 1251.64763187926 | 10046 | 0.036324405 |
| Col19a1 | -12.62392703 | 0.713553336 | 0.90992995 | 0.809929806 | 7.195279658 | 6.8172E-05 | 0.5 | 0.000771026 | FALSE | 0 | relative convergence (4) | 2948.65622199589 | 2999.16560663073 | -1467.32811099794 | 2934.65622199589 | 10046 | 0.022445616 |
| Gm28439 | -17.18338217 | 3.852395168 | 4.068546398 | 2.585293996 | 0.60763216 | 2.32478059 | 0.5 | 1.12663E-09 | FALSE | 0 | relative convergence (4) | 1893.57862945352 | 1944.08801408835 | -939.789314726758 | 1879.57862945352 | 10046 | 1.0211E-06 |
| 4933424G06Rik | -12.85949105 | 1.35813836 | 1.334361696 | 0.643617555 | 0.426644627 | 0.804606881 | 0.5 | 2.52561E-13 | FALSE | 0 | relative convergence (4) | 3642.80459147281 | 3693.31397610765 | -1814.40229573641 | 3628.80459147281 | 10046 | 5.72261E-10 |
| Rpl31 | -8.212286141 | 0.342004322 | 0.10731323 | 0.075625244 | 0.068788173 | 0.000106863 | 0.5 | 0.000180109 | FALSE | 0 | relative convergence (4) | 31765.4145096116 | 31815.9238942465 | -15875.7072548058 | 31751.4145096116 | 10046 | 0.00887169 |
| Rnf152 | -10.72955672 | 0.351286512 | 0.170274312 | 0.168074391 | 2.145708874 | 7.36944E-05 | 0.5 | 0.000461466 | FALSE | 0 | relative convergence (4) | 8909.34015587855 | 8959.84954051339 | -4447.67007793928 | 8895.34015587855 | 10046 | 0.01659689 |
| Tnfrsf11a | -10.9097186 | 0.608683448 | 0.355288008 | 0.228343961 | 2.555786175 | 0.338305329 | 0.5 | 0.002198967 | FALSE | 0 | relative convergence (4) | 8842.04826303241 | 8892.55764766725 | -4414.0241315162 | 8828.04826303241 | 10046 | 0.042707071 |
| Ralb | -10.43721221 | 0.226888733 | -0.041272069 | -0.035695728 | 0.259941911 | 0.061844248 | 0.5 | 1.57933E-05 | FALSE | 0 | relative convergence (4) | 10036.5535352544 | 10087.0629198893 | -5011.27676762721 | 10022.5535352544 | 10046 | 0.001747709 |
| Lypd1 | -12.9285255 | 0.936169599 | -0.295726772 | 0.494656547 | 38.57861661 | 0.335829346 | 0.5 | 0.000170625 | FALSE | 0 | relative convergence (4) | 1920.32379623269 | 1970.83318086753 | -953.161898116345 | 1906.32379623269 | 10046 | 0.008623216 |
| Mcm6 | -10.90147733 | 0.379327722 | 0.102906774 | 0.286062389 | 0.844033985 | 0.053994114 | 0.5 | 0.002094323 | FALSE | 0 | relative convergence (4) | 7963.83136919363 | 8014.34075382847 | -3974.91568459682 | 7949.83136919363 | 10046 | 0.041424452 |
| Synpo2l | -12.94976603 | 0.845081876 | 0.301093394 | -0.321089828 | 1.176102047 | 2.07051E-07 | 0.5 | 1.79219E-08 | FALSE | 0 | relative convergence (4) | 1995.64515064643 | 2046.15453528126 | -990.822575323213 | 1981.64515064643 | 10046 | 7.614E-06 |
| Cacna1d | -9.813766893 | 0.501949734 | 0.404946121 | 0.034869409 | 1.968095277 | 0.19376723 | 0.5 | 0.000907028 | FALSE | 0 | relative convergence (4) | 15979.9839447072 | 16030.4933293421 | -7982.99197235361 | 15965.9839447072 | 10046 | 0.024833336 |
| Rft1 | -11.37451994 | 0.245288963 | -0.103809995 | 0.313609117 | 0.45931083 | 4.22134E-05 | 0.5 | 0.001602036 | FALSE | 0 | relative convergence (4) | 5663.16325884017 | 5713.67264347501 | -2824.58162942008 | 5649.16325884017 | 10046 | 0.035386541 |
| Adamts6 | -9.221519245 | 0.286961254 | 0.233974299 | 0.133099769 | 1.194041769 | 9.99727E-05 | 0.5 | 0.000154133 | FALSE | 0 | relative convergence (4) | 21514.7976632109 | 21565.3070478457 | -10750.3988316055 | 21500.7976632109 | 10046 | 0.007937274 |
| Pde4b | -9.261764462 | 0.35285475 | 0.034595056 | 0.136846708 | 2.091493005 | 0.343762705 | 0.5 | 0.002372318 | FALSE | 0 | relative convergence (4) | 20697.6799474272 | 20748.1893320621 | -10341.8399737136 | 20683.6799474272 | 10046 | 0.044488461 |
| Acot11 | -10.84863882 | 0.390822017 | 0.354302485 | 0.201043958 | 0.619807334 | 0.097777441 | 0.5 | 0.00165424 | FALSE | 0 | relative convergence (4) | 8554.56582160868 | 8605.07520624351 | -4270.28291080434 | 8540.56582160868 | 10046 | 0.035925543 |
| Trabd2b | -9.345539952 | 0.260231872 | 0.142706983 | 0.128467944 | 1.359179377 | 0.312908913 | 0.5 | 0.002095335 | FALSE | 0 | relative convergence (4) | 20518.7346242369 | 20569.2440088717 | -10252.3673121184 | 20504.7346242369 | 10046 | 0.041424452 |
| P3h1 | -10.60361094 | 0.262806567 | -0.007749749 | 0.245871878 | 0.239729233 | 0.256685205 | 0.5 | 1.49275E-06 | FALSE | 0 | relative convergence (4) | 9519.65430838315 | 9570.16369301799 | -4752.82715419158 | 9505.65430838315 | 10046 | 0.000302895 |
| Faim | -11.41393578 | 0.403731657 | 0.302344225 | 0.339442902 | 0.130313731 | 2.29255E-12 | 0.5 | 6.91999E-08 | FALSE | 0 | relative convergence (4) | 5837.56414590088 | 5888.07353053572 | -2911.78207295044 | 5823.56414590088 | 10046 | 2.29457E-05 |
| Mras | -10.77554073 | 0.178245571 | 0.339410481 | 0.158628041 | 0.368360165 | 0.000382164 | 0.5 | 0.002542412 | FALSE | 0 | relative convergence (4) | NA | NA | NA | NA | 10046 | 0.046519642 |
| Tmem108 | -12.09908512 | 0.952493386 | 0.495499024 | 0.645746684 | 7.463475734 | 0.127659993 | 0.5 | 0.001525044 | FALSE | 0 | relative convergence (4) | 4645.50650910703 | 4696.01589374186 | -2315.75325455351 | 4631.50650910703 | 10046 | 0.034562383 |
| Rpl29 | -8.381392468 | 0.563210101 | 0.302028638 | 0.808378232 | 0.09058328 | 0.189442807 | 0.5 | 0.001420779 | FALSE | 0 | relative convergence (4) | 32341.6899652807 | 32392.1993499155 | -16163.8449826403 | 32327.6899652807 | 10046 | 0.032905435 |
| Tmem117 | -12.13397784 | 0.787454892 | 0.477997483 | -0.046101369 | 4.672532862 | 0.113840302 | 0.5 | 3.35046E-08 | FALSE | 0 | relative convergence (4) | 3794.88353536227 | 3845.39291999711 | -1890.44176768114 | 3780.88353536227 | 10046 | 1.23107E-05 |
| Tuba1b | -9.426793034 | 0.727685129 | -0.257743118 | 0.733322281 | 0.179036678 | 0.075540987 | 0.5 | 0.000518621 | FALSE | 0 | relative convergence (4) | 21382.0953644732 | 21432.604749108 | -10684.0476822366 | 21368.0953644732 | 10046 | 0.017788246 |

| | | | | | | | | | | | | | | | | |
|---|---|---|---|---|---|---|---|---|---|---|---|---|---|---|---|---|
| Mcrs1 | -10.69964301 | 0.254708414 | 0.32652343 | 0.532865249 | 0.046037468 | 0.145949675 | 0.5 | 3.39948E-06 | FALSE | 0 | relative convergence (4) | 9181.31625056645 | 9231.82563520128 | -4583.65812528322 | 9167.31625056645 | 10046 | 0.000550189 |
| Smarcd1 | -11.06451715 | 0.354416541 | 0.407719859 | 0.217425508 | 0.108495834 | 4.19848E-05 | 0.5 | 0.000148141 | FALSE | 0 | relative convergence (4) | 7173.8802097424 | 7224.38959437724 | -3579.9401048712 | 7159.8802097424 | 10046 | 0.00786014 |
| Tns2 | -9.742632664 | 0.399319428 | 0.276530267 | 0.457935154 | 0.188751735 | 0.110622522 | 0.5 | 0.001489017 | FALSE | 0 | relative convergence (4) | 16840.8904179173 | 16891.3998025521 | -8413.44520895864 | 16826.8904179173 | 10046 | 0.034022167 |
| C030004G16Rik | -13.20333534 | 1.392471701 | 1.695042522 | 1.240909334 | 1.85116977 | 0.228981935 | 0.5 | 0.000715617 | FALSE | 0 | relative convergence (4) | 3126.18502651575 | 3176.69441115059 | -1556.09251325788 | 3112.18502651575 | 10046 | 0.021619574 |
| Ccdc68 | -11.94278371 | 1.170107318 | 1.065787785 | 0.620457568 | 1.138393705 | 0.000189335 | 0.5 | 0.000118479 | FALSE | 0 | relative convergence (4) | 6084.08622013114 | 6134.59560476598 | -3035.04311006557 | 6070.08622013114 | 10046 | 0.006547632 |
| Rgs3 | -10.99444638 | 0.584689832 | 0.17155663 | 0.103183498 | 0.578342228 | 8.15088E-05 | 0.5 | 4.62224E-05 | FALSE | 0 | relative convergence (4) | 8114.53172681753 | 8165.04111145237 | -4050.26586340877 | 8100.53172681753 | 10046 | 0.003570416 |
| Tnc | -12.11326147 | 1.587032898 | 0.80724716 | 1.05502566 | 4.745357129 | 0.000287256 | 0.5 | 1.18808E-05 | FALSE | 0 | relative convergence (4) | 6286.70395804962 | 6337.21334268446 | -3136.35197902481 | 6272.70395804962 | 10046 | 0.001416239 |
| Kank4 | -13.22317879 | 0.979945483 | -0.101046828 | 0.396976786 | 4.154952682 | 3.51776E-14 | 0.5 | 3.49078E-08 | FALSE | 0 | relative convergence (4) | NA | NA | NA | NA | 10046 | 1.24887E-05 |
| Gm49980 | -12.8744307 | 0.752998984 | 0.476228836 | 0.077273027 | 2.803174726 | 8.9602E-07 | 0.5 | 7.40642E-07 | FALSE | 0 | relative convergence (4) | 2213.92867606423 | 2264.43806069907 | -1099.96433803212 | 2199.92867606423 | 10046 | 0.000165066 |
| Arhgap26 | -12.16741541 | 0.636257434 | 0.486965574 | 0.27488634 | 2.571087068 | 0.281369201 | 0.5 | 6.70463E-08 | FALSE | 0 | relative convergence (4) | 3628.00616122256 | 3678.5155458574 | -1807.00308061128 | 3614.00616122256 | 10046 | 2.27874E-05 |
| Fgf1 | -10.88254261 | 0.723367817 | 0.892393176 | 0.448259018 | 1.474183032 | 0.317504351 | 0.5 | 3.54242E-05 | FALSE | 0 | relative convergence (4) | 9827.33904090776 | 9877.8484255426 | -4906.66952045388 | 9813.33904090776 | 10046 | 0.002972791 |
| Ndst3 | -12.36071313 | 1.471128195 | 1.2958031 | 0.659481578 | 5.262620482 | 0.331983596 | 0.5 | 1.16467E-05 | FALSE | 0 | relative convergence (4) | 5057.41988918476 | 5107.9292738196 | -2521.70994459238 | 5043.41988918476 | 10046 | 0.001407788 |
| Ppa2 | -10.63943405 | 0.226947986 | 0.259388899 | 0.42833982 | 0.116600805 | 0.078842947 | 0.5 | 4.55095E-05 | FALSE | 0 | relative convergence (4) | 9451.66113512812 | 9502.17051976296 | -4718.83056756406 | 9437.66113512812 | 10046 | 0.003564746 |
| Ccn1 | -9.678767685 | 0.838466085 | 0.472860327 | 0.515307702 | 1.979775001 | 0.363430999 | 0.5 | 0.001134995 | FALSE | 0 | relative convergence (4) | 18633.7506469272 | 18684.2600315621 | -9309.87532346361 | 18619.7506469272 | 10046 | 0.028521727 |
| Rgs5 | -10.89731847 | 1.427181085 | 1.016877321 | 0.800493939 | 11.59128585 | 0.000334659 | 0.5 | 0.001294796 | FALSE | 0 | relative convergence (4) | 10167.4233517827 | 10217.9327364176 | -5076.71167589137 | 10153.4233517827 | 10046 | 0.030955819 |
| Casq1 | -12.76951581 | 1.382351157 | 0.097508337 | 0.439948634 | 3.450795796 | 0.39587991 | 0.5 | 2.25813E-05 | FALSE | 0 | relative convergence (4) | 3827.04747831509 | 3877.55686294992 | -1906.52373915754 | 3813.04747831509 | 10046 | 0.002205153 |
| Igsf9 | -10.96849241 | 0.777758613 | 0.937989283 | 0.944366101 | 2.275400425 | 0.328194469 | 0.5 | 0.001022083 | FALSE | 0 | relative convergence (4) | 9712.18595083649 | 9762.69533547132 | -4849.09297541824 | 9698.18595083649 | 10046 | 0.026670294 |
| Spout1 | -11.49467393 | 0.263087469 | 0.120020468 | 0.245176435 | 0.229841049 | 0.023841421 | 0.5 | 0.001177571 | FALSE | 0 | relative convergence (4) | 5253.49430524519 | 5304.00368988003 | -2619.74715262259 | 5239.49430524519 | 10046 | 0.029152159 |
| D330023K18Rik | -11.72778473 | 0.288668779 | 0.288981437 | 0.530611883 | 0.290983661 | 0.075159556 | 0.5 | 0.001431018 | FALSE | 0 | relative convergence (4) | 4709.54964527321 | 4760.05902990805 | -2347.77482263661 | 4695.54964527321 | 10046 | 0.033030026 |
| Ak1 | -10.88436406 | 0.412100961 | 0.301460832 | 0.628049889 | 0.357464137 | 0.248194472 | 0.5 | 0.002588605 | FALSE | 0 | relative convergence (4) | 8949.43994543591 | 8999.94933007075 | -4467.71997271796 | 8935.43994543591 | 10046 | 0.047048242 |
| Niban2 | -10.14379894 | 0.403396408 | 0.008641756 | -0.005156014 | 0.457111001 | 0.1188949 | 0.5 | 7.44328E-06 | FALSE | 0 | relative convergence (4) | 12686.3779340897 | 12736.8873187245 | -6336.18896704484 | 12672.3779340897 | 10046 | 0.001004954 |
| Ifngr2 | -10.31940531 | 0.451019857 | 0.211608664 | 0.151427454 | 0.206633633 | 0.160192803 | 0.5 | 5.63178E-05 | FALSE | 0 | relative convergence (4) | 11782.493981144 | 11833.0033657789 | -5884.24699057202 | 11768.493981144 | 10046 | 0.004094334 |
| Erg | -10.84745568 | 0.5664009 | 0.893289333 | 0.576598036 | 2.236050017 | 0.344807426 | 0.5 | 0.002535665 | FALSE | 0 | relative convergence (4) | 9801.94279891616 | 9852.45218355099 | -4893.97139945808 | 9787.94279891616 | 10046 | 0.046458711 |
| Gm29721 | -13.25534122 | 0.642406397 | -0.429162248 | 0.730916216 | 0.221824598 | 0.378850468 | 0.5 | 0.00047997 | FALSE | 0 | relative convergence (4) | 1647.32743354184 | 1697.83681817668 | -816.663716770922 | 1633.32743354184 | 10046 | 0.017081661 |
| Pnldc1 | -12.43419451 | 0.776419321 | 0.258781249 | 0.569980952 | 1.50321541 | 0.367057922 | 0.5 | 0.000790968 | FALSE | 0 | relative convergence (4) | 3463.91692094289 | 3514.42630557773 | -1724.95846047144 | 3449.91692094289 | 10046 | 0.022830586 |
| Eif2b5 | -10.76236041 | 0.226732328 | -0.046757134 | 0.1854646 | 0.151342823 | 0.118883131 | 0.5 | 9.93835E-05 | FALSE | 0 | relative convergence (4) | 8459.43806521931 | 8509.94744985414 | -4222.71903260965 | 8445.43806521931 | 10046 | 0.005737733 |
| Eif4g1 | -8.887499534 | 0.177672349 | 0.148302535 | 0.224536199 | 0.070888667 | 0.13904463 | 0.5 | 5.34029E-06 | FALSE | 0 | relative convergence (4) | 23188.9094466502 | 23239.418831285 | -11587.4547233251 | 23174.9094466502 | 10046 | 0.000772354 |
| P3h2 | -10.82413414 | 0.70183069 | 0.201884823 | 0.440849299 | 3.166838839 | 0.343705782 | 0.5 | 8.33153E-06 | FALSE | 0 | relative convergence (4) | 9665.16195300657 | 9715.6713376414 | -4825.58097650328 | 9651.16195300657 | 10046 | 0.001099681 |
| Pdia5 | -11.68294879 | 0.294375116 | 0.271515563 | 0.411144195 | 1.296029515 | 0.203122318 | 0.5 | 0.001956385 | FALSE | 0 | relative convergence (4) | 4732.09601940979 | 4782.60540404463 | -2359.0480097049 | 4718.09601940979 | 10046 | 0.039578945 |
| Npr3 | -11.84752021 | 0.757544567 | -0.188472835 | 0.811492691 | 4.418523908 | 0.238563593 | 0.5 | 6.82522E-05 | FALSE | 0 | relative convergence (4) | 5008.23819567261 | 5058.74758030745 | -2497.11909783631 | 4994.23819567261 | 10046 | 0.004593511 |
| Cdh6 | -10.44295709 | 1.018733154 | 1.034508429 | 0.709576249 | 2.994459312 | 0.000414265 | 0.5 | 0.001651812 | FALSE | 0 | relative convergence (4) | 14593.4786554941 | 14643.988040129 | -7289.73932774706 | 14579.4786554941 | 10046 | 0.035925543 |
| Cthrc1 | -11.33442565 | 0.844477903 | 0.048589453 | 1.051286658 | 6.730263845 | 0.853756021 | 0.5 | 0.000153216 | FALSE | 0 | relative convergence (4) | 8409.77847992588 | 8460.28786456071 | -4197.88923996294 | 8395.77847992588 | 10046 | 0.007920063 |
| Abra | -12.6360696 | 1.318439889 | 1.474623713 | 1.28690744 | 6.245829403 | 0.450662675 | 0.5 | 0.001178244 | FALSE | 0 | relative convergence (4) | 4256.28631220962 | 4306.79569684446 | -2121.14315610481 | 4242.28631220962 | 10046 | 0.029152159 |
| Rspo2 | -13.03894972 | 1.148929513 | -0.059604116 | -0.031753641 | 43.36914483 | 0.382169851 | 0.5 | 6.12755E-05 | FALSE | 0 | relative convergence (4) | 1879.54840604051 | 1930.05779067535 | -932.774203020257 | 1865.54840604051 | 10046 | 0.004309979 |
| Tnfrsf11b | -11.57526914 | 0.988812079 | 0.380872908 | 0.738184796 | 4.031002709 | 0.000137075 | 0.5 | 2.96381E-05 | FALSE | 0 | relative convergence (4) | 6911.87009113207 | 6962.3794757669 | -3448.93504556603 | 6897.87009113207 | 10046 | 0.002616432 |
| Ccn3 | -8.87422887 | 1.127883382 | 0.725633688 | 0.661828997 | 2.964413492 | 0.000937381 | 0.5 | 0.000749452 | FALSE | 0 | relative convergence (4) | 30873.9900856079 | 30924.4994702428 | -15429.995042804 | 30859.9900856079 | 10046 | 0.022101521 |
| Kif26b | -11.78552282 | 1.301764284 | 0.777965821 | 0.425293139 | 3.082793381 | 0.173516249 | 0.5 | 2.9613E-09 | FALSE | 0 | relative convergence (4) | 6369.18425458952 | 6419.69363922435 | -3177.59212729476 | 6355.18425458952 | 10046 | 1.91709E-06 |
| Coq8a | -11.58047625 | 0.739681316 | 0.763536086 | 0.381666214 | 0.671169223 | 0.000234628 | 0.5 | 0.002439648 | FALSE | 0 | relative convergence (4) | 6356.19313570868 | 6406.70252034352 | -3171.09656785434 | 6342.19313570868 | 10046 | 0.045328879 |
| Ephx1 | -9.690689729 | 0.203132311 | -0.027321872 | 0.085553943 | 0.424792831 | 0.081918482 | 0.5 | 3.49652E-07 | FALSE | 0 | relative convergence (4) | NA | NA | NA | NA | 10046 | 8.9689E-05 |
| Smyd2 | -10.90697746 | 0.204295172 | 0.073274659 | 0.055081343 | 0.356803596 | 0.080387816 | 0.5 | 0.001893494 | FALSE | 0 | relative convergence (4) | NA | NA | NA | NA | 10046 | 0.038593779 |
| Adamtsl5 | -9.491258164 | 0.272240284 | 0.103444584 | 0.394106473 | 0.428030256 | 0.118077559 | 0.5 | 1.29579E-08 | FALSE | 0 | relative convergence (4) | 18365.0447980882 | 18415.554182723 | -9175.5223990441 | 18351.0447980882 | 10046 | 5.94537E-06 |
| Klf16 | -12.05629648 | 0.333737699 | 0.137831353 | 0.243580875 | 0.364651748 | 0.000243927 | 0.5 | 0.001500846 | FALSE | 0 | relative convergence (4) | 3536.69723258181 | 3587.20661721665 | -1761.34861629091 | 3522.69723258181 | 10046 | 0.034234912 |
| Csnk1g2 | -9.955905282 | 0.23974666 | 0.029494366 | 0.497251701 | 0.032292207 | 0.172238877 | 0.5 | 0.000529547 | FALSE | 0 | relative convergence (4) | 14171.2841909318 | 14221.7935755667 | -7078.64209546592 | 14157.2841909318 | 10046 | 0.017889669 |
| Sppl2b | -11.66844249 | 0.30404772 | -0.060776287 | 0.419781075 | 0.219114629 | 0.190877903 | 0.5 | 0.000737394 | FALSE | 0 | relative convergence (4) | 4756.80889488088 | 4807.31827951571 | -2371.40444744044 | 4742.80889488088 | 10046 | 0.022032696 |
| Zbtb7a | -9.296902796 | 0.244605464 | -0.139437044 | 0.148411088 | 0.191810378 | 0.077665469 | 0.5 | 1.76261E-05 | FALSE | 0 | relative convergence (4) | 19352.2239308673 | 19402.7333155021 | -9669.11196543365 | 19338.2239308673 | 10046 | 0.001872079 |
| Ankrd24 | -12.09910222 | 0.372751866 | 0.445912848 | 0.174075626 | 0.280406861 | 0.085777897 | 0.5 | 0.00054166 | FALSE | 0 | relative convergence (4) | 3535.96869974141 | 3586.47808437625 | -1760.9843498707 | 3521.96869974141 | 10046 | 0.017960652 |
| Nuak1 | -10.32543943 | 0.308371442 | 0.049538297 | 0.045485394 | 0.939611491 | 0.086186017 | 0.5 | 0.001639905 | FALSE | 0 | relative convergence (4) | 11563.7233421086 | 11614.2327267434 | -5774.86167105429 | 11549.7233421086 | 10046 | 0.035925543 |
| Arhgap1 | -10.11927656 | 0.166768668 | -0.197983217 | 0.082620189 | 0.104008048 | 2.84943E-05 | 0.5 | 0.001886102 | FALSE | 0 | relative convergence (4) | 12127.2143045337 | 12177.7236891685 | -6056.60715226685 | 12113.2143045337 | 10046 | 0.038593779 |
| Cry2 | -10.28544332 | 0.311281575 | 0.136430431 | 0.199113441 | 0.559359602 | 0.167612264 | 0.5 | 0.002489025 | FALSE | 0 | relative convergence (4) | 11012.580972867 | 11063.0903575018 | -5499.2904864335 | 10998.580972867 | 10046 | 0.045789294 |
| Fjx1 | -12.32578454 | 1.07562773 | -0.009808393 | 0.326485964 | 1.290095772 | 0.000168156 | 0.5 | 0.00178959 | FALSE | 0 | relative convergence (4) | 4231.45827501519 | 4281.96765965003 | -2108.7291375076 | 4217.45827501519 | 10046 | 0.037841758 |
| Bdnf | -11.68375209 | 1.094005724 | 1.169296046 | 0.721016018 | 2.778930589 | 0.233654016 | 0.5 | 0.000885905 | FALSE | 0 | relative convergence (4) | 7041.55207577658 | 7092.06146041142 | -3513.77603788829 | 7027.55207577658 | 10046 | 0.024502896 |
| Bmf | -11.92812473 | 0.461135486 | 0.471817543 | 0.275229096 | 0.595556258 | 1.26212E-22 | 0.5 | 2.05927E-06 | FALSE | 1 | singular convergence (7) | NA | NA | NA | NA | 10046 | 0.000384031 |
| Ehd4 | -9.590138038 | 0.38881733 | 0.113628879 | -0.030465409 | 0.832177786 | 0.181490404 | 0.5 | 0.000300227 | FALSE | 0 | relative convergence (4) | 17936.9329197589 | 17987.4423043937 | -8961.46645987943 | 17922.9329197589 | 10046 | 0.012794924 |
| Mical1 | -10.19173607 | 0.23700164 | 0.01759787 | 0.026661877 | 0.233945358 | 4.34994E-06 | 0.5 | 0.000134489 | FALSE | 0 | relative convergence (4) | 11930.2025438128 | 11980.7119284477 | -5958.10127190641 | 11916.2025438128 | 10046 | 0.007255479 |
| Gja1 | -10.17061923 | 0.652246713 | -0.131066629 | -0.095238428 | 1.406161499 | 0.000222379 | 0.5 | 0.000764332 | FALSE | 0 | relative convergence (4) | 13507.2934494498 | 13557.8028340846 | -6746.64672472489 | 13493.2934494498 | 10046 | 0.022357569 |
| E130311K13Rik | -11.65205077 | 0.35969602 | 0.023882707 | 0.313192247 | 0.355864723 | 0.000189422 | 0.5 | 0.000804032 | FALSE | 0 | relative convergence (4) | 4934.68211602809 | 4985.19150066293 | -2460.34105801405 | 4920.68211602809 | 10046 | 0.023109554 |
| Gask1b | -9.099004711 | 0.380690049 | -0.077185819 | 0.120164461 | 0.850537043 | 0.000152822 | 0.5 | 5.69012E-06 | FALSE | 0 | relative convergence (4) | 23148.9167771832 | 23199.4261618181 | -11567.4583885916 | 23134.9167771832 | 10046 | 0.000814286 |
| Nes | -10.50573504 | 1.019190559 | 1.227347746 | 0.909096627 | 1.503886972 | 0.395901658 | 0.5 | 0.002525645 | FALSE | 0 | relative convergence (4) | 14659.4096303448 | 14709.9190149796 | -7322.7048151724 | 14645.4096303448 | 10046 | 0.046337578 |
| Igfbp2 | -12.92560128 | 3.372382503 | 1.000494926 | 1.152165611 | 14.15556876 | 0.000508038 | 0.5 | 1.06098E-11 | FALSE | 0 | relative convergence (4) | 6234.37486666327 | 6284.88425129811 | -3110.18743333164 | 6220.37486666327 | 10046 | 1.803E-08 |
| Arpc2 | -7.935478805 | 0.226017492 | -0.091071581 | 0.110064963 | 0.234749589 | 0.068826462 | 0.5 | 0.002058901 | FALSE | 0 | relative convergence (4) | 35946.5411400631 | 35997.050524698 | -17966.2705700316 | 35932.5411400631 | 10046 | 0.041284309 |
| Tmbim1 | -9.924901344 | 0.386757684 | 0.078604798 | 0.251351343 | 0.138567344 | 0.053698162 | 0.5 | 1.15684E-05 | FALSE | 0 | relative convergence (4) | 14520.6904510235 | 14571.1998356583 | -7253.34522551174 | 14506.6904510235 | 10046 | 0.001407788 |
| Slc4a3 | -10.67889229 | 0.491427533 | 0.429862688 | 0.308813436 | 0.293821963 | 0.000134699 | 0.5 | 7.04042E-05 | FALSE | 0 | relative convergence (4) | 9775.54343460006 | 9826.05281923489 | -4880.77171730003 | 9761.54343460006 | 10046 | 0.004714997 |
| Kcne4 | -11.084462 | 0.408935027 | -0.140664202 | 0.206265237 | 1.813839446 | 0.163409341 | 0.5 | 0.000945471 | FALSE | 0 | relative convergence (4) | 7537.87174672335 | 7588.38113135819 | -3761.93587336167 | 7523.87174672335 | 10046 | 0.025604932 |
| Serpine2 | -9.19067738 | 1.095444273 | 0.927938478 | 0.778958865 | 2.62440598 | 0.316385829 | 0.5 | 0.000788587 | FALSE | 0 | relative convergence (4) | 27543.5000180396 | 27594.0094026745 | -13764.7500090198 | 27529.5000180396 | 10046 | 0.022810305 |
| Sphkap | -10.83375631 | 1.104070099 | 0.808213528 | 1.174480253 | 5.484929407 | 0.000297626 | 0.5 | 1.56209E-06 | FALSE | 0 | relative convergence (4) | 11276.4359591877 | 11326.9453438225 | -5631.21797959385 | 11262.4359591877 | 10046 | 0.000309149 |
| Mlph | -13.97187344 | 1.152035411 | 0.259677018 | 0.302546816 | 3.647106611 | 5.45063E-05 | 0.5 | 0.000227687 | FALSE | 0 | relative convergence (4) | 1122.17600655232 | 1172.68539118716 | -554.088003276162 | 1108.17600655232 | 10046 | 0.010456617 |
| Klhl30 | -13.14873113 | 1.119521758 | 0.269452393 | 1.127368502 | 1.139026788 | 0.478384086 | 0.5 | 0.000774578 | FALSE | 0 | relative convergence (4) | 2735.06494801125 | 2785.57433264609 | -1360.53247400562 | 2721.06494801125 | 10046 | 0.022500836 |
| Tnnt2 | -10.63344483 | 1.187580461 | 1.23944062 | 0.498773623 | 7.149731964 | 0.000344517 | 0.5 | 0.000404572 | FALSE | 0 | relative convergence (4) | 12705.1332558753 | 12755.6426405102 | -6345.56662793767 | 12691.1332558753 | 10046 | 0.01528425 |
| Qsox1 | -9.621961168 | 0.61738187 | -0.153287999 | 0.021876726 | 0.378147688 | 0.000482684 | 0.5 | 0.002584011 | FALSE | 0 | relative convergence (4) | 18133.9763661698 | 18184.4857508047 | -9059.98818308492 | 18119.9763661698 | 10046 | 0.047048242 |
| Ptprs | -8.957777741 | 0.121362721 | -0.033474889 | -0.077261099 | 0.193058683 | 4.25292E-05 | 0.5 | 0.002299114 | FALSE | 0 | relative convergence (4) | 22474.4074128978 | 22524.9167975326 | -11230.2037064489 | 22460.4074128978 | 10046 | 0.043776544 |
| Gpr108 | -10.34338038 | 0.153452209 | 0.175173524 | 0.337976272 | 8.20765E-07 | 0.156329551 | 0.5 | 0.000806936 | FALSE | 0 | relative convergence (4) | NA | NA | NA | NA | 10046 | 0.023118855 |
| Fhl2 | -12.06800251 | 0.80148117 | 0.397213814 | 0.402149407 | 2.416835465 | 0.000133243 | 0.5 | 0.000973486 | FALSE | 0 | relative convergence (4) | 4700.45130661778 | 4750.96069125262 | -2343.22565330889 | 4686.45130661778 | 10046 | 0.026001063 |
| Dusp6 | -11.82457148 | 0.603014403 | -0.229649287 | -0.261135589 | 1.294771292 | 0.117515576 | 0.5 | 0.000653872 | FALSE | 0 | relative convergence (4) | 4195.17325687628 | 4245.68264151112 | -2090.58662843814 | 4181.17325687628 | 10046 | 0.020286488 |
| Lrig3 | -10.43156569 | 0.171870973 | -0.001575899 | 0.038805164 | 0.485034209 | 2.31394E-07 | 0.5 | 0.000907846 | FALSE | 0 | relative convergence (4) | 10267.8538450502 | 10318.363229685 | -5126.9269225251 | 10253.8538450502 | 10046 | 0.024833336 |
| Vcp | -9.06779621 | 0.401066361 | 0.092676724 | 0.408577806 | 0.068345321 | 0.208415452 | 0.5 | 2.16399E-06 | FALSE | 0 | relative convergence (4) | 22537.5947949375 | 22588.1041795724 | -11261.7973974688 | 22523.5947949375 | 10046 | 0.000387099 |
| Tln1 | -8.58046576 | 0.36825272 | 0.449346342 | 0.066462883 | 0.407984089 | 0.000253175 | 0.5 | 0.000404535 | FALSE | 0 | relative convergence (4) | 28829.0428433436 | 28879.5522279785 | -14407.5214216718 | 28815.0428433436 | 10046 | 0.01528425 |
| Cspg4 | -10.1527774 | 1.253993703 | 1.064878628 | 0.734124128 | 1.591190017 | 0.349250859 | 0.5 | 0.000818937 | FALSE | 0 | relative convergence (4) | 18545.1689163297 | 18595.6783009646 | -9265.58445816486 | 18531.1689163297 | 10046 | 0.023340564 |
| Stoml1 | -11.22324188 | 0.319227048 | 0.014680824 | 0.249933135 | 0.110371917 | 7.74356E-05 | 0.5 | 4.06322E-06 | FALSE | 0 | relative convergence (4) | 6485.61771425898 | 6536.12709889382 | -3235.80885712949 | 6471.61771425898 | 10046 | 0.000627722 |
| Scamp3 | -10.60518392 | 0.23563982 | 0.047687306 | 0.305053666 | 0.064922231 | 5.20852E-05 | 0.5 | 0.001428338 | FALSE | 0 | relative convergence (4) | 9543.92643006916 | 9594.43581470399 | -4764.96321503458 | 9529.92643006916 | 10046 | 0.033024238 |
| Efna4 | -13.38753437 | 0.617778376 | 0.981016974 | 0.943681783 | 1.288633999 | 0.365258757 | 0.5 | 0.000958012 | FALSE | 0 | relative convergence (4) | 1726.79326461057 | 1777.30264924541 | -856.396632305284 | 1712.79326461057 | 10046 | 0.025803244 |
| Pmvk | -11.34622502 | 0.389798305 | 0.533306218 | 0.608700412 | 0.673348821 | 0.202462159 | 0.5 | 0.000827529 | FALSE | 0 | relative convergence (4) | 6444.96976627349 | 6495.47915090833 | -3215.48488313675 | 6430.96976627349 | 10046 | 0.023438025 |
| Cdc42se1 | -10.20129366 | 0.403732541 | 0.03265414 | 0.347559398 | 0.196994394 | 7.84921E-05 | 0.5 | 0.00112082 | FALSE | 0 | relative convergence (4) | 12513.6074524588 | 12564.1168370937 | -6249.80372622941 | 12499.6074524588 | 10046 | 0.028322583 |
| Sema3b | -10.679791 | 0.242007653 | -0.191845199 | -0.037946948 | 0.668327115 | 5.62958E-05 | 0.5 | 0.002441697 | FALSE | 0 | relative convergence (4) | 8862.67190283938 | 8913.18128747422 | -4424.33595141969 | 8848.67190283938 | 10046 | 0.045328879 |
| Mir6236 | -8.042244205 | 0.839071725 | 2.395584484 | 0.649206082 | 0.540788748 | 0.883805194 | 0.5 | 0.000107897 | FALSE | 0 | relative convergence (4) | 42585.2204047116 | 42635.7297893464 | -21285.6102023558 | 42571.2204047116 | 10046 | 0.006137479 |
| Ptpn23 | -10.95216886 | 0.205750604 | 0.022743613 | 0.263031259 | 0.089445192 | 0.070284399 | 0.5 | 0.001620429 | FALSE | 0 | relative convergence (4) | 7525.28594838538 | 7575.79533302022 | -3755.64297419269 | 7511.28594838538 | 10046 | 0.035704588 |
| Stac | -11.49319367 | 1.061659144 | 1.289135673 | 1.036227261 | 3.264849689 | 0.000328384 | 0.5 | 0.000850396 | FALSE | 0 | relative convergence (4) | 8240.23910425815 | 8290.74848889298 | -4113.11955212907 | 8226.23910425815 | 10046 | 0.023839755 |
| Susd5 | -10.62503154 | 1.286851154 | 1.34382029 | 1.135369601 | 1.624288378 | 0.411002074 | 0.5 | 0.002411496 | FALSE | 0 | relative convergence (4) | 15190.3483851067 | 15240.8577697416 | -7588.17419255337 | 15176.3483851067 | 10046 | 0.045033366 |
| Utrn | -7.958032338 | 0.1952745 | 0.029385401 | -0.072180709 | 0.407298628 | 0.151479755 | 0.5 | 1.29833E-05 | FALSE | 0 | relative convergence (4) | 36236.2736369774 | 36286.7830216122 | -18111.1368184887 | 36222.2736369774 | 10046 | 0.001495832 |
| Chmp4c | -11.47648324 | 0.89827658 | 1.249016898 | 0.812727445 | 1.045917432 | 0.410988451 | 0.5 | 0.001114655 | FALSE | 0 | relative convergence (4) | 7684.55503061233 | 7735.06441524716 | -3835.27751530616 | 7670.55503061233 | 10046 | 0.028271897 |
| Postn | -8.206801752 | 1.032940754 | -0.15303857 | -0.038650518 | 1.837977121 | 0.001230033 | 0.5 | 9.93948E-05 | FALSE | 0 | relative convergence (4) | 39109.006720663 | 39159.5161052978 | -19547.5033603315 | 39095.006720663 | 10046 | 0.005737733 |
| Alg5 | -10.42086039 | 0.152600211 | -0.109073197 | -0.068485485 | 0.052444289 | 0.017502746 | 0.5 | 0.001645267 | FALSE | 0 | relative convergence (4) | 10065.7929238056 | 10116.3023084405 | -5025.89646190281 | 10051.7929238056 | 10046 | 0.035925543 |
| Itga7 | -11.59869404 | 1.216025314 | 1.17522484 | 0.912588852 | 0.888727987 | 0.264133456 | 0.5 | 0.000879084 | FALSE | 0 | relative convergence (4) | 8157.09187300481 | 8207.60125763964 | -4071.5459365024 | 8143.09187300481 | 10046 | 0.024439962 |
| Castor1 | -11.32915751 | 0.296878343 | 0.009719283 | 0.216839199 | 0.447276202 | 0.116959121 | 0.5 | 9.42098E-05 | FALSE | 0 | relative convergence (4) | 6117.64003552756 | 6168.1494201624 | -3051.82001776378 | 6103.64003552756 | 10046 | 0.005564624 |
| Oaz2 | -9.696931992 | 0.273828577 | 0.130491939 | 0.350717809 | 0.154518485 | 0.156271868 | 0.5 | 0.000650155 | FALSE | 0 | relative convergence (4) | 16123.8586740275 | 16174.3680586624 | -8054.92933701377 | 16109.8586740275 | 10046 | 0.020286488 |
| Fam81a | -12.87202782 | 1.656488653 | 0.660529261 | 0.763646211 | 2.023383274 | 0.000276983 | 0.5 | 0.000251907 | FALSE | 0 | relative convergence (4) | 3848.72001181327 | 3899.22939644811 | -1917.36000590664 | 3834.72001181327 | 10046 | 0.011207292 |
| Rsl24d1 | -11.0966409 | 0.366492717 | -0.410422037 | 0.114212445 | 0.039103763 | 0.227789926 | 0.5 | 2.09683E-07 | FALSE | 0 | relative convergence (4) | 6682.20806532505 | 6732.71744995988 | -3334.10403266252 | 6668.20806532505 | 10046 | 6.06518E-05 |
| Unc13c | -11.51575179 | 0.69146279 | -0.072164446 | 0.004951391 | 12.00452646 | 0.271793273 | 0.5 | 1.07818E-08 | FALSE | 0 | relative convergence (4) | 5180.60324709971 | 5231.11263173455 | -2583.30162354985 | 5166.60324709971 | 10046 | 5.27123E-06 |
| Pdk1 | -11.89867613 | 0.647084685 | 0.217597258 | 0.298026889 | 0.725370605 | 3.97609E-05 | 0.5 | 2.8499E-08 | FALSE | 0 | relative convergence (4) | 4436.38295554577 | 4486.89234018061 | -2211.19147777289 | 4422.38295554577 | 10046 | 1.10698E-05 |
| Gap43 | -11.2926653 | 0.461199567 | 1.627245447 | 0.865647855 | 3.970227927 | 0.527806914 | 0.5 | 0.001295613 | FALSE | 0 | relative convergence (4) | 6829.54492224005 | 6880.05430687489 | -3407.77246112002 | 6815.54492224005 | 10046 | 0.030955819 |
| Phldb2 | -9.179869914 | 0.386406223 | 0.249301884 | 0.039284717 | 0.751542268 | 9.29818E-05 | 0.5 | 0.000742306 | FALSE | 0 | relative convergence (4) | 22203.7044073028 | 22254.2137919377 | -11094.8522036514 | 22189.7044073028 | 10046 | 0.022034183 |
| Gm29686 | -13.99463508 | 2.205816817 | 1.291749788 | 1.071000171 | 2.030148772 | 0.000196318 | 0.5 | 0.000152216 | FALSE | 0 | relative convergence (4) | 2369.11876682765 | 2419.62815146248 | -1177.55938341382 | 2355.11876682765 | 10046 | 0.007920063 |

### Table S8. SMC Differentially Expressed Genes: Hypoxia versus Normoxia

**Smooth Muscle Cells: Genes that positively correlate with chronic hypoxia**

| gene | Beta0 | Beta1 | Disp | Sigma | PiC | PiI | P.Beta | P.Pi | Hessian | Convergence | Message | AIC | BIC | logLik | deviance | df.resid | fdrP |
|---|---|---|---|---|---|---|---|---|---|---|---|---|---|---|---|---|---|
| Pitpnm1 | -11.43118091 | 0.308955457 | 0.249488231 | 0.470409488 | 0.23272473 | 0.145736118 | 0.5 | 0.000306637 | FALSE | 0 | relative convergence (4) | 7735.1817809098 | 7790.42416502688 | -3860.5908904549 | 7721.1817809098 | 19760 | 0.009665553 |
| Cd248 | -11.11940544 | 0.629636832 | 0.296037169 | 0.335061536 | 1.032512502 | 0.290958964 | 0.5 | 0.002978537 | FALSE | 0 | relative convergence (4) | 9854.85876458958 | 9910.10114870665 | -4920.42938229479 | 9840.85876458958 | 19760 | 0.045836632 |
| Fibp | -10.79251895 | 0.445876966 | 0.13145159 | 0.294880044 | 5.22201E-07 | 1.25467E-17 | 0.5 | 0.000217733 | FALSE | 0 | relative convergence (4) | 11872.5673953498 | 11927.8097794669 | -5929.2836976749 | 11858.5673953498 | 19760 | 0.00755519 |
| Ehbp1l1 | -9.365684391 | 0.325617591 | 0.279747167 | 0.566081126 | 0.095641107 | 0.216207788 | 0.5 | 0.001180407 | FALSE | 0 | relative convergence (4) | 28335.0633479368 | 28390.3057320539 | -14160.5316739684 | 28321.0633479368 | 19760 | 0.024540046 |
| Ppp2r5b | -11.74816629 | 0.454607658 | 0.028427464 | 0.355560883 | 5.51509E-07 | 7.7867E-58 | 0.5 | 0.001423228 | FALSE | 1 | singular convergence (7) | NA | NA | NA | NA | 19760 | 0.027788342 |
| Piezo2 | -11.16077863 | 0.964455571 | 0.678424342 | 0.386078625 | 7.611609715 | 0.392705716 | 0.5 | 9.06404E-10 | FALSE | 0 | relative convergence (4) | 9880.27355292077 | 9935.51593703785 | -4933.13677646039 | 9866.27355292077 | 19760 | 3.27341E-07 |
| Spire1 | -10.37688485 | 0.301478453 | -0.049666091 | 0.102088784 | 1.176911441 | 0.014622159 | 0.5 | 6.06335E-09 | FALSE | 0 | relative convergence (4) | 15159.7067133233 | 15214.9490974403 | -7572.85335666164 | 15145.7067133233 | 19760 | 1.59668E-06 |
| Ccdc68 | -11.26614958 | 0.559213289 | 0.346722632 | 0.381579598 | 1.081410496 | 7.27625E-05 | 0.5 | 6.91237E-05 | FALSE | 0 | relative convergence (4) | 8915.6263960299 | 8970.86878014697 | -4450.81319801495 | 8901.6263960299 | 19760 | 0.003200455 |
| Rab27b | -11.19979797 | 0.429820371 | 0.084402245 | -0.029744639 | 2.037722308 | 0.00015249 | 0.5 | 0.00158559 | FALSE | 0 | relative convergence (4) | 8642.53401856449 | 8697.77640268157 | -4314.26700928225 | 8628.53401856449 | 19760 | 0.02995035 |
| Wtip | -10.20063304 | 0.864233433 | 0.36158237 | 0.704905359 | 0.274392666 | 0.000286588 | 0.5 | 9.1678E-06 | FALSE | 0 | relative convergence (4) | 20662.9816177441 | 20718.2240018612 | -10324.4908088721 | 20648.9816177441 | 19760 | 0.000640227 |
| Uba2 | -9.563961384 | 0.218689578 | 0.049471532 | 0.367000362 | 0.066274756 | 8.73664E-05 | 0.5 | 6.10744E-05 | FALSE | 0 | relative convergence (4) | 24888.6078043598 | 24943.8501884769 | -12437.3039021799 | 24874.6078043598 | 19760 | 0.002902183 |
| Gpi1 | -8.926148123 | 0.363099316 | 0.072201587 | 0.385912156 | 0.086400126 | 0.115053962 | 0.5 | 6.87747E-05 | FALSE | 0 | relative convergence (4) | 35546.4269867962 | 35601.6693709133 | -17766.2134933981 | 35532.4269867962 | 19760 | 0.003196001 |
| 5430431A17Rik | -10.77726444 | 0.811540163 | 0.856406627 | 0.787571945 | 4.990464213 | 0.000178354 | 0.5 | 2.85076E-05 | FALSE | 0 | relative convergence (4) | 13440.8253525333 | 13496.0677366504 | -6713.41267626664 | 13426.8253525333 | 19760 | 0.00159441 |
| Atf5 | -10.31113144 | 0.835125326 | 0.146002004 | 0.505660328 | 1.911490514 | 0.000402103 | 0.5 | 0.001274781 | FALSE | 0 | relative convergence (4) | 18694.2908772683 | 18749.5332613854 | -9340.14543863416 | 18680.2908772683 | 19760 | 0.025947241 |
| Tbc1d17 | -11.12245425 | 0.533873803 | 0.125847702 | 0.424820563 | 0.158105404 | 0.000136272 | 0.5 | 1.96593E-05 | FALSE | 0 | relative convergence (4) | 9980.17087181191 | 10035.413255929 | -4983.08543590595 | 9966.17087181191 | 19760 | 0.001224104 |
| Ap2a1 | -10.26281039 | 0.350110662 | 0.06908313 | 0.202027596 | 0.10621614 | 9.0028E-05 | 0.5 | 0.000638754 | FALSE | 0 | relative convergence (4) | 16615.1672171573 | 16670.4096012744 | -8300.58360857867 | 16601.1672171573 | 19760 | 0.015987823 |
| Nucb1 | -9.229832556 | 0.382710119 | -0.033365222 | 0.111494108 | 0.075125633 | 0.000129133 | 0.5 | 9.08847E-05 | FALSE | 0 | relative convergence (4) | 29606.6692622385 | 29661.9116463556 | -14796.3346311193 | 29592.6692622385 | 19760 | 0.003881022 |
| Dbp | -11.43145559 | 1.90896355 | -0.431923817 | -0.957948514 | 2.247393058 | 0.000264149 | 0.5 | 1.30757E-06 | FALSE | 0 | relative convergence (4) | 9284.62290664709 | 9339.86529076416 | -4635.31145332355 | 9270.62290664709 | 19760 | 0.000131172 |
| Nomo1 | -10.4033629 | 0.317412949 | -0.031583452 | 0.243604533 | 0.236772292 | 0.112023087 | 0.5 | 2.14526E-09 | FALSE | 0 | relative convergence (4) | 14931.69811185 | 14986.9404959671 | -7458.84905592501 | 14917.69811185 | 19760 | 7.13527E-07 |
| Ldha | -9.077062301 | 0.536078841 | -0.094682877 | 0.461255283 | 0.166224705 | 0.000311638 | 0.5 | 0.003179781 | FALSE | 0 | relative convergence (4) | 34231.5765242021 | 34286.8189083192 | -17108.7882621011 | 34217.5765242021 | 19760 | 0.047396742 |
| Tmem86a | -11.75865853 | 0.680758453 | 0.087501347 | 0.200160517 | 0.594206604 | 0.255159886 | 0.5 | 0.000163866 | FALSE | 0 | relative convergence (4) | 6202.02635060464 | 6257.26873472171 | -3094.01317530232 | 6188.02635060464 | 19760 | 0.006182877 |
| BC051226 | -12.61724752 | 0.492780262 | 0.451977734 | 0.443278965 | 0.267771481 | 7.11434E-05 | 0.5 | 0.00185594 | FALSE | 0 | relative convergence (4) | 3303.30155794274 | 3358.54394205981 | -1644.65077897137 | 3289.30155794274 | 19760 | 0.033078264 |
| Skiv2l | -11.09748296 | 0.48129624 | 0.187830072 | 0.588305356 | 0.166752506 | 0.000223727 | 0.5 | 4.39923E-07 | FALSE | 0 | relative convergence (4) | 10195.3906143111 | 10250.6329984282 | -5090.69530715556 | 10181.3906143111 | 19760 | 5.56062E-05 |
| Kmt5c | -11.77476125 | 0.347093305 | 0.305669686 | 0.519599706 | 0.32296054 | 0.22709315 | 0.5 | 0.000570412 | FALSE | 0 | relative convergence (4) | NA | NA | NA | NA | 19760 | 0.014816861 |
| Zfp628 | -12.02486327 | 0.364900842 | 0.454360321 | 0.59703519 | 0.239937447 | 0.326928143 | 0.5 | 0.000943397 | FALSE | 0 | relative convergence (4) | 4933.8742844137 | 4989.11666853078 | -2459.93714220685 | 4919.8742844137 | 19760 | 0.020957009 |
| Inafm1 | -10.99674565 | 0.346337556 | 0.412221834 | 0.675642222 | 0.237247225 | 0.20738017 | 0.5 | 3.5276E-07 | FALSE | 0 | relative convergence (4) | NA | NA | NA | NA | 19760 | 4.69356E-05 |
| Bbc3 | -12.03970329 | 0.960737162 | 0.333825704 | 0.258764473 | 0.993585018 | 0.000175634 | 0.5 | 0.001489683 | FALSE | 0 | relative convergence (4) | 6672.0646826237 | 6727.30706674077 | -3329.03234131185 | 6658.0646826237 | 19760 | 0.028659961 |
| Abhd4 | -11.17309106 | 0.2503303 | -0.474636801 | 0.019731283 | 0.018314525 | 1.14529E-06 | 0.5 | 0.002596209 | FALSE | 0 | relative convergence (4) | 8250.07208978462 | 8305.3144739017 | -4118.03604489231 | 8236.07208978462 | 19760 | 0.041539338 |
| Lrp10 | -9.704702043 | 0.116279528 | 0.050791654 | 0.209015384 | 0.079373145 | 0.045168148 | 0.5 | 0.0022007 | FALSE | 0 | relative convergence (4) | NA | NA | NA | NA | 19760 | 0.0371883 |
| Gm4022 | -13.01328152 | 0.753756143 | 0.288526238 | 0.265914172 | 0.488352202 | 0.468715461 | 0.5 | 0.001769214 | FALSE | 0 | relative convergence (4) | 2764.85008272504 | 2820.09246684212 | -1375.42504136252 | 2750.85008272504 | 19760 | 0.032456986 |
| Ifngr2 | -10.15056146 | 0.424321194 | 0.072813847 | 0.371422686 | 0.223491416 | 0.078363959 | 0.5 | 1.09374E-10 | FALSE | 0 | relative convergence (4) | 18168.5334321854 | 18223.7758163025 | -9077.26671609272 | 18154.5334321854 | 19760 | 5.52997E-08 |
| Ets2 | -10.0599749 | 0.207606336 | -0.033976046 | -0.00746686 | 0.340607739 | 0.087350537 | 0.5 | 7.76773E-06 | FALSE | 0 | relative convergence (4) | NA | NA | NA | NA | 19760 | 0.000564277 |
| Tmem181a | -11.65430619 | 0.379089124 | 0.2102639 | 0.247361525 | 0.823048837 | 0.06401963 | 0.5 | 0.000453316 | FALSE | 0 | relative convergence (4) | NA | NA | NA | NA | 19760 | 0.012768105 |
| Tmem181b-ps | -12.35280854 | 0.546872306 | 0.193058096 | 0.04851744 | 2.01685015 | 0.099354259 | 0.5 | 0.000131368 | FALSE | 0 | relative convergence (4) | 3822.4359886352 | 3877.67837275227 | -1904.2179943176 | 3808.4359886352 | 19760 | 0.005133354 |
| Slc22a1 | -11.97979087 | 0.810617628 | 0.886699533 | 0.720451452 | 2.822888388 | 0.000151561 | 0.5 | 0.002316517 | FALSE | 0 | relative convergence (4) | 6714.04460969421 | 6769.28699381128 | -3350.0223048471 | 6700.04460969421 | 19760 | 0.038503205 |
| Smoc2 | -12.7687633 | 1.253097673 | -0.49693502 | 0.709622941 | 10.74048933 | 0.467758207 | 0.5 | 4.1564E-16 | FALSE | 0 | relative convergence (4) | NA | NA | NA | NA | 19760 | 4.37808E-13 |
| Dynlt2a2 | -10.65390177 | 0.379200883 | 0.111700954 | 0.35934314 | 0.524726364 | 0.000139058 | 0.5 | 0.000929749 | FALSE | 0 | relative convergence (4) | 13528.8831142695 | 13584.1254983866 | -6757.44155713475 | 13514.8831142695 | 19760 | 0.020763307 |
| Cds1 | -12.38350384 | 0.414059472 | 0.142486412 | -0.141747771 | 3.807236284 | 0.06474156 | 0.5 | 0.002193172 | FALSE | 0 | relative convergence (4) | NA | NA | NA | NA | 19760 | 0.037110703 |
| Sparcl1 | -8.141120167 | 0.518137146 | 0.492189182 | 0.417834854 | 1.339055509 | 0.211577271 | 0.5 | 0.000577366 | FALSE | 0 | relative convergence (4) | 57599.9879924724 | 57655.2303765894 | -28792.9939962362 | 57585.9879924724 | 19760 | 0.014863345 |
| Crlf2 | -10.73475336 | 0.493968818 | -0.10248243 | 0.521706121 | 0.619709347 | 0.000107847 | 0.5 | 0.001781865 | FALSE | 0 | relative convergence (4) | 12655.8776504057 | 12711.1200345228 | -6320.93882520287 | 12641.8776504057 | 19760 | 0.032507478 |
| Crybb3 | -12.67421501 | 1.034374157 | 0.096426994 | 1.038142433 | 3.283326707 | 0.408810717 | 0.5 | 1.7333E-05 | FALSE | 0 | relative convergence (4) | 4198.49473386026 | 4253.73711797733 | -2092.24736693013 | 4184.49473386026 | 19760 | 0.001100949 |
| 2900026A02Rik | -10.00523078 | 0.972535444 | 0.33114862 | 0.472117598 | 0.531456982 | 0.000441076 | 0.5 | 2.31445E-09 | FALSE | 0 | relative convergence (4) | 23111.863639155 | 23167.106023272 | -11548.9318195775 | 23097.863639155 | 19760 | 7.13527E-07 |
| Kctd10 | -9.305942223 | 0.362140801 | 0.075695691 | 0.409087419 | 0.148851158 | 0.000176162 | 0.5 | 0.003345231 | FALSE | 0 | relative convergence (4) | 29312.6035438401 | 29367.8459279572 | -14649.3017719201 | 29298.6035438401 | 19760 | 0.048271342 |
| Cabp1 | -12.20472771 | 0.637656645 | 0.402093065 | 0.591333834 | 9.030755794 | 0.279704921 | 0.5 | 0.000795314 | FALSE | 0 | relative convergence (4) | 4247.38416428494 | 4302.62654840202 | -2116.69208214247 | 4233.38416428494 | 19760 | 0.018650788 |
| Rtf2 | -10.21555212 | 0.139089327 | 0.23345909 | 0.266580313 | 5.88545E-08 | 0.056520616 | 0.5 | 0.003314116 | FALSE | 1 | false convergence (8) | 16621.0329468564 | 16676.2753309734 | -8303.51647342818 | 16607.0329468564 | 19760 | 0.04814991 |
| Prelid3b | -11.17579951 | 0.410584537 | -0.050078829 | 0.416658531 | 0.05699857 | 0.11995582 | 0.5 | 6.62774E-06 | FALSE | 0 | relative convergence (4) | 9210.45653515203 | 9265.6989192691 | -4598.22826757601 | 9196.45653515203 | 19760 | 0.000501645 |
| Phactr3 | -12.59529968 | 0.435117384 | 0.402585446 | 0.190031376 | 2.841491661 | 0.434656738 | 0.5 | 0.003205879 | FALSE | 0 | relative convergence (4) | NA | NA | NA | NA | 19760 | 0.047617281 |
| Flywch1 | -10.80543476 | 0.215345664 | 0.215801429 | 0.310603374 | 0.279477397 | 0.129263585 | 0.5 | 0.001401556 | FALSE | 0 | relative convergence (4) | 11737.06511339 | 11792.3074975071 | -5861.532556695 | 11723.06511339 | 19760 | 0.027509351 |
| Tmem204 | -11.85518829 | 0.512653805 | -0.127430456 | -0.168675388 | 2.513030871 | 0.268062911 | 0.5 | 5.7016E-06 | FALSE | 0 | relative convergence (4) | 5323.08366284082 | 5378.32604695789 | -2654.54183142041 | 5309.08366284082 | 19760 | 0.000453259 |
| Lemd2 | -10.92565753 | 0.336025938 | 0.160202558 | 0.642449254 | 7.40413E-07 | 0.147282935 | 0.5 | 0.000159116 | FALSE | 1 | false convergence (8) | 11080.9105049873 | 11136.1528891043 | -5533.45525249363 | 11066.9105049873 | 19760 | 0.006025079 |
| Nudt3 | -10.70279878 | 0.314784403 | -0.118119556 | 0.288666028 | 0.116045541 | 8.65521E-05 | 0.5 | 0.000181716 | FALSE | 0 | relative convergence (4) | 12445.4323188221 | 12500.6747029392 | -6215.71615941106 | 12431.4323188221 | 19760 | 0.006676989 |
| Erf | -11.04219332 | 0.935253451 | 0.025948355 | 0.486903351 | 0.292240675 | 0.142589444 | 0.5 | 0.001338945 | FALSE | 0 | relative convergence (4) | 12383.2980600681 | 12438.5404441852 | -6184.64903003407 | 12369.2980600681 | 19760 | 0.026855569 |
| 2310022A10Rik | -11.33061019 | 0.273651412 | -0.150653042 | 0.157359517 | 0.275635681 | 0.118456828 | 0.5 | 0.001107722 | FALSE | 0 | relative convergence (4) | 7583.43265635959 | 7638.67504047666 | -3784.71632817979 | 7569.43265635959 | 19760 | 0.02345328 |
| Zfp607b | -12.87194494 | 0.607250302 | 0.012054395 | 0.675735409 | 1.86834583 | 1.45608E-13 | 0.5 | 0.000305109 | FALSE | 0 | relative convergence (4) | 2695.89889750504 | 2751.14128162211 | -1340.94944875252 | 2681.89889750504 | 19760 | 0.00964145 |
| Samd4b | -9.909902656 | 0.168746857 | 0.042063956 | 0.005527631 | 0.110065185 | 0.075497424 | 0.5 | 0.00252111 | FALSE | 0 | relative convergence (4) | 19575.3428630058 | 19630.5852471229 | -9780.67143150291 | 19561.3428630058 | 19760 | 0.04085491 |
| Sirt2 | -9.824930287 | 0.147907182 | 0.42071745 | 0.302018934 | 0.094908456 | 0.156016029 | 0.5 | 0.002992767 | FALSE | 0 | relative convergence (4) | 21200.6630489255 | 21255.9054330426 | -10593.3315244627 | 21186.6630489255 | 19760 | 0.045964239 |
| Fam98c | -11.25072423 | 0.498530308 | 0.130277267 | 0.243948949 | 0.135895045 | 0.000100112 | 0.5 | 0.000131583 | FALSE | 0 | relative convergence (4) | 8996.2553581196 | 9051.49774223667 | -4491.1276790598 | 8982.2553581196 | 19760 | 0.005133354 |
| Aplp1 | -11.37984028 | 0.573413827 | 0.049413695 | 0.325327337 | 0.615668163 | 0.000317553 | 0.5 | 0.002479313 | FALSE | 0 | relative convergence (4) | 8366.83724801208 | 8422.07963212915 | -4176.41862400604 | 8352.83724801208 | 19760 | 0.0404368 |
| Szrd1 | -10.10487357 | 0.270414901 | -0.015725723 | 0.214963729 | 0.053146754 | 0.000670087 | 0.5 | 0.000202297 | FALSE | 0 | relative convergence (4) | 18057.9688924762 | 18113.2112765933 | -9021.98444623811 | 18043.9688924762 | 19760 | 0.00716257 |
| Fhad1 | -12.52847949 | 0.636894762 | 0.540329451 | 0.080102194 | 5.190860783 | 0.205902271 | 0.5 | 0.003425422 | FALSE | 0 | relative convergence (4) | 3351.95287640528 | 3407.19526052236 | -1668.97643820264 | 3337.95287640528 | 19760 | 0.048805208 |
| Pgd | -10.69730765 | 0.368033344 | 0.097294251 | 0.294249553 | 0.381763426 | 0.121207981 | 0.5 | 0.002335232 | FALSE | 0 | relative convergence (4) | 12667.431831177 | 12722.6742152941 | -6326.71591558851 | 12653.431831177 | 19760 | 0.038690898 |
| Eno1 | -9.604407068 | 0.535586073 | 0.296459138 | 0.157674336 | 0.118986745 | 0.209446734 | 0.5 | 1.92449E-07 | FALSE | 0 | relative convergence (4) | 24716.1480002162 | 24771.3903843332 | -12351.0740001081 | 24702.1480002162 | 19760 | 2.70284E-05 |
| Icmt | -11.3206843 | 0.272831742 | 0.18207216 | 0.663842252 | 5.81303E-07 | 0.197679974 | 0.5 | 0.00081269 | FALSE | 1 | false convergence (8) | NA | NA | NA | NA | 19760 | 0.018917867 |
| Vwa1 | -12.09858192 | 0.765426191 | 0.370788589 | 0.789464954 | 1.804456267 | 0.266510741 | 0.5 | 1.54365E-06 | FALSE | 0 | relative convergence (4) | 5575.50685506032 | 5630.74923917739 | -2780.75342753016 | 5561.50685506032 | 19760 | 0.000152436 |
| Clptm1l | -10.0667487 | 0.243299763 | 0.067871743 | 0.263861627 | 0.009363856 | 2.51875E-05 | 0.5 | 0.001212471 | FALSE | 0 | relative convergence (4) | 18300.497479296 | 18355.7398634131 | -9143.248739648 | 18286.497479296 | 19760 | 0.025041877 |
| Cast | -8.670320209 | 0.152452656 | 0.027730742 | 0.067135923 | 0.153570949 | 0.100603867 | 0.5 | 2.29089E-09 | FALSE | 0 | relative convergence (4) | 39742.4103399737 | 39797.6527240907 | -19864.2051699868 | 39728.4103399737 | 19760 | 7.13527E-07 |
| Tmem250-ps | -10.88980769 | 0.330405802 | -0.00659879 | 0.276110318 | 0.077505914 | 6.03461E-05 | 0.5 | 0.001368194 | FALSE | 0 | relative convergence (4) | 10672.5626003608 | 10727.8049844779 | -5329.28130018042 | 10658.5626003608 | 19760 | 0.027149094 |
| Acan | -12.75923962 | 2.101997876 | 0.084762199 | 0.035305011 | 4.826795167 | 0.139695793 | 0.5 | 2.1081E-08 | FALSE | 0 | relative convergence (4) | 4494.64415169436 | 4549.88653581144 | -2240.32207584718 | 4480.64415169436 | 19760 | 5.08105E-06 |
| Leprotl1 | -10.29269819 | 0.264699961 | 0.078522302 | 0.083409471 | 0.044151482 | 0.000753856 | 0.5 | 0.003396665 | FALSE | 0 | relative convergence (4) | 15856.3250103535 | 15911.5673944706 | -7921.16250517677 | 15842.3250103535 | 19760 | 0.048622696 |
| Naf1 | -10.59765995 | 0.495041328 | 0.373360958 | 0.484841956 | 0.468018015 | 0.120100641 | 0.5 | 6.5041E-06 | FALSE | 0 | relative convergence (4) | 14704.8100005194 | 14760.0523846364 | -7345.40500025969 | 14690.8100005194 | 19760 | 0.000495252 |
| Vcp | -9.111860502 | 0.344300286 | 0.089368154 | 0.380847187 | 0.07100832 | 0.222888282 | 0.5 | 0.000363153 | FALSE | 0 | relative convergence (4) | 32285.722278729 | 32340.9646628461 | -16135.8611393645 | 32271.722278729 | 19760 | 0.010929185 |
| Fam214b | -10.493014 | 0.506913422 | -0.015596975 | 0.125912014 | 0.350642584 | 0.000234835 | 0.5 | 0.001203172 | FALSE | 0 | relative convergence (4) | 15288.0221083421 | 15343.2644924592 | -7637.01105417105 | 15274.0221083421 | 19760 | 0.024890502 |
| Fam124a | -11.45384184 | 0.28931452 | 0.500217299 | 0.348486024 | 2.008509564 | 0.197685371 | 0.5 | 0.003126692 | FALSE | 0 | relative convergence (4) | 7601.07183479808 | 7656.31421891515 | -3793.53591739904 | 7587.07183479808 | 19760 | 0.046881829 |
| Mtmr9 | -10.84809987 | 0.215463759 | -0.097442094 | 0.223808275 | 0.105400905 | 9.06772E-15 | 0.5 | 0.000236743 | FALSE | 0 | relative convergence (4) | 10827.8342553678 | 10883.0766394848 | -5406.91712768388 | 10813.8342553678 | 19760 | 0.007979828 |
| Dock5 | -11.19142372 | 0.652518768 | 0.591538784 | 0.622587943 | 1.898006181 | 0.23347803 | 0.5 | 2.21519E-16 | FALSE | 0 | relative convergence (4) | 9924.94566370603 | 9980.1880478231 | -4955.47283185301 | 9910.94566370603 | 19760 | 2.80001E-13 |
| Loxl2 | -11.32130442 | 0.412320025 | 0.268351293 | 0.308025387 | 0.724488382 | 0.372516307 | 0.5 | 0.00153052 | FALSE | 0 | relative convergence (4) | 8047.76682381062 | 8103.00920792769 | -4016.88341190531 | 8033.76682381062 | 19760 | 0.029189644 |
| Bmp1 | -10.24906059 | 0.246758953 | 0.221264355 | 0.321368531 | 0.449519358 | 0.33856278 | 0.5 | 0.001681425 | FALSE | 0 | relative convergence (4) | 16336.5140443551 | 16391.7564284721 | -8161.25702217753 | 16322.5140443551 | 19760 | 0.031439656 |
| Slc12a4 | -10.35175947 | 0.404653112 | -0.107686091 | 0.240979476 | 0.235839459 | 0.000189724 | 0.5 | 0.002680139 | FALSE | 0 | relative convergence (4) | 15738.124846255 | 15793.367230372 | -7862.06242312749 | 15724.124846255 | 19760 | 0.042293334 |
| Bcar1 | -11.31834315 | 0.773173529 | -0.179180149 | 0.531412674 | 0.651075438 | 0.319948891 | 0.5 | 0.000665847 | FALSE | 0 | relative convergence (4) | 10008.2370290505 | 10063.4794131675 | -4997.11851452523 | 9994.23702905046 | 19760 | 0.016318899 |
| Abhd16a | -10.52742599 | 0.318113135 | 0.251109275 | 0.65162478 | 0.143804408 | 0.201401552 | 0.5 | 8.09991E-05 | FALSE | 0 | relative convergence (4) | 14336.7966691503 | 14392.0390532674 | -7161.39833457516 | 14322.7966691503 | 19760 | 0.003542659 |
| Bag6 | -10.07434555 | 0.307332277 | 0.247166793 | 0.525914455 | 0.015580198 | 0.166364835 | 0.5 | 0.000133907 | FALSE | 0 | relative convergence (4) | 18613.0431637401 | 18668.2855478571 | -9299.52158187003 | 18599.0431637401 | 19760 | 0.005191988 |
| Ptchd4 | -12.01475434 | 0.624464007 | 0.586810472 | 0.696857951 | 25.037666 | 0.171453876 | 0.5 | 0.000334348 | FALSE | 0 | relative convergence (4) | 4677.15788277481 | 4732.40026689189 | -2331.57894138741 | 4663.15788277481 | 19760 | 0.010307706 |
| Vegfa | -11.8147747 | 1.19203263 | -0.378474218 | 0.308834515 | 1.232683588 | 0.15459495 | 0.5 | 5.54654E-06 | FALSE | 0 | relative convergence (4) | 7786.4779848672 | 7841.72036898428 | -3886.2389924336 | 7772.4779848672 | 19760 | 0.000443724 |
| Klhdc3 | -10.72451365 | 0.467248281 | -0.060932994 | 0.011626657 | 0.036088811 | 0.092284714 | 0.5 | 7.01787E-09 | FALSE | 0 | relative convergence (4) | 12099.5075197185 | 12154.7499038356 | -6042.75375985924 | 12085.5075197185 | 19760 | 1.81033E-06 |
| Egln1 | -9.638214042 | 0.118561584 | 0.334629536 | 0.416600966 | 0.113929937 | 0.153556106 | 0.5 | 0.001792307 | FALSE | 0 | relative convergence (4) | NA | NA | NA | NA | 19760 | 0.032596778 |
| Olfm2 | -13.5696754 | 1.192554029 | -0.129880309 | 0.056305353 | 14.46498767 | 0.750043186 | 0.5 | 0.000676159 | FALSE | 0 | relative convergence (4) | 2440.88928534327 | 2496.13166946035 | -1213.44464267164 | 2426.88928534327 | 19760 | 0.016473588 |
| Ppan | -11.65663033 | 0.325576847 | -0.031968232 | 0.472491271 | 0.319173769 | 0.066292745 | 0.5 | 0.000430678 | FALSE | 0 | relative convergence (4) | NA | NA | NA | NA | 19760 | 0.012372213 |
| Keap1 | -10.16297217 | 0.201732355 | 0.159399201 | 0.326010498 | 0.121082021 | 0.077217427 | 0.5 | 1.62135E-05 | FALSE | 0 | relative convergence (4) | 17200.9984332512 | 17256.2408173683 | -8593.49921662562 | 17186.9984332512 | 19760 | 0.001045605 |
| Yipf2 | -12.14629184 | 0.425475956 | -0.061940527 | 0.29211036 | 0.390261153 | 0.003115378 | 0.5 | 4.95731E-05 | FALSE | 0 | relative convergence (4) | 4324.26205386712 | 4379.5044379842 | -2155.13102693356 | 4310.26205386712 | 19760 | 0.002466945 |
| Ldlr | -10.41490529 | 0.343623674 | -0.703478328 | 0.021290563 | 1.89400614 | 0.305397962 | 0.5 | 0.003445949 | FALSE | 0 | relative convergence (4) | 15513.1603189398 | 15568.4027030569 | -7749.58015946992 | 15499.1603189398 | 19760 | 0.048995276 |
| Prkcsh | -10.26735588 | 0.248703002 | 0.319573264 | 0.613444242 | 0.063698551 | 0.189297854 | 0.5 | 1.35065E-05 | FALSE | 0 | relative convergence (4) | 16566.6979277883 | 16621.9403119054 | -8276.34896389417 | 16552.6979277883 | 19760 | 0.000889179 |
| Cldn3 | -13.47847483 | 1.367359583 | 0.292126803 | 0.671912567 | 6.945703813 | 0.000267397 | 0.5 | 0.000264393 | FALSE | 0 | relative convergence (4) | 2296.97778349101 | 2352.22016760809 | -1141.48889174551 | 2282.97778349101 | 19760 | 0.008689634 |
| Stx1a | -10.61762026 | 0.591723814 | 0.614385771 | 0.305274865 | 1.116452598 | 0.000138749 | 0.5 | 4.6724E-05 | FALSE | 0 | relative convergence (4) | 14028.1503009889 | 14083.392685106 | -7007.07515049446 | 14014.1503009889 | 19760 | 0.002352952 |
| Hip1 | -8.350157358 | 0.387164064 | 0.42166802 | 0.379135547 | 0.482154998 | 0.125726085 | 0.5 | 1.92345E-05 | FALSE | 0 | relative convergence (4) | 49404.5048585071 | 49459.7472426241 | -24695.2524292535 | 49390.5048585071 | 19760 | 0.001203584 |
| Tmem120a | -11.40190404 | 0.565961626 | 0.314257633 | 0.399984901 | 0.588499642 | 0.179536709 | 0.5 | 0.000110786 | FALSE | 0 | relative convergence (4) | 8347.76145693576 | 8403.00384105284 | -4166.88072846788 | 8333.76145693576 | 19760 | 0.004495057 |
| Serpine1 | -9.659812701 | 1.093746885 | 0.19446127 | 0.414405831 | 2.280247332 | 0.10368921 | 0.5 | 0.001458299 | FALSE | 0 | relative convergence (4) | 28664.9517930035 | 28720.1941771206 | -14325.4758965018 | 28650.9517930035 | 19760 | 0.028271316 |
| Cnpy4 | -11.47381057 | 0.334882208 | 0.312436639 | 0.434720304 | 0.069442142 | 0.257601545 | 0.5 | 7.95304E-05 | FALSE | 0 | relative convergence (4) | 7274.92318348779 | 7330.16556760486 | -3630.46159174389 | 7260.92318348779 | 19760 | 0.003514912 |
| Pdgfa | -9.869795107 | 0.572422499 | 0.337891442 | 0.255682169 | 0.203238213 | 0.000364447 | 0.5 | 0.002110865 | FALSE | 0 | relative convergence (4) | 22915.0536950354 | 22970.2960791525 | -11450.5268475177 | 22901.0536950354 | 19760 | 0.036153564 |
| Gpr146 | -10.98375319 | 0.476065795 | 0.352563228 | 0.005910076 | 0.296935643 | 0.000120119 | 0.5 | 0.001261523 | FALSE | 0 | relative convergence (4) | 10133.4924680657 | 10188.7348521828 | -5059.74623403285 | 10119.4924680657 | 19760 | 0.025843835 |
| Micall2 | -12.92891792 | 0.986264889 | 0.123786514 | -0.307301379 | 2.844119844 | 0.224066124 | 0.5 | 0.000914771 | FALSE | 0 | relative convergence (4) | 3088.05636997274 | 3143.29875408981 | -1537.02818498637 | 3074.05636997274 | 19760 | 0.020574218 |
| Mafk | -10.64444526 | 0.790467626 | 0.057950739 | 0.369586214 | 0.380901976 | 0.251444097 | 0.5 | 4.64801E-05 | FALSE | 0 | relative convergence (4) | 15518.7061303773 | 15573.9485144944 | -7752.35306518867 | 15504.7061303773 | 19760 | 0.002352952 |
| Plbd2 | -10.59610981 | 0.49260973 | -0.039433153 | 0.272177128 | 0.092498208 | 5.57881E-05 | 0.5 | 1.74699E-07 | FALSE | 0 | relative convergence (4) | 13432.8139967724 | 13488.0563808894 | -6709.40699838618 | 13418.8139967724 | 19760 | 2.50931E-05 |
| Cyb5r3 | -8.657024601 | 0.60060084 | 0.044100764 | 0.368306152 | 0.289933767 | 0.000476487 | 0.5 | 0.000746025 | FALSE | 0 | relative convergence (4) | 43392.9047041435 | 43448.1470882606 | -21689.4523520718 | 43378.9047041435 | 19760 | 0.017691841 |
| Pim3 | -11.2847475 | 0.936274828 | -0.229930092 | 0.340597859 | 0.424329559 | 0.000328822 | 0.5 | 0.003292734 | FALSE | 0 | relative convergence (4) | 11727.4139804975 | 11782.6563646146 | -5856.70699024876 | 11713.4139804975 | 19760 | 0.047991161 |
| Plxnb2 | -10.12468411 | 0.530536785 | 0.373876868 | 0.692017505 | 0.227784158 | 0.22835287 | 0.5 | 0.000255105 | FALSE | 0 | relative convergence (4) | 19181.0413565112 | 19236.2837406283 | -9583.5206782556 | 19167.0413565112 | 19760 | 0.008493035 |
| Tmem117 | -11.25173037 | 0.540328903 | -0.021503354 | 0.080110201 | 4.609450109 | 0.10596692 | 0.5 | 4.84342E-09 | FALSE | 0 | relative convergence (4) | 8409.52954078235 | 8464.77192489942 | -4197.76477039117 | 8395.52954078235 | 19760 | 1.30257E-06 |
| Mcrs1 | -10.79156898 | 0.375154422 | -0.061934687 | 0.373044751 | 1.21614E-07 | 1.01856E-33 | 0.5 | 0.002680108 | FALSE | 0 | relative convergence (4) | NA | NA | NA | NA | 19760 | 0.042293334 |
| Fmnl3 | -10.28054931 | 0.287653326 | 0.132330734 | 0.302443706 | 0.565300353 | 0.033327216 | 0.5 | 0.001398459 | FALSE | 0 | relative convergence (4) | 16706.7532395985 | 16761.9956237156 | -8346.37661979927 | 16692.7532395985 | 19760 | 0.027509351 |
| Cycs | -10.53815238 | 0.579753061 | 0.386888532 | 0.547977425 | 0.070438752 | 0.257822931 | 0.5 | 0.000729648 | FALSE | 0 | relative convergence (4) | 15641.8830859137 | 15697.1254700308 | -7813.94154295685 | 15627.8830859137 | 19760 | 0.017382746 |
| Prr15 | -12.88964746 | 0.804107328 | 0.637647379 | 0.463528345 | 3.170870001 | 0.212543368 | 0.5 | 0.000501374 | FALSE | 0 | relative convergence (4) | 3051.45460869516 | 3106.69699281224 | -1518.72730434758 | 3037.45460869516 | 19760 | 0.013570369 |
| Fkbp14 | -11.10247218 | 0.313992198 | 0.145296241 | 0.414919795 | 0.454877226 | 0.175713961 | 0.5 | 2.43294E-05 | FALSE | 0 | relative convergence (4) | 9169.83576041446 | 9225.07814453154 | -4577.91788020723 | 9155.83576041446 | 19760 | 0.001417162 |
| Rnf181 | -11.20409548 | 0.408246544 | -0.01833988 | 0.406388688 | 0.034165153 | 1.37771E-08 | 0.5 | 0.0017554 | FALSE | 0 | relative convergence (4) | 9098.00328086243 | 9153.2456649795 | -4542.00164043121 | 9084.00328086243 | 19760 | 0.032278359 |
| Dok1 | -11.53683271 | 0.604022358 | 0.286470311 | 0.636073543 | 0.07246768 | 0.184273436 | 0.5 | 3.42074E-08 | FALSE | 0 | relative convergence (4) | 7408.8696730525 | 7464.11205716958 | -3697.43483652625 | 7394.8696730525 | 19760 | 7.08823E-06 |
| Bscl2 | -10.56577929 | 0.357371199 | -0.196014718 | 0.308608936 | 0.157113583 | 0.000108624 | 0.5 | 9.17763E-05 | FALSE | 0 | relative convergence (4) | 13679.0961769999 | 13734.338561117 | -6832.54808849997 | 13665.0961769999 | 19760 | 0.003905901 |
| Fads3 | -10.46327691 | 0.476948253 | 0.178194809 | 0.201184023 | 0.264750711 | 0.169615637 | 0.5 | 0.000717223 | FALSE | 0 | relative convergence (4) | 15280.9480175626 | 15336.1904016797 | -7633.47400878131 | 15266.9480175626 | 19760 | 0.017202474 |
| Ddb1 | -9.379378582 | 0.326199065 | 0.095018227 | 0.363635695 | 0.08700485 | 0.114384724 | 0.5 | 4.30427E-05 | FALSE | 0 | relative convergence (4) | 27711.9924850255 | 27767.2348691426 | -13848.9962425127 | 27697.9924850255 | 19760 | 0.002202673 |
| Tmem132a | -11.08816501 | 0.713210994 | 0.078528278 | 0.549982286 | 0.700711183 | 0.290124457 | 0.5 | 0.000341407 | FALSE | 0 | relative convergence (4) | 10577.8618691541 | 10633.1042532712 | -5281.93093457704 | 10563.8618691541 | 19760 | 0.010476057 |
| Prpf19 | -10.17305633 | 0.215991492 | 0.316157931 | 0.505230609 | 0.017143752 | 0.120496358 | 0.5 | 3.52861E-06 | FALSE | 0 | relative convergence (4) | 17414.9465899604 | 17470.1889740775 | -8700.47329498019 | 17400.9465899604 | 19760 | 0.000309191 |
| Pcsk5 | -10.17453138 | 1.085772118 | 0.406230125 | 0.523780369 | 2.382498693 | 0.000184276 | 0.5 | 5.92348E-08 | FALSE | 0 | relative convergence (4) | 21446.0879606806 | 21501.3303447977 | -10716.0439803403 | 21432.0879606806 | 19760 | 1.06961E-05 |
| Anxa1 | -9.746705295 | 0.664940638 | 0.164131248 | 0.718668667 | 0.553321655 | 0.134684041 | 0.5 | 0.000221436 | FALSE | 0 | relative convergence (4) | 25650.4059226707 | 25705.6483067877 | -12818.2029613353 | 25636.4059226707 | 19760 | 0.007626583 |
| Zfand5 | -8.871788318 | 0.37264545 | 0.027607549 | -0.033949214 | 0.263326649 | 0.00011464 | 0.5 | 0.000602747 | FALSE | 0 | relative convergence (4) | 37701.5315728514 | 37756.7739569685 | -18843.7657864257 | 37687.5315728514 | 19760 | 0.015298645 |
| Tjp2 | -10.20368328 | 0.280354009 | 0.045046959 | 0.096111959 | 0.958169976 | 0.106148636 | 0.5 | 0.000686488 | FALSE | 0 | relative convergence (4) | 17012.2344158477 | 17067.4767999647 | -8499.11720792383 | 16998.2344158477 | 19760 | 0.016686934 |
| Lrrc59 | -10.19367282 | 0.203688872 | -0.167393279 | 0.072971478 | 0.108362177 | 0.075268259 | 0.5 | 1.66325E-05 | FALSE | 0 | relative convergence (4) | NA | NA | NA | NA | 19760 | 0.001067181 |
| Ppp1r9b | -10.22018312 | 0.307414894 | -0.170622724 | 0.228628952 | 0.177099623 | 0.000148144 | 0.5 | 0.002432886 | FALSE | 0 | relative convergence (4) | 16466.8311663884 | 16522.0735505055 | -8226.4155831942 | 16452.8311663884 | 19760 | 0.03988544 |

| | | | | | | | | | | | | | | | | |
|---|---|---|---|---|---|---|---|---|---|---|---|---|---|---|---|---|
| Comp | -12.86358651 | 0.885179146 | -0.178478429 | 0.722345116 | 25.71390422 | 0.195365401 | 0.5 | 3.10513E-06 | FALSE | 0relative convergence (4) | 2604.70977751516 | 2659.95216163223 | -1295.35488875758 | 2590.70977751516 | 19760 | 0.000284412 |
| Fam32a | -10.00471173 | 0.223525979 | 0.004824246 | 0.355821371 | 0.012614464 | 0.070854817 | 0.5 | 1.60285E-07 | FALSE | 0relative convergence (4) | NA | NA | NA | NA | 19760 | 2.32874E-05 |
| Slc35e1 | -11.11789216 | 0.271911824 | 0.00933537 | 0.377787414 | 0.166940001 | 3.29508E-10 | 0.5 | 3.70346E-05 | FALSE | 0relative convergence (4) | 9305.64448927437 | 9360.88687339144 | -4645.82224463718 | 9291.64448927437 | 19760 | 0.001991987 |
| Apbb2 | -10.04694314 | 0.332181842 | 0.351492485 | 0.211579859 | 0.891388878 | 0.146232017 | 0.5 | 1.44887E-07 | FALSE | 0relative convergence (4) | 18762.619808683 | 18817.8621928001 | -9374.30990434151 | 18748.619808683 | 19760 | 2.12951E-05 |
| Utp3 | -10.30418252 | 0.217203737 | 0.315753898 | 0.288647595 | 0.121402636 | 0.0162764 | 0.5 | 9.61774E-06 | FALSE | 0relative convergence (4) | 15946.4850930682 | 16001.7274771852 | -7966.24254653409 | 15932.4850930682 | 19760 | 0.000664307 |
| Ereg | -12.47490476 | 0.528317748 | 0.414585657 | 0.567545061 | 5.949354069 | 0.409436373 | 0.5 | 0.000360176 | FALSE | 0relative convergence (4) | 3952.58911505034 | 4007.83149916741 | -1969.29455752517 | 3938.58911505034 | 19760 | 0.010891437 |
| Art3 | -11.89903627 | 0.680341793 | 0.691694915 | 0.487980976 | 9.864776625 | 6.77386E-05 | 0.5 | 1.09296E-05 | FALSE | 0relative convergence (4) | 5802.89443650088 | 5858.13682061796 | -2894.44721825044 | 5788.89443650088 | 19760 | 0.000730954 |
| Anxa3 | -9.53731663 | 0.344469579 | 0.202410726 | 0.358987809 | 0.486153678 | 0.00014661 | 0.5 | 0.002518737 | FALSE | 0relative convergence (4) | 26658.847669599 | 26714.090053716 | -13322.4238347995 | 26644.847669599 | 19760 | 0.04085491 |
| Mir703 | -10.04291524 | 1.012422977 | -0.322862899 | 1.224510023 | 0.064213652 | 0.454254701 | 0.5 | 0.003368586 | FALSE | 0relative convergence (4) | 25738.3861243324 | 25793.6285084495 | -12862.1930621662 | 25724.3861243324 | 19760 | 0.048385151 |
| Bmp3 | -9.444813452 | 0.165123097 | 0.320934269 | 0.249302956 | 1.085847613 | 0.316668908 | 0.5 | 0.003116474 | FALSE | 0relative convergence (4) | 27998.6756259348 | 28053.9180100519 | -13992.3378129674 | 27984.6756259348 | 19760 | 0.04683975 |
| Srebf1 | -11.11111551 | 0.323126129 | -0.000172739 | 0.459809789 | 0.071320031 | 0.102534793 | 0.5 | 0.003295596 | FALSE | 0relative convergence (4) | 9540.97520667397 | 9596.21759079105 | -4763.48760333699 | 9526.97520667397 | 19760 | 0.047991161 |
| Flii | -9.574253944 | 0.194117846 | 0.271503576 | 0.43641829 | 0.056344875 | 0.152880142 | 0.5 | 5.45703E-08 | FALSE | 0relative convergence (4) | 24349.4802555081 | 24404.7226396252 | -12167.7401277541 | 24335.4802555081 | 19760 | 1.01437E-05 |
| Trim16 | -12.2025231 | 0.50955835 | -0.034693148 | -0.264484832 | 2.752820375 | 7.54542E-16 | 0.5 | 4.25643E-06 | FALSE | 1 singular convergence (7) | 4031.43817051644 | 4086.68055463352 | -2008.71908525822 | 4017.43817051644 | 19760 | 0.000361083 |
| Gm12295 | -10.76681521 | 0.476134065 | 0.439765797 | 0.419066445 | 4.116771526 | 0.290293537 | 0.5 | 2.77918E-09 | FALSE | 0relative convergence (4) | NA | NA | NA | NA | 19760 | 8.16948E-07 |
| Vamp2 | -9.981780445 | 0.375255146 | -0.043966184 | 0.378056088 | 0.093185587 | 0.039098994 | 0.5 | 0.000614867 | FALSE | 0relative convergence (4) | 20163.7404925856 | 20218.9828767026 | -10074.8702462928 | 20149.7404925856 | 19760 | 0.015513677 |
| Ankrd1 | -12.02497486 | 0.484712224 | 0.206026143 | 0.362169636 | 10.18628474 | 1.96535E-16 | 0.5 | 3.97426E-05 | FALSE | 0relative convergence (4) | 4699.56968507831 | 4754.81206919539 | -2342.78484253916 | 4685.56968507831 | 19760 | 0.002114008 |
| Myof | -9.134018328 | 0.200243849 | 0.309442934 | 0.160772811 | 0.36147729 | 0.157818578 | 0.5 | 5.3615E-05 | FALSE | 0relative convergence (4) | 31259.8705411051 | 31315.1129252222 | -15622.9352705525 | 31245.8705411051 | 19760 | 0.002613534 |
| Pdlim1 | -8.762048543 | 0.40642378 | -0.028320588 | 0.491633286 | 0.351795701 | 0.109130243 | 0.5 | 7.99686E-12 | FALSE | 0relative convergence (4) | 40582.1177354786 | 40637.3601195957 | -20284.0588677393 | 40568.1177354786 | 19760 | 4.59669E-09 |
| Sfxn3 | -10.19346014 | 0.272302636 | -0.335316711 | 0.049748369 | 0.111396107 | 0.15239629 | 0.5 | 0.003135068 | FALSE | 0relative convergence (4) | 16858.767093646 | 16914.0094777631 | -8422.383546823 | 16844.767093646 | 19760 | 0.046896167 |
| Zfp688 | -11.96350794 | 0.344046513 | 0.261282623 | 0.325482872 | 0.147697681 | 3.21779E-16 | 0.5 | 0.000417357 | FALSE | 0relative convergence (4) | NA | NA | NA | NA | 19760 | 0.012144681 |
| Armc5 | -11.56345042 | 0.484318891 | 0.210765303 | 0.351068796 | 0.384935322 | 0.000101623 | 0.5 | 0.00064464 | FALSE | 0relative convergence (4) | 7293.77655246385 | 7349.01893658092 | -3639.88827623192 | 7279.77655246385 | 19760 | 0.016008339 |
| Pwwp2b | -12.38538707 | 0.645296681 | 0.081429761 | 0.383453905 | 0.189186736 | 9.82721E-19 | 0.5 | 0.002272514 | FALSE | 0relative convergence (4) | 3910.82885203326 | 3966.07123615033 | -1948.41442601663 | 3896.82885203326 | 19760 | 0.038024376 |
| Ric8a | -10.7530645 | 0.211417937 | 0.021186472 | 0.386787249 | 0.045852124 | 0.071008621 | 0.5 | 0.000585301 | FALSE | 0relative convergence (4) | 11659.6877950169 | 11714.930179134 | -5822.84389750846 | 11645.6877950169 | 19760 | 0.014952615 |
| Pgghg | -11.35611206 | 0.296946563 | 0.044455951 | 0.395230112 | 0.454102169 | 0.120801901 | 0.5 | 0.000359981 | FALSE | 0relative convergence (4) | 7686.18733697445 | 7741.42972109152 | -3836.09366848722 | 7672.18733697445 | 19760 | 0.010891437 |
| Cend1 | -13.24238217 | 0.969172362 | 0.88983875 | 0.953181014 | 1.048699848 | 0.169670621 | 0.5 | 0.001026145 | FALSE | 0relative convergence (4) | 2662.68146816449 | 2717.92385228156 | -1324.34073408224 | 2648.68146816449 | 19760 | 0.0222927 |
| Tspan4 | -10.98976714 | 0.787987037 | 0.02642166 | 0.234385036 | 0.314709603 | 0.001620695 | 0.5 | 0.00017938 | FALSE | 0relative convergence (4) | 11384.0469151203 | 11439.2892992374 | -5685.02345756015 | 11370.0469151203 | 19760 | 0.00664915 |
| Ifitm10 | -12.63470106 | 0.875633034 | 0.617101526 | 0.26731228 | 2.18216208 | 8.3956E-05 | 0.5 | 0.000754973 | FALSE | 0relative convergence (4) | 3508.27774485583 | 3563.52012897291 | -1747.13887242792 | 3494.27774485583 | 19760 | 0.017870515 |
| Cdc42ep4 | -11.32172147 | 0.463393262 | 0.221092453 | 0.191351385 | 0.22273772 | 0.000155863 | 0.5 | 0.002548944 | FALSE | 0relative convergence (4) | 8570.68162905323 | 8625.9240131703 | -4278.34081452661 | 8556.68162905323 | 19760 | 0.041129195 |
| Trim47 | -9.867780713 | 0.474713245 | 0.097388419 | 0.234883371 | 0.304134286 | 0.000130941 | 0.5 | 0.001097958 | FALSE | 0relative convergence (4) | 21587.9283156116 | 21643.1706997287 | -10786.9641578058 | 21573.9283156116 | 19760 | 0.023295895 |
| Sphk1 | -13.38158978 | 1.363028612 | -1.38619555 | 0.066525338 | 4.475109295 | 0.000158356 | 0.5 | 0.000815497 | FALSE | 0relative convergence (4) | 2787.4081749063 | 2842.65055902337 | -1386.70408745315 | 2773.4081749063 | 19760 | 0.018941462 |
| Rhbdf2 | -11.07471593 | 0.588502591 | -0.152478552 | 0.248148374 | 0.867745647 | 0.282576818 | 0.5 | 0.001113698 | FALSE | 0relative convergence (4) | 11449.9040289582 | 11505.1464130753 | -5717.95201447912 | 11435.9040289582 | 19760 | 0.023540377 |
| Timp2 | -7.947142564 | 0.320660784 | -0.167913863 | 0.128876045 | 0.276607428 | 0.000117977 | 0.5 | 0.000497165 | FALSE | 0relative convergence (4) | 56247.1293605946 | 56302.3717447117 | -28116.5646802973 | 56233.1293605946 | 19760 | 0.013514343 |
| Slc38a10 | -10.1461181 | 0.257288741 | -0.045284648 | 0.37971212 | 0.109655052 | 0.145576583 | 0.5 | 2.97518E-08 | FALSE | 0relative convergence (4) | 17274.8853932099 | 17330.1277773269 | -8630.44269660493 | 17260.8853932099 | 19760 | 6.48383E-06 |
| Hgs | -10.92292562 | 0.24948882 | 0.020595238 | 0.229926007 | 2.39505E-08 | 3.7349E-54 | 0.5 | 2.80441E-05 | FALSE | 1 singular convergence (7) | NA | NA | NA | NA | 19760 | 0.001582488 |
| Arhgdia | -8.911957797 | 0.387918388 | -0.075235526 | 0.27086488 | 0.08098897 | 0.00010944 | 0.5 | 6.19248E-05 | FALSE | 0relative convergence (4) | 35768.729324222 | 35823.971708339 | -17877.364662111 | 35754.729324222 | 19760 | 0.002920631 |
| Mafg | -10.50360344 | 0.430957768 | 0.008223153 | 0.232584797 | 0.311766854 | 0.000144257 | 0.5 | 0.000262619 | FALSE | 0relative convergence (4) | 14475.299085349 | 14530.541469466 | -7230.64954267448 | 14461.299085349 | 19760 | 0.008667129 |
| Aspscr1 | -10.94182056 | 0.196763209 | -0.158743427 | 0.118178871 | 0.32051908 | 5.30342E-08 | 0.5 | 0.001726071 | FALSE | 0relative convergence (4) | 10098.3918155332 | 10153.6341996503 | -5042.1959077666 | 10084.3918155332 | 19760 | 0.032037491 |
| Slc16a3 | -11.85374135 | 0.988314385 | 0.155657947 | 0.111522575 | 0.656783638 | 0.000198115 | 0.5 | 0.002020016 | FALSE | 0relative convergence (4) | 7254.20493024467 | 7309.44731436174 | -3620.10246512233 | 7240.20493024467 | 19760 | 0.034928874 |
| Chrdl1 | -13.21396617 | 1.334344309 | 0.966526434 | 1.86143934 | 14.03412983 | 0.000123619 | 0.5 | 3.42916E-07 | FALSE | 0relative convergence (4) | 3098.69994206407 | 3153.94232618114 | -1542.34997103203 | 3084.69994206407 | 19760 | 4.61113E-05 |
| Pak3 | -11.60074405 | 0.483295022 | 0.154557198 | 0.01078674 | 10.0767353 | 0.00016122 | 0.5 | 0.002536942 | FALSE | 0relative convergence (4) | 6400.60821612742 | 6455.85060024449 | -3193.30410806371 | 6386.60821612742 | 19760 | 0.041058836 |
| Sat1 | -9.061803842 | 0.541558403 | 0.339596658 | 0.175750121 | 0.790739066 | 0.000395502 | 0.5 | 0.000973942 | FALSE | 0relative convergence (4) | 36674.3349202597 | 36729.5773043767 | -18330.1674601298 | 36660.3349202597 | 19760 | 0.021484515 |
| Map7d2 | -11.4176956 | 0.584756063 | 0.506228394 | 0.352233421 | 1.593599381 | 0.429779665 | 0.5 | 7.5789E-05 | FALSE | 0relative convergence (4) | 8762.83029722107 | 8818.07268133815 | -4374.41514861054 | 8748.83029722107 | 19760 | 0.003446511 |
| mt-Ts1 | -12.94478216 | 0.950998354 | 0.287687806 | 2.099489516 | 0.368325189 | 0.652857099 | 0.5 | 0.000585565 | FALSE | 0relative convergence (4) | 3948.02022699458 | 4003.26261111165 | -1967.01011349729 | 3934.02022699458 | 19760 | 0.014952615 |
| mt-Td | -12.76965066 | 0.94673122 | -0.248601653 | 1.216682169 | 0.311392888 | 0.72069767 | 0.5 | 6.79617E-08 | FALSE | 0relative convergence (4) | 4272.61178113989 | 4327.85416525696 | -2129.30589056994 | 4258.61178113989 | 19760 | 1.20991E-05 |
| Aamp | -9.827350044 | 0.28058367 | -0.248749096 | 0.161155511 | 0.06180422 | 0.055478936 | 0.5 | 0.003188785 | FALSE | 0relative convergence (4) | 20787.196219929 | 20842.438604046 | -10386.5981099645 | 20773.196219929 | 19760 | 0.047419102 |
| Tmbim1 | -9.98621983 | 0.548426087 | 0.035764592 | 0.236422503 | 0.173082536 | 0.000322709 | 0.5 | 5.64063E-05 | FALSE | 0relative convergence (4) | 20225.4363839563 | 20280.6787680734 | -10105.7181919781 | 20211.4363839563 | 19760 | 0.002721283 |
| Kcne4 | -11.12826302 | 0.450985746 | -0.017046637 | 0.505763875 | 1.370244227 | 0.133240762 | 0.5 | 7.6922E-07 | FALSE | 0relative convergence (4) | 9528.10863925731 | 9583.35102337438 | -4757.05431962865 | 9514.10863925731 | 19760 | 8.6812E-05 |
| Serpine2 | -7.894652123 | 0.583851865 | 0.13068674 | 0.36335242 | 0.558674956 | 0.000640187 | 0.5 | 0.000790743 | FALSE | 0relative convergence (4) | 62883.1258156427 | 62938.3681997598 | -31434.5629078214 | 62869.1258156427 | 19760 | 0.018612647 |
| Col4a4 | -11.53755449 | 0.71101987 | 0.539746361 | 0.494581091 | 3.859797865 | 0.148320999 | 0.5 | 5.51717E-14 | FALSE | 0relative convergence (4) | 7385.6345716657 | 7440.87695578278 | -3685.81728583285 | 7371.6345716657 | 19760 | 4.64913E-11 |
| Col4a3 | -11.71399329 | 0.650742102 | 0.953929601 | 0.884027431 | 5.799992692 | 8.48061E-05 | 0.5 | 3.57041E-08 | FALSE | 0relative convergence (4) | 7012.18547420574 | 7067.42785832282 | -3499.09273710287 | 6998.18547420574 | 19760 | 7.27902E-06 |
| Sphkap | -10.44831912 | 1.334141529 | 1.351042144 | 1.700375464 | 3.784167775 | 0.315975058 | 0.5 | 2.60779E-13 | FALSE | 0relative convergence (4) | 19936.8626966836 | 19992.1050808007 | -9961.43134834182 | 19922.8626966836 | 19760 | 1.93897E-10 |
| Itm2c | -9.419323292 | 0.635898949 | -0.121186844 | 0.36254852 | 0.113749849 | 0.000657539 | 0.5 | 0.000145337 | FALSE | 0relative convergence (4) | 28124.3673340004 | 28179.6097181175 | -14055.1836670002 | 28110.3673340004 | 19760 | 0.005566843 |
| Ube2f | -10.39294023 | 0.266213458 | -0.086345714 | 0.238729015 | 0.373381274 | 0.091618197 | 0.5 | 5.105E-07 | FALSE | 0relative convergence (4) | 15207.8674480995 | 15263.1098322166 | -7596.93372404974 | 15193.8674480995 | 19760 | 6.3262E-05 |
| Klhl30 | -11.97977888 | 0.631348433 | 0.263021312 | 0.418548686 | 0.626109472 | 0.107984274 | 0.5 | 0.003236754 | FALSE | 0relative convergence (4) | 5866.20770630403 | 5921.4500904211 | -2926.10385315201 | 5852.20770630403 | 19760 | 0.047683649 |
| Hes6 | -11.2190423 | 0.267745436 | 0.047866124 | 0.176586151 | 0.141063529 | 7.48004E-05 | 0.5 | 0.003213711 | FALSE | 0relative convergence (4) | 8528.60030766259 | 8583.84269177966 | -4257.30015383129 | 8514.60030766259 | 19760 | 0.047622618 |
| Nox4 | -8.334985486 | 0.374246667 | 0.464791294 | 0.574669435 | 1.15571114 | 0.352863543 | 0.5 | 9.31052E-19 | FALSE | 0relative convergence (4) | 52307.6707668636 | 52362.9131509807 | -26146.8353834318 | 52293.6707668636 | 19760 | 1.96142E-15 |
| Tmem126b | -11.39693901 | 0.274032925 | 0.168251001 | 0.178843044 | 2.05759E-08 | 2.30544E-09 | 0.5 | 0.001966753 | FALSE | 1false convergence (8) | 7437.7071123825 | 7492.94949649957 | -3711.85355619125 | 7423.7071123825 | 19760 | 0.034431792 |
| Lrrc32 | -11.57532761 | 0.45578375 | 0.653363031 | 0.539755886 | 1.128456994 | 0.309124167 | 0.5 | 0.000286066 | FALSE | 0relative convergence (4) | 7506.09880216197 | 7561.34118627904 | -3746.04940108098 | 7492.09880216197 | 19760 | 0.009224176 |
| P2ry2 | -12.74355509 | 0.707039566 | 0.167335021 | 0.146660485 | 0.952620091 | 0.181322531 | 0.5 | 1.64892E-06 | FALSE | 0relative convergence (4) | 2862.40265172079 | 2917.64503583786 | -1424.2013258604 | 2848.40265172079 | 19760 | 0.000161568 |
| Inppl1 | -10.72146091 | 0.233877655 | 0.412796905 | 0.479381337 | 0.075765623 | 0.264784425 | 0.5 | 0.000110954 | FALSE | 0relative convergence (4) | 12028.8131624087 | 12084.0555465257 | -6007.40658120433 | 12014.8131624087 | 19760 | 0.004495057 |
| Gja1 | -9.850975303 | 0.60963031 | -0.403729365 | 0.024051527 | 1.314591307 | 0.000185915 | 0.5 | 0.000135533 | FALSE | 0relative convergence (4) | 22169.7197015359 | 22224.9620856529 | -11077.8598507679 | 22155.7197015359 | 19760 | 0.005238966 |
| Psap | -7.727041382 | 0.34780208 | -0.128998467 | 0.321577084 | 0.233495433 | 0.000167092 | 0.5 | 0.00329322 | FALSE | 0relative convergence (4) | 62110.835576781 | 62166.0779608981 | -31048.4177883905 | 62096.835576781 | 19760 | 0.047991161 |
| Unc5b | -11.69925498 | 1.105802833 | -0.108557349 | 0.175396164 | 0.969334442 | 0.000481602 | 0.5 | 0.000103432 | FALSE | 0relative convergence (4) | 8494.01530234591 | 8549.25768646298 | -4240.00765117295 | 8480.01530234591 | 19760 | 0.004248961 |
| Sar1a | -9.637725242 | 0.285972141 | -0.028890003 | 0.290398815 | 1.97295E-05 | 0.000754135 | 0.5 | 0.000869543 | FALSE | 0relative convergence (4) | 23888.0838936914 | 23943.3262778084 | -11937.0419468457 | 23874.0838936914 | 19760 | 0.019803641 |
| Egr2 | -12.61363043 | 0.931281043 | -1.102605455 | 0.355388311 | 3.695923147 | 0.368998517 | 0.5 | 0.003355334 | FALSE | 0relative convergence (4) | 4584.17019747879 | 4639.41258159586 | -2285.08509873939 | 4570.17019747879 | 19760 | 0.048324936 |
| Mif | -9.553269486 | 0.6937176 | 0.094762945 | 0.545437053 | 0.161256404 | 0.281753129 | 0.5 | 0.001589928 | FALSE | 0relative convergence (4) | 29043.9199572785 | 29099.1623413956 | -14514.9599786393 | 29029.9199572785 | 19760 | 0.02995035 |
| Bcor | -9.521711607 | 0.294560232 | -0.057303599 | 0.072948858 | 0.5417114 | 0.13544511 | 0.5 | 0.000808958 | FALSE | 0relative convergence (4) | 26203.1407475708 | 26258.3831316878 | -13094.5703737854 | 26189.1407475708 | 19760 | 0.018900617 |
| Slc25a1 | -10.60759227 | 0.25372138 | 0.080937612 | -0.017575284 | 0.162108146 | 5.28453E-05 | 0.5 | 9.25541E-07 | FALSE | 0relative convergence (4) | 12574.0700846135 | 12629.3124687306 | -6280.03504230675 | 12560.0700846135 | 19760 | 9.83096E-05 |
| Eif4g1 | -9.003101285 | 0.259932585 | 0.12469195 | 0.222751384 | 0.116424007 | 0.122245046 | 0.5 | 2.10766E-05 | FALSE | 0relative convergence (4) | 33674.0245359951 | 33729.2669201122 | -16830.0122679976 | 33660.0245359951 | 19760 | 0.001299175 |
| P3h2 | -9.994687124 | 0.753497433 | 0.422059002 | 0.830591127 | 1.918840631 | 0.000145577 | 0.5 | 0.000231193 | FALSE | 0relative convergence (4) | 22336.3029500829 | 22391.5453341999 | -11161.1514750414 | 22322.3029500829 | 19760 | 0.007855603 |
| Plaat1 | -12.38531149 | 0.597775497 | 0.188156741 | 0.581147323 | 1.235588044 | 0.707796141 | 0.5 | 0.001861785 | FALSE | 0relative convergence (4) | 4801.61127091958 | 4856.85365503666 | -2393.80563545979 | 4787.61127091958 | 19760 | 0.033098403 |
| Bdh1 | -11.81507788 | 0.613232709 | 0.081785842 | 0.772435201 | 3.166921485 | 0.379643143 | 0.5 | 0.000195261 | FALSE | 0relative convergence (4) | 6736.50446295554 | 6791.74684707261 | -3361.25223147777 | 6722.50446295554 | 19760 | 0.007041463 |
| Pcyt1a | -9.975239324 | 0.144820192 | -0.149645166 | -0.047552771 | 0.328155197 | 0.100936802 | 0.5 | 0.001965261 | FALSE | 0relative convergence (4) | NA | NA | NA | NA | 19760 | 0.034431792 |
| Tnk2 | -10.11890391 | 0.215237249 | -0.110325902 | 0.34153674 | 0.108455724 | 0.064514057 | 0.5 | 0.000583843 | FALSE | 0relative convergence (4) | 17629.438975302 | 17684.681359419 | -8807.71948765099 | 17615.438975302 | 19760 | 0.014952615 |
| Pdia5 | -12.27063529 | 0.454291403 | 0.249436179 | 0.325368441 | 1.769293836 | 0.211444346 | 0.5 | 0.000424142 | FALSE | 0relative convergence (4) | 3863.8329320515 | 3919.07531616857 | -1924.91646602575 | 3849.8329320515 | 19760 | 0.012212186 |
| Cep164 | -10.78462596 | 0.188332204 | 0.464703261 | 0.066404032 | 0.357302108 | 4.70758E-05 | 0.5 | 0.00132572 | FALSE | 0relative convergence (4) | 11697.2969497427 | 11752.5393338598 | -5841.64847487135 | 11683.2969497427 | 19760 | 0.02664085 |
| Ap5s1 | -12.07415217 | 0.382334775 | 0.185326787 | 0.140168283 | 3.82111E-07 | 5.18133E-09 | 0.5 | 0.000159207 | FALSE | 1 singular convergence (7) | NA | NA | NA | NA | 19760 | 0.006025079 |
| Cds2 | -9.676886191 | 0.194233135 | 0.142340623 | 0.21855625 | 0.114774665 | 0.066877641 | 0.5 | 2.93153E-07 | FALSE | 0relative convergence (4) | NA | NA | NA | NA | 19760 | 4.02767E-05 |
| Bmp2 | -11.70308305 | 0.98701244 | 0.115525184 | 0.487088656 | 2.14679581 | 8.91038E-05 | 0.5 | 4.05524E-07 | FALSE | 0relative convergence (4) | 7179.77267283824 | 7235.01505695531 | -3582.88633641912 | 7165.77267283824 | 19760 | 5.28435E-05 |
| Mkks | -11.89092871 | 0.367630752 | 0.243569944 | 0.401338766 | 3.18757E-08 | 0.190184585 | 0.5 | 0.000697897 | FALSE | 0relative convergence (4) | 5499.36264526972 | 5554.6050293868 | -2742.68132263486 | 5485.36264526972 | 19760 | 0.016895984 |
| Lig4 | -12.6601244 | 0.494889865 | 0.155138217 | 0.639815381 | 1.70053487 | 0.094247326 | 0.5 | 0.000585062 | FALSE | 0relative convergence (4) | 3144.91309279725 | 3200.15547691433 | -1565.45654639863 | 3130.91309279725 | 19760 | 0.014952615 |
| Plat | -10.97304949 | 0.836918007 | 0.441519055 | 0.53406012 | 1.181485969 | 0.000601443 | 0.5 | 0.002642786 | FALSE | 0relative convergence (4) | 12981.9003406396 | 13037.1427247567 | -6483.95017031981 | 12967.9003406396 | 19760 | 0.041913198 |
| Fgfr1 | -9.015985416 | 0.579897428 | 0.175222259 | 0.157012792 | 0.387012799 | 0.123843949 | 0.5 | 3.36817E-06 | FALSE | 0relative convergence (4) | 35279.2516765733 | 35334.4940606904 | -17632.6258382867 | 35265.2516765733 | 19760 | 0.000299815 |
| Emp1 | -11.18291779 | 1.085189333 | 0.13979004 | 0.413774956 | 6.344233945 | 0.000170302 | 0.5 | 0.002612503 | FALSE | 0relative convergence (4) | 10228.5383146339 | 10283.780698751 | -5107.26915731695 | 10214.5383146339 | 19760 | 0.041641908 |
| Ppp1r12c | -9.173473741 | 0.338133751 | 0.093075463 | 0.371656039 | 0.080597615 | 0.170944658 | 0.5 | 0.000325991 | FALSE | 0relative convergence (4) | 30762.2249258788 | 30817.4673099958 | -15374.1124629394 | 30748.2249258788 | 19760 | 0.010109006 |
| Cops7a | -11.13885476 | 0.43792659 | -0.407104728 | 0.319201818 | 0.049289432 | 0.003166756 | 0.5 | 0.002410895 | FALSE | 0relative convergence (4) | 9401.01692880298 | 9456.25931292005 | -4693.50846440149 | 9387.01692880298 | 19760 | 0.039707351 |
| Gapdh | -8.584296225 | 0.846273477 | 0.658307196 | 0.825029599 | 0.119181021 | 0.449555354 | 0.5 | 0.000542578 | FALSE | 0relative convergence (4) | 45758.9071731651 | 45814.1495572822 | -22872.4535865825 | 45744.9071731651 | 19760 | 0.014287889 |
| Ltbr | -11.02025521 | 0.612173764 | -0.091599296 | 0.360565747 | 0.026910686 | 0.000161842 | 0.5 | 0.00302728 | FALSE | 0relative convergence (4) | NA | NA | NA | NA | 19760 | 0.046198077 |
| Pgk1 | -9.846523273 | 0.472948624 | -0.119780131 | 0.412553856 | 0.121804709 | 0.071306641 | 0.5 | 6.17727E-07 | FALSE | 0relative convergence (4) | 22209.3076278347 | 22264.5500119518 | -11097.6538139174 | 22195.3076278347 | 19760 | 7.3661E-05 |
| Tceal6 | -12.68660273 | 1.141222845 | 0.717110468 | 1.409796734 | 4.801266536 | 0.517643002 | 0.5 | 0.000103535 | FALSE | 0relative convergence (4) | 4861.72514573076 | 4916.96752984783 | -2423.86257286538 | 4847.72514573076 | 19760 | 0.004248961 |
| Ak4 | -12.80224942 | 0.843136807 | 0.350598061 | 0.288046164 | 3.145631663 | 0.346114946 | 0.5 | 9.30292E-05 | FALSE | 0relative convergence (4) | 3420.70997085479 | 3475.95235497186 | -1703.35498542739 | 3406.70997085479 | 19760 | 0.003945936 |
| Leprot | -9.74265893 | 0.20777322 | 0.18018298 | 0.312615508 | 0.092676309 | 0.120287357 | 0.5 | 5.8674E-06 | FALSE | 0relative convergence (4) | 22119.5904618662 | 22174.8328459833 | -11052.7952309331 | 22105.5904618662 | 19760 | 0.000460646 |
| Podn | -13.03632828 | 0.852388558 | 0.473221709 | 1.015691505 | 6.154055705 | 0.462162861 | 0.5 | 3.54689E-06 | FALSE | 0relative convergence (4) | 2496.25909119731 | 2551.50147531438 | -1241.12954559865 | 2482.25909119731 | 19760 | 0.000309191 |
| Mast2 | -9.600174291 | 0.285202329 | -0.004244601 | 0.067993839 | 0.631233538 | 0.127165889 | 0.5 | 0.000961383 | FALSE | 0relative convergence (4) | 25470.6384907323 | 25525.8808748494 | -12728.3192453661 | 25456.6384907323 | 19760 | 0.021281762 |
| Akr1a1 | -8.583089086 | 0.256540737 | -0.174421593 | 0.28054891 | 0.061739421 | 9.96808E-05 | 0.5 | 0.002265296 | FALSE | 0relative convergence (4) | 40698.5928449607 | 40753.8352290777 | -20342.2964224803 | 40684.5928449607 | 19760 | 0.037975246 |
| Spry4 | -12.12126305 | 0.668119105 | 0.232883904 | 0.570853501 | 0.947712556 | 0.244727011 | 0.5 | 0.002817343 | FALSE | 0relative convergence (4) | 5318.84444525763 | 5374.0868293747 | -2652.42222262881 | 5304.84444525763 | 19760 | 0.043856178 |
| Fgf1 | -10.20596584 | 0.580195549 | 0.463653509 | 0.521811344 | 0.749068135 | 0.13134157 | 0.5 | 7.20734E-08 | FALSE | 0relative convergence (4) | 18442.7990838805 | 18498.0414679975 | -9214.39954194023 | 18428.7990838805 | 19760 | 1.26529E-05 |
| C030004G16Rik | -12.0725544 | 0.874672931 | 0.87119256 | 0.836669199 | 1.662887157 | 0.329165699 | 0.5 | 3.97143E-16 | FALSE | 0relative convergence (4) | 5467.69553365438 | 5522.93791777145 | -2726.84776682719 | 5453.69553365438 | 19760 | 4.37808E-13 |
| Eif1a | -10.03298265 | 0.417714927 | -0.17271686 | -0.023113961 | 0.435744393 | 0.175789914 | 0.5 | 0.000316675 | FALSE | 0relative convergence (4) | 19706.4816811291 | 19761.7240652462 | -9846.24084056455 | 19692.4816811291 | 19760 | 0.009885647 |
| Ppic | -10.14281988 | 0.530885109 | -0.060083662 | 0.300264763 | 0.213580632 | 0.315928551 | 0.5 | 0.002071952 | FALSE | 0relative convergence (4) | 18981.2618385416 | 19036.5042226587 | -9483.63091927079 | 18967.2618385416 | 19760 | 0.035631944 |
| Fosl2 | -9.919777809 | 0.524620945 | -0.234015232 | 0.364733605 | 0.639272726 | 0.190274209 | 0.5 | 0.000286027 | FALSE | 0relative convergence (4) | 22569.4627996879 | 22624.705183805 | -11277.731399844 | 22555.4627996879 | 19760 | 0.009224176 |
| Mxd4 | -9.302974731 | 0.146502399 | 0.071375193 | 0.178137406 | 0.101446615 | 0.104863047 | 0.5 | 0.000476038 | FALSE | 0relative convergence (4) | 27970.5212488119 | 28025.763632929 | -13978.260624406 | 27956.5212488119 | 19760 | 0.013181948 |
| Tnip2 | -10.77048378 | 0.216879312 | -0.096302798 | 0.17115141 | 0.464598522 | 0.130312946 | 0.5 | 0.000844898 | FALSE | 0relative convergence (4) | 11589.4959216612 | 11644.7383057782 | -5787.74796083058 | 11575.4959216612 | 19760 | 0.019417291 |
| Wdr1 | -8.998859576 | 0.228600248 | -0.355946969 | 0.026317405 | 0.0693623 | 0.111416795 | 0.5 | 1.04548E-07 | FALSE | 0relative convergence (4) | 33244.2848973088 | 33299.5272814259 | -16615.1424486544 | 33230.2848973088 | 19760 | 1.67276E-05 |
| C1qtnf7 | -11.0158961 | 0.612331791 | 0.331663155 | 0.339634661 | 1.198661417 | 0.000145555 | 0.5 | 3.66165E-05 | FALSE | 0relative convergence (4) | 10671.6675287909 | 10726.909912908 | -5328.83376439544 | 10657.6675287909 | 19760 | 0.001986408 |
| Cttnbp2nl | -9.992447375 | 0.263702153 | 0.054627106 | 0.222714144 | 0.577731653 | 0.21447718 | 0.5 | 0.001269521 | FALSE | 0relative convergence (4) | 19291.6203330186 | 19346.8627171356 | -9638.81016650928 | 19277.6203330186 | 19760 | 0.025893222 |
| Ndst3 | -11.1630172 | 0.80889565 | 0.702652321 | 0.669645378 | 4.72490882 | 0.367016934 | 0.5 | 1.02131E-06 | FALSE | 0relative convergence (4) | 9475.43147682538 | 9530.67386094245 | -4730.71573841269 | 9461.43147682538 | 19760 | 0.000106689 |
| Col18a1 | -8.011148357 | 0.801703824 | 0.223972016 | 0.224255385 | 0.468251772 | 0.00050346 | 0.5 | 5.16681E-06 | FALSE | 0relative convergence (4) | 60296.6195099342 | 60351.8618940513 | -30141.3097549671 | 60282.6195099342 | 19760 | 0.000421345 |
| Pfkl | -10.4796795 | 0.76086722 | -0.064353734 | 0.34086983 | 0.172557097 | 0.000211445 | 0.5 | 2.12098E-05 | FALSE | 0relative convergence (4) | 16320.9905571181 | 16376.2329412352 | -8153.49527855904 | 16306.9905571181 | 19760 | 0.001299175 |
| Ilvbl | -11.25596483 | 0.222912091 | 0.159775641 | 0.36148965 | 0.000883213 | 0.063870243 | 0.5 | 0.003179405 | FALSE | 0relative convergence (4) | NA | NA | NA | NA | 19760 | 0.047396742 |
| Tmem259 | -10.29173372 | 0.38506828 | 0.191593185 | 0.563290535 | 0.032397054 | 0.00016012 | 0.5 | 0.000499488 | FALSE | 0relative convergence (4) | 17292.6174467132 | 17347.8598308302 | -8639.30872335658 | 17278.6174467132 | 19760 | 0.013548333 |
| Sbno2 | -10.19595938 | 0.61110597 | -0.009959859 | 0.597041991 | 1.793689973 | 0.342479927 | 0.5 | 0.000948959 | FALSE | 0relative convergence (4) | 20791.9339768075 | 20847.1763609246 | -10388.9669884038 | 20777.9339768075 | 19760 | 0.021043585 |
| Adamtsl5 | -9.857206763 | 0.407690395 | 0.323043092 | 0.429836512 | 0.301440766 | 0.171126904 | 0.5 | 7.538E-05 | FALSE | 0relative convergence (4) | 22178.8564503256 | 22234.0988344427 | -11082.4282251628 | 22164.8564503256 | 19760 | 0.003446511 |
| Csnk1g2 | -10.09587127 | 0.289334034 | 0.170275474 | 0.481529928 | 3.78577E-12 | 0.218568122 | 0.5 | 4.62159E-05 | FALSE | 1false convergence (8) | 18662.9869945037 | 18718.2293786208 | -9324.49349725184 | 18648.9869945037 | 19760 | 0.002352952 |
| Sppl2b | -11.84265728 | 0.453521387 | 0.259890007 | 0.397311864 | 0.240161158 | 0.240577203 | 0.5 | 4.82927E-06 | FALSE | 0relative convergence (4) | NA | NA | NA | NA | 19760 | 0.000398968 |
| Pld2 | -10.93567252 | 0.27262816 | -0.084723851 | 0.277772892 | 0.840861156 | 0.204234478 | 0.5 | 0.000196494 | FALSE | 0relative convergence (4) | 10104.1581041164 | 10159.4004882334 | -5045.07905205818 | 10090.1581041164 | 19760 | 0.007041463 |
| P2rx1 | -11.98029672 | 0.69678864 | -0.193261353 | 0.577425827 | 6.211464235 | 9.23514E-05 | 0.5 | 3.42711E-07 | FALSE | 0relative convergence (4) | 5309.26085743563 | 5364.5032415527 | -2647.63042871781 | 5295.26085743563 | 19760 | 4.61113E-05 |
| Serpinf1 | -11.73756855 | 0.622867899 | -0.176157803 | 0.317779277 | 11.67386138 | 0.336809159 | 0.5 | 3.24446E-05 | FALSE | 0relative convergence (4) | 5455.3081496699 | 5510.55053378697 | -2720.65407483495 | 5441.3081496699 | 19760 | 0.001783045 |
| Coro6 | -12.42058092 | 0.438459683 | 0.480874062 | 0.520423206 | 0.232314831 | 0.278555625 | 0.5 | 0.00093242 | FALSE | 0relative convergence (4) | 3690.62850588687 | 3745.87089000395 | -1838.31425294344 | 3676.62850588687 | 19760 | 0.020786217 |
| Git1 | -11.25011171 | 0.449759676 | 0.102963257 | 0.189486461 | 0.239688609 | 0.000110416 | 0.5 | 1.02537E-05 | FALSE | 0relative convergence (4) | 8741.78237052585 | 8797.02475464293 | -4363.89118526293 | 8727.78237052585 | 19760 | 0.000700579 |
| Tmem97 | -12.09535495 | 0.380122215 | -0.112249327 | 0.271801539 | 0.223518447 | 0.099212012 | 0.5 | 0.000644244 | FALSE | 0relative convergence (4) | 4500.37025979063 | 4555.6126439077 | -2243.18512989531 | 4486.37025979063 | 19760 | 0.016008339 |
| Gm26532 | -10.33160437 | 1.313296336 | -0.459963401 | 1.141474269 | 2.591867692 | 0.251946962 | 0.5 | 8.29479E-07 | FALSE | 0relative convergence (4) | 20739.3128575308 | 20794.5552416479 | -10362.6564287654 | 20725.3128575308 | 19760 | 9.0531E-05 |
| Ier2 | -9.502888431 | 1.100124091 | -0.028273297 | 0.953900466 | 0.664780005 | 0.001157982 | 0.5 | 0.003016591 | FALSE | 0relative convergence (4) | 35141.5924221073 | 35196.8348062244 | -17563.7962110537 | 35127.5924221073 | 19760 | 0.046161876 |
| Dhps | -10.84296172 | 0.199318455 | 0.349059729 | 0.273100174 | 0.25041397 | 0.128622364 | 0.5 | 0.00239301 | FALSE | 0relative convergence (4) | NA | NA | NA | NA | 19760 | 0.039487782 |
| Kifc3 | -11.74234537 | 0.727754731 | 0.142877178 | 0.246296448 | 0.863900426 | 8.36281E-05 | 0.5 | 1.37657E-05 | FALSE | 0relative convergence (4) | 6241.04174126102 | 6296.28412537809 | -3113.52087063051 | 6227.04174126102 | 19760 | 0.000901546 |
| Cdh11 | -10.71403983 | 0.515283442 | 0.429689999 | 0.437839407 | 2.459314282 | 7.80946E-05 | 0.5 | 6.67405E-05 | FALSE | 0relative convergence (4) | 12952.1133696609 | 13007.355753778 | -6469.05668483047 | 12938.1133696609 | 19760 | 0.003112913 |

| | | | | | | | | | | | | | | | | | |
|---|---|---|---|---|---|---|---|---|---|---|---|---|---|---|---|---|---|
| B3gnt9 | -11.92927916 | 0.471504805 | -0.056338586 | 0.367274962 | 0.802399295 | 0.000414919 | 0.5 | 5.51151E-07 | FALSE | 0 | relative convergence (4) | NA | NA | NA | NA | 19760 | 6.6986E-05 |
| Fbxl8 | -12.69440066 | 0.499552828 | 0.27948142 | 0.826915921 | 0.598899855 | 0.335084038 | 0.5 | 0.000655238 | FALSE | 0 | relative convergence (4) | 3306.97110125407 | 3362.21348537114 | -1646.48555062704 | 3292.97110125407 | 19760 | 0.016144653 |
| Map3k8 | -10.84138362 | 0.449036524 | -0.191192056 | -0.038019672 | 1.181994489 | 0.095506018 | 0.5 | 0.001297761 | FALSE | 0 | relative convergence (4) | 11717.7202972272 | 11772.9626813443 | -5851.86014861359 | 11703.7202972272 | 19760 | 0.026245924 |
| Cables1 | -10.46130959 | 0.473885773 | 0.174239452 | 0.087688947 | 1.072793999 | 9.9739E-05 | 0.5 | 0.000457947 | FALSE | 0 | relative convergence (4) | 14557.0449378146 | 14612.2873219317 | -7271.5224689073 | 14543.0449378146 | 19760 | 0.012847334 |
| Pdk1 | -11.35810896 | 0.448626457 | -0.12632428 | 0.020153657 | 1.133430236 | 9.00126E-05 | 0.5 | 5.08255E-05 | FALSE | 0 | relative convergence (4) | 7740.08940968155 | 7795.33179379862 | -3863.04470484077 | 7726.08940968155 | 19760 | 0.00250951 |
| Itprid2 | -9.950387349 | 0.344750631 | 0.044086619 | 0.167794137 | 0.313269947 | 0.000108165 | 0.5 | 0.002630686 | FALSE | 0 | relative convergence (4) | 20286.8950515377 | 20342.1374356547 | -10136.4475257688 | 20272.8950515377 | 19760 | 0.041826258 |
| Serping1 | -9.196492187 | 0.396903598 | 0.025651527 | 0.226766394 | 0.472837748 | 0.000268662 | 0.5 | 0.003424934 | FALSE | 0 | relative convergence (4) | 31742.5884009837 | 31797.8307851008 | -15864.2942004918 | 31728.5884009837 | 19760 | 0.048805208 |
| Ube2l6 | -12.246626 | 0.607876871 | -0.092562955 | 0.323629041 | 0.496958953 | 6.72248E-05 | 0.5 | 0.000224177 | FALSE | 0 | relative convergence (4) | 4199.38907941705 | 4254.63146353413 | -2092.69453970853 | 4185.38907941705 | 19760 | 0.007697248 |
| Cry2 | -10.43886559 | 0.746290132 | 0.090310266 | 0.277645923 | 0.623027101 | 0.13109108 | 0.5 | 4.16998E-05 | FALSE | 0 | relative convergence (4) | 16423.679738743 | 16478.9221228601 | -8204.8398693715 | 16409.679738743 | 19760 | 0.002169077 |
| Fjx1 | -11.76435725 | 0.737087787 | -0.334214371 | -0.007391771 | 1.590684766 | 0.000184612 | 0.5 | 0.0010651 | FALSE | 0 | relative convergence (4) | 6377.99363150529 | 6433.23601562237 | -3181.99681575265 | 6363.99363150529 | 19760 | 0.022857158 |
| Gne | -10.4778506 | 0.335434691 | 0.314155983 | 0.339679562 | 0.61056253 | 0.188166465 | 0.5 | 0.000420263 | FALSE | 0 | relative convergence (4) | 15113.3794479108 | 15168.6218320279 | -7549.68972395542 | 15099.3794479108 | 19760 | 0.012155882 |
| Col15a1 | -8.570703368 | 0.693192361 | -0.354869582 | -0.324424944 | 0.513173924 | 0.000696798 | 0.5 | 0.000605758 | FALSE | 0 | relative convergence (4) | 43898.8291401216 | 43954.0715242387 | -21942.4145700608 | 43884.8291401216 | 19760 | 0.015344252 |
| Rgs3 | -10.70234919 | 0.613914117 | -0.060931335 | 0.226815021 | 0.32840887 | 0.000152196 | 0.5 | 4.14421E-07 | FALSE | 0 | relative convergence (4) | 12775.318480641 | 12830.560864758 | -6380.65924032048 | 12761.318480641 | 19760 | 5.34519E-05 |
| Tnc | -10.90513608 | 0.852860753 | 0.581936666 | 1.223319872 | 3.911147685 | 0.420522393 | 0.5 | 1.71457E-08 | FALSE | 0 | relative convergence (4) | 12870.1116263847 | 12925.3540105017 | -6428.05581319233 | 12856.1116263847 | 19760 | 4.33443E-06 |
| Dmac1 | -10.17700125 | 0.191950578 | 0.118376735 | 0.251068566 | 0.064754932 | 4.52351E-05 | 0.5 | 0.000481832 | FALSE | 0 | relative convergence (4) | 17138.3801454212 | 17193.6225295383 | -8562.19007271061 | 17124.3801454212 | 19760 | 0.013219656 |
| Srxn1 | -11.48391538 | 0.275995966 | 0.097228084 | 0.21459238 | 0.275324372 | 0.204274698 | 0.5 | 0.001869937 | FALSE | 0 | relative convergence (4) | 7149.06427502732 | 7204.30665914439 | -3567.53213751366 | 7135.06427502732 | 19760 | 0.033196628 |
| Rem1 | -12.40308771 | 0.804482576 | 0.766224806 | 0.521878153 | 0.719950016 | 6.60348E-05 | 0.5 | 9.2596E-08 | FALSE | 0 | relative convergence (4) | 4181.30975149405 | 4236.55213561113 | -2083.65487574703 | 4167.30975149405 | 19760 | 1.50701E-05 |
| H13 | -9.589850747 | 0.274485641 | -0.089014692 | 0.17057418 | 0.236300458 | 0.000175199 | 0.5 | 2.69487E-08 | FALSE | 0 | relative convergence (4) | 24224.1488593904 | 24279.3912435075 | -12105.0744296952 | 24210.1488593904 | 19760 | 6.1933E-06 |
| Cox4i2 | -10.90058998 | 0.953970268 | 0.079712064 | 0.440768751 | 1.30973196 | 0.000255518 | 0.5 | 0.000699486 | FALSE | 0 | relative convergence (4) | 13280.6742963858 | 13335.9166805029 | -6633.33714819291 | 13266.6742963858 | 19760 | 0.016895984 |
| Chmp4b | -9.150686975 | 0.132405431 | -0.190759882 | -0.165607262 | 0.046576819 | 0.153151499 | 0.5 | 1.08489E-05 | FALSE | 0 | relative convergence (4) | NA | NA | NA | NA | 19760 | 0.000729416 |
| BC029722 | -11.19587037 | 0.450771203 | 0.271518418 | 0.461691052 | 0.040500365 | 0.129470829 | 0.5 | 5.76507E-10 | FALSE | 0 | relative convergence (4) | 9274.04445865958 | 9329.28684277666 | -4630.02222932979 | 9260.04445865958 | 19760 | 2.2772E-07 |
| Eif6 | -10.09016633 | 0.277693374 | 0.116310297 | 0.270462615 | 3.1633E-07 | 0.089436074 | 0.5 | 0.000647904 | FALSE | 0 | relative convergence (4) | 18408.1275565121 | 18463.3699406292 | -9197.06377825605 | 18394.1275565121 | 19760 | 0.016008528 |
| Rpn2 | -8.838845949 | 0.242058668 | 0.028471178 | 0.122716544 | 0.053018602 | 7.96876E-05 | 0.5 | 0.00027718 | FALSE | 0 | relative convergence (4) | 35926.0984550792 | 35981.3408391963 | -17956.0492275396 | 35912.0984550792 | 19760 | 0.00900658 |
| Ccn5 | -10.99980293 | 0.995759796 | 0.056305632 | 0.998232488 | 2.823443116 | 0.00047187 | 0.5 | 5.61855E-05 | FALSE | 0 | relative convergence (4) | 13235.3213484347 | 13290.5637325517 | -6610.66067421733 | 13221.3213484347 | 19760 | 0.002721016 |
| Gm10824 | -12.63347972 | 0.517004629 | 0.007246651 | 0.233712557 | 3.080135163 | 0.126548414 | 0.5 | 0.000886036 | FALSE | 0 | relative convergence (4) | 3015.00840033835 | 3070.25078445542 | -1500.50420016917 | 3001.00840033835 | 19760 | 0.020106815 |
| Ccn2 | -8.171401053 | 0.616421815 | 0.563898863 | 1.164901012 | 1.120334691 | 0.449498412 | 0.5 | 0.000388224 | FALSE | 0 | relative convergence (4) | 59259.468966705 | 59314.7113508221 | -29622.7344833525 | 59245.468966705 | 19760 | 0.011465309 |
| Enpp1 | -10.43575696 | 0.207922751 | 0.014793 | 0.180205546 | 0.769818128 | 7.92247E-11 | 0.5 | 5.8064E-05 | FALSE | 0 | relative convergence (4) | 14466.78444285 | 14522.0268269671 | -7226.392221425 | 14452.78444285 | 19760 | 0.002769544 |
| Spryd3 | -10.86799548 | 0.302146559 | 0.105060232 | 0.089173296 | 0.050030916 | 0.129871436 | 0.5 | 0.000326303 | FALSE | 0 | relative convergence (4) | 10828.7770511685 | 10884.0194352856 | -5407.38852558427 | 10814.7770511685 | 19760 | 0.010109006 |
| Snai2 | -11.76232172 | 0.446822476 | -0.064762221 | 0.105598072 | 1.227332506 | 0.000165998 | 0.5 | 0.002898779 | FALSE | 0 | relative convergence (4) | 5840.10626657137 | 5895.34865068845 | -2913.05313328569 | 5826.10626657137 | 19760 | 0.044902662 |
| Atp1b3 | -8.93190459 | 0.231187463 | 0.099438144 | 0.044945004 | 0.12381378 | 8.96342E-05 | 0.5 | 9.74961E-05 | FALSE | 0 | relative convergence (4) | 35054.1821078824 | 35109.4244919995 | -17520.0910539412 | 35040.1821078824 | 19760 | 0.004062794 |
| Dusp7 | -11.12577784 | 0.501800436 | 0.074551443 | 0.261254928 | 0.431053606 | 7.23045E-05 | 0.5 | 0.001098446 | FALSE | 0 | relative convergence (4) | 9239.18579071658 | 9294.42817483365 | -4612.59289535829 | 9225.18579071658 | 19760 | 0.023295895 |
| Rrp9 | -11.73700006 | 0.291199656 | 0.044090864 | 0.378358441 | 0.207903269 | 0.301031074 | 0.5 | 0.002781041 | FALSE | 0 | relative convergence (4) | 6146.56427538086 | 6201.80665949793 | -3066.28213769043 | 6132.56427538086 | 19760 | 0.043505401 |
| Tex264 | -10.56134637 | 0.506411278 | 0.262161557 | 0.337692296 | 0.004895191 | 2.71425E-17 | 0.5 | 0.000743621 | FALSE | 0 | relative convergence (4) | NA | NA | NA | NA | 19760 | 0.017667988 |
| Rassf1 | -10.23237736 | 0.721587285 | -0.116264238 | 0.573660367 | 0.63161882 | 0.000215568 | 0.5 | 0.00052444 | FALSE | 0 | relative convergence (4) | 19808.4649145768 | 19863.7072986938 | -9897.23245728839 | 19794.4649145768 | 19760 | 0.013955614 |
| Gmppb | -11.98155263 | 0.38881769 | 0.282211675 | 0.31571157 | 0.577297189 | 0.179121211 | 0.5 | 0.000324174 | FALSE | 0 | relative convergence (4) | 4940.71550633301 | 4995.95789045008 | -2463.3577531665 | 4926.71550633301 | 19760 | 0.010092499 |
| Ptprz1 | -9.04733187 | 0.581381661 | 0.260437062 | 0.487389871 | 1.62314668 | 0.000294268 | 0.5 | 0.00109096 | FALSE | 0 | relative convergence (4) | 36371.2237678352 | 36426.4661519523 | -18178.6118839176 | 36357.2237678352 | 19760 | 0.023215041 |
| Arf5 | -9.755560024 | 0.417159792 | 0.162778933 | -0.079817691 | 0.021643923 | 1.28711E-15 | 0.5 | 0.00097628 | FALSE | 0 | relative convergence (4) | 22892.3853038024 | 22947.6276879195 | -11439.1926519012 | 22878.3853038024 | 19760 | 0.021498575 |
| Impdh1 | -11.21475341 | 0.414392693 | -0.046209154 | 0.282436354 | 0.117463962 | 0.20695027 | 0.5 | 0.000262565 | FALSE | 0 | relative convergence (4) | 8843.02955205211 | 8898.27193616918 | -4414.51477602605 | 8829.02955205211 | 19760 | 0.008667129 |
| F630206G17Rik | -13.12946709 | 0.795903383 | 0.312599484 | -0.441452652 | 0.687886677 | 0.225832759 | 0.5 | 0.000231072 | FALSE | 0 | relative convergence (4) | 2211.87838397881 | 2267.12076809589 | -1098.93919198941 | 2197.87838397881 | 19760 | 0.007855603 |
| Kif3a | -10.64102384 | 0.278145054 | 0.306358006 | 0.20638807 | 0.566851264 | 0.126785585 | 0.5 | 4.82183E-06 | FALSE | 0 | relative convergence (4) | NA | NA | NA | NA | 19760 | 0.000398968 |
| Nr1d1 | -9.637943951 | 0.869505587 | -0.392132986 | -0.773609945 | 0.532184579 | 0.000277213 | 0.5 | 0.000720857 | FALSE | 0 | relative convergence (4) | 24676.9836469784 | 24732.2260310955 | -12331.4918234892 | 24662.9836469784 | 19760 | 0.017224257 |
| Jup | -10.19194387 | 0.455383324 | 0.051707661 | -0.036865863 | 0.312748095 | 0.000172226 | 0.5 | 0.00070027 | FALSE | 0 | relative convergence (4) | 17412.4157043922 | 17467.6580885092 | -8699.20785219609 | 17398.4157043922 | 19760 | 0.016895984 |
| Vat1 | -10.27921156 | 0.607265333 | 0.067554823 | 0.360890209 | 0.302286127 | 0.000111325 | 0.5 | 0.000109826 | FALSE | 0 | relative convergence (4) | 17674.3847761086 | 17729.6271602257 | -8830.19238805429 | 17660.3847761086 | 19760 | 0.004478069 |
| 4930417O22Rik | -12.69392934 | 0.746192899 | 0.108380115 | -0.05266048 | 1.997952207 | 6.58678E-05 | 0.5 | 4.79413E-09 | FALSE | 0 | relative convergence (4) | 2945.16122509148 | 3000.40360920856 | -1465.58061254574 | 2931.16122509148 | 19760 | 1.30257E-06 |
| Sost | -7.815635714 | 0.731539965 | 0.026078735 | 0.020925736 | 1.583697935 | 0.186715425 | 0.5 | 0.000235165 | FALSE | 0 | relative convergence (4) | 65028.1571468487 | 65083.3995309658 | -32507.0785734243 | 65014.1571468487 | 19760 | 0.007947823 |
| Ckap4 | -9.306823418 | 0.294016749 | -0.350869347 | -0.14252586 | 0.222807465 | 0.166466128 | 0.5 | 7.71723E-05 | FALSE | 0 | relative convergence (4) | 28529.2600687376 | 28584.5024528546 | -14257.6300343688 | 28515.2600687376 | 19760 | 0.003459068 |
| Thbs1 | -11.0976396 | 0.74116093 | -0.189573424 | 0.023275663 | 16.11815913 | 0.415281889 | 0.5 | 0.002997199 | FALSE | 0 | relative convergence (4) | 9126.54326656556 | 9181.78565068263 | -4556.27163328278 | 9112.54326656556 | 19760 | 0.045976448 |
| Ehd4 | -8.702031472 | 0.31695697 | -0.097932518 | 0.009302267 | 0.548666221 | 0.108450379 | 0.5 | 0.000483187 | FALSE | 0 | relative convergence (4) | 39610.4635316831 | 39665.7059158002 | -19798.2317658416 | 39596.4635316831 | 19760 | 0.013219656 |
| Gap43 | -13.45787951 | 1.483393279 | 3.198143936 | 1.980682509 | 0.926596032 | 0.563551286 | 0.5 | 1.15731E-06 | FALSE | 0 | relative convergence (4) | 4270.96496423403 | 4326.20734835111 | -2128.48248211702 | 4256.96496423403 | 19760 | 0.000117971 |
| Gm29686 | -13.11979563 | 1.351029603 | 0.065458255 | 0.736776764 | 5.160652612 | 0.000102091 | 0.5 | 1.12611E-06 | FALSE | 0 | relative convergence (4) | 2911.47007317164 | 2966.71245728871 | -1448.73503658582 | 2897.47007317164 | 19760 | 0.000115723 |
| Col8a1 | -12.78692257 | 0.856011558 | 0.616507725 | 0.534858151 | 21.59082978 | 0.420729344 | 0.5 | 3.45917E-06 | FALSE | 0 | relative convergence (4) | NA | NA | NA | NA | 19760 | 0.000305761 |
| Optn | -10.88513103 | 0.296236995 | -0.021655362 | 0.080841795 | 0.774051427 | 7.30184E-05 | 0.5 | 0.002120294 | FALSE | 0 | relative convergence (4) | 10879.1158035185 | 10934.3581876356 | -5432.55790175927 | 10865.1158035185 | 19760 | 0.036265928 |
| A930004D18Rik | -12.85240728 | 0.531328553 | 0.114133548 | 0.409659736 | 2.210580792 | 2.03934E-22 | 0.5 | 0.000290515 | FALSE | 1 | singular convergence (7) | 2591.17307710159 | 2646.41546121866 | -1288.5865385508 | 2577.17307710159 | 19760 | 0.009343798 |
| Rnf152 | -10.7266415 | 0.435265565 | 0.255534075 | 0.579289924 | 2.397619064 | 0.262850792 | 0.5 | 7.69728E-10 | FALSE | 0 | relative convergence (4) | NA | NA | NA | NA | 19760 | 2.94829E-07 |
| Ralb | -10.59901242 | 0.397436977 | 0.015279882 | 0.050283554 | 0.318266934 | 6.21121E-05 | 0.5 | 7.33199E-07 | FALSE | 0 | relative convergence (4) | 13129.4026044359 | 13184.644988553 | -6557.70130221795 | 13115.4026044359 | 19760 | 8.42512E-05 |
| Gpr39 | -12.77716456 | 1.008028327 | 0.148694123 | -0.255262311 | 20.53691645 | 0.000159956 | 0.5 | 0.000369195 | FALSE | 0 | relative convergence (4) | 2737.25306076821 | 2792.49544488528 | -1361.6265303841 | 2723.25306076821 | 19760 | 0.011084623 |
| Mcm6 | -10.49422921 | 0.226037219 | -0.324889306 | 0.028472488 | 0.897554264 | 0.19783568 | 0.5 | 0.001288054 | FALSE | 0 | relative convergence (4) | 14128.2841522273 | 14183.5265363443 | -7057.14207611364 | 14114.2841522273 | 19760 | 0.02612257 |
| Mapkapk2 | -9.505178432 | 0.214499908 | 0.002006463 | 0.280379294 | 0.301992409 | 0.164294368 | 0.5 | 2.57543E-09 | FALSE | 0 | relative convergence (4) | NA | NA | NA | NA | 19760 | 7.75083E-07 |
| Phlda3 | -10.93222511 | 0.787702998 | 0.192905318 | 0.24835109 | 1.264035259 | 0.000220388 | 0.5 | 0.000215157 | FALSE | 0 | relative convergence (4) | 12296.3051247067 | 12351.5475088238 | -6141.15256235337 | 12282.3051247067 | 19760 | 0.00751266 |
| Atp23 | -12.49180031 | 0.434578992 | 0.316308066 | 0.470766587 | 0.406120051 | 0.000521382 | 0.5 | 0.000439819 | FALSE | 0 | relative convergence (4) | 3495.38733468377 | 3550.62971880085 | -1740.69366734189 | 3481.38733468377 | 19760 | 0.0125725 |
| Tspan31 | -10.34752245 | 0.268646418 | -0.162026415 | 0.331924546 | 0.03532044 | 0.078409758 | 0.5 | 0.001487923 | FALSE | 0 | relative convergence (4) | 15409.8668784217 | 15465.1092625388 | -7697.93343921086 | 15395.8668784217 | 19760 | 0.028659961 |
| Ndufa4l2 | -13.07400544 | 1.157620629 | 0.308603344 | 0.825449928 | 35.17001355 | 0.356806468 | 0.5 | 6.08542E-06 | FALSE | 0 | relative convergence (4) | 2400.85895756566 | 2456.10134168274 | -1193.42947878283 | 2386.85895756566 | 19760 | 0.000474581 |
| Lrp1 | -8.679819837 | 0.373298333 | 0.051542678 | 0.065519571 | 0.186306878 | 0.000342857 | 0.5 | 0.003356748 | FALSE | 0 | relative convergence (4) | 39776.2324118727 | 39831.4747959898 | -19881.1162059364 | 39762.2324118727 | 19760 | 0.048324936 |
| Nab2 | -11.35532762 | 0.473442419 | -0.101855691 | 0.494395247 | 0.176605956 | 0.175509513 | 0.5 | 1.86604E-05 | FALSE | 0 | relative convergence (4) | 8074.62322048041 | 8129.86560459748 | -4030.31161024021 | 8060.62322048041 | 19760 | 0.001173472 |
| Slc35e4 | -11.21549029 | 0.446535089 | 0.115045669 | -0.025848294 | 0.164472911 | 9.63089E-05 | 0.5 | 0.00063516 | FALSE | 0 | relative convergence (4) | 8709.77853461835 | 8765.02091873543 | -4347.88926730918 | 8695.77853461835 | 19760 | 0.015929398 |
| Gas2l1 | -10.35766217 | 0.432334871 | -0.210724578 | -0.073355479 | 0.055424125 | 0.000143644 | 0.5 | 0.001279142 | FALSE | 0 | relative convergence (4) | 15423.301316524 | 15478.5437006411 | -7704.65065826199 | 15409.301316524 | 19760 | 0.025994142 |
| Hif1a | -9.706620683 | 0.239709005 | -0.202810159 | -0.233812294 | 0.715168888 | 0.145988911 | 0.5 | 0.001891728 | FALSE | 0 | relative convergence (4) | 23162.3953031862 | 23217.6376873033 | -11574.1976515931 | 23148.3953031862 | 19760 | 0.033489412 |
| Ltbp2 | -11.37808551 | 1.654873214 | 0.483597526 | 0.546139305 | 2.878812699 | 0.000252966 | 0.5 | 3.50071E-17 | FALSE | 0 | relative convergence (4) | 10643.8298046112 | 10699.0721887283 | -5314.91490230562 | 10629.8298046112 | 19760 | 5.53112E-14 |
| Pgf | -13.4346138 | 1.334500582 | 1.090317821 | 1.280591989 | 10.08021785 | 5.97441E-05 | 0.5 | 1.06987E-06 | FALSE | 0 | relative convergence (4) | 2594.38427072048 | 2649.62665483756 | -1290.19213536024 | 2580.38427072048 | 19760 | 0.000110846 |
| Pxdc1 | -10.59721554 | 0.533540652 | 0.234574112 | 0.839751813 | 0.463865458 | 0.235743318 | 0.5 | 8.23405E-06 | FALSE | 0 | relative convergence (4) | 15387.5990831005 | 15442.8414672175 | -7686.79954155023 | 15373.5990831005 | 19760 | 0.00058471 |
| Gmpr | -11.74106768 | 0.549196592 | 0.321812387 | 0.209595743 | 0.634437701 | 0.060543198 | 0.5 | 8.4686E-05 | FALSE | 0 | relative convergence (4) | 6294.05239760303 | 6349.2947817201 | -3140.02619880151 | 6280.05239760303 | 19760 | 0.003665861 |
| Kif13a | -8.845845196 | 0.272743623 | 0.163932161 | 0.266983916 | 0.39742573 | 0.074196573 | 0.5 | 0.003075165 | FALSE | 0 | relative convergence (4) | 38009.2084913562 | 38064.4508754733 | -18997.6042456781 | 37995.2084913562 | 19760 | 0.046495311 |
| Bicd2 | -10.2386742 | 0.233007149 | 0.025940498 | 0.339774292 | 0.103844136 | 0.136213925 | 0.5 | 0.000666708 | FALSE | 0 | relative convergence (4) | 16216.9289836015 | 16272.1713677186 | -8101.46449180075 | 16202.9289836015 | 19760 | 0.016318899 |
| Prelid1 | -10.02622082 | 0.569832517 | -0.173087653 | 0.643201776 | 2.43601E-07 | 1.69949E-36 | 0.5 | 0.000644453 | FALSE | 1 | singular convergence (7) | 20697.8128140186 | 20753.0551981356 | -10341.9064070093 | 20683.8128140186 | 19760 | 0.016008339 |
| Tmem222 | -11.22168408 | 0.414126824 | 0.031345008 | 0.419743823 | 1.32527E-06 | 0.124716405 | 0.5 | 0.003230426 | FALSE | 1 | false convergence (8) | 9158.34477667308 | 9213.58716079016 | -4572.17238833654 | 9144.34477667308 | 19760 | 0.047645961 |
| Trnp1 | -13.08491728 | 1.269373991 | 1.216878993 | 1.509217301 | 3.611735956 | 7.1598E-05 | 0.5 | 3.82327E-10 | FALSE | 0 | relative convergence (4) | 3337.20168566937 | 3392.44406978644 | -1661.60084283468 | 3323.20168566937 | 19760 | 1.61087E-07 |
| Ldlrap1 | -11.29221329 | 0.482084821 | 0.160673764 | 0.125550676 | 0.460996804 | 0.037810374 | 0.5 | 6.19332E-11 | FALSE | 0 | relative convergence (4) | 8375.21559088965 | 8430.45797500672 | -4180.60779544482 | 8361.21559088965 | 19760 | 3.26181E-08 |
| Maco1 | -9.468526768 | 0.345895237 | -0.264561714 | -0.052509716 | 0.500170008 | 0.19725124 | 0.5 | 5.84889E-06 | FALSE | 0 | relative convergence (4) | 27731.6365559159 | 27786.878940033 | -13858.818277958 | 27717.6365559159 | 19760 | 0.000460646 |
| Nbl1 | -10.80802932 | 0.647279815 | -0.175239719 | 0.437725467 | 0.45090601 | 0.000224572 | 0.5 | 0.00081964 | FALSE | 0 | relative convergence (4) | 12999.3148648598 | 13054.5572489769 | -6492.65743242992 | 12985.3148648598 | 19760 | 0.018974814 |
| Capzb | -8.378408678 | 0.191114362 | -0.22795741 | 0.138270266 | 0.034774907 | 0.088352579 | 0.5 | 0.001002448 | FALSE | 0 | relative convergence (4) | 44615.6626767347 | 44670.9050608518 | -22300.8313383674 | 44601.6626767347 | 19760 | 0.021884185 |
| Gask1b | -9.016105074 | 0.38411962 | -0.147717239 | 0.159368041 | 0.640233606 | 0.10409768 | 0.5 | 6.65239E-07 | FALSE | 0 | relative convergence (4) | 35042.6417695397 | 35097.8841536568 | -17514.3208847699 | 35028.6417695397 | 19760 | 7.78576E-05 |
| Pdgfc | -10.06298677 | 0.604510106 | 0.292178162 | 0.343078428 | 1.973331753 | 0.000157257 | 0.5 | 0.000264676 | FALSE | 0 | relative convergence (4) | 19449.9720101496 | 19505.2143942666 | -9717.98600507478 | 19435.9720101496 | 19760 | 0.008689634 |
| Arfip1 | -11.15696026 | 0.398073275 | -0.510463576 | 0.209857602 | 0.019980033 | 0.240087275 | 0.5 | 3.81598E-05 | FALSE | 0 | relative convergence (4) | 9196.15287385677 | 9251.39525797384 | -4591.07643692838 | 9182.15287385677 | 19760 | 0.002043813 |
| Zbtb7b | -11.25849112 | 0.503042423 | -0.078836412 | 0.111117536 | 0.047126628 | 0.20263726 | 0.5 | 8.00057E-12 | FALSE | 0 | relative convergence (4) | 8725.14657366602 | 8780.38895778309 | -4355.57328683301 | 8711.14657366602 | 19760 | 4.59669E-09 |
| Spry2 | -12.11630204 | 0.720966965 | 0.050013922 | 0.20856397 | 1.187088961 | 0.000238466 | 0.5 | 0.000590528 | FALSE | 0 | relative convergence (4) | 5301.90538995546 | 5357.14777407253 | -2643.95269497773 | 5287.90538995546 | 19760 | 0.015048933 |
| Dnajc3 | -8.899467875 | 0.300243735 | 0.172032792 | 0.173480734 | 0.17518189 | 0.000347601 | 0.5 | 0.000333371 | FALSE | 0 | relative convergence (4) | 35851.9547913889 | 35907.197175506 | -17918.9773956945 | 35837.9547913889 | 19760 | 0.010302711 |
| Brix1 | -10.72945592 | 0.219317745 | 0.264535377 | 0.353371434 | 0.066288722 | 0.095809245 | 0.5 | 0.000451547 | FALSE | 0 | relative convergence (4) | 12138.2823035368 | 12193.5246876538 | -6062.14115176839 | 12124.2823035368 | 19760 | 0.012768105 |
| Rgs7bp | -9.873160808 | 0.336925168 | 0.364127629 | 0.334373701 | 1.777690493 | 0.118625493 | 0.5 | 3.05547E-08 | FALSE | 0 | relative convergence (4) | 21846.130967494 | 21901.3733516111 | -10916.065483747 | 21832.130967494 | 19760 | 6.51266E-06 |
| Fst | -13.14834578 | 0.933275123 | 0.298124909 | 0.128168131 | 19.43012151 | 0.000140047 | 0.5 | 0.001774874 | FALSE | 0 | relative convergence (4) | 2327.15783915131 | 2382.40022326838 | -1156.57891957565 | 2313.15783915131 | 19760 | 0.032466577 |
| Nr1d2 | -9.902066104 | 0.586542746 | -0.398472947 | -0.732001706 | 0.335051848 | 0.000144217 | 0.5 | 1.34752E-06 | FALSE | 0 | relative convergence (4) | 20455.8070225145 | 20511.0494066316 | -10220.9035112572 | 20441.8070225145 | 19760 | 0.000134116 |
| Atp6v0b | -9.029928579 | 0.36116621 | 0.243578989 | 0.302609918 | 0.042596867 | 0.000124892 | 0.5 | 0.003148022 | FALSE | 0 | relative convergence (4) | 34330.3472257043 | 34385.5896098214 | -17158.1736128522 | 34316.3472257043 | 19760 | 0.047034279 |
| Ppcs | -11.96609224 | 0.333163044 | 0.403598115 | 0.41186273 | 0.256939996 | 5.58067E-05 | 0.5 | 0.000700435 | FALSE | 0 | relative convergence (4) | NA | NA | NA | NA | 19760 | 0.016895984 |
| Fhl3 | -11.83519002 | 0.506983674 | 0.025812341 | 0.028848861 | 0.507593123 | 0.278502335 | 0.5 | 0.00149808 | FALSE | 0 | relative convergence (4) | 6022.52318105597 | 6077.76556517305 | -3004.26159052799 | 6008.52318105597 | 19760 | 0.02873404 |
| Map7d1 | -9.157788037 | 0.405714516 | 0.035445243 | 0.1289819 | 0.114763735 | 0.091203076 | 0.5 | 0.001634218 | FALSE | 0 | relative convergence (4) | 31487.1157691659 | 31542.358153283 | -15736.5578845829 | 31473.1157691659 | 19760 | 0.030693186 |
| Gja4 | -12.99178987 | 1.364784119 | 0.94027729 | 0.977615637 | 2.678386008 | 0.087070895 | 0.5 | 1.37165E-19 | FALSE | 0 | relative convergence (4) | NA | NA | NA | NA | 19760 | 3.46754E-16 |
| Col16a1 | -11.69415211 | 0.370812473 | 0.458656166 | 0.788296317 | 0.917276512 | 0.354347801 | 0.5 | 0.000108997 | FALSE | 0 | relative convergence (4) | NA | NA | NA | NA | 19760 | 0.004458666 |
| F8a | -11.80051899 | 0.325688879 | -0.083233101 | 0.289011809 | 0.009933943 | 0.085875363 | 0.5 | 0.000826882 | FALSE | 0 | relative convergence (4) | 5598.58337652192 | 5653.825760639 | -2792.29168826096 | 5584.58337652192 | 19760 | 0.019072615 |
| Ubl4a | -10.93684694 | 0.281094355 | 0.001302713 | 0.199578326 | 0.171953201 | 5.63045E-05 | 0.5 | 0.000250589 | FALSE | 0 | relative convergence (4) | 10313.9429901706 | 10369.1853742877 | -5149.97149508532 | 10299.9429901706 | 19760 | 0.008401721 |
| Susd5 | -9.235922206 | 0.687831913 | 0.303109575 | 0.35120184 | 0.291498765 | 0.000472603 | 0.5 | 0.001424592 | FALSE | 0 | relative convergence (4) | 32886.0603202122 | 32941.3027043293 | -16436.0301601061 | 32872.0603202122 | 19760 | 0.027788342 |
| Cmtm8 | -12.73984342 | 0.520403628 | 0.061688169 | -0.135608585 | 8.683501067 | 0.140560952 | 0.5 | 0.002790259 | FALSE | 0 | relative convergence (4) | 2692.23640783469 | 2747.47879195176 | -1339.11820391734 | 2678.23640783469 | 19760 | 0.043595643 |
| Cmc1 | -10.93697318 | 0.302995424 | 0.036522009 | 0.119979328 | 0.370703624 | 7.43755E-05 | 0.5 | 0.001531071 | FALSE | 0 | relative convergence (4) | 10439.3158769646 | 10494.5582610817 | -5212.6579384823 | 10425.3158769646 | 19760 | 0.029189644 |
| Snrk | -9.7183765 | 0.221021984 | 0.319900365 | 0.329314003 | 0.226753028 | 0.127926613 | 0.5 | 7.83165E-06 | FALSE | 0 | relative convergence (4) | 22691.9375011044 | 22747.1798852215 | -11338.9687505522 | 22677.9375011044 | 19760 | 0.000565669 |
| Plec | -9.245081801 | 0.267355707 | 0.340362109 | 0.183373531 | 0.213372248 | 0.112018218 | 0.5 | 0.000552352 | FALSE | 0 | relative convergence (4) | 30089.871659709 | 30145.114043826 | -15037.9358298545 | 30075.871659709 | 19760 | 0.014515042 |
| Scx | -12.51402649 | 1.622850148 | 0.05244397 | 1.150592893 | 4.909764344 | 0.000618937 | 0.5 | 2.29819E-08 | FALSE | 0 | relative convergence (4) | 5141.61502564004 | 5196.85740975711 | -2563.80751282002 | 5127.61502564004 | 19760 | 5.37947E-06 |
| Cdc42ep1 | -12.56522016 | 0.715348032 | 0.517757285 | 0.915362805 | 1.137277155 | 0.240092674 | 0.5 | 0.000542495 | FALSE | 0 | relative convergence (4) | 4027.04868483331 | 4082.29106895038 | -2006.52434241665 | 4013.04868483331 | 19760 | 0.014287889 |
| Maff | -11.73931123 | 1.163668495 | 0.200502414 | 0.574225162 | 0.75829724 | 0.413173921 | 0.5 | 0.000926371 | FALSE | 0 | relative convergence (4) | 9323.10166873922 | 9378.34405285629 | -4654.55083436961 | 9309.10166873922 | 19760 | 0.020761215 |
| Csnk1e | -10.20184408 | 0.334887437 | 0.084941075 | 0.248630293 | 0.167128881 | 0.149571242 | 0.5 | 0.002554305 | FALSE | 0 | relative convergence (4) | 17658.1878806181 | 17713.4302647352 | -8822.09394030904 | 17644.1878806181 | 19760 | 0.041129195 |
| Pdgfb | -13.08369436 | 1.214140168 | 0.186885387 | 0.207013532 | 10.14602047 | 9.8799E-05 | 0.5 | 4.83557E-05 | FALSE | 0 | relative convergence (4) | 2404.92964300592 | 2460.172027123 | -1195.46482150296 | 2390.92964300592 | 19760 | 0.002415871 |
| Tef | -10.3128254 | 0.522169257 | 0.089358043 | -0.21746531 | 0.439126048 | 0.192596446 | 0.5 | 0.000418915 | FALSE | 0 | relative convergence (4) | 15833.5801906628 | 15888.8225747798 | -7909.79009533139 | 15819.5801906628 | 19760 | 0.012144681 |
| Naga | -11.55327724 | 0.548645657 | -0.140734051 | 0.326643911 | 0.254111314 | 0.000166949 | 0.5 | 0.001820512 | FALSE | 0 | relative convergence (4) | 7343.22780810568 | 7398.47019222275 | -3664.61390405284 | 7329.22780810568 | 19760 | 0.03290476 |
| Serhl | -11.53540685 | 0.311515206 | -0.039326785 | 0.588465735 | 0.012581327 | 0.133746659 | 0.5 | 0.000493137 | FALSE | 0 | relative convergence (4) | 7097.17112895513 | 7152.4135130722 | -3541.58556447756 | 7083.17112895513 | 19760 | 0.013433732 |
| Atp8b2 | -11.32427235 | 0.428237854 | -0.328505337 | 0.228432315 | 0.37867203 | 0.104905605 | 0.5 | 0.001172014 | FALSE | 0 | relative convergence (4) | 7990.81794984575 | 8046.06033396282 | -3988.40897492288 | 7976.81794984575 | 19760 | 0.024445962 |
| Npr1 | -9.76880568 | 0.330815659 | 0.413050474 | 0.727069982 | 0.460909034 | 0.180167066 | 0.5 | 0.001357619 | FALSE | 0 | relative convergence (4) | 23742.1411843199 | 23797.383568437 | -11864.07059216 | 23728.1411843199 | 19760 | 0.027058402 |
| S100a4 | -11.38030878 | 1.336344503 | -0.375914032 | 1.453640923 | 1.264567743 | 0.000642116 | 0.5 | 0.001950969 | FALSE | 0 | relative convergence (4) | 14314.3332494825 | 14369.5756335996 | -7150.16662474125 | 14300.3332494825 | 19760 | 0.034250338 |
| Cdc42se1 | -9.916423149 | 0.258866755 | -0.047561947 | 0.370509831 | 0.100866424 | 0.067754497 | 0.5 | 0.002335534 | FALSE | 0 | relative convergence (4) | 20243.2136045791 | 20298.4559886961 | -10114.6068022895 | 20229.2136045791 | 19760 | 0.038690898 |
| Aph1a | -10.52658852 | 0.279923185 | 0.093252356 | 0.27872977 | 8.7975E-10 | 0.137333133 | 0.5 | 0.003289636 | FALSE | 1 | false convergence (8) | 13778.2679682865 | 13833.5103524036 | -6882.13398414327 | 13764.2679682865 | 19760 | 0.047991161 |
| Plekho1 | -9.602821993 | 0.369584808 | 0.21978809 | 0.464330992 | 0.252184463 | 0.000159386 | 0.5 | 0.00086779 | FALSE | 0 | relative convergence (4) | 25583.4829747255 | 25638.7253588425 | -12784.7414873627 | 25569.4829747255 | 19760 | 0.019803641 |
| Akt1 | -10.02222874 | 0.417271011 | -0.079110123 | 0.416252688 | 0.095632954 | 0.070597908 | 0.5 | 0.003379851 | FALSE | 0 | relative convergence (4) | 19876.8283631892 | 19932.0707473063 | -9931.41418159461 | 19862.8283631892 | 19760 | 0.048491851 |
| Clba1 | -13.22308466 | 0.759423793 | 0.711819479 | 0.668188629 | 1.90987E-08 | 0.314497701 | 0.5 | 1.02347E-05 | FALSE | 1 | false convergence (8) | NA | NA | NA | NA | 19760 | 0.000700579 |
| Akr1c14 | -12.11176519 | 0.461584546 | 0.218046842 | 0.375018344 | 0.665537108 | 7.6258E-05 | 0.5 | 0.000476594 | FALSE | 0 | relative convergence (4) | 4658.92858443617 | 4714.17096855324 | -2322.46429221808 | 4644.92858443617 | 19760 | 0.013181948 |
| Klf6 | -8.370582192 | 0.647968023 | -0.078714954 | 0.296041598 | 0.444818882 | 0.179174555 | 0.5 | 0.002862358 | FALSE | 0 | relative convergence (4) | 51662.4199981849 | 51717.662382302 | -25824.2099990925 | 51648.4199981849 | 19760 | 0.044447422 |
| Serpinb6b | -10.22249818 | 0.219671435 | 0.240312658 | 0.221689533 | 0.882427766 | 8.03241E-05 | 0.5 | 0.002418881 | FALSE | 0 | relative convergence (4) | 16856.599651 | 16911.842035117 | -8421.29982549999 | 16842.599651 | 19760 | 0.039707351 |
| Sulf1 | -8.587648328 | 0.555744472 | 0.21107332 | 0.378168017 | 0.939527312 | 0.000306612 | 0.5 | 5.37594E-05 | FALSE | 0 | relative convergence (4) | 46652.8344148553 | 46708.0767989724 | -23319.4172074277 | 46638.8344148553 | 19760 | 0.002613534 |
| Sbspon | -12.9884304 | 1.030448855 | 0.678848512 | 1.03669757 | 5.205025874 | 5.58646E-05 | 0.5 | 2.00367E-09 | FALSE | 0 | relative convergence (4) | 2972.62891158303 | 3027.87129570011 | -1479.31445579152 | 2958.62891158303 | 19760 | 6.84497E-07 |
| Col19a1 | -11.79657055 | 0.714104768 | 0.833701351 | 0.511601828 | 9.245914624 | 0.000244816 | 0.5 | 0.002042828 | FALSE | 0 | relative convergence (4) | 6139.28168124427 | 6194.52406536134 | -3062.64084062213 | 6125.28168124427 | 19760 | 0.035275055 |
| Ccdc115 | -10.97864464 | 0.307860805 | -0.147729115 | 0.019743904 | 0.057102889 | 0.100010523 | 0.5 | 0.000482976 | FALSE | 0 | relative convergence (4) | 10013.2810522986 | 10068.5234364157 | -4999.64052614932 | 9999.28105229864 | 19760 | 0.013219656 |

| | | | | | | | | | | | | | | | | |
|---|---|---|---|---|---|---|---|---|---|---|---|---|---|---|---|---|
| Sema4c | -11.72322624 | 0.529416882 | 0.143530684 | -0.05205056 | 0.178277264 | 0.000174144 | 0.5 | 0.002151039 | FALSE | 0 | relative convergence (4) | 6280.5472312256 | 6335.78961534267 | -3133.2736156128 | 6266.5472312256 | 19760 | 0.036544526 |
| Actr1b | -10.13071607 | 0.210775994 | -0.098163603 | 0.097985868 | 0.07478939 | 0.083091973 | 0.5 | 0.000856601 | FALSE | 0 | relative convergence (4) | 17114.1264616688 | 17169.3688457859 | -8550.0632308344 | 17100.1264616688 | 19760 | 0.019650525 |
| 4933424G06Rik | -12.53752374 | 1.290239616 | 1.735142306 | 0.680079558 | 0.611494127 | 0.592469114 | 0.5 | 9.08373E-06 | FALSE | 0 | relative convergence (4) | 6350.79349047944 | 6406.03587459651 | -3168.39674523972 | 6336.79349047944 | 19760 | 0.00063788 |
| Inpp4a | -9.483516197 | 0.171420665 | 0.02049743 | 0.174520592 | 0.479187376 | 7.76173E-05 | 0.5 | 0.002067118 | FALSE | 0 | relative convergence (4) | 25999.1804162075 | 26054.4228003245 | -12992.5902081037 | 25985.1804162075 | 19760 | 0.035597229 |
| Sh3gl1 | -10.82316841 | 0.404839965 | 0.083903669 | 0.327876237 | 0.11551464 | 0.000156346 | 0.5 | 0.003466476 | FALSE | 0 | relative convergence (4) | 11775.021941881 | 11830.2643259981 | -5880.51097094049 | 11761.021941881 | 19760 | 0.049121358 |
| Ticam1 | -12.15716398 | 0.335164725 | -0.020481798 | 0.415945039 | 0.335324624 | 0.000168698 | 0.5 | 0.00194604 | FALSE | 0 | relative convergence (4) | 4279.55650773445 | 4334.79889185152 | -2132.77825386722 | 4265.55650773445 | 19760 | 0.034211336 |
| Washc1 | -11.16501828 | 0.302255261 | 0.035257832 | 0.30071134 | 0.015564501 | 0.21427037 | 0.5 | 4.00852E-05 | FALSE | 0 | relative convergence (4) | NA | NA | NA | NA | 19760 | 0.002119988 |
| Lbh | -9.506106853 | 0.473138119 | -0.081503454 | 0.392458502 | 0.610708508 | 0.117581972 | 0.5 | 0.000787912 | FALSE | 0 | relative convergence (4) | 27780.4605658935 | 27835.7029500106 | -13883.2302829468 | 27766.4605658935 | 19760 | 0.018580603 |
| Fez2 | -10.16791665 | 0.442872363 | 0.315689088 | 0.428363971 | 0.234932397 | 0.000243577 | 0.5 | 1.37034E-07 | FALSE | 0 | relative convergence (4) | 18311.2903356596 | 18366.5327197767 | -9148.6451678298 | 18297.2903356596 | 19760 | 2.05818E-05 |
| Odc1 | -9.810433579 | 0.305485716 | -0.258961333 | 0.14424977 | 0.615618688 | 0.337685153 | 0.5 | 0.002306849 | FALSE | 0 | relative convergence (4) | 22857.6019111249 | 22912.844295242 | -11421.8009555625 | 22843.6019111249 | 19760 | 0.038467779 |
| Egln3 | -11.33055824 | 0.911795115 | 0.454010282 | 0.457930404 | 1.780913294 | 0.000169088 | 0.5 | 0.000185652 | FALSE | 0 | relative convergence (4) | 10012.3711215698 | 10067.6135056868 | -4999.18556078488 | 9998.37112156976 | 19760 | 0.006782217 |
| Dusp12 | -11.20300039 | 0.234472518 | 0.083847947 | 0.264068541 | 0.000484306 | 0.001105177 | 0.5 | 0.000666843 | FALSE | 0 | relative convergence (4) | NA | NA | NA | NA | 19760 | 0.016318899 |
| Sdhc | -9.41800312 | 0.334786496 | 0.172974049 | 0.327803309 | 0.104562267 | 0.000115595 | 0.5 | 0.000573216 | FALSE | 0 | relative convergence (4) | 27849.4774052977 | 27904.7197894148 | -13917.7387026489 | 27835.4774052977 | 19760 | 0.014816861 |
| Casq1 | -11.60233668 | 0.878008833 | -0.273414491 | -0.488001042 | 2.045816228 | 0.000351134 | 0.5 | 0.003031014 | FALSE | 0 | relative convergence (4) | 7273.02732954651 | 7328.26971366358 | -3629.51366477325 | 7259.02732954651 | 19760 | 0.046198077 |
| Kif26b | -11.11642202 | 1.084050488 | 0.26472945 | 0.652942598 | 2.317568722 | 9.74071E-05 | 0.5 | 8.67051E-08 | FALSE | 0 | relative convergence (4) | 11183.9391010177 | 11239.1814851347 | -5584.96955050884 | 11169.9391010177 | 19760 | 1.48102E-05 |
| Rab3gap2 | -9.904656695 | 0.27010676 | -0.128986587 | 0.061401344 | 0.786843011 | 0.000138793 | 0.5 | 0.001399695 | FALSE | 0 | relative convergence (4) | 20358.1938099601 | 20413.4361940772 | -10172.09690498 | 20344.1938099601 | 19760 | 0.027509351 |
| Tgfb2 | -9.040589681 | 0.569308802 | -0.155243904 | 0.258569634 | 1.379318231 | 0.273261869 | 0.5 | 7.08387E-06 | FALSE | 0 | relative convergence (4) | 35942.879837522 | 35998.122221639 | -17964.439918761 | 35928.879837522 | 19760 | 0.000520582 |
| Layn | -11.50609226 | 0.592177 | 0.347451041 | 0.11612856 | 1.261445188 | 7.15337E-05 | 0.5 | 0.000112199 | FALSE | 0 | relative convergence (4) | 7498.44894710775 | 7553.69133122483 | -3742.22447355388 | 7484.44894710775 | 19760 | 0.004516553 |
| Stoml1 | -11.25939787 | 0.3104892 | 0.017199671 | 0.251492742 | 4.05045E-07 | 4.32391E-07 | 0.5 | 6.83008E-06 | FALSE | 1 | false convergence (8) | NA | NA | NA | NA | 19760 | 0.000507837 |
| Uaca | -10.51385979 | 0.467802601 | 0.27195983 | 0.435961239 | 1.047846246 | 0.284698061 | 0.5 | 8.74935E-07 | FALSE | 0 | relative convergence (4) | 14218.7892604841 | 14274.0316446011 | -7102.39463024203 | 14204.7892604841 | 19760 | 9.37219E-05 |
| Pusl1 | -12.29501654 | 0.520608668 | -0.072560312 | -0.089048602 | 1.27592E-06 | 1.88211E-71 | 0.5 | 0.0003027 | FALSE | 1 | singular convergence (7) | 3930.40180790766 | 3985.64419202473 | -1958.20090395383 | 3916.40180790766 | 19760 | 0.009613402 |
| Sdf4 | -9.257465994 | 0.272893256 | -0.193685075 | 0.313105015 | 0.105602029 | 0.000119333 | 0.5 | 0.000240463 | FALSE | 0 | relative convergence (4) | 29292.7672227518 | 29348.0096068689 | -14639.3836113759 | 29278.7672227518 | 19760 | 0.008083643 |
| Chpf2 | -11.39494045 | 0.439273496 | -0.071441692 | 0.424432873 | 0.457776286 | 8.31307E-05 | 0.5 | 1.95964E-06 | FALSE | 0 | relative convergence (4) | 7840.77418169197 | 7896.01656580905 | -3913.38709084599 | 7826.77418169197 | 19760 | 0.000187651 |
| Tmem214 | -10.61750919 | 0.309501521 | -0.286713044 | 0.158279179 | 0.13301904 | 0.000276502 | 0.5 | 0.000125416 | FALSE | 0 | relative convergence (4) | 12619.3131781985 | 12674.5555623156 | -6302.65658909924 | 12605.3131781985 | 19760 | 0.00495394 |
| Cgref1 | -13.16384739 | 1.133738717 | 0.444635632 | 0.58153538 | 1.786159199 | 0.258935544 | 0.5 | 2.12384E-05 | FALSE | 0 | relative convergence (4) | 2751.34964110386 | 2806.59202522093 | -1368.67482055193 | 2737.34964110386 | 19760 | 0.001299175 |
| Synpo2l | -12.57475481 | 0.883091225 | -0.234595221 | 0.478622501 | 1.395684448 | 0.25719771 | 0.5 | 0.000890084 | FALSE | 0 | relative convergence (4) | 3441.03214702868 | 3496.27453114575 | -1713.51607351434 | 3427.03214702868 | 19760 | 0.020162478 |
| Comtd1 | -12.23392054 | 0.357365602 | 0.575822368 | 0.627627442 | 0.593879443 | 0.298698541 | 0.5 | 0.003452202 | FALSE | 0 | relative convergence (4) | 4549.05731493823 | 4604.29969905531 | -2267.52865746912 | 4535.05731493823 | 19760 | 0.049029022 |
| Nisch | -8.862401783 | 0.128849421 | -0.001213812 | 0.139398181 | 0.081037015 | 0.081370389 | 0.5 | 0.001073899 | FALSE | 0 | relative convergence (4) | 35232.760129187 | 35288.0025133041 | -17609.3800645935 | 35218.760129187 | 19760 | 0.022904515 |
| Psmc1 | -9.920129904 | 0.318776697 | -0.07985088 | 0.290371538 | 0.09842229 | 0.000123706 | 0.5 | 0.000315564 | FALSE | 0 | relative convergence (4) | 20334.1050902509 | 20389.347474368 | -10160.0525451255 | 20320.1050902509 | 19760 | 0.009885647 |
| Fbln5 | -8.027337139 | 0.319055326 | -0.216652184 | 0.094863404 | 0.561704713 | 0.000193011 | 0.5 | 0.002589566 | FALSE | 0 | relative convergence (4) | 55891.6121911313 | 55946.8545752484 | -27938.8060955657 | 55877.6121911313 | 19760 | 0.041485569 |
| Anxa5 | -8.290993244 | 0.303907167 | -0.042246356 | 0.069184137 | 0.211004866 | 0.000117792 | 0.5 | 0.001219052 | FALSE | 0 | relative convergence (4) | 48114.1965385895 | 48169.4389227066 | -24050.0982692947 | 48100.1965385895 | 19760 | 0.025136724 |
| Fgf2 | -10.41409379 | 0.619478607 | 0.0975789 | 0.261901296 | 0.422883782 | 0.124119974 | 0.5 | 5.48591E-07 | FALSE | 0 | relative convergence (4) | 16065.7019635355 | 16120.9443476526 | -8025.85098176777 | 16051.7019635355 | 19760 | 6.6986E-05 |
| Postn | -7.923672561 | 0.822565252 | 0.008769613 | 0.069225061 | 0.796382326 | 0.000349781 | 0.5 | 8.55355E-07 | FALSE | 0 | relative convergence (4) | 62508.8379969188 | 62564.0803810359 | -31247.4189984594 | 62494.8379969188 | 19760 | 9.24076E-05 |
| Alg5 | -10.66819363 | 0.183711264 | 0.226763762 | 0.283968641 | 0.120511707 | 1.34339E-11 | 0.5 | 0.000719295 | FALSE | 0 | relative convergence (4) | 12460.5137500437 | 12515.7561341608 | -6223.25687502187 | 12446.5137500437 | 19760 | 0.017219481 |
| Dclk1 | -11.5064197 | 0.630654844 | -0.117550221 | -0.021168313 | 11.35777822 | 0.208225836 | 0.5 | 2.94436E-08 | FALSE | 0 | relative convergence (4) | 6474.27846893474 | 6529.52085305181 | -3230.13923446737 | 6460.27846893474 | 19760 | 6.48383E-06 |
| Pfn2 | -11.48676917 | 0.389206887 | -0.168368931 | 0.129905743 | 0.441981682 | 0.235247589 | 0.5 | 0.00025259 | FALSE | 0 | relative convergence (4) | 7388.21889087782 | 7443.4612749949 | -3687.10944543891 | 7374.21889087782 | 19760 | 0.008446383 |
| Fpgs | -11.90015584 | 0.348742667 | -0.11417193 | 0.32628888 | 0.08423612 | 0.186916619 | 0.5 | 0.000793604 | FALSE | 0 | relative convergence (4) | 5289.88745526996 | 5345.12983938703 | -2637.94372763498 | 5275.88745526996 | 19760 | 0.018645272 |
| Ptrh1 | -13.00918314 | 0.582070702 | 0.198823211 | 0.360109163 | 1.032871947 | 5.6081E-05 | 0.5 | 0.002979194 | FALSE | 0 | relative convergence (4) | 2404.70866754787 | 2459.95105166495 | -1195.35433377394 | 2390.70866754787 | 19760 | 0.045836632 |
| Gorasp2 | -9.453322485 | 0.124732252 | -0.341626372 | -0.131615073 | 0.125973426 | 0.100807161 | 0.5 | 0.000459831 | FALSE | 0 | relative convergence (4) | 25491.5018537549 | 25546.744237872 | -12738.7509268774 | 25477.5018537549 | 19760 | 0.012859003 |
| Mapk3 | -10.25470859 | 0.513388613 | 0.066968964 | 0.430812968 | 0.090651994 | 0.000143703 | 0.5 | 0.000129506 | FALSE | 0 | relative convergence (4) | 17396.3113748156 | 17451.5537589327 | -8691.15568740779 | 17382.3113748156 | 19760 | 0.005083733 |
| Ykt6 | -10.78944969 | 0.23072779 | -0.358501848 | -0.000150603 | 0.209492086 | 0.067290276 | 0.5 | 0.000625812 | FALSE | 0 | relative convergence (4) | 10916.9883371843 | 10972.2307213014 | -5451.49416859214 | 10902.9883371843 | 19760 | 0.015757487 |
| Rhbdf1 | -11.19380451 | 0.506809589 | 0.24794721 | 0.80049022 | 0.114628419 | 0.182609189 | 0.5 | 0.000715365 | FALSE | 0 | relative convergence (4) | 10004.285390363 | 10059.5277744801 | -4995.14269518152 | 9990.28539036303 | 19760 | 0.017190528 |
| Slit3 | -8.247083951 | 0.558582928 | 0.352339737 | 0.123582654 | 1.088228287 | 0.000188746 | 0.5 | 0.002664674 | FALSE | 0 | relative convergence (4) | 53510.7023422377 | 53565.9447263548 | -26748.3511711189 | 53496.7023422377 | 19760 | 0.042154537 |
| Odr4 | -9.878949597 | 0.143956202 | 0.148298424 | 0.097465616 | 0.197079863 | 4.30817E-05 | 0.5 | 0.001756923 | FALSE | 0 | relative convergence (4) | 20277.6203409813 | 20332.8627250983 | -10131.8101704906 | 20263.6203409813 | 19760 | 0.032278359 |
| Cryzl2 | -12.50633508 | 0.663257367 | 0.481663191 | 0.731129634 | 1.318930869 | 0.051295264 | 0.5 | 9.27731E-08 | FALSE | 0 | relative convergence (4) | 3787.53667483809 | 3842.77905895517 | -1886.76833741905 | 3773.53667483809 | 19760 | 1.50701E-05 |
| Dusp27 | -13.17796167 | 0.730745138 | 0.605561927 | 0.903000935 | 4.443506185 | 0.106152181 | 0.5 | 0.000345703 | FALSE | 0 | relative convergence (4) | 2269.80869344804 | 2325.05107756511 | -1127.90434672402 | 2255.80869344804 | 19760 | 0.010580347 |
| Rgs5 | -9.414145467 | 0.744634065 | 0.104207657 | 0.328994806 | 7.783178968 | 0.246962836 | 0.5 | 9.29958E-08 | FALSE | 0 | relative convergence (4) | 26102.4965847043 | 26157.7389688214 | -13044.2482923521 | 26088.4965847043 | 19760 | 1.50701E-05 |
| Rgs4 | -9.829969735 | 0.461045458 | -0.111119671 | 0.155869399 | 1.327527319 | 0.224708435 | 0.5 | 0.001314422 | FALSE | 0 | relative convergence (4) | 22347.4803166838 | 22402.7227008009 | -11166.7401583419 | 22333.4803166838 | 19760 | 0.02649808 |
| Ccn3 | -8.016391751 | 0.569047223 | 0.328263937 | 0.476338843 | 1.276174264 | 0.000258884 | 0.5 | 2.59063E-06 | FALSE | 0 | relative convergence (4) | 60077.1484625481 | 60132.3908466652 | -30031.5742312741 | 60063.1484625481 | 19760 | 0.00024437 |
| 9130401M01Rik | -11.37000809 | 0.538103054 | 0.114303303 | 0.399458162 | 1.07073E-07 | 0.118535736 | 0.5 | 0.002225655 | FALSE | 0 | relative convergence (4) | 8558.00364961096 | 8613.24603372804 | -4272.00182480548 | 8544.00364961096 | 19760 | 0.037509711 |
| Sqle | -11.65471741 | 0.350374224 | -0.455392641 | -0.062213845 | 1.847249876 | 0.369862374 | 0.5 | 0.000570885 | FALSE | 0 | relative convergence (4) | 6662.74466834711 | 6717.98705246419 | -3324.37233417356 | 6648.74466834711 | 19760 | 0.014816861 |
| Kcnq3 | -12.66359889 | 0.811216768 | 0.913291174 | 0.688457826 | 24.4182726 | 0.250150188 | 0.5 | 6.4078E-06 | FALSE | 0 | relative convergence (4) | NA | NA | NA | NA | 19760 | 0.000490876 |
| Lynx1 | -10.73457723 | 0.266235271 | 0.072159968 | 0.62102937 | 0.280858029 | 0.268696363 | 0.5 | 0.000934093 | FALSE | 0 | relative convergence (4) | 12576.0227244671 | 12631.2651085842 | -6281.01136223354 | 12562.0227244671 | 19760 | 0.020786854 |
| Bnip2 | -8.800956104 | 0.328573583 | 0.135098665 | 0.256819481 | 0.118581284 | 0.000150114 | 0.5 | 0.00330612 | FALSE | 0 | relative convergence (4) | 37801.8734111193 | 37857.1157952364 | -18893.9367055596 | 37787.8734111193 | 19760 | 0.048089022 |
| Mapk6 | -10.08873468 | 0.393892028 | -0.714596912 | -0.481253095 | 0.930696226 | 0.368862511 | 0.5 | 0.000811129 | FALSE | 0 | relative convergence (4) | 19858.3143235995 | 19913.5567077165 | -9922.15716179973 | 19844.3143235995 | 19760 | 0.018916356 |
| Cd109 | -10.71847167 | 0.716433249 | 0.414762004 | 0.292939731 | 1.910443149 | 0.113749678 | 0.5 | 1.70695E-28 | FALSE | 0 | relative convergence (4) | NA | NA | NA | NA | 19760 | 1.07879E-24 |
| Tent5a | -9.59731538 | 0.570467765 | -0.172525725 | 0.400627622 | 1.52181178 | 0.000163252 | 0.5 | 0.003536059 | FALSE | 0 | relative convergence (4) | 26363.6279725528 | 26418.8703566699 | -13174.8139862764 | 26349.6279725528 | 19760 | 0.049995288 |
| Tmeff2 | -12.57101486 | 0.555257103 | 0.322531671 | 0.438630805 | 33.01280327 | 0.198295466 | 0.5 | 0.003271011 | FALSE | 0 | relative convergence (4) | 2978.3345775044 | 3033.57696162148 | -1482.1672887522 | 2964.3345775044 | 19760 | 0.047964711 |
| Fn1 | -8.313385821 | 0.677321258 | -0.080867917 | 0.148684818 | 0.930014752 | 0.001455982 | 0.5 | 2.29662E-05 | FALSE | 0 | relative convergence (4) | 53246.3638901554 | 53301.6062742725 | -26616.1819450777 | 53232.3638901554 | 19760 | 0.001362875 |
| Igfbp2 | -13.20399039 | 3.084937119 | 0.874575653 | 1.286318079 | 12.11060498 | 0.000222518 | 0.5 | 4.56656E-26 | FALSE | 0 | relative convergence (4) | 5125.66739282979 | 5180.90977694687 | -2555.8336964149 | 5111.66739282979 | 19760 | 1.92404E-22 |
| Arpc1b | -8.552819936 | 0.426594149 | -0.04974746 | 0.3478814 | 0.1778625 | 0.000247558 | 0.5 | 0.001670157 | FALSE | 0 | relative convergence (4) | 43165.2413998353 | 43220.4837839524 | -21575.6206999176 | 43151.2413998353 | 19760 | 0.031275238 |
| Flt1 | -10.50038727 | 0.451970692 | 0.561370353 | 0.012687514 | 2.767855275 | 0.209232538 | 0.5 | 1.23933E-05 | FALSE | 0 | relative convergence (4) | 14691.6441943422 | 14746.8865784593 | -7338.82209717109 | 14677.6441943422 | 19760 | 0.000824481 |
| Hmgb1 | -8.742340424 | 0.282005088 | 0.176176772 | -0.105080553 | 0.092186174 | 0.178026181 | 0.5 | 0.001270079 | FALSE | 0 | relative convergence (4) | 37755.3452399194 | 37810.5876240365 | -18870.6726199597 | 37741.3452399194 | 19760 | 0.025893222 |
| Pon2 | -9.945515406 | 0.325601023 | 0.06298786 | 0.131059637 | 0.211911101 | 9.11497E-05 | 0.5 | 0.001837189 | FALSE | 0 | relative convergence (4) | 20232.0538084708 | 20287.2961925878 | -10109.0269042354 | 20218.0538084708 | 19760 | 0.032939108 |
| Fam241a | -10.99921153 | 0.335250295 | 0.210556591 | 0.228377343 | 0.535452362 | 6.53966E-05 | 0.5 | 0.001289595 | FALSE | 0 | relative convergence (4) | 10303.4812081614 | 10358.7235922784 | -5144.74060408068 | 10289.4812081614 | 19760 | 0.02612257 |
| Enpep | -12.17706651 | 0.797429293 | 0.209929532 | 0.686569658 | 5.710141072 | 9.72571E-05 | 0.5 | 0.001774149 | FALSE | 0 | relative convergence (4) | 4547.48226015439 | 4602.72464427146 | -2266.74113007719 | 4533.48226015439 | 19760 | 0.032466577 |
| Pla2g12a | -11.47768997 | 0.365590662 | 0.078354035 | 0.341934635 | 0.282102174 | 7.69573E-05 | 0.5 | 0.001249416 | FALSE | 0 | relative convergence (4) | 7352.58917912704 | 7407.83156324412 | -3669.29458956352 | 7338.58917912704 | 19760 | 0.025637365 |
| Casp6 | -11.72515313 | 0.299495543 | 0.060447311 | 0.236455672 | 0.480595015 | 7.73356E-05 | 0.5 | 0.000770079 | FALSE | 0 | relative convergence (4) | NA | NA | NA | NA | 19760 | 0.018194015 |
| Slc9b2 | -13.23389498 | 0.769032944 | 0.904762421 | 0.426498489 | 4.281464169 | 0.486374388 | 0.5 | 0.001858035 | FALSE | 0 | relative convergence (4) | 2474.12786937274 | 2529.37025348982 | -1230.06393468637 | 2460.12786937274 | 19760 | 0.033078264 |

### Table S9. Macrophage Differentially Expressed Genes: Hypoxia versus Normoxia

**Macrophages: Genes that positively correlate with chronic hypoxia**

| gene | Beta0 | Beta1 | Disp | Sigma | PiC | PiI | P.Beta | P.Pi | Hessian | Convergence | Message | AIC | BIC | logLik | deviance | df.resid | fdrP |
|---|---|---|---|---|---|---|---|---|---|---|---|---|---|---|---|---|---|
| Lpl | -11.85387923 | 1.538070501 | 0.408291341 | -0.777837306 | 4.293551656 | 0.758548981 | 0.5 | 5.03279E-05 | FALSE | 0 | relative convergence (4) | 1703.59702826717 | 1744.70421431762 | -844.798514133585 | 1689.59702826717 | 2617 | 0.004411533 |
| Smad1 | -9.790605044 | 0.450243683 | 0.000924129 | 0.024264207 | 0.814624025 | 1.24535E-05 | 0.5 | 0.000502199 | FALSE | 0 | relative convergence (4) | 2579.24027319923 | 2620.34745924967 | -1282.62013659961 | 2565.24027319923 | 2617 | 0.020506977 |
| Nfix | -8.44064627 | 0.333074743 | 0.32261638 | 0.166090875 | 0.334957932 | 0.157732265 | 0.5 | 0.000474799 | FALSE | 0 | relative convergence (4) | 5608.23905099018 | 5649.34623704063 | -2797.11952549509 | 5594.23905099018 | 2617 | 0.019833976 |
| Ebf3 | -12.53621939 | 1.953423287 | -0.158418832 | -0.143423302 | 7.123555644 | 1.82647E-31 | 0.5 | 1.7659E-05 | FALSE | 1 | singular convergence (7) | NA | NA | NA | NA | 2617 | 0.002121854 |
| Bnip3 | -10.77686421 | 0.691610735 | -0.637099549 | 0.308343611 | 1.069333432 | 3.2769E-14 | 0.5 | 0.000456064 | FALSE | 0 | relative convergence (4) | 1512.75157096418 | 1553.85875701463 | -749.375785482091 | 1498.75157096418 | 2617 | 0.019567969 |
| Tnnt2 | -12.19546297 | 1.468727663 | 0.433460681 | 0.812453564 | 4.486474401 | 0.37244045 | 0.5 | 0.000907752 | FALSE | 0 | relative convergence (4) | NA | NA | NA | NA | 2617 | 0.029468944 |
| Serpine2 | -10.47231125 | 1.262733114 | 0.099876448 | 0.200881358 | 6.251439605 | 8.53768E-10 | 0.5 | 6.58248E-10 | FALSE | 0 | relative convergence (4) | 2208.20059842733 | 2249.30778447778 | -1097.10029921366 | 2194.20059842733 | 2617 | 6.39937E-07 |
| Sphkap | -12.49869385 | 1.875673784 | 1.047426241 | -0.52848706 | 15.09846117 | 0.487153361 | 0.5 | 0.000910321 | FALSE | 0 | relative convergence (4) | 855.104411600076 | 896.211597650524 | -420.552205800038 | 841.104411600076 | 2617 | 0.029468944 |
| Efhd1 | -11.31808933 | 0.986653162 | 0.089013833 | 0.030205808 | 3.590634966 | 1.49124E-18 | 0.5 | 0.00019802 | FALSE | 0 | relative convergence (4) | NA | NA | NA | NA | 2617 | 0.011385075 |
| Arl4c | -11.88641626 | 1.491771893 | 0.866108257 | 0.64877732 | 2.175653114 | 1.50397E-37 | 0.5 | 2.86068E-06 | FALSE | 1 | singular convergence (7) | NA | NA | NA | NA | 2617 | 0.000546287 |
| Dsel | -11.11909444 | 0.752299531 | 0.627850858 | 0.472016065 | 0.508217201 | 1.39127E-08 | 0.5 | 0.000702666 | FALSE | 0 | relative convergence (4) | 1336.91679426584 | 1378.02398031629 | -661.458397132919 | 1322.91679426584 | 2617 | 0.0252158 |
| Neurl1b | -12.8248947 | 1.912808639 | 0.053109822 | -0.212272407 | 7.832418784 | 6.43029E-30 | 0.5 | 0.000120232 | FALSE | 1 | singular convergence (7) | 603.041938576245 | 644.149124626692 | -294.520969288122 | 589.041938576245 | 2617 | 0.007936823 |
| Mmp14 | -10.36187009 | 0.840515107 | -0.276745527 | 0.116108783 | 0.618177843 | 0.168437836 | 0.5 | 7.68058E-05 | FALSE | 0 | relative convergence (4) | 2207.62141958075 | 2248.7286056312 | -1096.81070979038 | 2193.62141958075 | 2617 | 0.006039419 |
| Ephx2 | -8.947538577 | 0.486135067 | 0.192381923 | 0.358227736 | 1.381954285 | 8.93297E-17 | 0.5 | 2.55234E-06 | FALSE | 0 | relative convergence (4) | NA | NA | NA | NA | 2617 | 0.000496267 |
| Rcan2 | -10.02842935 | 0.777074599 | 0.116835053 | 0.104498204 | 5.98137444 | 1.58201E-08 | 0.5 | 2.19001E-05 | FALSE | 0 | relative convergence (4) | 2407.937766469 | 2449.04495251945 | -1196.9688832345 | 2393.937766469 | 2617 | 0.002545645 |
| Slc29a1 | -9.468691869 | 0.445966 | -0.281498634 | 0.143755707 | 0.36717115 | 5.15921E-07 | 0.5 | 2.34302E-05 | FALSE | 0 | relative convergence (4) | 3188.82186505418 | 3229.92905110463 | -1587.41093252709 | 3174.82186505418 | 2617 | 0.002637498 |
| Skiv2l | -11.4764954 | 1.037652431 | 0.044422783 | 1.056980779 | 0.604506901 | 0.189283401 | 0.5 | 0.000822824 | FALSE | 0 | relative convergence (4) | 1224.69712656085 | 1265.8043126113 | -605.348563280426 | 1210.69712656085 | 2617 | 0.027757967 |
| Ncam1 | -8.291473312 | 0.489657008 | 0.092577418 | 0.192189764 | 1.392864714 | 0.223794695 | 0.5 | 5.49771E-05 | FALSE | 0 | relative convergence (4) | 6404.50676884345 | 6445.6139548939 | -3195.25338442173 | 6390.50676884345 | 2617 | 0.004666069 |
| Acat1 | -10.05592469 | 0.453294512 | 0.179318564 | 0.396083628 | 0.489623261 | 3.41184E-17 | 0.5 | 0.001093336 | FALSE | 1 | singular convergence (7) | 2313.30003700433 | 2354.40722305478 | -1149.65001850216 | 2299.30003700433 | 2617 | 0.033694905 |
| Nlgn2 | -12.17906057 | 1.144137332 | 0.878301609 | 0.962455553 | 1.42478107 | 4.07802E-05 | 0.5 | 0.001782779 | FALSE | 0 | relative convergence (4) | 792.563061995277 | 833.670248045725 | -389.281530997639 | 778.563061995277 | 2617 | 0.046816888 |
| Pimreg | -12.33595167 | 1.55601827 | -1.110936618 | -0.561329812 | 5.103047753 | 2.23191E-18 | 0.5 | 9.69693E-05 | FALSE | 0 | relative convergence (4) | 670.775231226057 | 711.882417276505 | -328.387615613029 | 656.775231226057 | 2617 | 0.006959665 |
| Ift81 | -11.67499769 | 0.959615198 | 0.865747632 | 0.929027636 | 2.792952321 | 7.88119E-12 | 0.5 | 0.001478569 | FALSE | 0 | relative convergence (4) | 1039.10449545636 | 1080.2116815068 | -512.552247728178 | 1025.10449545636 | 2617 | 0.041282833 |
| Gtf2ird1 | -10.62502791 | 0.585712955 | 0.122108416 | 0.239232711 | 0.23114015 | 1.29755E-05 | 0.5 | 0.000782361 | FALSE | 0 | relative convergence (4) | 1642.12643529178 | 1683.23362134222 | -814.063217645888 | 1628.12643529178 | 2617 | 0.026987214 |
| Stx1a | -12.18903656 | 1.49222242 | 0.738462905 | 0.769604472 | 0.729604752 | 3.62412E-05 | 0.5 | 2.68706E-05 | FALSE | 0 | relative convergence (4) | 903.985721308196 | 945.092907358643 | -444.992860654098 | 889.985721308196 | 2617 | 0.002872579 |
| Upk3b | -13.34615987 | 2.526917138 | -17.74156706 | -1.119830445 | 60.53029092 | 8.29943E-08 | 0.5 | 0.000720937 | FALSE | 0 | relative convergence (4) | NA | NA | NA | NA | 2617 | 0.025580576 |
| Gm10801 | -9.132420859 | 1.061420715 | 0.178025349 | -2.25250194 | 1.782854889 | 0.698821482 | 0.5 | 0.001079455 | FALSE | 0 | relative convergence (4) | 4848.15848769884 | 4889.26567374929 | -2417.07924384942 | 4834.15848769884 | 2617 | 0.033459969 |
| Gm10800 | -11.93447329 | 1.937935658 | 0.577515797 | -1.877605155 | 3.315233506 | 0.658309088 | 0.5 | 6.55067E-06 | FALSE | 0 | relative convergence (4) | 1231.8109017133 | 1272.91808776374 | -608.905450856648 | 1217.8109017133 | 2617 | 0.001015259 |
| Pamr1 | -12.63307673 | 1.713669477 | 0.519177342 | -0.45244619 | 8.154556575 | 3.76283E-13 | 0.5 | 0.000223602 | FALSE | 0 | relative convergence (4) | 624.465049531038 | 665.572235581486 | -305.232524765519 | 610.465049531038 | 2617 | 0.012373469 |
| Fmn1 | -11.86529926 | 1.018278001 | -0.486012687 | 0.110780772 | 3.639591714 | 2.21142E-05 | 0.5 | 0.00185478 | FALSE | 0 | relative convergence (4) | 765.649983565359 | 806.757169615806 | -375.824991782679 | 751.649983565359 | 2617 | 0.047748837 |
| Thbs1 | -10.65295898 | 1.278483092 | -0.341458811 | -0.001480018 | 6.204819 | 1.16397E-11 | 0.5 | 6.2383E-09 | FALSE | 0 | relative convergence (4) | NA | NA | NA | NA | 2617 | 4.16952E-06 |
| Osbpl3 | -9.966700116 | 0.471140259 | -0.023951754 | 0.08961064 | 1.444671102 | 9.12498E-79 | 0.5 | 0.00124137 | FALSE | 1 | singular convergence (7) | NA | NA | NA | NA | 2617 | 0.036978292 |
| Jazf1 | -10.53834986 | 0.635148786 | -0.139750582 | 0.31270619 | 3.098347242 | 5.70713E-13 | 0.5 | 0.001374688 | FALSE | 0 | relative convergence (4) | 1724.88008910372 | 1765.98727515417 | -855.440044551859 | 1710.88008910372 | 2617 | 0.039317425 |
| Scrn1 | -12.78737542 | 1.695311431 | 1.345835592 | 0.95892074 | 0.878787716 | 6.03725E-06 | 0.5 | 0.000314828 | FALSE | 0 | relative convergence (4) | 657.822866762852 | 698.930052813299 | -321.911433381426 | 643.822866762852 | 2617 | 0.015744333 |
| Htra3 | -11.55693063 | 1.227047509 | -17.97234144 | -0.436889663 | 7.494698422 | 1.71186E-31 | 0.5 | 9.79134E-05 | FALSE | 1 | singular convergence (7) | 965.306957135927 | 1006.41414318637 | -475.653478567963 | 951.306957135927 | 2617 | 0.006980571 |
| 4632427E13Rik | -9.976511004 | 0.536580135 | -0.089660838 | 0.508385064 | 0.636941867 | 8.06804E-85 | 0.5 | 8.23693E-05 | FALSE | 1 | singular convergence (7) | NA | NA | NA | NA | 2617 | 0.00637051 |
| Dtx3 | -10.9971329 | 0.783479632 | 0.635427309 | 0.617235144 | 0.416204008 | 1.51521E-05 | 0.5 | 0.000142715 | FALSE | 0 | relative convergence (4) | 1473.39705906877 | 1514.50424511922 | -729.698529534387 | 1459.39705906877 | 2617 | 0.009164459 |
| Selenbp1 | -11.49636569 | 0.847283897 | 0.668098857 | 0.922253535 | 1.246803372 | 8.50867E-07 | 0.5 | 0.001635959 | FALSE | 0 | relative convergence (4) | 1117.40283158254 | 1158.51001763299 | -551.701415791269 | 1103.40283158254 | 2617 | 0.044076646 |
| Zfp687 | -12.60262139 | 1.362766639 | 1.155932725 | 1.281565538 | 0.483457553 | 1.80234E-05 | 0.5 | 0.001710676 | FALSE | 0 | relative convergence (4) | 645.79242011633 | 686.899606166778 | -315.896210058165 | 631.79242011633 | 2617 | 0.045507393 |
| Ttc3 | -8.665965675 | 0.348088121 | -0.121265352 | 0.234985179 | 0.284024657 | 0.077250408 | 0.5 | 0.001769593 | FALSE | 0 | relative convergence (4) | 4968.21141197256 | 5009.31859802301 | -2477.10570598628 | 4954.21141197256 | 2617 | 0.046725984 |
| Agrn | -9.847193733 | 0.431949148 | -0.297354411 | 0.090305292 | 0.611636493 | 6.40786E-16 | 0.5 | 0.000917632 | FALSE | 1 | singular convergence (7) | NA | NA | NA | NA | 2617 | 0.029468944 |
| Chpf2 | -11.29406078 | 0.80519483 | -0.70094002 | 1.255120526 | 0.643438914 | 2.70722E-05 | 0.5 | 0.000831658 | FALSE | 0 | relative convergence (4) | 1310.05612502923 | 1351.16331107968 | -648.028062514615 | 1296.05612502923 | 2617 | 0.027901639 |
| Cox6b2 | -12.20915259 | 1.412312406 | -0.474245398 | 0.36260307 | 5.302322941 | 2.5012E-30 | 0.5 | 0.000224467 | FALSE | 0 | relative convergence (4) | 759.81587921824 | 800.923065268688 | -372.90793960912 | 745.81587921824 | 2617 | 0.012373469 |
| Mfsd5 | -11.39621364 | 0.775632804 | 0.310940959 | 0.561087392 | 1.44422E-09 | 5.40243E-05 | 0.5 | 0.001636639 | FALSE | 1 | false convergence (8) | 1083.66423748297 | 1124.77142353341 | -534.832118741483 | 1069.66423748297 | 2617 | 0.044076646 |
| Ddx50 | -9.254615417 | 0.313695067 | -0.044816573 | 0.044378721 | 0.127266812 | 3.22865E-05 | 0.5 | 0.000659579 | FALSE | 0 | relative convergence (4) | 3402.37915786355 | 3443.486343914 | -1694.18957893178 | 3388.37915786355 | 2617 | 0.024322553 |
| Col18a1 | -9.36678043 | 0.787073666 | -0.708136038 | -0.016489174 | 2.801898464 | 1.21568E-11 | 0.5 | 2.49394E-09 | FALSE | 0 | relative convergence (4) | 3631.20388706844 | 3672.31107311889 | -1808.60194353422 | 3617.20388706844 | 2617 | 1.77801E-06 |
| Nsmf | -11.11611581 | 0.927337461 | 0.50635952 | 1.064045457 | 0.190015822 | 0.292393309 | 0.5 | 0.000634067 | FALSE | 0 | relative convergence (4) | 1377.13058576869 | 1418.23777181914 | -681.565292884344 | 1363.13058576869 | 2617 | 0.023732387 |
| Fstl3 | -12.47436829 | 1.388924042 | 0.673119453 | 0.528927511 | 3.205993938 | 2.06916E-05 | 0.5 | 0.000878075 | FALSE | 0 | relative convergence (4) | 627.971517344917 | 669.078703395364 | -306.985758672458 | 613.971517344917 | 2617 | 0.029161894 |
| Tmem259 | -10.23378819 | 0.646748868 | -0.121043418 | 0.706775392 | 0.148784766 | 4.8332E-16 | 0.5 | 5.30837E-06 | FALSE | 0 | relative convergence (4) | 2301.82362298188 | 2342.93080903233 | -1143.91181149094 | 2287.82362298188 | 2617 | 0.00087335 |
| Lmnb2 | -11.73612269 | 1.09852786 | -1.6850664 | -1.006582031 | 5.383247057 | 1.11892E-16 | 0.5 | 0.000689619 | FALSE | 1 | singular convergence (7) | NA | NA | NA | NA | 2617 | 0.025018224 |
| Eef2 | -7.659539394 | 0.291952026 | 0.004747979 | 0.330566811 | 0.162516241 | 0.176968395 | 0.5 | 0.001788166 | FALSE | 0 | relative convergence (4) | 7496.9128551167 | 7538.02004116715 | -3741.45642755835 | 7482.9128551167 | 2617 | 0.046816888 |
| Hmg20b | -10.40388013 | 0.623085581 | 0.340595705 | 0.180139422 | 0.367703892 | 1.13178E-25 | 0.5 | 7.83839E-05 | FALSE | 1 | singular convergence (7) | 1891.06448277009 | 1932.17166882054 | -938.532241385047 | 1877.06448277009 | 2617 | 0.00611852 |
| Nfic | -8.909129704 | 0.569861182 | 0.206126155 | 0.317347275 | 0.309153103 | 0.161202784 | 0.5 | 0.000412212 | FALSE | 0 | relative convergence (4) | 4501.34958741392 | 4542.45677346436 | -2243.67479370696 | 4487.34958741392 | 2617 | 0.018354797 |
| Ccn5 | -13.30382835 | 2.208957687 | 0.636454473 | -0.207389405 | 16.84301653 | 7.35274E-18 | 0.5 | 0.000225841 | FALSE | 0 | relative convergence (4) | 493.069205345666 | 534.176391396114 | -239.534602672833 | 479.069205345666 | 2617 | 0.012385355 |
| Gpx7 | -10.83611759 | 1.035147922 | 0.22015967 | 0.43305402 | 1.316644117 | 0.000128862 | 0.5 | 3.84445E-05 | FALSE | 0 | relative convergence (4) | 1792.17007647896 | 1833.27726252941 | -889.08503823948 | 1778.17007647896 | 2617 | 0.003638283 |
| Eri3 | -9.647304346 | 0.376780314 | -0.075009154 | 0.00975396 | 0.429351346 | 9.25941E-44 | 0.5 | 0.001140073 | FALSE | 1 | singular convergence (7) | 2769.11762954677 | 2810.22481559722 | -1377.55881477339 | 2755.11762954677 | 2617 | 0.034636191 |
| P3h1 | -11.25599233 | 0.803744617 | -0.186239546 | 0.343215302 | 0.068645044 | 4.43664E-05 | 0.5 | 0.00047468 | FALSE | 0 | relative convergence (4) | 1158.55716837521 | 1199.66435442565 | -572.278584187603 | 1144.55716837521 | 2617 | 0.019833976 |
| Trp53inp2 | -11.35439379 | 0.828692904 | -0.428508062 | 0.157225999 | 0.334420841 | 2.07166E-17 | 0.5 | 0.000641396 | FALSE | 0 | relative convergence (4) | 1019.12403626354 | 1060.23122231399 | -502.562018131771 | 1005.12403626354 | 2617 | 0.02373389 |
| Soga1 | -10.69779715 | 0.712113205 | 0.435345446 | 0.576604105 | 0.507439788 | 4.78622E-06 | 0.5 | 9.13539E-05 | FALSE | 0 | relative convergence (4) | 1682.69800234717 | 1723.80518839762 | -834.349001173585 | 1668.69800234717 | 2617 | 0.00670765 |
| Jph2 | -11.07179529 | 0.836419788 | -0.820138934 | -0.330969695 | 6.303531324 | 4.13872E-44 | 0.5 | 0.0008047 | FALSE | 1 | singular convergence (7) | NA | NA | NA | NA | 2617 | 0.027405921 |
| 2310022B05Rik | -10.74321606 | 0.794254144 | -0.474841384 | -0.278475738 | 1.402223293 | 1.59137E-15 | 0.5 | 4.56094E-05 | FALSE | 0 | relative convergence (4) | 1509.61660750726 | 1550.7237935577 | -747.808303753628 | 1495.61660750726 | 2617 | 0.004201778 |
| Sprtn | -11.95314209 | 1.061293848 | -0.066109775 | 0.677484538 | 2.53765E-07 | 0.186394271 | 0.5 | 0.001841025 | FALSE | 0 | relative convergence (4) | 859.901551195665 | 901.008737246113 | -422.950775597833 | 845.901551195665 | 2617 | 0.047670506 |
| Plekha4 | -11.81185335 | 1.249106964 | 0.229484955 | 0.683249195 | 1.739997056 | 4.7851E-09 | 0.5 | 5.15196E-05 | FALSE | 0 | relative convergence (4) | 1011.63326299525 | 1052.7404490457 | -498.816631497625 | 997.63326299525 | 2617 | 0.004444998 |
| Dbp | -10.39665247 | 0.947048635 | -0.216014017 | -0.154855911 | 1.391124512 | 0.432778729 | 0.5 | 1.97879E-05 | FALSE | 0 | relative convergence (4) | 1953.27981348821 | 1994.38699953866 | -969.639906744107 | 1939.27981348821 | 2617 | 0.002325406 |
| Pls3 | -8.724716419 | 0.379854189 | -0.02983819 | 0.298081982 | 1.19949161 | 2.91996E-06 | 0.5 | 4.86419E-05 | FALSE | 0 | relative convergence (4) | 4954.25316580439 | 4995.36035185483 | -2470.12658290219 | 4940.25316580439 | 2617 | 0.00437123 |
| Sost | -9.959162205 | 0.662325319 | 0.050662804 | -0.168948387 | 6.222610601 | 1.8101E-06 | 0.5 | 0.000322934 | FALSE | 0 | relative convergence (4) | 2330.51999869718 | 2371.62718474763 | -1158.25999934859 | 2316.51999869718 | 2617 | 0.015988216 |
| Cd300lg | -13.02288466 | 2.052439745 | 1.113480669 | -1.520848054 | 10.93024499 | 0.8509116 | 0.5 | 0.000617494 | FALSE | 0 | relative convergence (4) | 1035.29589839253 | 1076.40308444298 | -510.647949196264 | 1021.29589839253 | 2617 | 0.0234166 |
| Mrc2 | -12.02341956 | 1.27626095 | 0.755824894 | -1.408628846 | 2.895814502 | 3.13267E-05 | 0.5 | 0.000183691 | FALSE | 0 | relative convergence (4) | 777.621588042272 | 818.72877409272 | -381.810794021136 | 763.621588042272 | 2617 | 0.010793387 |
| Eda | -10.11905364 | 0.679681847 | -0.572927265 | 0.397509186 | 4.940987543 | 4.93206E-40 | 0.5 | 0.000361738 | FALSE | 1 | singular convergence (7) | NA | NA | NA | NA | 2617 | 0.017185402 |
| Frk | -12.43504014 | 1.485921503 | -0.940048863 | 0.95957566 | 3.645078343 | 2.97173E-05 | 0.5 | 0.000371994 | FALSE | 0 | relative convergence (4) | 710.272258635342 | 751.37944468579 | -348.136129317671 | 696.272258635342 | 2617 | 0.017447801 |
| Thbs3 | -11.4531208 | 0.986260933 | -1.935606552 | -1.568980436 | 1.891498704 | 3.44733E-05 | 0.5 | 0.000205494 | FALSE | 0 | relative convergence (4) | 929.322345509839 | 970.429531560287 | -457.66117275492 | 915.322345509839 | 2617 | 0.011689136 |
| Pbxip1 | -9.526368572 | 0.545878405 | 0.017995146 | 0.334253147 | 1.191996591 | 2.47035E-06 | 0.5 | 5.23478E-06 | FALSE | 0 | relative convergence (4) | 3302.93299171181 | 3344.04017776226 | -1644.46649585591 | 3288.93299171181 | 2617 | 0.00087335 |
| Git1 | -10.60596445 | 0.735359481 | 0.4179649 | 0.933770256 | 0.570497475 | 0.251973849 | 0.5 | 0.001128442 | FALSE | 0 | relative convergence (4) | 1937.29716944981 | 1978.40435550025 | -961.648584724903 | 1923.29716944981 | 2617 | 0.034478733 |
| Flot2 | -10.04438004 | 0.472435864 | 0.544222001 | 0.475647902 | 0.142047395 | 2.23996E-28 | 0.5 | 0.000347941 | FALSE | 1 | singular convergence (7) | NA | NA | NA | NA | 2617 | 0.016685586 |
| Prkacb | -9.614993625 | 0.43561619 | -0.112357666 | 0.105392878 | 0.199970795 | 1.19378E-09 | 0.5 | 7.2962E-05 | FALSE | 1 | singular convergence (7) | 2944.77320410196 | 2985.88039015241 | -1465.38660205098 | 2930.77320410196 | 2617 | 0.005779673 |
| N4bp3 | -12.46916547 | 1.328588346 | 0.669353753 | 1.131420198 | 2.703702686 | 4.22522E-19 | 0.5 | 0.001446853 | FALSE | 0 | relative convergence (4) | 673.910216280449 | 715.017402330897 | -329.955108140225 | 659.910216280449 | 2617 | 0.040761691 |
| Samd14 | -13.69628106 | 2.335253002 | 1.92305916 | 1.518172436 | 0.894930477 | 4.37664E-07 | 0.5 | 0.001295273 | FALSE | 0 | relative convergence (4) | 505.372457829283 | 546.479643879731 | -245.686228914641 | 491.372457829283 | 2617 | 0.037949721 |
| Naglu | -11.09610888 | 0.728025149 | -0.094527699 | 0.332445122 | 1.55056E-06 | 1.4766E-154 | 0.5 | 0.000625067 | FALSE | 1 | singular convergence (7) | 1269.01355255169 | 1310.12073860214 | -627.506776275845 | 1255.01355255169 | 2617 | 0.02353686 |
| Tmem132a | -11.43831752 | 1.06684567 | 0.223011747 | 0.140773881 | 0.978286419 | 8.01626E-13 | 0.5 | 3.09845E-05 | FALSE | 0 | relative convergence (4) | 1127.42810161363 | 1168.53528766408 | -556.714050806814 | 1113.42810161363 | 2617 | 0.00312593 |
| Prpf19 | -10.29625051 | 0.524076777 | -0.706005684 | 0.324903423 | 0.250142978 | 2.9452E-16 | 0.5 | 0.00044674 | FALSE | 0 | relative convergence (4) | NA | NA | NA | NA | 2617 | 0.019263881 |
| Nbeal1 | -8.936946654 | 0.45698413 | 0.057424659 | 0.307192848 | 0.383865684 | 1.11494E-16 | 0.5 | 9.42803E-08 | FALSE | 0 | relative convergence (4) | 4533.75955909019 | 4574.86674514064 | -2259.8797795451 | 4519.75955909019 | 2617 | 3.36078E-05 |
| Fn1 | -7.895910218 | 1.109926392 | -0.257412426 | -0.313859115 | 1.433675396 | 0.053365471 | 0.5 | 2.94941E-06 | FALSE | 0 | relative convergence (4) | 8577.09834234318 | 8618.20552839363 | -4281.54917117159 | 8563.09834234318 | 2617 | 0.00055335 |
| Igfbp2 | -13.67178607 | 3.217639006 | -16.2365037 | -20.03853772 | 30.95960369 | 1.46199E-20 | 0.5 | 5.02208E-05 | FALSE | 1 | singular convergence (7) | NA | NA | NA | NA | 2617 | 0.004411533 |
| Enah | -9.926247028 | 0.609407947 | 0.021224155 | -0.234333157 | 4.458623822 | 9.35241E-57 | 0.5 | 0.000336191 | FALSE | 1 | singular convergence (7) | 2372.54778370378 | 2413.65496975422 | -1179.27389185189 | 2358.54778370378 | 2617 | 0.016491856 |
| Susd4 | -13.26240311 | 2.026375767 | 0.541399261 | -0.83975404 | 13.71138254 | 3.87894E-05 | 0.5 | 0.001564234 | FALSE | 0 | relative convergence (4) | 463.070599004108 | 504.177785054555 | -224.535299502054 | 449.070599004108 | 2617 | 0.042782392 |
| G0s2 | -11.643224 | 1.48036365 | 1.292953967 | -0.355709281 | 9.491755963 | 1.22994E-12 | 0.5 | 1.23643E-05 | FALSE | 0 | relative convergence (4) | 1150.23644286046 | 1191.34362891091 | -568.11822143023 | 1136.23644286046 | 2617 | 0.001632398 |
| Hacd1 | -11.53697654 | 1.335078153 | 0.344928314 | 0.384251052 | 2.48822963 | 2.89471E-11 | 0.5 | 9.18692E-07 | FALSE | 0 | relative convergence (4) | 1220.63771996514 | 1261.74490601559 | -603.31885998257 | 1206.63771996514 | 2617 | 0.000218322 |
| Tc2n | -12.735702 | 1.754061552 | -18.30279306 | 0.19302073 | 8.22307261 | 1.79019E-14 | 0.5 | 0.000460395 | FALSE | 0 | relative convergence (4) | NA | NA | NA | NA | 2617 | 0.01961538 |
| Fbln5 | -8.574631715 | 0.750712954 | -0.374355923 | 0.329548708 | 0.990054071 | 0.272170264 | 0.5 | 7.15034E-06 | FALSE | 0 | relative convergence (4) | 7124.33258082006 | 7165.43976687051 | -3555.16629041003 | 7110.33258082006 | 2617 | 0.001076982 |
| Trf | -10.76929183 | 0.962584862 | 0.738813211 | 0.15862514 | 3.635655054 | 1.13438E-12 | 0.5 | 8.83422E-06 | FALSE | 0 | relative convergence (4) | NA | NA | NA | NA | 2617 | 0.001259642 |
| Dpysl3 | -9.449855355 | 0.675749636 | 0.088879791 | 0.060757527 | 1.431745854 | 1.76315E-14 | 0.5 | 1.4676E-08 | FALSE | 0 | relative convergence (4) | NA | NA | NA | NA | 2617 | 8.26029E-06 |
| Ppic | -9.86344543 | 0.718187464 | 0.26488963 | -0.005681123 | 0.438441676 | 0.336105142 | 0.5 | 1.40452E-05 | FALSE | 0 | relative convergence (4) | 3242.39707557472 | 3283.50426162517 | -1614.19853778736 | 3228.39707557472 | 2617 | 0.001767053 |
| Prrc1 | -9.989251199 | 0.439871743 | -0.152165788 | 0.051152459 | 0.316097593 | 5.27606E-35 | 0.5 | 0.000883293 | FALSE | 1 | singular convergence (7) | 2254.36411788598 | 2295.47130393643 | -1120.18205894299 | 2240.36411788598 | 2617 | 0.029244378 |
| Pdgfrb | -10.99808798 | 0.952051326 | 0.124850671 | 0.192385469 | 1.904912654 | 2.29379E-09 | 0.5 | 1.16424E-05 | FALSE | 0 | relative convergence (4) | 1453.96676130775 | 1495.0739473582 | -719.983380653874 | 1439.96676130775 | 2617 | 0.001556298 |
| Csf1r | -13.73589258 | 2.437629204 | 2.199949533 | 1.224852986 | 19.54695276 | 2.33738E-05 | 0.5 | 0.001794925 | FALSE | 0 | relative convergence (4) | 477.617466731786 | 518.724652782234 | -231.808733365893 | 463.617466731786 | 2617 | 0.046816888 |
| Ablim3 | -12.01302745 | 1.811884958 | -0.606905845 | -0.050073276 | 8.176240412 | 0.40718285 | 0.5 | 0.000694121 | FALSE | 0 | relative convergence (4) | 1250.42658954225 | 1291.5337755927 | -618.213294771125 | 1236.42658954225 | 2617 | 0.025077451 |
| Fech | -10.47382199 | 0.570543971 | 0.674783404 | 0.081635451 | 0.158755083 | 2.89085E-05 | 0.5 | 0.000409311 | FALSE | 0 | relative convergence (4) | 1843.13506920246 | 1884.2422552529 | -914.567534601228 | 1829.13506920246 | 2617 | 0.018354797 |
| Tubb6 | -11.46738765 | 1.661776971 | -0.290851146 | -0.76093537 | 5.110349807 | 2.76106E-12 | 0.5 | 9.37324E-10 | FALSE | 0 | relative convergence (4) | NA | NA | NA | NA | 2617 | 8.35312E-07 |
| Gm7694 | -11.99132358 | 1.538582647 | -0.689886314 | 1.39481551 | 1.405907299 | 2.55611E-05 | 0.5 | 2.54292E-06 | FALSE | 0 | relative convergence (4) | 1118.17512826032 | 1159.28231431077 | -552.08756413016 | 1104.17512826032 | 2617 | 0.000496267 |
| Sccpdh | -12.01754242 | 1.088954543 | -1.383965594 | -0.155786748 | 0.949366216 | 1.03995E-05 | 0.5 | 0.001052883 | FALSE | 0 | relative convergence (4) | 691.131191795547 | 732.238377845995 | -338.565595897773 | 677.131191795547 | 2617 | 0.032848934 |
| Actn1 | -8.40397703 | 0.461724975 | 0.089411509 | -0.050656508 | 0.827892289 | 0.063747656 | 0.5 | 0.000509969 | FALSE | 0 | relative convergence (4) | 5881.21759736322 | 5922.32478341367 | -2933.60879868161 | 5867.21759736322 | 2617 | 0.020736137 |
| Abcc9 | -11.34524562 | 1.091620202 | -0.468331129 | -0.281765942 | 10.32977438 | 3.83464E-77 | 0.5 | 0.000247452 | FALSE | 1 | singular convergence (7) | 1059.3667904212 | 1100.47397647165 | -522.683395210601 | 1045.3667904212 | 2617 | 0.013432772 |
| Stk38l | -10.92265181 | 0.708499157 | 0.455417679 | 0.451068442 | 1.711849856 | 3.95705E-33 | 0.5 | 0.001053599 | FALSE | 1 | singular convergence (7) | 1442.98374097232 | 1484.09092702277 | -714.491870486162 | 1428.98374097232 | 2617 | 0.032848934 |
| Papss2 | -11.56225133 | 1.197703812 | 1.102553391 | 0.638594337 | 2.948186952 | 4.637E-05 | 0.5 | 2.2878E-05 | FALSE | 0 | relative convergence (4) | 1211.62334546424 | 1252.73053151468 | -598.811672732118 | 1197.62334546424 | 2617 | 0.002602733 |
| Arhgdia | -8.47305362 | 0.297560592 | 0.060333733 | 0.106490792 | 0.063047801 | 0.137941361 | 0.5 | 0.000470885 | FALSE | 0 | relative convergence (4) | NA | NA | NA | NA | 2617 | 0.019833976 |
| Lrrc45 | -12.08058862 | 1.309305004 | 0.474686363 | 0.679202757 | 0.712594559 | 4.29092E-06 | 0.5 | 0.000121121 | FALSE | 0 | relative convergence (4) | 886.758475923073 | 927.865661973521 | -436.379237961536 | 872.758475923073 | 2617 | 0.007946425 |
| Nid2 | -12.4330455 | 1.932247517 | -0.229375246 | -0.682856518 | 3.429362331 | 6.93739E-15 | 0.5 | 2.42896E-06 | FALSE | 0 | relative convergence (4) | 810.982669667847 | 852.089855718295 | -398.491334833924 | 796.982669667847 | 2617 | 0.0004901 |
| Mdk | -11.98649228 | 1.176867965 | -0.313769563 | -0.740388345 | 2.349761236 | 4.14109E-05 | 0.5 | 0.000502415 | FALSE | 0 | relative convergence (4) | 740.849201547675 | 781.956387598122 | -363.424600773837 | 726.849201547675 | 2617 | 0.020506977 |
| Dsp | -11.68216539 | 1.366166334 | 0.598245014 | -0.047653213 | 2.991304701 | 5.27302E-05 | 0.5 | 0.000898868 | FALSE | 0 | relative convergence (4) | 1063.9292716364 | 1105.03645768685 | -524.9646358182 | 1049.9292716364 | 2617 | 0.029468944 |
| Sema4d | -9.665915631 | 0.427375593 | -0.041552447 | 0.019885719 | 1.04479565 | 4.76154E-07 | 0.5 | 0.000605922 | FALSE | 0 | relative convergence (4) | 2779.04535906615 | 2820.15254511659 | -1382.52267953307 | 2765.04535906615 | 2617 | 0.023059524 |
| Epdr1 | -12.45229431 | 1.539438746 | 0.452794293 | 0.805825265 | 0.820703193 | 3.36406E-05 | 0.5 | 0.000125292 | FALSE | 0 | relative convergence (4) | 730.229461617216 | 771.336647667664 | -358.114730808608 | 716.229461617216 | 2617 | 0.008169952 |
| Mylk4 | -11.2518237 | 1.060790242 | 0.440260685 | -0.960564623 | 12.23645996 | 4.31449E-14 | 0.5 | 0.000311832 | FALSE | 0 | relative convergence (4) | 1098.54785205253 | 1139.65503810298 | -542.273926026267 | 1084.54785205253 | 2617 | 0.015744333 |
| Ank2 | -10.9179434 | 0.798271117 | 0.408140276 | -0.715163101 | 6.312935823 | 3.33357E-85 | 0.5 | 0.001303675 | FALSE | 1 | singular convergence (7) | 1311.64922532658 | 1352.75641137703 | -648.824612663292 | 1297.64922532658 | 2617 | 0.037987732 |
| Papss1 | -10.67834451 | 0.564858855 | 0.467453041 | 0.099956021 | 0.263253839 | 2.11401E-13 | 0.5 | 0.001619789 | FALSE | 0 | relative convergence (4) | NA | NA | NA | NA | 2617 | 0.043853235 |
| Khdrbs3 | -11.10408063 | 0.770250652 | 0.043290863 | 0.241991452 | 1.278379843 | 6.71889E-06 | 0.5 | 0.000568943 | FALSE | 0 | relative convergence (4) | 1237.01485308434 | 1278.12203913479 | -611.50742654217 | 1223.01485308434 | 2617 | 0.022368649 |
| H1f0 | -9.614772558 | 0.593144963 | -0.175720615 | 0.286003584 | 1.179698609 | 5.27689E-26 | 0.5 | 1.90614E-06 | FALSE | 1 | singular convergence (7) | 3169.89916646707 | 3211.00635251752 | -1577.94958323354 | 3155.89916646707 | 2617 | 0.000407685 |
| Cpeb4 | -9.830118683 | 0.420450725 | 0.082043524 | 0.125196546 | 0.221481147 | 1.83875E-07 | 0.5 | 0.000542384 | FALSE | 0 | relative convergence (4) | 2572.05597125457 | 2613.16315730502 | -1279.02798562729 | 2558.05597125457 | 2617 | 0.021642733 |
| Sh3pxd2b | -11.29085074 | 0.967234609 | 0.229792267 | 0.032393494 | 1.571051984 | 7.48325E-06 | 0.5 | 6.49877E-05 | FALSE | 0 | relative convergence (4) | 1164.19219991229 | 1205.29938596274 | -575.096099956146 | 1150.19219991229 | 2617 | 0.00526499 |
| Slco2b1 | -12.31826448 | 1.324528792 | 0.281884927 | 0.76959278 | 5.762739062 | 7.29183E-05 | 0.5 | 0.001154846 | FALSE | 0 | relative convergence (4) | 716.896083542436 | 758.003269592884 | -351.448041771218 | 702.896083542436 | 2617 | 0.034886796 |
| Irag1 | -11.17030184 | 0.889670239 | -0.295153555 | 0.096537124 | 6.54046297 | 3.62931E-06 | 0.5 | 0.000820452 | FALSE | 0 | relative convergence (4) | 1176.37610635012 | 1217.48329240057 | -581.188053175061 | 1162.37610635012 | 2617 | 0.027757967 |
| Uhrf1bp1l | -9.198189327 | 0.33865528 | 0.100033652 | 0.255153045 | 0.630881726 | 1.98015E-12 | 0.5 | 0.000779106 | FALSE | 0 | relative convergence (4) | 3771.55822485673 | 3812.66541090718 | -1878.77911242836 | 3757.55822485673 | 2617 | 0.026963606 |
| Tbc1d30 | -11.12918064 | 1.044969333 | 0.56764794 | 0.79756805 | 2.293826339 | 5.54353E-05 | 0.5 | 8.00403E-06 | FALSE | 0 | relative convergence (4) | 1527.50699789008 | 1568.61418394053 | -756.753498945041 | 1513.50699789008 | 2617 | 0.001188821 |
| Vcan | -10.79223904 | 1.103727275 | -0.258505097 | -0.671696995 | 3.834830122 | 4.44293E-06 | 0.5 | 2.15004E-07 | FALSE | 0 | relative convergence (4) | 1635.60650956671 | 1676.71369561716 | -810.803254783355 | 1621.60650956671 | 2617 | 6.5693E-05 |
| Wdr41 | -11.35971629 | 0.786933111 | 0.347962899 | 0.748030341 | 0.802354699 | 1.56502E-39 | 0.5 | 0.001551781 | FALSE | 1 | singular convergence (7) | NA | NA | NA | NA | 2617 | 0.042550623 |
| Arhgef28 | -9.563283439 | 0.659455457 | -0.224378019 | -0.021203788 | 0.8895247 | 3.97863E-12 | 0.5 | 2.71991E-08 | FALSE | 1 | singular convergence (7) | NA | NA | NA | NA | 2617 | 1.38508E-05 |

| | | | | | | | | | | | | | | | | |
|---|---|---|---|---|---|---|---|---|---|---|---|---|---|---|---|---|
| Twsg1 | -10.01349475 | 0.558570832 | -0.100675198 | -0.295837069 | 0.66225085 | 4.63718E-30 | 0.5 | 4.59704E-05 | FALSE | 1 | singular convergence (7) | NA | NA | NA | NA | 2617 | 0.004201778 |
| Hnrnpll | -10.04829471 | 0.453785957 | -0.25923554 | 0.268181366 | 0.455847504 | 8.45814E-21 | 0.5 | 0.001042492 | FALSE | 1 | singular convergence (7) | 2299.49330457397 | 2340.60049062442 | -1142.74665228698 | 2285.49330457397 | 2617 | 0.032754541 |
| Rhoq | -9.676235348 | 0.377098311 | -0.336090782 | -0.252224963 | 0.302255638 | 4.06457E-26 | 0.5 | 0.001021805 | FALSE | 1 | singular convergence (7) | 2610.24716846742 | 2651.35435451787 | -1298.12358423371 | 2596.24716846742 | 2617 | 0.032321335 |
| Ip6k1 | -10.51409951 | 0.659957539 | 0.346020518 | 0.664831142 | 0.182264341 | 8.17909E-07 | 0.5 | 5.39779E-05 | FALSE | 0 | relative convergence (4) | 2006.70664406511 | 2047.81383011555 | -996.353322032553 | 1992.70664406511 | 2617 | 0.004617918 |
| Mir6236 | -7.157929982 | 1.290235991 | 3.250994329 | 0.467928382 | 0.756096232 | 1.18568547 | 0.5 | 0.000493556 | FALSE | 0 | relative convergence (4) | 10808.8098285023 | 10849.9170145528 | -5397.40491425116 | 10794.8098285023 | 2617 | 0.020378703 |
| Susd5 | -11.84429532 | 1.13020337 | -0.259295621 | -0.986437418 | 9.953632116 | 7.16153E-10 | 0.5 | 0.001249073 | FALSE | 0 | relative convergence (4) | 750.840617967509 | 791.947804017956 | -368.420308983754 | 736.840617967509 | 2617 | 0.037014976 |
| Fat4 | -10.44186456 | 0.651169928 | -0.045683539 | -0.004594739 | 1.595830453 | 2.13131E-40 | 0.5 | 0.000178288 | FALSE | 1 | singular convergence (7) | 1817.93978354794 | 1859.04696959839 | -901.969891773972 | 1803.93978354794 | 2617 | 0.010700445 |
| Zcchc24 | -10.3482214 | 0.515396002 | 0.184429626 | 0.350891497 | 0.701513715 | 1.25332E-24 | 0.5 | 0.001136551 | FALSE | 1 | singular convergence (7) | 1930.65391385753 | 1971.76109990797 | -958.326956928763 | 1916.65391385753 | 2617 | 0.034627578 |
| Pard6g | -13.71794866 | 2.366127414 | 2.183014121 | -0.387928392 | 6.98430734 | 3.40593E-05 | 0.5 | 0.001488485 | FALSE | 0 | relative convergence (4) | 463.078106833584 | 504.185292884032 | -224.539053416792 | 449.078106833584 | 2617 | 0.041282833 |
| Sucla2 | -10.37574202 | 0.509842017 | -0.021253647 | 0.167143852 | 0.405567347 | 2.5844E-21 | 0.5 | 0.001301796 | FALSE | 0 | relative convergence (4) | 1837.44614025105 | 1878.5533263015 | -911.723070125525 | 1823.44614025105 | 2617 | 0.037987732 |
| Dgkh | -9.849465255 | 0.538551749 | 0.218090078 | 0.120266925 | 1.568314529 | 1.90364E-54 | 0.5 | 0.000161251 | FALSE | 1 | singular convergence (7) | 2667.1886339954 | 2708.29582004585 | -1326.5943169977 | 2653.1886339954 | 2617 | 0.009962568 |
| Dab2 | -9.566559827 | 0.474714207 | 0.303858911 | 0.106006791 | 1.445610984 | 1.87961E-12 | 0.5 | 0.000212397 | FALSE | 0 | relative convergence (4) | 3146.27037174749 | 3187.37755779794 | -1566.13518587375 | 3132.27037174749 | 2617 | 0.012017835 |
| Vgll3 | -11.98113402 | 1.165006429 | 0.614018799 | 0.593863215 | 5.221526917 | 3.56859E-05 | 0.5 | 0.000784835 | FALSE | 0 | relative convergence (4) | 833.850153611404 | 874.957339661852 | -409.925076805702 | 819.850153611404 | 2617 | 0.026987214 |
| Olfml2a | -10.45389683 | 0.903000236 | -0.636960911 | 0.537945936 | 0.998135026 | 3.23607E-05 | 0.5 | 7.24738E-08 | FALSE | 0 | relative convergence (4) | 2146.16539373819 | 2187.27257978864 | -1066.0826968691 | 2132.16539373819 | 2617 | 3.10014E-05 |
| Grb14 | -12.49323125 | 1.647361196 | 0.411009043 | 0.534692592 | 5.814135321 | 2.77315E-10 | 0.5 | 0.000117072 | FALSE | 0 | relative convergence (4) | 709.727123543802 | 750.834309594249 | -347.863561771901 | 695.727123543802 | 2617 | 0.007824811 |
| 2610203C22Rik | -13.69505529 | 2.406968855 | -16.86952302 | 0.001811996 | 12.69096394 | 1.29062E-20 | 0.5 | 0.001506265 | FALSE | 1 | singular convergence (7) | 414.586206720047 | 455.693392770495 | -200.293103360024 | 400.586206720047 | 2617 | 0.04162273 |
| Uaca | -9.487930142 | 0.670718479 | 0.629488547 | 0.42357658 | 0.565483775 | 4.44083E-05 | 0.5 | 6.4021E-10 | FALSE | 0 | relative convergence (4) | 3692.1970027119 | 3733.30418876235 | -1839.09850135595 | 3678.1970027119 | 2617 | 6.39937E-07 |
| Rasl12 | -11.37171653 | 0.89262648 | -0.165626433 | -0.025926841 | 6.89607429 | 1.74535E-05 | 0.5 | 0.001531962 | FALSE | 0 | relative convergence (4) | 1010.79779402685 | 1051.9049800773 | -498.398897013425 | 996.797794026851 | 2617 | 0.042115158 |
| Mtus2 | -10.90710778 | 0.740774884 | -0.067248157 | -0.312765296 | 5.74053742 | 1.09287E-15 | 0.5 | 0.00185187 | FALSE | 0 | relative convergence (4) | 1281.86276030076 | 1322.96994635121 | -633.931380150382 | 1267.86276030076 | 2617 | 0.047748837 |
| Gm44250 | -11.29399566 | 1.103314524 | 0.355786234 | 0.93663576 | 0.475267652 | 0.000117279 | 0.5 | 0.001044445 | FALSE | 0 | relative convergence (4) | 1226.10325590549 | 1267.21044195594 | -606.051627952745 | 1212.10325590549 | 2617 | 0.032754541 |
| Impdh1 | -11.10095731 | 0.689850379 | 0.587803382 | 0.325292972 | 0.550111392 | 8.03325E-16 | 0.5 | 0.00172693 | FALSE | 0 | relative convergence (4) | NA | NA | NA | NA | 2617 | 0.045712338 |
| Bpgm | -10.24074861 | 0.745938537 | 0.328369811 | 0.270462613 | 1.514433248 | 2.16565E-12 | 0.5 | 3.2326E-06 | FALSE | 0 | relative convergence (4) | NA | NA | NA | NA | 2617 | 0.000596025 |
| Ammecr1l | -10.8651006 | 0.806635087 | 0.35362842 | 0.856945556 | 0.472255919 | 1.78744E-05 | 0.5 | 3.33032E-05 | FALSE | 0 | relative convergence (4) | 1644.65865851474 | 1685.76584456519 | -815.32932925737 | 1630.65865851474 | 2617 | 0.003328452 |
| Camk4 | -12.03799717 | 1.739204848 | 0.355437298 | 0.346434387 | 5.74434084 | 0.000180864 | 0.5 | 0.000583298 | FALSE | 0 | relative convergence (4) | 1237.94218007037 | 1279.04936612082 | -611.971090035186 | 1223.94218007037 | 2617 | 0.02251911 |
| Osr1 | -11.14021016 | 1.031069591 | -0.699581203 | 0.084498031 | 3.889188451 | 3.18021E-07 | 0.5 | 3.77607E-05 | FALSE | 0 | relative convergence (4) | 1295.89721867905 | 1337.0044047295 | -640.948609339525 | 1281.89721867905 | 2617 | 0.003605472 |
| Sox11 | -12.7848336 | 1.693380645 | 0.112962368 | -1.338223452 | 5.727025115 | 1.25626E-05 | 0.5 | 0.000594118 | FALSE | 0 | relative convergence (4) | 545.304125474621 | 586.411311525069 | -265.65206273731 | 531.304125474621 | 2617 | 0.022854306 |
| Sntg2 | -9.961355373 | 0.58181811 | 0.449525777 | 0.043043368 | 2.287857212 | 8.22864E-50 | 0.5 | 0.000162099 | FALSE | 1 | singular convergence (7) | NA | NA | NA | NA | 2617 | 0.009962568 |
| Heatr5a | -9.789324584 | 0.390117159 | -0.110506656 | 0.448723815 | 0.537131311 | 4.29359E-24 | 0.5 | 0.001894088 | FALSE | 1 | singular convergence (7) | 2684.60335114686 | 2725.7105371973 | -1335.30167557343 | 2670.60335114686 | 2617 | 0.04857405 |
| Sec23a | -9.99283346 | 0.472481982 | -0.2935078 | 0.158313556 | 0.229321886 | 1.65893E-09 | 0.5 | 0.000274273 | FALSE | 0 | relative convergence (4) | 2305.80390493631 | 2346.91109098676 | -1145.90195246816 | 2291.80390493631 | 2617 | 0.014307704 |

### Table S10. Gene Enrichment Results: Hypoxia versus Normoxia

| GO.biological.process.complete | GOsimple | negLog10fdr | cellType | index | GOsimpleApp |
|---|---|---|---|---|---|
| regulation of MAP kinase activity (GO:0043405) | GO:0043405 | 4.946921557 | SMC | 1 | SMC-GO:0043405 |
| cellular response to vascular endothelial growth factor stimulus (GO:0035924) | GO:0035924 | 4.749579998 | SMC | 2 | SMC-GO:0035924 |
| response to hypoxia (GO:0001666) | GO:0001666 | 4.372634143 | SMC | 3 | SMC-GO:0001666 |
| positive regulation of blood vessel endothelial cell migration (GO:0043536) | GO:0043536 | 4.290730039 | SMC | 4 | SMC-GO:0043536 |
| fibroblast migration (GO:0010761) | GO:0010761 | 2.872895202 | SMC | 5 | SMC-GO:0010761 |
| regulation of blood coagulation (GO:0030193) | GO:0030193 | 2.372634143 | SMC | 6 | SMC-GO:0030193 |
| cellular response to mechanical stimulus (GO:0071260) | GO:0071260 | 2.270025714 | SMC | 7 | SMC-GO:0071260 |
| cellular response to stress (GO:0033554) | GO:0033554 | 2.211124884 | SMC | 8 | SMC-GO:0033554 |
| fibroblast growth factor receptor signaling pathway (GO:0008543) | GO:0008543 | 2 | SMC | 9 | SMC-GO:0008543 |
| positive regulation of chemokine (C-X-C motif) ligand 2 production (GO:2000343) | GO:2000343 | 1.970616222 | SMC | 10 | SMC-GO:2000343 |
| canonical glycolysis (GO:0061621) | GO:0061621 | 1.879426069 | SMC | 11 | SMC-GO:0061621 |
| muscle cell differentiation (GO:0042692) | GO:0042692 | 1.876148359 | SMC | 12 | SMC-GO:0042692 |
| regulation of smooth muscle cell migration (GO:0014910) | GO:0014910 | 1.844663963 | SMC | 13 | SMC-GO:0014910 |
| bone mineralization (GO:0030282) | GO:0030282 | 1.744727495 | SMC | 14 | SMC-GO:0030282 |
| regulation of vascular associated smooth muscle cell dedifferentiation (GO:1905174) | GO:1905174 | 1.716698771 | SMC | 15 | SMC-GO:1905174 |
| bone growth (GO:0098868) | GO:0098868 | 1.581698709 | SMC | 16 | SMC-GO:0098868 |
| regulation of blood vessel remodeling (GO:0060312) | GO:0060312 | 1.469800302 | SMC | 17 | SMC-GO:0060312 |
| regulation of phospholipid metabolic process (GO:1903725) | GO:1903725 | 1.460923901 | SMC | 18 | SMC-GO:1903725 |
| collagen fibril organization (GO:0030199) | GO:0030199 | 1.449771647 | SMC | 19 | SMC-GO:0030199 |
| sensory perception of mechanical stimulus (GO:0050954) | GO:0050954 | 1.41453927 | SMC | 20 | SMC-GO:0050954 |
| regulation of ERK1 and ERK2 cascade (GO:0070372) | GO:0070372 | 1.387216143 | SMC | 21 | SMC-GO:0070372 |
| regulation of platelet-derived growth factor receptor signaling pathway (GO:0010640) | GO:0010640 | 1.367542708 | SMC | 22 | SMC-GO:0010640 |
| regulation of leukocyte chemotaxis (GO:0002688) | GO:0002688 | 1.341988603 | SMC | 23 | SMC-GO:0002688 |
| BMP signaling pathway (GO:0030509) | GO:0030509 | 1.317854924 | SMC | 24 | SMC-GO:0030509 |
| regulation of cell communication (GO:0010646) | GO:0010646 | 6.313363731 | Fib | 25 | Fib-GO:0010646 |
| regulation of signaling (GO:0023051) | GO:0023051 | 6.220403509 | Fib | 26 | Fib-GO:0023051 |
| protein hydroxylation (GO:0018126) | GO:0018126 | 2.896196279 | Fib | 27 | Fib-GO:0018126 |
| extracellular matrix organization (GO:0030198) | GO:0030198 | 2.521433504 | Fib | 28 | Fib-GO:0030198 |
| blood vessel morphogenesis (GO:0048514) | GO:0048514 | 2.434152181 | Fib | 29 | Fib-GO:0048514 |
| cell adhesion (GO:0007155) | GO:0007155 | 2.057991947 | Fib | 30 | Fib-GO:0007155 |
| regulation of cell population proliferation (GO:0042127) | GO:0042127 | 1.790484985 | Fib | 31 | Fib-GO:0042127 |
| angiogenesis (GO:0001525) | GO:0001525 | 1.774690718 | Fib | 32 | Fib-GO:0001525 |
| glycolytic process (GO:0006096) | GO:0006096 | 1.684029655 | Fib | 33 | Fib-GO:0006096 |
| collagen fibril organization (GO:0030199) | GO:0030199 | 1.482804102 | Fib | 34 | Fib-GO:0030199 |
| heparan sulfate proteoglycan metabolic process (GO:0030201) | GO:0030201 | 1.460923901 | Fib | 35 | Fib-GO:0030201 |
| regulation of ossification (GO:0030278) | GO:0030278 | 1.355561411 | Fib | 36 | Fib-GO:0030278 |
| bone development (GO:0060348) | GO:0060348 | 4.457174573 | Mac | 37 | Mac-GO:0060348 |
| ossification (GO:0001503) | GO:0001503 | 2.756961951 | Mac | 38 | Mac-GO:0001503 |
| tissue development (GO:0009888) | GO:0009888 | 1.30980392 | Mac | 39 | Mac-GO:0009888 |

**Table S11.** NICHES intercellular communication list of genes.

cluster 4 = NOX-HOX signaling archetype

cluster 6 = HOX-NOX recovery signaling archetype

cluster 7 = HOX-NOX with exercise recovery signaling archetype

| gene-gene pair | p_val | avg_log2FC | pct.1 | pct.2 | p_val_adj | cluster | gene-gene pair |
|---|---|---|---|---|---|---|---|
| Sema3f Nrp1 | 3.48E-204 | 5.423050014 | 0.949 | 0.043 | 5.88E-201 | 4 | Sema3f Nrp1 |
| Vwf Sirpa | 7.55E-183 | 3.354128179 | 0.967 | 0.144 | 1.27E-179 | 4 | Vwf Sirpa |
| Sema3f Plxna1 | 1.03E-181 | 5.011472035 | 0.889 | 0.04 | 1.74E-178 | 4 | Sema3f Plxna1 |
| Vegfc Lyve1 | 1.47E-175 | 5.463996591 | 0.862 | 0.035 | 2.48E-172 | 4 | Vegfc Lyve1 |
| Sema6d Trem2 | 3.11E-173 | 4.609992559 | 0.874 | 0.053 | 5.25E-170 | 4 | Sema6d Trem2 |
| Vegfa Nrp1 | 1.09E-168 | 3.982160029 | 0.906 | 0.203 | 1.84E-165 | 4 | Vegfa Nrp1 |
| Edn1 Ednrb | 4.59E-168 | 6.125643279 | 0.825 | 0.018 | 7.75E-165 | 4 | Edn1 Ednrb |
| Ntn1 Unc5a | 5.40E-166 | 5.605979406 | 0.829 | 0.036 | 9.12E-163 | 4 | Ntn1 Unc5a |
| F8 Asgr2 | 2.98E-165 | 4.139131813 | 0.873 | 0.078 | 5.03E-162 | 4 | F8 Asgr2 |
| Kitl Epor | 1.79E-164 | 4.496629645 | 0.852 | 0.031 | 3.02E-161 | 4 | Kitl Epor |
| Vegfa Sirpa | 1.32E-157 | 3.163829372 | 0.916 | 0.21 | 2.23E-154 | 4 | Vegfa Sirpa |
| Sema3f Nrp2 | 6.66E-157 | 3.020070993 | 0.933 | 0.127 | 1.12E-153 | 4 | Sema3f Nrp2 |
| Bgn Tlr4 | 1.69E-156 | 1.998944246 | 0.973 | 0.568 | 2.86E-153 | 4 | Bgn Tlr4 |
| Fgf12 Fgfr1 | 3.05E-155 | 4.884614101 | 0.813 | 0.053 | 5.15E-152 | 4 | Fgf12 Fgfr1 |
| Col1a2 Cd36 | 1.46E-148 | 3.015519807 | 0.9 | 0.144 | 2.46E-145 | 4 | Col1a2 Cd36 |
| Jag1 Notch1 | 2.53E-147 | 2.823183893 | 0.93 | 0.198 | 4.26E-144 | 4 | Jag1 Notch1 |
| Dll1 Notch1 | 2.21E-146 | 3.058500142 | 0.895 | 0.096 | 3.74E-143 | 4 | Dll1 Notch1 |
| Col4a4 Cd93 | 6.62E-143 | 5.025075232 | 0.753 | 0.025 | 1.12E-139 | 4 | Col4a4 Cd93 |
| Bgn Ly96 | 1.49E-139 | 1.891224582 | 0.975 | 0.51 | 2.52E-136 | 4 | Bgn Ly96 |
| Gnai2 Ednrb | 2.56E-139 | 2.49935807 | 0.946 | 0.122 | 4.31E-136 | 4 | Gnai2 Ednrb |
| Vegfc Nrp2 | 2.04E-138 | 2.9990653 | 0.87 | 0.139 | 3.44E-135 | 4 | Vegfc Nrp2 |
| Vegfa Nrp2 | 2.74E-137 | 2.791364864 | 0.891 | 0.18 | 4.63E-134 | 4 | Vegfa Nrp2 |
| Dll4 Notch1 | 4.90E-134 | 2.936575254 | 0.874 | 0.106 | 8.28E-131 | 4 | Dll4 Notch1 |
| Sema6d Tyrobp | 2.17E-133 | 2.556917856 | 0.904 | 0.19 | 3.67E-130 | 4 | Sema6d Tyrobp |
| Col1a2 Cd93 | 3.35E-132 | 3.376351636 | 0.802 | 0.092 | 5.66E-129 | 4 | Col1a2 Cd93 |
| Ntn4 Unc5a | 4.23E-132 | 4.519620057 | 0.745 | 0.059 | 7.13E-129 | 4 | Ntn4 Unc5a |
| Gnai2 P2ry12 | 2.79E-130 | 2.341114435 | 0.931 | 0.137 | 4.71E-127 | 4 | Gnai2 P2ry12 |
| Mfng Notch1 | 6.29E-130 | 2.556119522 | 0.903 | 0.149 | 1.06E-126 | 4 | Mfng Notch1 |
| Bmp4 Acvr2b | 2.85E-129 | 3.352885055 | 0.817 | 0.231 | 4.81E-126 | 4 | Bmp4 Acvr2b |
| Rps19 C5ar1 | 4.94E-129 | 2.641110513 | 0.853 | 0.111 | 8.34E-126 | 4 | Rps19 C5ar1 |
| Bmp6 Acvr2b | 9.50E-126 | 3.16916112 | 0.823 | 0.251 | 1.60E-122 | 4 | Bmp6 Acvr2b |
| Igfbp4 Lrp6 | 3.90E-125 | 1.89424119 | 0.922 | 0.487 | 6.58E-122 | 4 | Igfbp4 Lrp6 |
| Gnai2 C5ar1 | 1.03E-121 | 2.18370622 | 0.925 | 0.153 | 1.74E-118 | 4 | Gnai2 C5ar1 |
| Edn1 Kel | 6.03E-120 | 4.304548007 | 0.691 | 0.035 | 1.02E-116 | 4 | Edn1 Kel |
| Calr Scarf1 | 6.49E-117 | 3.144717649 | 0.762 | 0.102 | 1.09E-113 | 4 | Calr Scarf1 |
| Sema6d Plxna1 | 1.65E-116 | 2.483430492 | 0.844 | 0.312 | 2.79E-113 | 4 | Sema6d Plxna1 |
| Sema3g Nrp2 | 1.75E-116 | 2.794665289 | 0.802 | 0.124 | 2.95E-113 | 4 | Sema3g Nrp2 |
| Adam17 Notch1 | 1.50E-114 | 1.893266487 | 0.946 | 0.287 | 2.54E-111 | 4 | Adam17 Notch1 |
| Bmp4 Bmpr2 | 5.08E-113 | 2.714519649 | 0.786 | 0.226 | 8.58E-110 | 4 | Bmp4 Bmpr2 |

| | | | | | | | |
|---|---|---|---|---|---|---|---|
| Col5a1 Sdc3 | 6.62E-112 | 3.115596019 | 0.748 | 0.106 | 1.12E-108 | 4 | Col5a1 Sdc3 |
| Sema4c Plxnb2 | 1.15E-111 | 2.450846704 | 0.844 | 0.365 | 1.95E-108 | 4 | Sema4c Plxnb2 |
| Fn1 C5ar1 | 2.44E-111 | 3.141639737 | 0.738 | 0.081 | 4.12E-108 | 4 | Fn1 C5ar1 |
| Vegfa Egfr | 4.48E-110 | 3.17165578 | 0.765 | 0.233 | 7.56E-107 | 4 | Vegfa Egfr |
| App Tnfrsf21 | 5.62E-110 | 1.990590679 | 0.903 | 0.2 | 9.49E-107 | 4 | App Tnfrsf21 |
| Bmp6 Bmpr2 | 1.30E-109 | 2.659700843 | 0.786 | 0.233 | 2.19E-106 | 4 | Bmp6 Bmpr2 |
| Mfap5 Notch1 | 1.55E-107 | 3.073309471 | 0.733 | 0.106 | 2.61E-104 | 4 | Mfap5 Notch1 |
| Hsp90b1 Tlr4 | 1.26E-106 | 1.306608576 | 0.963 | 0.599 | 2.12E-103 | 4 | Hsp90b1 Tlr4 |
| Col4a3 Cd93 | 1.57E-106 | 5.086105719 | 0.609 | 0.018 | 2.64E-103 | 4 | Col4a3 Cd93 |
| Col4a1 Cd93 | 6.07E-106 | 3.510964446 | 0.684 | 0.068 | 1.02E-102 | 4 | Col4a1 Cd93 |
| Calm1 Abca1 | 1.45E-102 | 1.302256224 | 0.984 | 0.653 | 2.46E-99 | 4 | Calm1 Abca1 |
| Col1a1 Cd36 | 6.35E-100 | 2.878511953 | 0.715 | 0.102 | 1.07E-96 | 4 | Col1a1 Cd36 |
| Lin7c Abca1 | 5.80E-98 | 1.245540093 | 0.994 | 0.698 | 9.79E-95 | 4 | Lin7c Abca1 |
| Cp Slc40a1 | 1.29E-96 | 4.874492983 | 0.58 | 0.026 | 2.17E-93 | 4 | Cp Slc40a1 |
| Col1a1 Cd93 | 3.67E-94 | 3.147267687 | 0.655 | 0.076 | 6.19E-91 | 4 | Col1a1 Cd93 |
| Plat Lrp1 | 9.18E-94 | 2.210698577 | 0.766 | 0.226 | 1.55E-90 | 4 | Plat Lrp1 |
| B2m Hfe | 2.21E-93 | 1.365464823 | 0.942 | 0.249 | 3.73E-90 | 4 | B2m Hfe |
| Calm2 Abca1 | 9.03E-89 | 1.114272026 | 0.982 | 0.696 | 1.52E-85 | 4 | Calm2 Abca1 |
| Bgn Tlr2 | 8.79E-88 | 2.581928336 | 0.684 | 0.132 | 1.48E-84 | 4 | Bgn Tlr2 |
| Vegfc Itga9 | 1.28E-85 | 3.092583568 | 0.609 | 0.079 | 2.16E-82 | 4 | Vegfc Itga9 |
| Bmp4 Acvr1 | 2.11E-85 | 3.079988031 | 0.657 | 0.182 | 3.55E-82 | 4 | Bmp4 Acvr1 |
| Rgmb Bmpr2 | 8.26E-80 | 1.813786598 | 0.754 | 0.244 | 1.39E-76 | 4 | Rgmb Bmpr2 |
| Vegfc Itgb1 | 7.77E-79 | 1.559050619 | 0.88 | 0.271 | 1.31E-75 | 4 | Vegfc Itgb1 |
| Col5a2 Itgb1 | 7.68E-78 | 1.104393522 | 0.933 | 0.629 | 1.30E-74 | 4 | Col5a2 Itgb1 |
| Bmp6 Acvr1 | 1.87E-77 | 2.866131152 | 0.652 | 0.213 | 3.16E-74 | 4 | Bmp6 Acvr1 |
| Sema3f Plxna3 | 7.60E-77 | 3.763981633 | 0.523 | 0.043 | 1.28E-73 | 4 | Sema3f Plxna3 |
| Hsp90b1 Tlr7 | 2.11E-75 | 1.303090755 | 0.919 | 0.347 | 3.56E-72 | 4 | Hsp90b1 Tlr7 |
| Bmp6 Acvr2a | 2.31E-75 | 2.528859212 | 0.654 | 0.175 | 3.90E-72 | 4 | Bmp6 Acvr2a |
| Vcam1 Itga9 | 5.21E-75 | 2.646628433 | 0.609 | 0.132 | 8.80E-72 | 4 | Vcam1 Itga9 |
| Csf1 Csf1r | 6.59E-75 | 2.161808903 | 0.694 | 0.14 | 1.11E-71 | 4 | Csf1 Csf1r |
| Bmp4 Acvr2a | 7.30E-74 | 2.473768059 | 0.645 | 0.168 | 1.23E-70 | 4 | Bmp4 Acvr2a |
| Psen1 Notch1 | 3.16E-73 | 1.08360546 | 0.949 | 0.314 | 5.34E-70 | 4 | Psen1 Notch1 |
| Sema4d Plxnb2 | 3.23E-72 | 1.472318493 | 0.751 | 0.177 | 5.45E-69 | 4 | Sema4d Plxnb2 |
| Apoe Lrp1 | 7.29E-71 | 1.751161917 | 0.73 | 0.256 | 1.23E-67 | 4 | Apoe Lrp1 |
| Dkk2 Lrp6 | 2.79E-70 | 2.385611275 | 0.669 | 0.251 | 4.71E-67 | 4 | Dkk2 Lrp6 |
| Gnai2 Ccr5 | 1.76E-69 | 1.0868739 | 0.966 | 0.305 | 2.97E-66 | 4 | Gnai2 Ccr5 |
| Calm1 Glp2r | 3.26E-69 | 1.365574257 | 0.865 | 0.356 | 5.51E-66 | 4 | Calm1 Glp2r |
| Vegfa Itgb1 | 5.36E-69 | 1.239020483 | 0.904 | 0.462 | 9.05E-66 | 4 | Vegfa Itgb1 |
| Col4a4 Itgb1 | 7.85E-67 | 1.507011801 | 0.834 | 0.249 | 1.32E-63 | 4 | Col4a4 Itgb1 |
| Gas6 Mertk | 8.72E-67 | 2.085407261 | 0.657 | 0.234 | 1.47E-63 | 4 | Gas6 Mertk |
| Vcam1 Itgb1 | 7.60E-66 | 1.358105273 | 0.882 | 0.323 | 1.28E-62 | 4 | Vcam1 Itgb1 |
| Vegfb Nrp1 | 7.92E-66 | 1.933884681 | 0.643 | 0.18 | 1.34E-62 | 4 | Vegfb Nrp1 |
| Icam1 Egfr | 3.63E-65 | 2.796142727 | 0.564 | 0.13 | 6.12E-62 | 4 | Icam1 Egfr |
| Hsp90b1 Tlr2 | 1.71E-64 | 1.448203249 | 0.679 | 0.16 | 2.88E-61 | 4 | Hsp90b1 Tlr2 |
| Col4a2 Cd93 | 7.95E-64 | 3.148258047 | 0.496 | 0.061 | 1.34E-60 | 4 | Col4a2 Cd93 |
| F8 Lrp 1 | 8.39E-64 | 1.329388458 | 0.828 | 0.465 | 1.42E-60 | 4 | F8 Lrp1 |
| Fgf11 Fgfr1 | 2.28E-63 | 2.096062513 | 0.685 | 0.251 | 3.84E-60 | 4 | Fgf11 Fgfr1 |
| Hspg2 Itgb1 | 7.46E-60 | 0.819133811 | 0.963 | 0.713 | 1.26E-56 | 4 | Hspg2 Itgb1 |
| Apoe Lrp5 | 2.76E-59 | 2.485043201 | 0.582 | 0.17 | 4.66E-56 | 4 | Apoe Lrp5 |
| Vegfa Itga9 | 1.37E-58 | 2.048043016 | 0.622 | 0.251 | 2.30E-55 | 4 | Vegfa Itga9 |
| Ltf Tfrc | 4.53E-56 | 0.885284588 | 0.678 | 0.178 | 7.65E-53 | 4 | Ltf Tfrc |

| | | | | | | | |
|---|---|---|---|---|---|---|---|
| Sema6a Plxna4 | 1.69E-55 | 2.853367523 | 0.502 | 0.114 | 2.85E-52 | 4 | Sema6a Plxna4 |
| Pdgfb Lrp1 | 1.56E-53 | 3.148717932 | 0.403 | 0.031 | 2.64E-50 | 4 | Pdgfb Lrp1 |
| Ncam1 Fgfr1 | 2.39E-52 | 1.067531423 | 0.814 | 0.472 | 4.04E-49 | 4 | Ncam1 Fgfr1 |
| Adam15 Itgb1 | 3.25E-50 | 0.855757707 | 0.919 | 0.517 | 5.48E-47 | 4 | Adam15 Itgb1 |
| Ntn1 Adora2b | 5.55E-50 | 4.071825631 | 0.37 | 0.031 | 9.37E-47 | 4 | Ntn1 Adora2b |
| Tfpi Lrp1 | 1.31E-48 | 1.08793804 | 0.786 | 0.388 | 2.22E-45 | 4 | Tfpi Lrp1 |
| Epo Epor | 1.38E-48 | 2.228585948 | 0.459 | 0.076 | 2.33E-45 | 4 | Epo Epor |
| Vcam1 Itgb2 | 5.94E-48 | 1.159677038 | 0.82 | 0.252 | 1.00E-44 | 4 | Vcam1 Itgb2 |
| Icam2 Itgb2 | 1.91E-47 | 1.225219868 | 0.832 | 0.267 | 3.22E-44 | 4 | Icam2 Itgb2 |
| Jag2 Notch1 | 2.55E-46 | 2.52165192 | 0.451 | 0.083 | 4.31E-43 | 4 | Jag2 Notch1 |
| Adam15 Itga9 | 3.27E-45 | 1.436667123 | 0.634 | 0.287 | 5.52E-42 | 4 | Adam15 Itga9 |
| Col4a3 Itgb1 | 4.57E-43 | 1.29253179 | 0.672 | 0.223 | 7.71E-40 | 4 | Col4a3 Itgb1 |
| Plat Itgb2 | 8.58E-43 | 1.163485014 | 0.787 | 0.271 | 1.45E-39 | 4 | Plat Itgb2 |
| Calm1 Egfr | 2.89E-41 | 0.98339983 | 0.807 | 0.462 | 4.87E-38 | 4 | Calm1 Egfr |
| B2m Tfrc | 4.21E-41 | 0.851716752 | 0.91 | 0.46 | 7.11E-38 | 4 | B2m Tfrc |
| Lama5 Itgb1 | 9.69E-41 | 1.090063147 | 0.783 | 0.348 | 1.64E-37 | 4 | Lama5 Itgb1 |
| Col3a1 Itgb1 | 7.17E-39 | 0.957349503 | 0.846 | 0.446 | 1.21E-35 | 4 | Col3a1 Itgb1 |
| Gnas Adcy7 | 2.80E-38 | 0.589905071 | 0.987 | 0.807 | 4.72E-35 | 4 | Gnas Adcy7 |
| Gnas Adcy9 | 6.52E-38 | 0.84619051 | 0.856 | 0.612 | 1.10E-34 | 4 | Gnas Adcy9 |
| Gnai2 Egfr | 1.43E-37 | 0.873977385 | 0.813 | 0.498 | 2.41E-34 | 4 | Gnai2 Egfr |
| Calm2 Pde1b | 1.46E-37 | 0.630992285 | 0.943 | 0.545 | 2.47E-34 | 4 | Calm2 Pde1b |
| Hras Tlr2 | 3.21E-37 | 1.359678408 | 0.543 | 0.167 | 5.41E-34 | 4 | Hras Tlr2 |
| Col5a1 Itgb1 | 6.02E-37 | 1.024991954 | 0.748 | 0.368 | 1.02E-33 | 4 | Col5a1 Itgb1 |
| Col4a5 Cd93 | 6.18E-37 | 3.838944968 | 0.303 | 0.035 | 1.04E-33 | 4 | Col4a5 Cd93 |
| Nid1 Itgb1 | 9.23E-37 | 1.207138996 | 0.649 | 0.269 | 1.56E-33 | 4 | Nid1 Itgb1 |
| Ntn1 Dcc | 2.13E-36 | 0.96630855 | 0.562 | 0.183 | 3.60E-33 | 4 | Ntn1 Dcc |
| Cx3cl1 Cx3cr1 | 2.29E-35 | 1.850742663 | 0.376 | 0.078 | 3.87E-32 | 4 | Cx3cl1 Cx3cr1 |
| Adam9 Itgb1 | 4.17E-34 | 0.657553044 | 0.933 | 0.627 | 7.03E-31 | 4 | Adam9 Itgb1 |
| Ncam1 Ptpra | 4.86E-33 | 0.562412579 | 0.885 | 0.784 | 8.21E-30 | 4 | Ncam1 Ptpra |
| Calm1 Pde1b | 6.93E-33 | 0.691177613 | 0.942 | 0.53 | 1.17E-29 | 4 | Calm1 Pde1b |
| Egf Egfr | 9.45E-33 | 1.528124863 | 0.46 | 0.155 | 1.60E-29 | 4 | Egf Egfr |
| Tgfa Egfr | 6.39E-32 | 0.986128319 | 0.699 | 0.436 | 1.08E-28 | 4 | Tgfa Egfr |
| Gnai2 Adcy9 | 1.43E-31 | 0.681083111 | 0.849 | 0.583 | 2.41E-28 | 4 | Gnai2 Adcy9 |
| Mfap2 Notch1 | 2.03E-31 | 2.4851087 | 0.324 | 0.063 | 3.43E-28 | 4 | Mfap2 Notch1 |
| Jag1 Notch4 | 2.40E-31 | 4.34644 | 0.228 | 0.012 | 4.06E-28 | 4 | Jag1 Notch4 |
| Adam17 Itgb1 | 5.07E-31 | 0.469532269 | 0.96 | 0.769 | 8.55E-28 | 4 | Adam17 Itgb1 |
| Dll1 Notch4 | 8.17E-30 | 4.600090604 | 0.205 | 0.005 | 1.38E-26 | 4 | Dll1 Notch4 |
| Dll4 Notch4 | 6.85E-29 | 4.912308811 | 0.199 | 0.005 | 1.16E-25 | 4 | Dll4 Notch4 |
| Bmp4 Bmpr1a | 3.17E-27 | 1.731026407 | 0.445 | 0.198 | 5.35E-24 | 4 | Bmp4 Bmpr1a |
| Col3a1 Mag | 8.47E-27 | 2.339629964 | 0.258 | 0.038 | 1.43E-23 | 4 | Col3a1 Mag |
| Dll1 Notch2 | 1.17E-26 | 0.275327291 | 0.891 | 0.327 | 1.97E-23 | 4 | Dll1 Notch2 |
| Psen1 Notch4 | 2.03E-26 | 2.277316822 | 0.234 | 0.026 | 3.43E-23 | 4 | Psen1 Notch4 |
| Ltf Gp9 | 4.38E-26 | 1.360108383 | 0.246 | 0.031 | 7.39E-23 | 4 | Ltf Gp9 |
| Apoe Scarb1 | 1.02E-25 | 2.195418016 | 0.3 | 0.073 | 1.72E-22 | 4 | Apoe Scarb1 |
| Angptl2 Tie1 | 4.06E-25 | 4.419272308 | 0.183 | 0.008 | 6.85E-22 | 4 | Angptl2 Tie1 |
| Ahsg Insr | 4.22E-25 | 1.482234496 | 0.489 | 0.249 | 7.12E-22 | 4 | Ahsg Insr |
| Sfrp1 Fzd6 | 1.50E-24 | 2.800685201 | 0.217 | 0.025 | 2.54E-21 | 4 | Sfrp1 Fzd6 |
| Bmp6 Bmpr1a | 1.78E-24 | 1.601126874 | 0.439 | 0.211 | 3.01E-21 | 4 | Bmp6 Bmpr1a |
| Vegfc Flt1 | 1.24E-23 | 3.725462274 | 0.187 | 0.015 | 2.10E-20 | 4 | Vegfc Flt1 |
| Col1a1 Itgb1 | 2.11E-23 | 0.795900032 | 0.703 | 0.368 | 3.57E-20 | 4 | Col1a1 Itgb1 |
| Vegfa Kdr | 3.14E-23 | 5.9214435 | 0.162 | 0.005 | 5.31E-20 | 4 | Vegfa Kdr |

| | | | | | | | |
|---|---|---|---|---|---|---|---|
| Vegfc Kdr | 5.42E-23 | 5.898499899 | 0.16 | 0.005 | 9.14E-20 | 4 | Vegfc Kdr |
| Ptn Sdc3 | 5.75E-23 | 3.387197367 | 0.205 | 0.026 | 9.70E-20 | 4 | Ptn Sdc3 |
| Adam2 Itgb1 | 1.12E-22 | 0.683542619 | 0.783 | 0.546 | 1.88E-19 | 4 | Adam2 Itgb1 |
| Fn1 Mag | 1.43E-22 | 1.966585294 | 0.229 | 0.036 | 2.41E-19 | 4 | Fn1 Mag |
| Col1a2 Itgb1 | 2.41E-22 | 0.552451253 | 0.882 | 0.645 | 4.07E-19 | 4 | Col1a2 Itgb1 |
| Sema6d Kdr | 2.74E-22 | 5.062360503 | 0.166 | 0.01 | 4.62E-19 | 4 | Sema6d Kdr |
| Fn1 Itga4 | 2.82E-22 | 0.339144491 | 0.687 | 0.314 | 4.76E-19 | 4 | Fn1 Itga4 |
| Dll4 Notch2 | 4.36E-22 | 0.354350135 | 0.867 | 0.337 | 7.35E-19 | 4 | Dll4 Notch2 |
| Jag1 Notch2 | 8.98E-22 | 0.138077227 | 0.93 | 0.739 | 1.52E-18 | 4 | Jag1 Notch2 |
| Inha Acvr2b | 1.97E-21 | 1.697385622 | 0.387 | 0.167 | 3.32E-18 | 4 | Inha Acvr2b |
| Slit2 Dcc | 2.63E-21 | 1.028633515 | 0.412 | 0.152 | 4.44E-18 | 4 | Slit2 Dcc |
| Sema3a Nrp2 | 5.60E-21 | 3.149176667 | 0.193 | 0.026 | 9.45E-18 | 4 | Sema3a Nrp2 |
| Sema3a Nrp1 | 2.17E-20 | 3.886598399 | 0.196 | 0.033 | 3.67E-17 | 4 | Sema3a Nrp1 |
| Sema3c Plxnd1 | 2.41E-20 | 2.92558777 | 0.19 | 0.026 | 4.06E-17 | 4 | Sema3c Plxnd1 |
| Sema3c Nrp2 | 3.06E-20 | 2.376377005 | 0.207 | 0.035 | 5.16E-17 | 4 | Sema3c Nrp2 |
| Tctn1 Tmem67 | 3.39E-20 | 0.666693776 | 0.82 | 0.617 | 5.72E-17 | 4 | Tctn1 Tmem67 |
| Agrn Atp1a3 | 3.51E-20 | 3.502751189 | 0.196 | 0.033 | 5.92E-17 | 4 | Agrn Atp1a3 |
| Timp3 Kdr | 2.84E-19 | 3.082278331 | 0.163 | 0.017 | 4.80E-16 | 4 | Timp3 Kdr |
| Ptdss1 Scarb1 | 7.28E-19 | 1.454138292 | 0.361 | 0.162 | 1.23E-15 | 4 | Ptdss1 Scarb1 |
| Lama4 Itgb1 | 7.31E-19 | 0.600238487 | 0.718 | 0.47 | 1.23E-15 | 4 | Lama4 Itgb1 |
| L1cam Egfr | 1.87E-18 | 2.083773535 | 0.244 | 0.066 | 3.16E-15 | 4 | L1cam Egfr |
| Col1a1 Cd44 | 1.07E-17 | 1.141216234 | 0.429 | 0.215 | 1.81E-14 | 4 | Col1a1 Cd44 |
| Sema3d Nrp1 | 1.08E-17 | 3.085788619 | 0.187 | 0.036 | 1.83E-14 | 4 | Sema3d Nrp1 |
| Icam4 Itgb1 | 1.27E-17 | 0.495997287 | 0.423 | 0.155 | 2.15E-14 | 4 | Icam4 Itgb1 |
| Cthrc1 Fzd6 | 3.50E-17 | 2.930156472 | 0.175 | 0.031 | 5.92E-14 | 4 | Cthrc1 Fzd6 |
| App Ncstn | 3.55E-17 | 0.310018804 | 0.978 | 0.894 | 5.99E-14 | 4 | App Ncstn |
| Lamc1 Itgb1 | 4.96E-17 | 0.695768136 | 0.676 | 0.414 | 8.38E-14 | 4 | Lamc1 Itgb1 |
| Inha Acvr2a | 5.12E-17 | 1.256883235 | 0.307 | 0.112 | 8.64E-14 | 4 | Inha Acvr2a |
| Col4a1 Itgb1 | 1.05E-16 | 0.680463913 | 0.754 | 0.521 | 1.78E-13 | 4 | Col4a1 Itgb1 |
| Col4a4 Itga1 | 1.06E-16 | 2.491302481 | 0.165 | 0.026 | 1.79E-13 | 4 | Col4a4 Itga1 |
| Col18a1 Itgb1 | 1.17E-16 | 1.068884046 | 0.424 | 0.211 | 1.97E-13 | 4 | Col18a1 Itgb1 |
| Ptn Ptprb | 1.39E-16 | 4.212392827 | 0.13 | 0.01 | 2.34E-13 | 4 | Ptn Ptprb |
| Fn1 Robo4 | 2.96E-16 | 4.01821632 | 0.121 | 0.007 | 5.00E-13 | 4 | Fn1 Robo4 |
| Gnai2 Adcy7 | 2.98E-16 | 0.29657166 | 0.979 | 0.79 | 5.03E-13 | 4 | Gnai2 Adcy7 |
| Sema3c Nrp1 | 3.11E-16 | 2.228328127 | 0.21 | 0.056 | 5.26E-13 | 4 | Sema3c Nrp1 |
| Calm2 Egfr | 7.58E-16 | 0.460233911 | 0.805 | 0.521 | 1.28E-12 | 4 | Calm2 Egfr |
| Angpt1 Tie1 | 9.70E-16 | 5.349611274 | 0.109 | 0.003 | 1.64E-12 | 4 | Angpt1 Tie1 |
| Sema3a Plxna1 | 1.34E-15 | 2.806190241 | 0.18 | 0.041 | 2.25E-12 | 4 | Sema3a Plxna1 |
| Angpt1 Itgb1 | 1.54E-15 | 0.800816314 | 0.436 | 0.195 | 2.59E-12 | 4 | Angpt1 Itgb1 |
| App Lrp1 | 2.26E-15 | 0.340577319 | 0.891 | 0.602 | 3.82E-12 | 4 | App Lrp1 |
| Calm1 Insr | 2.37E-15 | 0.337487407 | 0.952 | 0.838 | 3.99E-12 | 4 | Calm1 Insr |
| Mfng Notch2 | 4.29E-15 | 0.162690619 | 0.901 | 0.396 | 7.24E-12 | 4 | Mfng Notch2 |
| Efnb2 Epha4 | 4.46E-15 | 2.69797066 | 0.211 | 0.066 | 7.53E-12 | 4 | Efnb2 Epha4 |
| Lama5 Sdc1 | 4.71E-15 | 2.215818882 | 0.216 | 0.066 | 7.95E-12 | 4 | Lama5 Sdc1 |
| Fgf12 Fgfr3 | 5.06E-15 | 4.303615816 | 0.105 | 0.003 | 8.55E-12 | 4 | Fgf12 Fgfr3 |
| Icam1 Cav1 | 5.95E-15 | 1.997142717 | 0.241 | 0.084 | 1.00E-11 | 4 | Icam1 Cav1 |
| Efnb2 Ephb4 | 7.48E-15 | 3.261563206 | 0.183 | 0.048 | 1.26E-11 | 4 | Efnb2 Ephb4 |
| Col4a6 Cd93 | 8.70E-15 | 3.457177108 | 0.135 | 0.018 | 1.47E-11 | 4 | Col4a6 Cd93 |
| Adam9 Itga6 | 1.39E-14 | 1.471812071 | 0.213 | 0.061 | 2.35E-11 | 4 | Adam9 Itga6 |
| Cd34 Selp | 1.50E-14 | 5.727941647 | 0.097 | 0.002 | 2.54E-11 | 4 | Cd34 Selp |
| Vwf Selp | 1.53E-14 | 5.582341259 | 0.097 | 0.002 | 2.58E-11 | 4 | Vwf Selp |

| | | | | | | | |
|---|---|---|---|---|---|---|---|
| Slit2 Robo4 | 2.41E-14 | 5.866996401 | 0.096 | 0.002 | 4.06E-11 | 4 | Slit2 Robo4 |
| Vegfa Flt1 | 2.62E-14 | 3.112628498 | 0.19 | 0.056 | 4.42E-11 | 4 | Vegfa Flt1 |
| Tnfsf10 Tnfrsf10b | 3.27E-14 | 3.280587054 | 0.115 | 0.01 | 5.51E-11 | 4 | Tnfsf10 Tnfrsf10b |
| Vcan Tlr1 | 5.72E-14 | 1.294695348 | 0.18 | 0.041 | 9.65E-11 | 4 | Vcan Tlr1 |
| Bmp8b Acvr2b | 7.60E-14 | 2.020743448 | 0.192 | 0.054 | 1.28E-10 | 4 | Bmp8b Acvr2b |
| Lamc1 Itgb4 | 9.94E-14 | 5.081025436 | 0.096 | 0.003 | 1.68E-10 | 4 | Lamc1 Itgb4 |
| Lamc1 Itga6 | 1.24E-13 | 1.535261481 | 0.175 | 0.043 | 2.10E-10 | 4 | Lamc1 Itga6 |
| Slit2 Sdc1 | 1.30E-13 | 1.956018 | 0.187 | 0.053 | 2.19E-10 | 4 | Slit2 Sdc1 |
| Vwf Itga2b | 2.08E-13 | 1.412011046 | 0.241 | 0.089 | 3.51E-10 | 4 | Vwf Itga2b |
| Lama5 Itgb4 | 2.35E-13 | 4.724378068 | 0.097 | 0.005 | 3.96E-10 | 4 | Lama5 Itgb4 |
| Gnai2 Igf1r | 2.54E-13 | 0.381633762 | 0.913 | 0.635 | 4.29E-10 | 4 | Gnai2 Igf1r |
| Lamb1 Itgb1 | 3.52E-13 | 1.09534369 | 0.247 | 0.094 | 5.94E-10 | 4 | Lamb1 Itgb1 |
| Fn1 Itga6 | 4.53E-13 | 1.696562209 | 0.192 | 0.056 | 7.64E-10 | 4 | Fn1 Itga6 |
| Vegfb Flt1 | 4.68E-13 | 2.492370216 | 0.156 | 0.036 | 7.91E-10 | 4 | Vegfb Flt1 |
| Lama4 Itga6 | 4.99E-13 | 1.663953194 | 0.169 | 0.043 | 8.43E-10 | 4 | Lama4 Itga6 |
| Jag2 Notch4 | 8.03E-13 | 4.828052236 | 0.09 | 0.003 | 1.36E-09 | 4 | Jag2 Notch4 |
| Ptn Plxnb2 | 8.09E-13 | 2.210892468 | 0.207 | 0.073 | 1.37E-09 | 4 | Ptn Plxnb2 |
| Icam1 Itgb2 | 9.21E-13 | 0.513029971 | 0.609 | 0.297 | 1.56E-09 | 4 | Icam1 Itgb2 |
| Bmp8b Acvr1 | 9.83E-13 | 1.836014135 | 0.156 | 0.036 | 1.66E-09 | 4 | Bmp8b Acvr1 |
| Inha Acvr1 | 2.24E-12 | 1.187507898 | 0.315 | 0.147 | 3.78E-09 | 4 | Inha Acvr1 |
| Bmp8b Bmpr2 | 2.25E-12 | 1.433983018 | 0.186 | 0.054 | 3.80E-09 | 4 | Bmp8b Bmpr2 |
| Sema3a Plxna4 | 2.56E-12 | 3.260016847 | 0.129 | 0.025 | 4.32E-09 | 4 | Sema3a Plxna4 |
| Lama5 Itga6 | 4.22E-12 | 1.405360988 | 0.174 | 0.048 | 7.12E-09 | 4 | Lama5 Itga6 |
| Tnc Ptprb | 9.21E-12 | 3.477839733 | 0.117 | 0.02 | 1.55E-08 | 4 | Tnc Ptprb |
| Col4a3 Itga1 | 2.27E-11 | 1.775331671 | 0.129 | 0.026 | 3.84E-08 | 4 | Col4a3 Itga1 |
| Adam2 Itga9 | 2.41E-11 | 0.682250172 | 0.532 | 0.348 | 4.06E-08 | 4 | Adam2 Itga9 |
| Col18a1 Kdr | 6.73E-11 | 4.791888197 | 0.076 | 0.003 | 1.14E-07 | 4 | Col18a1 Kdr |
| Adam2 Itga6 | 6.84E-11 | 1.244356235 | 0.175 | 0.056 | 1.15E-07 | 4 | Adam2 Itga6 |
| Bmp8b Acvr2a | 7.64E-11 | 1.241858977 | 0.159 | 0.045 | 1.29E-07 | 4 | Bmp8b Acvr2a |
| Hspg2 Sdc1 | 8.28E-11 | 1.565691357 | 0.276 | 0.147 | 1.40E-07 | 4 | Hspg2 Sdc1 |
| Vcan Tlr2 | 1.66E-10 | 2.108217326 | 0.13 | 0.031 | 2.80E-07 | 4 | Vcan Tlr2 |
| Gnai2 S1pr1 | 2.57E-10 | 0.406963647 | 0.855 | 0.419 | 4.33E-07 | 4 | Gnai2 S1pr1 |
| Lama1 Itgb1 | 2.69E-10 | 1.886931626 | 0.124 | 0.028 | 4.54E-07 | 4 | Lama1 Itgb1 |
| Calm1 Ptpra | 2.76E-10 | 0.20415288 | 0.985 | 0.922 | 4.67E-07 | 4 | Calm1 Ptpra |
| Icam4 Itga2b | 3.03E-10 | 1.209985621 | 0.109 | 0.02 | 5.12E-07 | 4 | Icam4 Itga2b |
| Sema4d Met | 3.09E-10 | 1.273861375 | 0.096 | 0.013 | 5.21E-07 | 4 | Sema4d Met |
| Hras Insr | 3.43E-10 | 0.492133359 | 0.772 | 0.606 | 5.78E-07 | 4 | Hras Insr |
| Psap Lrp1 | 1.36E-09 | 0.197771163 | 0.885 | 0.612 | 2.30E-06 | 4 | Psap Lrp1 |
| Inha Acvr1b | 1.42E-09 | 0.861962862 | 0.325 | 0.173 | 2.39E-06 | 4 | Inha Acvr1b |
| Gdf6 Bmpr2 | 1.63E-09 | 2.534432002 | 0.099 | 0.018 | 2.76E-06 | 4 | Gdf6 Bmpr2 |
| Cxcl12 Itgb1 | 1.90E-09 | 1.458558939 | 0.229 | 0.106 | 3.21E-06 | 4 | Cxcl12 Itgb1 |
| Gnai2 Adra2a | 2.07E-09 | 0.287072711 | 0.189 | 0.066 | 3.49E-06 | 4 | Gnai2 Adra2a |
| Fgf2 Sdc3 | 2.61E-09 | 1.084556507 | 0.27 | 0.137 | 4.41E-06 | 4 | Fgf2 Sdc3 |
| Vcan Itga4 | 3.94E-09 | 0.903748293 | 0.177 | 0.066 | 6.65E-06 | 4 | Vcan Itga4 |
| Lamb1 Itga6 | 5.78E-09 | 4.207545325 | 0.07 | 0.007 | 9.75E-06 | 4 | Lamb1 Itga6 |
| Pdgfb S1pr1 | 7.99E-09 | 0.678131408 | 0.382 | 0.193 | 1.35E-05 | 4 | Pdgfb S1pr1 |
| Serping1 Selp | 1.10E-08 | 9.776247877 | 0.052 | 0 | 1.85E-05 | 4 | Serping1 Selp |
| Efnb2 Ephb1 | 1.12E-08 | 3.462965845 | 0.072 | 0.008 | 1.89E-05 | 4 | Efnb2 Ephb1 |
| Sost Lrp5 | 1.25E-08 | 1.980039307 | 0.126 | 0.038 | 2.10E-05 | 4 | Sost Lrp5 |
| Papln Sirpa | 1.57E-08 | 2.100670933 | 0.159 | 0.063 | 2.65E-05 | 4 | Papln Sirpa |
| Nlgn2 Nrxn3 | 3.81E-08 | 2.51656266 | 0.085 | 0.017 | 6.43E-05 | 4 | Nlgn2 Nrxn3 |

| | | | | | | | |
|---|---|---|---|---|---|---|---|
| Rarres2 Cmklr1 | 4.95E-08 | 2.182668529 | 0.183 | 0.087 | 8.36E-05 | 4 | Rarres2 Cmklr1 |
| Tgm2 Itga9 | 5.18E-08 | 0.199834318 | 0.589 | 0.409 | 8.74E-05 | 4 | Tgm2 Itga9 |
| Cd24a Selp | 5.45E-08 | 2.203732042 | 0.064 | 0.007 | 9.19E-05 | 4 | Cd24a Selp |
| Lamb1 Itgb4 | 7.88E-08 | 9.02498033 | 0.046 | 0 | 0.000132947 | 4 | Lamb1 Itgb4 |
| Col4a5 Itgb1 | 8.32E-08 | 1.033399891 | 0.322 | 0.206 | 0.000140453 | 4 | Col4a5 Itgb1 |
| Thbs2 Itga4 | 8.73E-08 | 1.285742591 | 0.115 | 0.035 | 0.000147395 | 4 | Thbs2 Itga4 |
| Col4a5 Cd47 | 1.29E-07 | 0.805649152 | 0.324 | 0.208 | 0.000217788 | 4 | Col4a5 Cd47 |
| Egf Cav1 | 1.29E-07 | 1.092767708 | 0.21 | 0.102 | 0.000217953 | 4 | Egf Cav1 |
| Mdk Ptprb | 1.40E-07 | 2.555260642 | 0.108 | 0.033 | 0.000236341 | 4 | Mdk Ptprb |
| Adam10 Axl | 1.47E-07 | 0.144090475 | 0.214 | 0.419 | 0.000247617 | 4 | Adam10 Axl |
| Vegfc Flt4 | 2.11E-07 | 10.16171977 | 0.043 | 0 | 0.00035554 | 4 | Vegfc Flt4 |
| Fgf12 Fgfr2 | 2.30E-07 | 0.695189331 | 0.286 | 0.153 | 0.000388753 | 4 | Fgf12 Fgfr2 |
| Sema3a Plxna3 | 2.40E-07 | 1.141007803 | 0.112 | 0.035 | 0.000405116 | 4 | Sema3a Plxna3 |
| Serping1 Lrp1 | 2.47E-07 | 0.43867499 | 0.298 | 0.162 | 0.000416562 | 4 | Serping1 Lrp1 |
| Col6a1 Itga6 | 4.84E-07 | 1.636897333 | 0.103 | 0.031 | 0.000816613 | 4 | Col6a1 Itga6 |
| Col1a2 Flt4 | 6.20E-07 | 3.95012854 | 0.052 | 0.005 | 0.001046793 | 4 | Col1a2 Flt4 |
| Ntn1 Unc5b | 6.55E-07 | 2.400357888 | 0.094 | 0.028 | 0.001105437 | 4 | Ntn1 Unc5b |
| Scgb3a1 Marco | 8.20E-07 | 3.128969852 | 0.055 | 0.007 | 0.001384523 | 4 | Scgb3a1 Marco |
| Ntn4 Dcc | 8.23E-07 | 0.18993646 | 0.513 | 0.365 | 0.001388576 | 4 | Ntn4 Dcc |
| Nampt Insr | 8.50E-07 | 0.142938525 | 0.838 | 0.683 | 0.001435565 | 4 | Nampt Insr |
| Gdf6 Bmpr1a | 8.67E-07 | 2.379699052 | 0.069 | 0.013 | 0.001464068 | 4 | Gdf6 Bmpr1a |
| Sost Lrp6 | 1.04E-06 | 1.56135374 | 0.165 | 0.079 | 0.001757583 | 4 | Sost Lrp6 |
| Fgf11 Fgfr3 | 1.48E-06 | 1.901672609 | 0.087 | 0.025 | 0.002496251 | 4 | Fgf11 Fgfr3 |
| Mdk Sdc3 | 2.61E-06 | 1.888593292 | 0.154 | 0.076 | 0.004410895 | 4 | Mdk Sdc3 |
| Fn1 Flt4 | 2.68E-06 | 3.314348112 | 0.048 | 0.005 | 0.004522667 | 4 | Fn1 Flt4 |
| Bst1 Cav1 | 3.87E-06 | 1.19634688 | 0.117 | 0.045 | 0.006531912 | 4 | Bst1 Cav1 |
| Angpt1 Tek | 4.67E-06 | 1.206093201 | 0.108 | 0.04 | 0.00787991 | 4 | Angpt1 Tek |
| Col1a1 Flt4 | 5.51E-06 | 3.951487496 | 0.042 | 0.003 | 0.009296736 | 4 | Col1a1 Flt4 |
| Angpt2 Tie1 | 6.20E-06 | 5.86549265 | 0.037 | 0.002 | 0.010473053 | 4 | Angpt2 Tie1 |
| Lpl Gpihbp1 | 6.24E-06 | 5.526799377 | 0.037 | 0.002 | 0.010534863 | 4 | Lpl Gpihbp1 |
| Col1a2 Cd44 | 8.07E-06 | 0.489853884 | 0.552 | 0.477 | 0.013622233 | 4 | Col1a2 Cd44 |
| Angptl4 Tie1 | 1.03E-05 | 5.270085525 | 0.036 | 0.002 | 0.017314601 | 4 | Angptl4 Tie1 |
| Pigf Flt1 | 1.74E-05 | 0.445044177 | 0.192 | 0.099 | 0.029343481 | 4 | Pigf Flt1 |
| Vcan Selp | 1.75E-05 | 8.630472111 | 0.03 | 0 | 0.029568118 | 4 | Vcan Selp |
| Bmp8b Bmpr1a | 2.13E-05 | 0.67215163 | 0.1 | 0.038 | 0.036020971 | 4 | Bmp8b Bmpr1a |
| Gpc3 Cd81 | 2.62E-05 | 1.029400308 | 0.147 | 0.074 | 0.044162162 | 4 | Gpc3 Cd81 |
| Lama1 Itga6 | 2.89E-05 | 2.927112139 | 0.033 | 0.002 | 0.048707326 | 4 | Lama1 Itga6 |
| Col2a1 Itgb1 | 3.26E-260 | 7.869002908 | 0.984 | 0.007 | 5.50E-257 | 6 | Col2a1 Itgb1 |
| Col11a1 Itgb1 | 2.61E-258 | 7.489866359 | 0.984 | 0.01 | 4.41E-255 | 6 | Col11a1 Itgb1 |
| Chad Itgb1 | 2.70E-257 | 7.272500198 | 0.984 | 0.011 | 4.56E-254 | 6 | Chad Itgb1 |
| Bmp2 Bmpr1a | 1.75E-233 | 7.091795572 | 0.935 | 0.025 | 2.95E-230 | 6 | Bmp2 Bmpr1a |
| Edil3 Itgb5 | 1.97E-228 | 6.766813393 | 0.908 | 0.017 | 3.32E-225 | 6 | Edil3 Itgb5 |
| Bmp2 Bmpr2 | 1.59E-224 | 5.779836464 | 0.948 | 0.048 | 2.68E-221 | 6 | Bmp2 Bmpr2 |
| Spp1 Itgb1 | 6.80E-224 | 6.733661968 | 0.882 | 0.008 | 1.15E-220 | 6 | Spp1 Itgb1 |
| Pdgfc Pdgfrb | 9.76E-223 | 8.226831926 | 0.863 | 0.004 | 1.65E-219 | 6 | Pdgfc Pdgfrb |
| Edil3 Itgav | 1.81E-219 | 5.097115031 | 0.938 | 0.039 | 3.06E-216 | 6 | Edil3 Itgav |
| Bmp5 Bmpr1a | 1.18E-214 | 8.890913597 | 0.833 | 0.003 | 1.98E-211 | 6 | Bmp5 Bmpr1a |
| Spp1 Itgav | 3.58E-214 | 6.400991993 | 0.853 | 0.008 | 6.04E-211 | 6 | Spp1 Itgav |
| Rspo3 Lgr4 | 4.43E-212 | 9.184486139 | 0.833 | 0.007 | 7.48E-209 | 6 | Rspo3 Lgr4 |
| Bmp5 Bmpr2 | 1.26E-211 | 6.431720512 | 0.846 | 0.009 | 2.13E-208 | 6 | Bmp5 Bmpr2 |
| Rspo3 Lrp6 | 1.30E-211 | 5.101684686 | 0.922 | 0.04 | 2.20E-208 | 6 | Rspo3 Lrp6 |

| | | | | | | | |
|---|---|---|---|---|---|---|---|
| Ndp Lgr4 | 1.85E-208 | 13.27843023 | 0.807 | 0.001 | 3.12E-205 | 6 | Ndp Lgr4 |
| Bmp2 Acvr1 | 1.15E-205 | 6.225908099 | 0.869 | 0.034 | 1.93E-202 | 6 | Bmp2 Acvr1 |
| Myoc Fzd7 | 2.78E-203 | 10.76716406 | 0.791 | 0.001 | 4.70E-200 | 6 | Myoc Fzd7 |
| Hbegf Erbb2 | 3.06E-201 | 9.123268634 | 0.804 | 0.009 | 5.17E-198 | 6 | Hbegf Erbb2 |
| A2m Lrp1 | 1.76E-198 | 4.704664205 | 0.948 | 0.107 | 2.97E-195 | 6 | A2m Lrp1 |
| Col2a1 Itga2b | 3.42E-198 | 15.62699937 | 0.771 | 0 | 5.78E-195 | 6 | Col2a1 Itga2b |
| Myoc Fzd1 | 3.42E-198 | 15.50773178 | 0.771 | 0 | 5.78E-195 | 6 | Myoc Fzd1 |
| Wnt3a Fzd2 | 8.09E-195 | 8.741970303 | 0.775 | 0.005 | 1.37E-191 | 6 | Wnt3a Fzd2 |
| Bmp5 Acvr1 | 1.70E-193 | 7.028719405 | 0.775 | 0.005 | 2.88E-190 | 6 | Bmp5 Acvr1 |
| Wnt3a Ryk | 2.69E-193 | 7.049785111 | 0.824 | 0.031 | 4.55E-190 | 6 | Wnt3a Ryk |
| Spp1 Itga9 | 5.28E-191 | 7.64142493 | 0.765 | 0.005 | 8.91E-188 | 6 | Spp1 Itga9 |
| Bmp5 Acvr2b | 2.53E-190 | 5.912074055 | 0.781 | 0.008 | 4.27E-187 | 6 | Bmp5 Acvr2b |
| Bmp2 Acvr2b | 6.47E-190 | 5.053859527 | 0.866 | 0.052 | 1.09E-186 | 6 | Bmp2 Acvr2b |
| Calm2 Pde1a | 4.16E-189 | 4.903553332 | 0.859 | 0.044 | 7.02E-186 | 6 | Calm2 Pde1a |
| Mfge8 Pdgfrb | 6.90E-189 | 4.827522235 | 0.866 | 0.053 | 1.16E-185 | 6 | Mfge8 Pdgfrb |
| Pdgfa Pdgfrb | 1.02E-188 | 5.854586551 | 0.807 | 0.025 | 1.72E-185 | 6 | Pdgfa Pdgfrb |
| Wnt3a Lrp1 | 1.57E-188 | 5.976145518 | 0.837 | 0.044 | 2.65E-185 | 6 | Wnt3a Lrp1 |
| Tgfb3 Tgfbr3 | 4.10E-183 | 5.380748397 | 0.797 | 0.025 | 6.92E-180 | 6 | Tgfb3 Tgfbr3 |
| Lrpap1 Vldlr | 1.92E-178 | 4.868842202 | 0.82 | 0.043 | 3.23E-175 | 6 | Lrpap1 Vldlr |
| Rspo3 Fzd8 | 7.63E-178 | 9.614694565 | 0.722 | 0.007 | 1.29E-174 | 6 | Rspo3 Fzd8 |
| Fn1 Sdc2 | 5.85E-177 | 4.447962922 | 0.876 | 0.083 | 9.87E-174 | 6 | Fn1 Sdc2 |
| Wnt3a Atp6ap2 | 6.20E-176 | 4.454767859 | 0.843 | 0.055 | 1.05E-172 | 6 | Wnt3a Atp6ap2 |
| Col11a1 Ddr1 | 7.34E-176 | 11.03487541 | 0.699 | 0.001 | 1.24E-172 | 6 | Col11a1 Ddr1 |
| Col2a1 Ddr1 | 7.34E-176 | 11.00958187 | 0.699 | 0.001 | 1.24E-172 | 6 | Col2a1 Ddr1 |
| Timp1 Cd63 | 2.05E-174 | 4.952175713 | 0.833 | 0.064 | 3.46E-171 | 6 | Timp1 Cd63 |
| Rims1 Cacna1c | 1.53E-173 | 9.025566052 | 0.703 | 0.005 | 2.58E-170 | 6 | Rims1 Cacna1c |
| Hbegf Cd9 | 1.96E-173 | 7.044760665 | 0.735 | 0.019 | 3.31E-170 | 6 | Hbegf Cd9 |
| Wnt3a Fzd5 | 2.95E-173 | 6.747761793 | 0.752 | 0.025 | 4.99E-170 | 6 | Wnt3a Fzd5 |
| Nrg1 Erbb2 | 1.04E-172 | 6.584924002 | 0.755 | 0.027 | 1.76E-169 | 6 | Nrg1 Erbb2 |
| Wnt3a Fzd7 | 4.97E-172 | 9.797677665 | 0.703 | 0.007 | 8.39E-169 | 6 | Wnt3a Fzd7 |
| Fgf2 Sdc2 | 8.90E-172 | 6.000790212 | 0.765 | 0.032 | 1.50E-168 | 6 | Fgf2 Sdc2 |
| Tgfa Erbb2 | 1.48E-168 | 4.688022862 | 0.833 | 0.069 | 2.50E-165 | 6 | Tgfa Erbb2 |
| Lamb3 Cd151 | 3.62E-168 | 4.481160351 | 0.794 | 0.039 | 6.11E-165 | 6 | Lamb3 Cd151 |
| Wnt3a Lrp6 | 7.41E-168 | 4.292457393 | 0.83 | 0.057 | 1.25E-164 | 6 | Wnt3a Lrp6 |
| Hsp90b1 Erbb2 | 7.66E-168 | 4.903637543 | 0.846 | 0.087 | 1.29E-164 | 6 | Hsp90b1 Erbb2 |
| Rspo3 Sdc4 | 1.00E-167 | 7.035865445 | 0.709 | 0.014 | 1.70E-164 | 6 | Rspo3 Sdc4 |
| Wnt3a Fzd1 | 2.20E-167 | 9.039858884 | 0.68 | 0.004 | 3.71E-164 | 6 | Wnt3a Fzd1 |
| F13a1 Itgb1 | 8.08E-165 | 3.133097234 | 0.814 | 0.038 | 1.36E-161 | 6 | F13a1 Itgb1 |
| Hbegf Egfr | 4.86E-164 | 4.7532949 | 0.804 | 0.062 | 8.20E-161 | 6 | Hbegf Egfr |
| Col6a3 Itgb1 | 5.12E-164 | 3.034110367 | 0.984 | 0.218 | 8.64E-161 | 6 | Col6a3 Itgb1 |
| Serpine2 Lrp1 | 2.06E-160 | 3.30366491 | 0.948 | 0.153 | 3.47E-157 | 6 | Serpine2 Lrp1 |
| Wnt3a Fzd8 | 3.70E-159 | 9.820682873 | 0.654 | 0.005 | 6.24E-156 | 6 | Wnt3a Fzd8 |
| Tgfb3 Tgfbr1 | 1.81E-157 | 4.36869045 | 0.768 | 0.047 | 3.06E-154 | 6 | Tgfb3 Tgfbr1 |
| Dlk1 Notch2 | 5.90E-157 | 4.563260911 | 0.712 | 0.02 | 9.96E-154 | 6 | Dlk1 Notch2 |
| Rgma Neo1 | 2.30E-155 | 4.88896961 | 0.784 | 0.07 | 3.88E-152 | 6 | Rgma Neo1 |
| F13a1 Itga9 | 6.99E-152 | 4.241748947 | 0.696 | 0.021 | 1.18E-148 | 6 | F13a1 Itga9 |
| Sema4b Dcbld2 | 4.83E-151 | 4.897743906 | 0.686 | 0.021 | 8.16E-148 | 6 | Sema4b Dcbld2 |
| Tgfb2 Tgfbr3 | 4.59E-150 | 5.109967426 | 0.719 | 0.041 | 7.74E-147 | 6 | Tgfb2 Tgfbr3 |
| Myoc Fzd3 | 1.70E-149 | 11.01952518 | 0.608 | 0.001 | 2.87E-146 | 6 | Myoc Fzd3 |
| Col2a1 Itga1 | 1.91E-149 | 15.65190807 | 0.605 | 0 | 3.22E-146 | 6 | Col2a1 Itga1 |
| Calm1 Pde1a | 1.99E-149 | 4.144314338 | 0.752 | 0.048 | 3.37E-146 | 6 | Calm1 Pde1a |

| | | | | | | | |
|---|---|---|---|---|---|---|---|
| Fgf2 Gpc4 | 3.07E-149 | 5.932813938 | 0.693 | 0.034 | 5.18E-146 | 6 | Fgf2 Gpc4 |
| Dll3 Notch2 | 2.44E-148 | 4.493625134 | 0.66 | 0.013 | 4.11E-145 | 6 | Dll3 Notch2 |
| Serpinc1 Sdc2 | 1.10E-147 | 6.827276918 | 0.634 | 0.011 | 1.86E-144 | 6 | Serpinc1 Sdc2 |
| Sorbs1 Itgb5 | 7.00E-147 | 3.268906692 | 0.902 | 0.181 | 1.18E-143 | 6 | Sorbs1 Itgb5 |
| Sema4g Plxnb2 | 1.53E-146 | 3.659562137 | 0.791 | 0.065 | 2.58E-143 | 6 | Sema4g Plxnb2 |
| Fn1 Il17rc | 9.55E-141 | 4.831589172 | 0.748 | 0.08 | 1.61E-137 | 6 | Fn1 Il17rc |
| Col6a2 Itgb1 | 1.78E-136 | 2.567670535 | 0.958 | 0.231 | 3.01E-133 | 6 | Col6a2 Itgb1 |
| Ncam1 Cacna1c | 7.50E-134 | 4.05847057 | 0.686 | 0.038 | 1.27E-130 | 6 | Ncam1 Cacna1c |
| Col6a1 Itgb1 | 1.85E-133 | 2.408353982 | 0.974 | 0.279 | 3.13E-130 | 6 | Col6a1 Itgb1 |
| Bmp5 Acvr2a | 5.13E-131 | 5.877678807 | 0.572 | 0.008 | 8.67E-128 | 6 | Bmp5 Acvr2a |
| Tgfb3 Tgfbr2 | 1.63E-130 | 2.631456429 | 0.84 | 0.097 | 2.74E-127 | 6 | Tgfb3 Tgfbr2 |
| Spp1 Itga5 | 1.68E-130 | 7.422328551 | 0.549 | 0.003 | 2.83E-127 | 6 | Spp1 Itga5 |
| Col8a1 Itga1 | 1.62E-129 | 6.153348441 | 0.605 | 0.025 | 2.73E-126 | 6 | Col8a1 Itga1 |
| Bmp2 Acvr2a | 3.83E-129 | 5.7697842 | 0.631 | 0.037 | 6.47E-126 | 6 | Bmp2 Acvr2a |
| Rspo3 Lgr6 | 1.11E-128 | 11.23574476 | 0.533 | 0.001 | 1.88E-125 | 6 | Rspo3 Lgr6 |
| Lamb3 Itgb1 | 2.34E-128 | 2.403862029 | 0.912 | 0.129 | 3.96E-125 | 6 | Lamb3 Itgb1 |
| Ltbp3 Itgb5 | 8.35E-128 | 2.681425491 | 0.892 | 0.199 | 1.41E-124 | 6 | Ltbp3 Itgb5 |
| Lamc3 Itgb1 | 2.05E-125 | 3.694892888 | 0.667 | 0.042 | 3.46E-122 | 6 | Lamc3 Itgb1 |
| Vtn Itgav | 2.38E-125 | 6.886991908 | 0.542 | 0.007 | 4.02E-122 | 6 | Vtn Itgav |
| Vtn Itgb1 | 4.77E-125 | 5.595952239 | 0.559 | 0.01 | 8.05E-122 | 6 | Vtn Itgb1 |
| Vtn Itgb5 | 3.60E-124 | 6.608190653 | 0.526 | 0.003 | 6.08E-121 | 6 | Vtn Itgb5 |
| Fgf18 Fgfr2 | 2.83E-122 | 5.213902627 | 0.559 | 0.013 | 4.78E-119 | 6 | Fgf18 Fgfr2 |
| Farp2 Plxna2 | 1.06E-119 | 3.289803733 | 0.758 | 0.12 | 1.79E-116 | 6 | Farp2 Plxna2 |
| Vtn Cd47 | 1.22E-119 | 5.446097407 | 0.539 | 0.01 | 2.06E-116 | 6 | Vtn Cd47 |
| Spp1 Cd44 | 4.44E-118 | 4.773194168 | 0.523 | 0.007 | 7.50E-115 | 6 | Spp1 Cd44 |
| Fgf18 Fgfr1 | 5.58E-118 | 3.900387221 | 0.614 | 0.033 | 9.43E-115 | 6 | Fgf18 Fgfr1 |
| Thbs1 Itga2b | 9.48E-118 | 4.258198559 | 0.647 | 0.054 | 1.60E-114 | 6 | Thbs1 Itga2b |
| Calm1 Cacna1c | 1.25E-117 | 3.662576323 | 0.644 | 0.042 | 2.11E-114 | 6 | Calm1 Cacna1c |
| Lamb3 Itga3 | 6.02E-117 | 5.693350571 | 0.503 | 0.004 | 1.02E-113 | 6 | Lamb3 Itga3 |
| Wnt3a Apcdd1 | 1.38E-114 | 10.90132186 | 0.484 | 0.002 | 2.32E-111 | 6 | Wnt3a Apcdd1 |
| Fgf2 Fgfr2 | 5.72E-114 | 3.355857596 | 0.696 | 0.086 | 9.66E-111 | 6 | Fgf2 Fgfr2 |
| Col6a3 Itga1 | 8.38E-113 | 4.746332681 | 0.605 | 0.051 | 1.41E-109 | 6 | Col6a3 Itga1 |
| Tbc1d24 Neo1 | 9.22E-113 | 2.877174308 | 0.732 | 0.089 | 1.56E-109 | 6 | Tbc1d24 Neo1 |
| Hbegf Erbb4 | 9.26E-113 | 5.899956181 | 0.516 | 0.013 | 1.56E-109 | 6 | Hbegf Erbb4 |
| F2 Itga2b | 3.89E-112 | 7.310801327 | 0.49 | 0.006 | 6.56E-109 | 6 | F2 Itga2b |
| Gdf11 Acvr1b | 1.05E-110 | 3.159633075 | 0.663 | 0.067 | 1.77E-107 | 6 | Gdf11 Acvr1b |
| Gnai2 Tbxa2r | 9.78E-110 | 3.706860675 | 0.582 | 0.036 | 1.65E-106 | 6 | Gnai2 Tbxa2r |
| Fn1 Itgav | 4.05E-109 | 2.179011337 | 0.918 | 0.376 | 6.83E-106 | 6 | Fn1 Itgav |
| Thbs1 Itgb3 | 2.06E-107 | 3.577809707 | 0.585 | 0.04 | 3.48E-104 | 6 | Thbs1 Itgb3 |
| Rtn4 Cntnap1 | 3.53E-107 | 3.255613119 | 0.712 | 0.122 | 5.97E-104 | 6 | Rtn4 Cntnap1 |
| Il16 Grin2d | 8.33E-107 | 5.85049903 | 0.52 | 0.023 | 1.41E-103 | 6 | Il16 Grin2d |
| Amelx Lamp1 | 1.90E-106 | 5.735710903 | 0.487 | 0.01 | 3.21E-103 | 6 | Amelx Lamp1 |
| Fn1 Itga2b | 4.40E-106 | 3.588851295 | 0.742 | 0.151 | 7.43E-103 | 6 | Fn1 Itga2b |
| Col6a2 Itga1 | 6.30E-106 | 4.210009049 | 0.595 | 0.054 | 1.06E-102 | 6 | Col6a2 Itga1 |
| Fbn1 Itgav | 1.69E-105 | 1.891636804 | 0.951 | 0.398 | 2.85E-102 | 6 | Fbn1 Itgav |
| Gdf11 Acvr2b | 2.06E-105 | 2.916587499 | 0.644 | 0.06 | 3.48E-102 | 6 | Gdf11 Acvr2b |
| Pros1 Tyro3 | 2.10E-104 | 4.835915646 | 0.529 | 0.029 | 3.55E-101 | 6 | Pros1 Tyro3 |
| Fgf2 Fgfrl1 | 9.22E-104 | 7.38183519 | 0.448 | 0.003 | 1.56E-100 | 6 | Fgf2 Fgfrl1 |
| Vtn Itga2b | 2.18E-103 | 9.85663519 | 0.438 | 0.001 | 3.68E-100 | 6 | Vtn Itga2b |
| Fgf2 Sdc4 | 3.76E-103 | 3.651526999 | 0.641 | 0.079 | 6.34E-100 | 6 | Fgf2 Sdc4 |
| Pdap1 Pdgfrb | 6.16E-103 | 4.327354666 | 0.572 | 0.049 | 1.04E-99 | 6 | Pdap1 Pdgfrb |

| | | | | | | | |
|---|---|---|---|---|---|---|---|
| Inhba Tgfbr3 | 7.31E-102 | 3.997580062 | 0.552 | 0.037 | 1.23E-98 | 6 | Inhba Tgfbr3 |
| Col6a1 Itga1 | 8.85E-102 | 4.072217407 | 0.598 | 0.065 | 1.49E-98 | 6 | Col6a1 Itga1 |
| Nrg1 Erbb4 | 1.46E-101 | 5.170621484 | 0.507 | 0.024 | 2.47E-98 | 6 | Nrg1 Erbb4 |
| Fn1 Itga8 | 3.42E-101 | 4.12779981 | 0.605 | 0.07 | 5.77E-98 | 6 | Fn1 Itga8 |
| Tfpi Vldlr | 2.72E-100 | 3.830105079 | 0.556 | 0.037 | 4.60E-97 | 6 | Tfpi Vldlr |
| Tgfb2 Tgfbr1 | 6.14E-100 | 2.855239277 | 0.719 | 0.119 | 1.04E-96 | 6 | Tgfb2 Tgfbr1 |
| Tgfb3 Acvrl1 | 2.51E-99 | 4.139664144 | 0.539 | 0.036 | 4.24E-96 | 6 | Tgfb3 Acvrl1 |
| Fbn1 Itgb3 | 2.64E-98 | 2.994485217 | 0.693 | 0.113 | 4.46E-95 | 6 | Fbn1 Itgb3 |
| App Gpc1 | 1.42E-96 | 5.00120038 | 0.441 | 0.008 | 2.40E-93 | 6 | App Gpc1 |
| Fn1 Itgb3 | 1.88E-96 | 3.170626389 | 0.67 | 0.113 | 3.17E-93 | 6 | Fn1 Itgb3 |
| Ctf1 Il6st | 2.58E-96 | 3.463760133 | 0.601 | 0.067 | 4.35E-93 | 6 | Ctf1 Il6st |
| Pdgfc Pdgfra | 2.70E-96 | 8.299566261 | 0.431 | 0.007 | 4.55E-93 | 6 | Pdgfc Pdgfra |
| Dcn Erbb4 | 5.88E-96 | 5.049789717 | 0.51 | 0.034 | 9.93E-93 | 6 | Dcn Erbb4 |
| Myoc Fzd4 | 8.99E-96 | 10.15278531 | 0.412 | 0.002 | 1.52E-92 | 6 | Myoc Fzd4 |
| Mmp2 Sdc2 | 1.26E-95 | 3.60947457 | 0.614 | 0.079 | 2.13E-92 | 6 | Mmp2 Sdc2 |
| Calr Itga2b | 2.53E-95 | 3.11779065 | 0.752 | 0.179 | 4.27E-92 | 6 | Calr Itga2b |
| Thbs1 Itga3 | 4.73E-95 | 4.761795625 | 0.461 | 0.017 | 7.99E-92 | 6 | Thbs1 Itga3 |
| Vtn Itgb3 | 1.63E-93 | 13.06085203 | 0.395 | 0 | 2.76E-90 | 6 | Vtn Itgb3 |
| Fgf2 Fgfr1 | 3.95E-93 | 2.588349889 | 0.804 | 0.198 | 6.66E-90 | 6 | Fgf2 Fgfr1 |
| Tgm2 Tbxa2r | 6.34E-93 | 3.383332034 | 0.516 | 0.034 | 1.07E-89 | 6 | Tgm2 Tbxa2r |
| Serpinc1 Lrp1 | 8.53E-93 | 2.633953045 | 0.673 | 0.107 | 1.44E-89 | 6 | Serpinc1 Lrp1 |
| Fn1 Itgb8 | 9.19E-93 | 5.865869793 | 0.435 | 0.011 | 1.55E-89 | 6 | Fn1 Itgb8 |
| Nrg1 Gpc1 | 1.45E-91 | 8.980170865 | 0.395 | 0.002 | 2.44E-88 | 6 | Nrg1 Gpc1 |
| Lrpap1 Sort1 | 3.12E-91 | 3.678242248 | 0.559 | 0.059 | 5.26E-88 | 6 | Lrpap1 Sort1 |
| Lamc3 Itga3 | 1.83E-90 | 7.803476844 | 0.392 | 0.002 | 3.10E-87 | 6 | Lamc3 Itga3 |
| Fn1 Itga3 | 8.10E-90 | 4.588329272 | 0.523 | 0.051 | 1.37E-86 | 6 | Fn1 Itga3 |
| Gnai2 Ednra | 1.57E-89 | 4.015015478 | 0.467 | 0.022 | 2.65E-86 | 6 | Gnai2 Ednra |
| Col1a2 Itga2b | 1.67E-89 | 2.996367789 | 0.742 | 0.163 | 2.82E-86 | 6 | Col1a2 Itga2b |
| Gnai2 Ptpru | 3.92E-89 | 3.270787007 | 0.536 | 0.046 | 6.62E-86 | 6 | Gnai2 Ptpru |
| Tgfb2 Acvr1 | 6.39E-89 | 2.552022244 | 0.706 | 0.142 | 1.08E-85 | 6 | Tgfb2 Acvr1 |
| Calm2 Mylk | 1.91E-88 | 3.133781402 | 0.627 | 0.101 | 3.22E-85 | 6 | Calm2 Mylk |
| Tbc1d24 Cdon | 2.81E-87 | 2.911528866 | 0.595 | 0.071 | 4.74E-84 | 6 | Tbc1d24 Cdon |
| Cdh1 Igf1r | 7.33E-87 | 4.213544351 | 0.451 | 0.02 | 1.24E-83 | 6 | Cdh1 Igf1r |
| Ndp Fzd4 | 1.59E-86 | 6.542615278 | 0.386 | 0.004 | 2.68E-83 | 6 | Ndp Fzd4 |
| Timp2 Itga3 | 2.93E-86 | 3.87603562 | 0.529 | 0.058 | 4.94E-83 | 6 | Timp2 Itga3 |
| Calr Itga3 | 5.99E-86 | 4.054290305 | 0.526 | 0.058 | 1.01E-82 | 6 | Calr Itga3 |
| Bmp2 Eng | 1.13E-85 | 5.153419232 | 0.441 | 0.023 | 1.90E-82 | 6 | Bmp2 Eng |
| Itgb3bp Itgb5 | 3.75E-85 | 2.090816143 | 0.801 | 0.2 | 6.33E-82 | 6 | Itgb3bp Itgb5 |
| Vtn Itga8 | 4.62E-85 | 9.296252444 | 0.366 | 0.001 | 7.80E-82 | 6 | Vtn Itga8 |
| C4b Cd46 | 6.03E-85 | 3.382837665 | 0.477 | 0.03 | 1.02E-81 | 6 | C4b Cd46 |
| Has2 Cd44 | 1.41E-84 | 3.898640403 | 0.451 | 0.023 | 2.38E-81 | 6 | Has2 Cd44 |
| Pros1 Axl | 1.61E-84 | 2.086246838 | 0.768 | 0.153 | 2.71E-81 | 6 | Pros1 Axl |
| Gas6 Tyro3 | 2.78E-84 | 5.344811238 | 0.425 | 0.02 | 4.69E-81 | 6 | Gas6 Tyro3 |
| Tgfb1 Sdc2 | 4.58E-83 | 3.612556297 | 0.536 | 0.067 | 7.72E-80 | 6 | Tgfb1 Sdc2 |
| Rgma Bmpr2 | 1.95E-82 | 2.058290797 | 0.866 | 0.451 | 3.29E-79 | 6 | Rgma Bmpr2 |
| Fgf9 Fgfr2 | 2.35E-82 | 4.412693308 | 0.425 | 0.019 | 3.97E-79 | 6 | Fgf9 Fgfr2 |
| Thbs1 Sdc4 | 3.57E-82 | 2.585400803 | 0.621 | 0.095 | 6.03E-79 | 6 | Thbs1 Sdc4 |
| Col1a2 Itgb3 | 5.63E-82 | 2.608892049 | 0.667 | 0.125 | 9.51E-79 | 6 | Col1a2 Itgb3 |
| Shank1 Abca1 | 6.54E-82 | 2.808877359 | 0.542 | 0.057 | 1.10E-78 | 6 | Shank1 Abca1 |
| Psap Sort1 | 2.66E-81 | 3.207005274 | 0.552 | 0.07 | 4.50E-78 | 6 | Psap Sort1 |
| Tgfa Erbb4 | 4.13E-81 | 3.438609777 | 0.526 | 0.061 | 6.97E-78 | 6 | Tgfa Erbb4 |

| | | | | | | | |
|---|---|---|---|---|---|---|---|
| Pdgfd Pdgfrb | 1.48E-80 | 3.079926096 | 0.513 | 0.052 | 2.50E-77 | 6 | Pdgfd Pdgfrb |
| Vtn Itga5 | 1.05E-79 | 7.461019705 | 0.353 | 0.003 | 1.77E-76 | 6 | Vtn Itga5 |
| Cntn2 Cntnap2 | 1.23E-79 | 7.026881926 | 0.35 | 0.002 | 2.08E-76 | 6 | Cntn2 Cntnap2 |
| Thbs1 Lrp1 | 1.76E-79 | 2.062656914 | 0.797 | 0.204 | 2.98E-76 | 6 | Thbs1 Lrp1 |
| Sorbs1 Itga1 | 3.42E-79 | 3.062989285 | 0.598 | 0.11 | 5.78E-76 | 6 | Sorbs1 Itga1 |
| Cdh1 Egfr | 7.68E-79 | 3.849475934 | 0.415 | 0.019 | 1.30E-75 | 6 | Cdh1 Egfr |
| Inhba Bambi | 8.15E-79 | 4.966454896 | 0.379 | 0.01 | 1.38E-75 | 6 | Inhba Bambi |
| Pdgfa Pdgfra | 2.22E-77 | 5.292885307 | 0.402 | 0.021 | 3.75E-74 | 6 | Pdgfa Pdgfra |
| Efnb1 Erbb2 | 2.53E-77 | 5.147613133 | 0.399 | 0.02 | 4.27E-74 | 6 | Efnb1 Erbb2 |
| Col2a1 Itga10 | 3.83E-77 | 14.24970544 | 0.33 | 0 | 6.46E-74 | 6 | Col2a1 Itga10 |
| Gas6 Axl | 1.54E-76 | 2.856079727 | 0.598 | 0.101 | 2.59E-73 | 6 | Gas6 Axl |
| Gdf11 Acvr2a | 2.80E-76 | 3.274866906 | 0.477 | 0.044 | 4.72E-73 | 6 | Gdf11 Acvr2a |
| Hbegf Cd44 | 1.54E-75 | 3.014888646 | 0.546 | 0.078 | 2.61E-72 | 6 | Hbegf Cd44 |
| Col7a1 Itgb1 | 3.73E-75 | 3.463300053 | 0.487 | 0.051 | 6.30E-72 | 6 | Col7a1 Itgb1 |
| Cdh1 Ptprf | 9.88E-75 | 12.0136408 | 0.32 | 0 | 1.67E-71 | 6 | Cdh1 Ptprf |
| Cntn2 Nrp1 | 3.79E-74 | 4.449190177 | 0.337 | 0.004 | 6.40E-71 | 6 | Cntn2 Nrp1 |
| Col5a2 Ddr1 | 7.76E-74 | 2.095651223 | 0.533 | 0.06 | 1.31E-70 | 6 | Col5a2 Ddr1 |
| Serpinc1 Gpc1 | 6.19E-73 | 9.863366785 | 0.317 | 0.001 | 1.05E-69 | 6 | Serpinc1 Gpc1 |
| Col1a2 Itga11 | 6.79E-73 | 4.837917268 | 0.353 | 0.01 | 1.15E-69 | 6 | Col1a2 Itga11 |
| Wnt4 Fzd2 | 1.80E-72 | 5.589286856 | 0.32 | 0.002 | 3.03E-69 | 6 | Wnt4 Fzd2 |
| Cdh1 Ptprm | 2.91E-72 | 4.667618555 | 0.34 | 0.006 | 4.91E-69 | 6 | Cdh1 Ptprm |
| Vtn Itga3 | 4.03E-72 | 9.695660411 | 0.314 | 0.001 | 6.81E-69 | 6 | Vtn Itga3 |
| Efemp2 Plscr4 | 5.81E-72 | 4.053139493 | 0.454 | 0.046 | 9.81E-69 | 6 | Efemp2 Plscr4 |
| F2 F2r | 7.68E-72 | 5.546095211 | 0.346 | 0.009 | 1.30E-68 | 6 | F2 F2r |
| Lif Il6st | 1.10E-71 | 4.521392194 | 0.366 | 0.014 | 1.86E-68 | 6 | Lif Il6st |
| Angptl3 Itgav | 2.30E-71 | 5.825404179 | 0.343 | 0.009 | 3.88E-68 | 6 | Angptl3 Itgav |
| Vtn Pvr | 3.45E-71 | 6.566907579 | 0.324 | 0.004 | 5.83E-68 | 6 | Vtn Pvr |
| Col3a1 Ddr2 | 5.58E-71 | 2.666593094 | 0.595 | 0.111 | 9.42E-68 | 6 | Col3a1 Ddr2 |
| Mfge8 Itgb3 | 4.91E-70 | 2.186679452 | 0.686 | 0.179 | 8.30E-67 | 6 | Mfge8 Itgb3 |
| Mfge8 Itgav | 9.76E-70 | 1.367211771 | 0.941 | 0.496 | 1.65E-66 | 6 | Mfge8 Itgav |
| Fn1 Itga9 | 9.96E-70 | 1.928208558 | 0.807 | 0.384 | 1.68E-66 | 6 | Fn1 Itga9 |
| Hras Sdc2 | 1.98E-69 | 3.350777851 | 0.51 | 0.081 | 3.34E-66 | 6 | Hras Sdc2 |
| 1700013F07Rik Plscr4 | 1.10E-68 | 4.896358388 | 0.376 | 0.023 | 1.86E-65 | 6 | 1700013F07Rik Plscr4 |
| Fn1 Itga5 | 1.34E-68 | 2.789376565 | 0.562 | 0.111 | 2.25E-65 | 6 | Fn1 Itga5 |
| Fgf9 Fgfr1 | 2.12E-68 | 3.094042135 | 0.451 | 0.043 | 3.58E-65 | 6 | Fgf9 Fgfr1 |
| Calm2 Mylk2 | 2.90E-68 | 6.992664467 | 0.307 | 0.003 | 4.90E-65 | 6 | Calm2 Mylk2 |
| Thbs1 Itgb1 | 4.36E-68 | 1.537896267 | 0.83 | 0.243 | 7.35E-65 | 6 | Thbs1 Itgb1 |
| Dusp18 Cd151 | 2.62E-66 | 3.608254997 | 0.49 | 0.082 | 4.42E-63 | 6 | Dusp18 Cd151 |
| Adam2 Cd9 | 2.93E-66 | 2.049035201 | 0.605 | 0.112 | 4.95E-63 | 6 | Adam2 Cd9 |
| Fbn1 Itga5 | 3.65E-66 | 2.495454638 | 0.575 | 0.115 | 6.17E-63 | 6 | Fbn1 Itga5 |
| Fgf22 Fgfr2 | 1.15E-65 | 6.158442344 | 0.297 | 0.003 | 1.95E-62 | 6 | Fgf22 Fgfr2 |
| Sema3e Nrp1 | 1.65E-65 | 3.177651604 | 0.35 | 0.016 | 2.79E-62 | 6 | Sema3e Nrp1 |
| Apod Lepr | 3.41E-65 | 11.97095545 | 0.281 | 0 | 5.76E-62 | 6 | Apod Lepr |
| Penk Ogfr | 5.11E-65 | 3.267255487 | 0.337 | 0.012 | 8.63E-62 | 6 | Penk Ogfr |
| Nrg3 Erbb4 | 6.10E-65 | 9.17606109 | 0.284 | 0.001 | 1.03E-61 | 6 | Nrg3 Erbb4 |
| Lama3 Sdc2 | 1.24E-64 | 2.253978243 | 0.526 | 0.082 | 2.09E-61 | 6 | Lama3 Sdc2 |
| Fat4 Dchs1 | 2.97E-64 | 2.61572853 | 0.608 | 0.135 | 5.02E-61 | 6 | Fat4 Dchs1 |
| Fn1 Itgb1 | 4.36E-64 | 1.11911787 | 0.951 | 0.628 | 7.36E-61 | 6 | Fn1 Itgb1 |
| Serpine1 Itgb5 | 6.71E-64 | 4.833805795 | 0.34 | 0.018 | 1.13E-60 | 6 | Serpine1 Itgb5 |
| Rtn4 Lingo1 | 9.34E-64 | 4.944527472 | 0.31 | 0.008 | 1.58E-60 | 6 | Rtn4 Lingo1 |
| Calm1 Mylk | 1.17E-63 | 2.571643893 | 0.546 | 0.104 | 1.97E-60 | 6 | Calm1 Mylk |

| | | | | | | | |
|---|---|---|---|---|---|---|---|
| Bmp2 Bmpr1b | 1.28E-63 | 13.6886104 | 0.275 | 0 | 2.15E-60 | 6 | Bmp2 Bmpr1b |
| Col1a2 Itgav | 3.29E-63 | 1.451589252 | 0.912 | 0.422 | 5.56E-60 | 6 | Col1a2 Itgav |
| Fgf1 Fgfr2 | 4.10E-63 | 4.051717227 | 0.337 | 0.016 | 6.93E-60 | 6 | Fgf1 Fgfr2 |
| Dusp18 Itga7 | 1.60E-62 | 7.443995586 | 0.294 | 0.006 | 2.71E-59 | 6 | Dusp18 Itga7 |
| Col1a1 Ddr2 | 2.94E-62 | 2.851987568 | 0.513 | 0.092 | 4.97E-59 | 6 | Col1a1 Ddr2 |
| Vtn Itgb8 | 2.86E-61 | 12.19819006 | 0.265 | 0 | 4.82E-58 | 6 | Vtn Itgb8 |
| Col4a1 Itgb8 | 2.93E-61 | 5.032274991 | 0.317 | 0.013 | 4.94E-58 | 6 | Col4a1 Itgb8 |
| Tgfb1 Tgfbr3 | 4.01E-61 | 2.768123859 | 0.529 | 0.109 | 6.77E-58 | 6 | Tgfb1 Tgfbr3 |
| Pon2 Htr2a | 2.56E-60 | 3.393767746 | 0.324 | 0.016 | 4.32E-57 | 6 | Pon2 Htr2a |
| Cdh1 Lrp5 | 2.67E-60 | 3.675072644 | 0.33 | 0.017 | 4.50E-57 | 6 | Cdh1 Lrp5 |
| Dcn Egfr | 3.52E-60 | 1.757021426 | 0.81 | 0.273 | 5.93E-57 | 6 | Dcn Egfr |
| Efemp2 Lingo1 | 4.99E-60 | 7.186675109 | 0.265 | 0.001 | 8.42E-57 | 6 | Efemp2 Lingo1 |
| Dusp18 Itga3 | 9.30E-60 | 6.395774671 | 0.304 | 0.012 | 1.57E-56 | 6 | Dusp18 Itga3 |
| Tfpi F3 | 1.01E-59 | 3.905768255 | 0.307 | 0.011 | 1.71E-56 | 6 | Tfpi F3 |
| Adam10 Epha3 | 1.15E-58 | 2.652401309 | 0.359 | 0.026 | 1.94E-55 | 6 | Adam10 Epha3 |
| BC055324 Plscr4 | 1.80E-58 | 2.474929886 | 0.438 | 0.056 | 3.03E-55 | 6 | BC055324 Plscr4 |
| Calm1 Mylk2 | 1.99E-58 | 6.06786965 | 0.271 | 0.004 | 3.35E-55 | 6 | Calm1 Mylk2 |
| Adam17 Erbb4 | 2.01E-57 | 2.414827681 | 0.471 | 0.074 | 3.40E-54 | 6 | Adam17 Erbb4 |
| Bmp5 Bmpr1b | 2.23E-57 | 12.52491871 | 0.248 | 0 | 3.77E-54 | 6 | Bmp5 Bmpr1b |
| Angptl3 Itgb3 | 2.23E-57 | 11.76920372 | 0.248 | 0 | 3.77E-54 | 6 | Angptl3 Itgb3 |
| Sftpd Tlr4 | 2.59E-57 | 1.419054548 | 0.791 | 0.222 | 4.36E-54 | 6 | Sftpd Tlr4 |
| Tgfb2 Tgfbr2 | 2.84E-57 | 1.41196241 | 0.788 | 0.214 | 4.80E-54 | 6 | Tgfb2 Tgfbr2 |
| Adam10 Axl1 | 6.41E-57 | 0.953340328 | 0.761 | 0.17 | 1.08E-53 | 6 | Adam10 Axl |
| Tgfb3 Eng | 1.19E-56 | 2.860811792 | 0.379 | 0.038 | 2.00E-53 | 6 | Tgfb3 Eng |
| Gdf9 Bmpr1a | 2.75E-56 | 3.516353166 | 0.297 | 0.013 | 4.64E-53 | 6 | Gdf9 Bmpr1a |
| Efna1 Ephb6 | 5.95E-56 | 2.890133333 | 0.31 | 0.016 | 1.00E-52 | 6 | Efna1 Ephb6 |
| Col3a1 Ddr1 | 1.66E-55 | 2.54329996 | 0.422 | 0.059 | 2.80E-52 | 6 | Col3a1 Ddr1 |
| Ncam1 Robo1 | 4.75E-55 | 3.011750424 | 0.353 | 0.031 | 8.02E-52 | 6 | Ncam1 Robo1 |
| Col1a2 Itga1 | 3.52E-54 | 1.992556 | 0.578 | 0.138 | 5.94E-51 | 6 | Col1a2 Itga1 |
| Tgfb1 Itgb8 | 3.65E-54 | 4.630657433 | 0.291 | 0.014 | 6.16E-51 | 6 | Tgfb1 Itgb8 |
| Mdk Lrp1 | 8.48E-54 | 2.645734395 | 0.5 | 0.102 | 1.43E-50 | 6 | Mdk Lrp1 |
| Mdk Itgb1 | 1.29E-53 | 2.282864176 | 0.526 | 0.115 | 2.18E-50 | 6 | Mdk Itgb1 |
| Angptl1 Tek | 3.23E-53 | 6.518360998 | 0.248 | 0.004 | 5.44E-50 | 6 | Angptl1 Tek |
| Fgf1 Egfr | 6.40E-53 | 3.089609378 | 0.363 | 0.037 | 1.08E-49 | 6 | Fgf1 Egfr |
| Lama2 Itga7 | 8.62E-53 | 6.205237509 | 0.258 | 0.007 | 1.46E-49 | 6 | Lama2 Itga7 |
| Col1a1 Ddr1 | 1.05E-52 | 2.743467513 | 0.379 | 0.048 | 1.77E-49 | 6 | Col1a1 Ddr1 |
| Dusp18 Itga1 | 1.06E-52 | 4.250465158 | 0.33 | 0.031 | 1.78E-49 | 6 | Dusp18 Itga1 |
| Efna1 Epha1 | 1.14E-52 | 2.813494899 | 0.297 | 0.017 | 1.93E-49 | 6 | Efna1 Epha1 |
| Lrpap1 Lrp1 | 2.71E-52 | 1.224575489 | 0.948 | 0.629 | 4.57E-49 | 6 | Lrpap1 Lrp1 |
| Fgf1 Fgfr1 | 3.86E-52 | 2.686560484 | 0.382 | 0.043 | 6.52E-49 | 6 | Fgf1 Fgfr1 |
| Mdk Sdc4 | 7.08E-52 | 2.85698279 | 0.389 | 0.052 | 1.20E-48 | 6 | Mdk Sdc4 |
| Adam9 Itga3 | 1.24E-51 | 2.937252879 | 0.389 | 0.055 | 2.09E-48 | 6 | Adam9 Itga3 |
| Bdnf Ddr1 | 2.54E-51 | 4.009829009 | 0.255 | 0.007 | 4.30E-48 | 6 | Bdnf Ddr1 |
| Ecm1 Cachd1 | 3.81E-51 | 3.482300303 | 0.324 | 0.027 | 6.43E-48 | 6 | Ecm1 Cachd1 |
| Lama2 Itga3 | 3.18E-50 | 5.558519552 | 0.271 | 0.014 | 5.36E-47 | 6 | Lama2 Itga3 |
| Bmp7 Bmpr1a | 4.58E-50 | 5.477732722 | 0.232 | 0.003 | 7.74E-47 | 6 | Bmp7 Bmpr1a |
| Lama4 Itga3 | 6.21E-50 | 3.731252625 | 0.346 | 0.041 | 1.05E-46 | 6 | Lama4 Itga3 |
| Hsp90b1 Lrp1 | 7.16E-50 | 1.167955033 | 0.941 | 0.659 | 1.21E-46 | 6 | Hsp90b1 Lrp1 |
| Efna5 Ephb2 | 8.77E-50 | 3.808858116 | 0.281 | 0.016 | 1.48E-46 | 6 | Efna5 Ephb2 |
| Serpine1 Itgav | 1.61E-49 | 3.142910415 | 0.353 | 0.042 | 2.72E-46 | 6 | Serpine1 Itgav |
| Ptgs2 Cav1 | 1.86E-48 | 2.859047127 | 0.324 | 0.032 | 3.14E-45 | 6 | Ptgs2 Cav1 |

| | | | | | | | |
|---|---|---|---|---|---|---|---|
| Lama2 Itga1 | 9.42E-48 | 4.050285323 | 0.31 | 0.031 | 1.59E-44 | 6 | Lama2 Itga1 |
| Thbs1 Cd47 | 1.25E-47 | 0.995237102 | 0.804 | 0.253 | 2.11E-44 | 6 | Thbs1 Cd47 |
| Vegfa Gpc1 | 1.34E-47 | 2.566604184 | 0.239 | 0.007 | 2.27E-44 | 6 | Vegfa Gpc1 |
| Fgf1 Fgfrl1 | 2.16E-47 | 10.41899717 | 0.206 | 0 | 3.64E-44 | 6 | Fgf1 Fgfrl1 |
| Hspg2 Ptprs | 2.24E-47 | 1.147309392 | 0.918 | 0.442 | 3.78E-44 | 6 | Hspg2 Ptprs |
| Ngf Maged1 | 2.91E-47 | 3.825387009 | 0.301 | 0.028 | 4.91E-44 | 6 | Ngf Maged1 |
| Inhba Acvr1 | 4.76E-47 | 1.409018264 | 0.533 | 0.125 | 8.03E-44 | 6 | Inhba Acvr1 |
| Gnb3 Tgfbr1 | 1.30E-46 | 2.536508651 | 0.245 | 0.009 | 2.19E-43 | 6 | Gnb3 Tgfbr1 |
| Thbs1 Lrp5 | 1.31E-46 | 1.630259481 | 0.569 | 0.148 | 2.21E-43 | 6 | Thbs1 Lrp5 |
| Vegfa Tyro3 | 2.17E-46 | 3.250819098 | 0.294 | 0.026 | 3.66E-43 | 6 | Vegfa Tyro3 |
| Efna1 Epha3 | 2.63E-46 | 2.902202204 | 0.288 | 0.022 | 4.43E-43 | 6 | Efna1 Epha3 |
| Efna5 Epha1 | 1.03E-45 | 2.505914102 | 0.268 | 0.017 | 1.74E-42 | 6 | Efna5 Epha1 |
| Wnt3a Fzd6 | 1.35E-45 | 5.370472141 | 0.235 | 0.009 | 2.28E-42 | 6 | Wnt3a Fzd6 |
| Rgma Bmpr1b | 2.13E-45 | 5.262529161 | 0.248 | 0.013 | 3.60E-42 | 6 | Rgma Bmpr1b |
| Fbn1 Itgb1 | 4.27E-45 | 0.909506897 | 0.984 | 0.675 | 7.21E-42 | 6 | Fbn1 Itgb1 |
| Angptl3 Itga5 | 6.96E-45 | 6.156949171 | 0.209 | 0.003 | 1.17E-41 | 6 | Angptl3 Itga5 |
| Bmp7 Bmpr2 | 1.26E-44 | 4.026303916 | 0.239 | 0.01 | 2.13E-41 | 6 | Bmp7 Bmpr2 |
| Spp1 Itga4 | 1.49E-44 | 3.251069865 | 0.235 | 0.009 | 2.51E-41 | 6 | Spp1 Itga4 |
| Rtn4 Rtn4r | 1.54E-44 | 5.826812081 | 0.212 | 0.004 | 2.59E-41 | 6 | Rtn4 Rtn4r |
| Apoe Vldlr | 1.60E-44 | 2.647336881 | 0.33 | 0.04 | 2.69E-41 | 6 | Apoe Vldlr |
| Il16 Kcnj4 | 2.29E-44 | 5.445796245 | 0.216 | 0.005 | 3.87E-41 | 6 | Il16 Kcnj4 |
| Mdk Ptprz1 | 4.54E-44 | 5.143857696 | 0.219 | 0.006 | 7.66E-41 | 6 | Mdk Ptprz1 |
| Lamc1 Itga7 | 4.60E-44 | 5.181337432 | 0.248 | 0.016 | 7.76E-41 | 6 | Lamc1 Itga7 |
| F2 Thbd | 6.00E-44 | 3.984120309 | 0.258 | 0.018 | 1.01E-40 | 6 | F2 Thbd |
| Eda Eda2r | 6.65E-44 | 4.014484874 | 0.255 | 0.017 | 1.12E-40 | 6 | Eda Eda2r |
| Efna4 Epha3 | 1.25E-43 | 6.047067154 | 0.216 | 0.006 | 2.11E-40 | 6 | Efna4 Epha3 |
| Bmp7 Acvr1 | 1.72E-43 | 4.291488849 | 0.222 | 0.007 | 2.91E-40 | 6 | Bmp7 Acvr1 |
| Ncam1 Fgfr2 | 2.13E-43 | 1.386035883 | 0.768 | 0.291 | 3.60E-40 | 6 | Ncam1 Fgfr2 |
| Gdf9 Bmpr2 | 2.73E-43 | 2.909786643 | 0.307 | 0.036 | 4.62E-40 | 6 | Gdf9 Bmpr2 |
| Serpine1 Lrp1 | 3.39E-43 | 2.602711597 | 0.356 | 0.053 | 5.73E-40 | 6 | Serpine1 Lrp1 |
| Efnb1 Ephb3 | 3.45E-43 | 4.678431864 | 0.219 | 0.007 | 5.83E-40 | 6 | Efnb1 Ephb3 |
| Adam9 Itgb5 | 3.86E-43 | 1.432750849 | 0.644 | 0.214 | 6.52E-40 | 6 | Adam9 Itgb5 |
| Ctf1 Lifr | 4.46E-43 | 1.667978355 | 0.408 | 0.069 | 7.53E-40 | 6 | Ctf1 Lifr |
| Calr Itgav | 5.40E-43 | 0.921631896 | 0.925 | 0.552 | 9.12E-40 | 6 | Calr Itgav |
| Gnas Htr6 | 7.53E-43 | 11.74685266 | 0.186 | 0 | 1.27E-39 | 6 | Gnas Htr6 |
| Lama2 Itgb1 | 6.32E-42 | 2.115407042 | 0.513 | 0.144 | 1.07E-38 | 6 | Lama2 Itgb1 |
| Calm1 Adcyap1r1 | 6.60E-42 | 4.448643561 | 0.222 | 0.009 | 1.11E-38 | 6 | Calm1 Adcyap1r1 |
| Efnb1 Ephb6 | 1.06E-41 | 4.884759297 | 0.196 | 0.003 | 1.79E-38 | 6 | Efnb1 Ephb6 |
| Itgb3bp Itgb3 | 1.39E-41 | 1.440522531 | 0.601 | 0.178 | 2.34E-38 | 6 | Itgb3bp Itgb3 |
| Efna4 Epha1 | 2.01E-41 | 7.145545568 | 0.19 | 0.002 | 3.40E-38 | 6 | Efna4 Epha1 |
| Dlk1 Notch3 | 2.41E-41 | 11.76246226 | 0.18 | 0 | 4.07E-38 | 6 | Dlk1 Notch3 |
| Bmp7 Acvr2b | 4.15E-41 | 3.716987685 | 0.216 | 0.008 | 7.01E-38 | 6 | Bmp7 Acvr2b |
| Lamc2 Cd151 | 4.16E-41 | 4.47995726 | 0.239 | 0.017 | 7.02E-38 | 6 | Lamc2 Cd151 |
| Efnb1 Ephb2 | 4.47E-41 | 4.635200778 | 0.193 | 0.003 | 7.54E-38 | 6 | Efnb1 Ephb2 |
| Efnb3 Epha4 | 5.22E-41 | 4.920865216 | 0.206 | 0.006 | 8.82E-38 | 6 | Efnb3 Epha4 |
| Calm1 Fas | 5.88E-41 | 1.126390715 | 0.755 | 0.314 | 9.93E-38 | 6 | Calm1 Fas |
| Efna5 Epha3 | 1.63E-40 | 2.828218878 | 0.258 | 0.021 | 2.76E-37 | 6 | Efna5 Epha3 |
| Col1a1 Itga11 | 4.13E-40 | 4.20643412 | 0.212 | 0.009 | 6.97E-37 | 6 | Col1a1 Itga11 |
| Lif Lifr | 4.32E-40 | 2.761580101 | 0.235 | 0.014 | 7.29E-37 | 6 | Lif Lifr |
| Bdnf Sort1 | 1.42E-39 | 5.530331298 | 0.186 | 0.003 | 2.40E-36 | 6 | Bdnf Sort1 |
| Nppa Npr1 | 2.06E-39 | 3.780190225 | 0.219 | 0.011 | 3.47E-36 | 6 | Nppa Npr1 |

| | | | | | | | |
|---|---|---|---|---|---|---|---|
| Agrn Musk | 4.91E-39 | 3.310358558 | 0.242 | 0.019 | 8.28E-36 | 6 | Agrn Musk |
| Inhba Acvr1b | 7.08E-39 | 1.339650027 | 0.556 | 0.165 | 1.19E-35 | 6 | Inhba Acvr1b |
| Inhbb Acvr1 | 1.22E-38 | 2.040418249 | 0.454 | 0.124 | 2.06E-35 | 6 | Inhbb Acvr1 |
| Dusp18 Itgb1 | 1.40E-38 | 1.897705392 | 0.549 | 0.202 | 2.37E-35 | 6 | Dusp18 Itgb1 |
| Gdf9 Tgfbr1 | 1.77E-38 | 2.524907321 | 0.291 | 0.036 | 2.99E-35 | 6 | Gdf9 Tgfbr1 |
| Npy Fap | 5.13E-38 | 8.436892233 | 0.17 | 0.001 | 8.65E-35 | 6 | Npy Fap |
| Thbs1 Tnfrsf11b | 5.74E-38 | 7.415664508 | 0.17 | 0.001 | 9.70E-35 | 6 | Thbs1 Tnfrsf11b |
| Il16 Kcnj15 | 1.59E-37 | 7.233696903 | 0.176 | 0.003 | 2.69E-34 | 6 | Il16 Kcnj15 |
| Ngf Kidins220 | 4.11E-37 | 2.072509069 | 0.33 | 0.052 | 6.94E-34 | 6 | Ngf Kidins220 |
| Npy Npy1r | 7.45E-37 | 11.53388541 | 0.16 | 0 | 1.26E-33 | 6 | Npy Npy1r |
| Dll3 Notch3 | 7.45E-37 | 11.38777394 | 0.16 | 0 | 1.26E-33 | 6 | Dll3 Notch3 |
| Il16 Kcnd1 | 1.47E-36 | 1.972097787 | 0.33 | 0.054 | 2.49E-33 | 6 | Il16 Kcnd1 |
| Calr Lrp1 | 5.87E-36 | 1.041166715 | 0.922 | 0.637 | 9.91E-33 | 6 | Calr Lrp1 |
| Efnb3 Ephb3 | 1.01E-35 | 6.412439544 | 0.173 | 0.004 | 1.71E-32 | 6 | Efnb3 Ephb3 |
| Efnb3 Ephb4 | 1.04E-35 | 4.766194105 | 0.186 | 0.007 | 1.76E-32 | 6 | Efnb3 Ephb4 |
| Fn1 Itgb6 | 1.15E-35 | 5.908687392 | 0.16 | 0.001 | 1.95E-32 | 6 | Fn1 Itgb6 |
| Ntf3 Ntrk3 | 1.27E-35 | 5.961576583 | 0.173 | 0.004 | 2.14E-32 | 6 | Ntf3 Ntrk3 |
| Lpl Vldlr | 1.87E-35 | 4.476929296 | 0.199 | 0.011 | 3.15E-32 | 6 | Lpl Vldlr |
| Fbn1 Itgb6 | 4.60E-35 | 5.50569528 | 0.167 | 0.003 | 7.76E-32 | 6 | Fbn1 Itgb6 |
| Npnt Itga8 | 9.88E-35 | 3.925212931 | 0.222 | 0.02 | 1.67E-31 | 6 | Npnt Itga8 |
| Farp2 Plxna3 | 1.20E-34 | 3.020184198 | 0.533 | 0.26 | 2.03E-31 | 6 | Farp2 Plxna3 |
| Efna5 Ephb6 | 1.49E-34 | 1.498675617 | 0.222 | 0.018 | 2.51E-31 | 6 | Efna5 Ephb6 |
| Efnb1 Epha4 | 1.60E-34 | 1.32576887 | 0.268 | 0.032 | 2.71E-31 | 6 | Efnb1 Epha4 |
| Efnb1 Ephb4 | 2.28E-34 | 1.426925486 | 0.258 | 0.029 | 3.84E-31 | 6 | Efnb1 Ephb4 |
| Efnb3 Ephb2 | 2.79E-34 | 8.150238649 | 0.154 | 0.001 | 4.70E-31 | 6 | Efnb3 Ephb2 |
| Calm2 Adcy8 | 3.76E-34 | 3.516443982 | 0.176 | 0.006 | 6.34E-31 | 6 | Calm2 Adcy8 |
| Fgf2 Cd44 | 4.75E-34 | 1.479117248 | 0.5 | 0.153 | 8.02E-31 | 6 | Fgf2 Cd44 |
| Inhba Acvr2b1 | 5.14E-34 | 0.92031174 | 0.556 | 0.158 | 8.68E-31 | 6 | Inhba Acvr2b |
| Sema3b Nrp1 | 6.23E-34 | 1.76388252 | 0.288 | 0.041 | 1.05E-30 | 6 | Sema3b Nrp1 |
| Efnb3 Ephb6 | 6.99E-34 | 9.493987982 | 0.147 | 0 | 1.18E-30 | 6 | Efnb3 Ephb6 |
| Farp2 Plxna1 | 7.46E-34 | 1.434406654 | 0.712 | 0.431 | 1.26E-30 | 6 | Farp2 Plxna1 |
| Col4a1 Itga1 | 1.28E-33 | 1.721588015 | 0.415 | 0.111 | 2.16E-30 | 6 | Col4a1 Itga1 |
| Gnas Adcy1 | 1.64E-33 | 5.144410349 | 0.16 | 0.003 | 2.77E-30 | 6 | Gnas Adcy1 |
| Fn1 Tnfrsf11b | 1.79E-33 | 4.712553427 | 0.18 | 0.008 | 3.01E-30 | 6 | Fn1 Tnfrsf11b |
| Fgf14 Fgfr2 | 2.44E-33 | 1.124976361 | 0.536 | 0.169 | 4.12E-30 | 6 | Fgf14 Fgfr2 |
| Gnas Adcy8 | 3.42E-33 | 3.56778224 | 0.176 | 0.007 | 5.77E-30 | 6 | Gnas Adcy8 |
| Psen1 Notch3 | 3.53E-33 | 2.60688832 | 0.209 | 0.017 | 5.96E-30 | 6 | Psen1 Notch3 |
| Nrg4 Erbb2 | 3.84E-33 | 10.43266938 | 0.144 | 0 | 6.49E-30 | 6 | Nrg4 Erbb2 |
| Lamc2 Itga3 | 4.59E-33 | 7.474530011 | 0.157 | 0.003 | 7.75E-30 | 6 | Lamc2 Itga3 |
| Vegfa Ephb2 | 6.33E-33 | 2.877591941 | 0.212 | 0.018 | 1.07E-29 | 6 | Vegfa Ephb2 |
| Inhbb Acvr1b | 6.85E-33 | 1.804920153 | 0.464 | 0.148 | 1.16E-29 | 6 | Inhbb Acvr1b |
| Tgfb2 Eng | 7.23E-33 | 1.789707173 | 0.373 | 0.084 | 1.22E-29 | 6 | Tgfb2 Eng |
| Lamc2 Itgb1 | 8.14E-33 | 2.575929521 | 0.261 | 0.036 | 1.37E-29 | 6 | Lamc2 Itgb1 |
| Gnai2 Adcy1 | 8.80E-33 | 5.12690549 | 0.157 | 0.003 | 1.49E-29 | 6 | Gnai2 Adcy1 |
| Lamc1 Itga3 | 1.37E-32 | 3.06356047 | 0.258 | 0.038 | 2.30E-29 | 6 | Lamc1 Itga3 |
| Gnai2 Adcy8 | 2.04E-32 | 2.890629587 | 0.173 | 0.007 | 3.44E-29 | 6 | Gnai2 Adcy8 |
| Lin7c Kcnj4 | 2.66E-32 | 2.5923986 | 0.219 | 0.021 | 4.49E-29 | 6 | Lin7c Kcnj4 |
| Col2a1 Itga2 | 1.15E-31 | 13.12133539 | 0.137 | 0 | 1.95E-28 | 6 | Col2a1 Itga2 |
| Col11a1 Itga2 | 1.15E-31 | 13.00726094 | 0.137 | 0 | 1.95E-28 | 6 | Col11a1 Itga2 |
| Chad Itga2 | 1.15E-31 | 12.90676592 | 0.137 | 0 | 1.95E-28 | 6 | Chad Itga2 |
| Fgf1 Cd44 | 1.83E-31 | 1.593212423 | 0.252 | 0.032 | 3.09E-28 | 6 | Fgf1 Cd44 |

| | | | | | | | |
|---|---|---|---|---|---|---|---|
| Gnai2 F2r | 2.41E-31 | 1.391776466 | 0.467 | 0.135 | 4.08E-28 | 6 | Gnai2 F2r |
| Efna4 Epha6 | 2.57E-31 | 2.728718433 | 0.193 | 0.014 | 4.33E-28 | 6 | Efna4 Epha6 |
| Calm2 Grm7 | 3.10E-31 | 2.784411524 | 0.261 | 0.04 | 5.23E-28 | 6 | Calm2 Grm7 |
| Anxa1 Dysf | 3.87E-31 | 2.718365347 | 0.232 | 0.028 | 6.54E-28 | 6 | Anxa1 Dysf |
| Calm1 Adcy8 | 5.80E-31 | 2.946870658 | 0.167 | 0.007 | 9.79E-28 | 6 | Calm1 Adcy8 |
| Efna4 Epha4 | 6.79E-31 | 1.927360344 | 0.291 | 0.051 | 1.15E-27 | 6 | Efna4 Epha4 |
| Adipoq Adipor1 | 1.54E-30 | 2.474146312 | 0.219 | 0.023 | 2.61E-27 | 6 | Adipoq Adipor1 |
| Ngf Sort1 | 3.20E-30 | 3.637254561 | 0.167 | 0.008 | 5.41E-27 | 6 | Ngf Sort1 |
| Hhipl2 Cachd1 | 8.08E-30 | 4.620968738 | 0.144 | 0.003 | 1.36E-26 | 6 | Hhipl2 Cachd1 |
| Vegfb Tyro3 | 9.65E-30 | 4.689922558 | 0.199 | 0.021 | 1.63E-26 | 6 | Vegfb Tyro3 |
| Inha Tgfbr3 | 1.67E-29 | 2.776437774 | 0.314 | 0.073 | 2.82E-26 | 6 | Inha Tgfbr3 |
| Hsp90aa1 Egfr | 1.77E-29 | 1.017275368 | 0.676 | 0.27 | 2.99E-26 | 6 | Hsp90aa1 Egfr |
| Calr Mtnr1a | 2.14E-29 | 3.128982189 | 0.163 | 0.008 | 3.61E-26 | 6 | Calr Mtnr1a |
| Fgf2 Nrp1 | 3.77E-29 | 1.32960833 | 0.556 | 0.202 | 6.36E-26 | 6 | Fgf2 Nrp1 |
| Bmp7 Acvr2a | 5.44E-29 | 4.089260178 | 0.157 | 0.007 | 9.18E-26 | 6 | Bmp7 Acvr2a |
| Pdgfd Pdgfra | 7.69E-29 | 2.750330416 | 0.248 | 0.04 | 1.30E-25 | 6 | Pdgfd Pdgfra |
| Nrg4 Egfr | 8.99E-29 | 4.017306695 | 0.144 | 0.004 | 1.52E-25 | 6 | Nrg4 Egfr |
| Col8a1 Itga2 | 1.24E-28 | 7.125741331 | 0.137 | 0.003 | 2.09E-25 | 6 | Col8a1 Itga2 |
| Jag1 Notch3 | 1.40E-28 | 1.469096265 | 0.19 | 0.017 | 2.37E-25 | 6 | Jag1 Notch3 |
| Adam17 Itga5 | 1.63E-28 | 0.845980986 | 0.503 | 0.15 | 2.76E-25 | 6 | Adam17 Itga5 |
| App Cav1 | 2.08E-28 | 1.106744995 | 0.637 | 0.273 | 3.51E-25 | 6 | App Cav1 |
| Pla2g10 Pla2r1 | 5.47E-28 | 10.54921917 | 0.121 | 0 | 9.23E-25 | 6 | Pla2g10 Pla2r1 |
| Gnai2 Mtnr1a | 5.68E-28 | 2.948298667 | 0.157 | 0.008 | 9.59E-25 | 6 | Gnai2 Mtnr1a |
| Gdf9 Acvr2a | 2.15E-27 | 2.323226664 | 0.216 | 0.029 | 3.62E-24 | 6 | Gdf9 Acvr2a |
| Calm1 Grm3 | 3.60E-27 | 4.693331671 | 0.144 | 0.006 | 6.08E-24 | 6 | Calm1 Grm3 |
| Tgfb1 Cd109 | 4.29E-27 | 2.644355431 | 0.19 | 0.02 | 7.23E-24 | 6 | Tgfb1 Cd109 |
| Efna1 Epha7 | 4.46E-27 | 1.721896404 | 0.212 | 0.027 | 7.53E-24 | 6 | Efna1 Epha7 |
| F13a1 Itga4 | 4.96E-27 | 1.308424131 | 0.225 | 0.031 | 8.37E-24 | 6 | F13a1 Itga4 |
| Inhbb Acvr2b | 6.38E-27 | 1.360400701 | 0.448 | 0.14 | 1.08E-23 | 6 | Inhbb Acvr2b |
| Col6a3 Itga2 | 8.41E-27 | 5.827433675 | 0.137 | 0.005 | 1.42E-23 | 6 | Col6a3 Itga2 |
| Col5a2 Itga1 | 1.19E-26 | 0.907638855 | 0.461 | 0.138 | 2.01E-23 | 6 | Col5a2 Itga1 |
| Inhba Eng | 2.12E-26 | 1.196804822 | 0.294 | 0.06 | 3.58E-23 | 6 | Inhba Eng |
| Inhba Acvr2a | 2.28E-26 | 1.209772166 | 0.405 | 0.119 | 3.85E-23 | 6 | Inhba Acvr2a |
| Vtn Tnfrsf11b | 8.54E-26 | 10.41647359 | 0.111 | 0 | 1.44E-22 | 6 | Vtn Tnfrsf11b |
| Shank1 Sstr2 | 8.59E-26 | 5.627463509 | 0.121 | 0.002 | 1.45E-22 | 6 | Shank1 Sstr2 |
| Gnas Adora1 | 1.13E-25 | 3.028710932 | 0.134 | 0.005 | 1.91E-22 | 6 | Gnas Adora1 |
| Mdk Alk | 1.46E-25 | 5.143910405 | 0.124 | 0.003 | 2.47E-22 | 6 | Mdk Alk |
| Calm1 Grm7 | 1.52E-25 | 2.189410389 | 0.232 | 0.039 | 2.56E-22 | 6 | Calm1 Grm7 |
| Gnai2 S1pr3 | 1.65E-25 | 3.732940667 | 0.137 | 0.006 | 2.78E-22 | 6 | Gnai2 S1pr3 |
| Nid1 Ptprf | 2.85E-25 | 2.02162046 | 0.203 | 0.028 | 4.81E-22 | 6 | Nid1 Ptprf |
| Rgmb Neo1 | 2.93E-25 | 1.688615809 | 0.317 | 0.077 | 4.95E-22 | 6 | Rgmb Neo1 |
| Gnai2 Unc5b | 3.01E-25 | 1.802301077 | 0.307 | 0.073 | 5.09E-22 | 6 | Gnai2 Unc5b |
| Gdnf Slc44a5 | 4.58E-25 | 10.84418693 | 0.108 | 0 | 7.73E-22 | 6 | Gdnf Slc44a5 |
| Gnai2 Adora1 | 6.07E-25 | 3.014639008 | 0.131 | 0.005 | 1.02E-21 | 6 | Gnai2 Adora1 |
| Tgm2 Itgb3 | 7.61E-25 | 0.713170547 | 0.552 | 0.194 | 1.28E-21 | 6 | Tgm2 Itgb3 |
| Tgfb1 Itgb6 | 1.42E-24 | 4.748285682 | 0.124 | 0.004 | 2.40E-21 | 6 | Tgfb1 Itgb6 |
| Fgf1 Nrp1 | 1.91E-24 | 1.336842209 | 0.258 | 0.049 | 3.22E-21 | 6 | Fgf1 Nrp1 |
| Col6a2 Itga2 | 2.16E-24 | 4.957557179 | 0.134 | 0.007 | 3.64E-21 | 6 | Col6a2 Itga2 |
| Vtn Itgb6 | 2.45E-24 | 10.5947339 | 0.105 | 0 | 4.13E-21 | 6 | Vtn Itgb6 |
| Dll3 Notch1 | 7.48E-24 | 3.579358664 | 0.144 | 0.01 | 1.26E-20 | 6 | Dll3 Notch1 |
| Agrp Mc5r | 8.04E-24 | 5.404584415 | 0.124 | 0.005 | 1.36E-20 | 6 | Agrp Mc5r |

| | | | | | | | |
|---|---|---|---|---|---|---|---|
| Col18a1 Gpc4 | 1.02E-23 | 2.768988158 | 0.219 | 0.039 | 1.73E-20 | 6 | Col18a1 Gpc4 |
| Efna5 Epha7 | 1.10E-23 | 1.046763854 | 0.199 | 0.028 | 1.86E-20 | 6 | Efna5 Epha7 |
| Col6a1 Itga2 | 1.44E-23 | 4.598839332 | 0.137 | 0.009 | 2.44E-20 | 6 | Col6a1 Itga2 |
| Col18a1 Gpc1 | 2.15E-23 | 3.908790684 | 0.114 | 0.003 | 3.63E-20 | 6 | Col18a1 Gpc1 |
| Lamb3 Itga2 | 2.29E-23 | 3.707103225 | 0.127 | 0.006 | 3.87E-20 | 6 | Lamb3 Itga2 |
| Pomc Mc5r | 3.13E-23 | 5.649734211 | 0.118 | 0.004 | 5.29E-20 | 6 | Pomc Mc5r |
| Kng1 Sdc2 | 5.37E-23 | 7.984395543 | 0.108 | 0.002 | 9.07E-20 | 6 | Kng1 Sdc2 |
| Calm2 Pde1c | 1.01E-22 | 1.406251021 | 0.373 | 0.12 | 1.71E-19 | 6 | Calm2 Pde1c |
| Hspa1a Grin2d | 1.34E-22 | 3.770704146 | 0.15 | 0.014 | 2.26E-19 | 6 | Hspa1a Grin2d |
| Col1a1 Itgav | 2.52E-22 | 1.419180229 | 0.529 | 0.281 | 4.25E-19 | 6 | Col1a1 Itgav |
| Efna4 Epha7 | 5.20E-22 | 3.874276575 | 0.127 | 0.008 | 8.77E-19 | 6 | Efna4 Epha7 |
| Ccl25 Ackr2 | 5.96E-22 | 3.920984171 | 0.137 | 0.011 | 1.01E-18 | 6 | Ccl25 Ackr2 |
| Gdf9 Fxyd6 | 8.01E-22 | 8.062353507 | 0.098 | 0.001 | 1.35E-18 | 6 | Gdf9 Fxyd6 |
| Gdf9 Bmpr1b | 8.54E-22 | 6.865653144 | 0.098 | 0.001 | 1.44E-18 | 6 | Gdf9 Bmpr1b |
| Egf Erbb2 | 8.90E-22 | 3.451987757 | 0.219 | 0.047 | 1.50E-18 | 6 | Egf Erbb2 |
| Bmp7 Eng | 1.34E-21 | 3.991628819 | 0.111 | 0.004 | 2.26E-18 | 6 | Bmp7 Eng |
| Adipoq Adipor2 | 1.67E-21 | 1.917853078 | 0.17 | 0.022 | 2.82E-18 | 6 | Adipoq Adipor2 |
| Tnc Itga7 | 1.75E-21 | 5.882107987 | 0.101 | 0.002 | 2.96E-18 | 6 | Tnc Itga7 |
| Nrg4 Erbb4 | 1.96E-21 | 10.54548989 | 0.092 | 0 | 3.32E-18 | 6 | Nrg4 Erbb4 |
| Olfm2 Robo2 | 4.37E-21 | 7.583891303 | 0.095 | 0.001 | 7.38E-18 | 6 | Olfm2 Robo2 |
| Lamc3 Itga2 | 1.04E-20 | 10.38765276 | 0.088 | 0 | 1.76E-17 | 6 | Lamc3 Itga2 |
| Col1a1 Itga1 | 1.94E-20 | 1.689174661 | 0.333 | 0.113 | 3.28E-17 | 6 | Col1a1 Itga1 |
| Wnt5a Fzd2 | 2.39E-20 | 5.633108997 | 0.105 | 0.004 | 4.03E-17 | 6 | Wnt5a Fzd2 |
| Gnas Lhcgr | 2.50E-20 | 2.003463984 | 0.16 | 0.021 | 4.22E-17 | 6 | Gnas Lhcgr |
| Il18 Il1rapl1 | 4.60E-20 | 1.911662151 | 0.131 | 0.011 | 7.76E-17 | 6 | Il18 Il1rapl1 |
| Ereg Erbb2 | 5.50E-20 | 9.07562383 | 0.085 | 0 | 9.28E-17 | 6 | Ereg Erbb2 |
| Gnai2 Cav1 | 8.04E-20 | 0.755439096 | 0.601 | 0.273 | 1.36E-16 | 6 | Gnai2 Cav1 |
| Col1a2 Itga2 | 1.16E-19 | 3.304014053 | 0.127 | 0.011 | 1.96E-16 | 6 | Col1a2 Itga2 |
| Inhbb Acvr2a | 1.30E-19 | 1.619348469 | 0.346 | 0.118 | 2.20E-16 | 6 | Inhbb Acvr2a |
| Wnt4 Fzd6 | 2.87E-19 | 3.129500617 | 0.105 | 0.005 | 4.85E-16 | 6 | Wnt4 Fzd6 |
| Lama5 Itga3 | 4.00E-19 | 2.594994553 | 0.212 | 0.05 | 6.74E-16 | 6 | Lama5 Itga3 |
| Wnt5a Fzd1 | 4.57E-19 | 5.031713019 | 0.095 | 0.003 | 7.72E-16 | 6 | Wnt5a Fzd1 |
| Hspg2 Itga2 | 5.89E-19 | 2.769849648 | 0.131 | 0.013 | 9.94E-16 | 6 | Hspg2 Itga2 |
| Calm1 Calcr | 7.26E-19 | 2.693161951 | 0.124 | 0.011 | 1.23E-15 | 6 | Calm1 Calcr |
| Fn1 Itga2 | 9.23E-19 | 3.678263339 | 0.131 | 0.014 | 1.56E-15 | 6 | Fn1 Itga2 |
| Wnt5a Ror1 | 1.08E-18 | 5.814254128 | 0.105 | 0.006 | 1.82E-15 | 6 | Wnt5a Ror1 |
| Nlgn2 Nrxn2 | 1.12E-18 | 4.469108552 | 0.105 | 0.006 | 1.89E-15 | 6 | Nlgn2 Nrxn2 |
| Fgf14 Fgfr11 | 1.20E-18 | 0.62468394 | 0.624 | 0.274 | 2.03E-15 | 6 | Fgf14 Fgfr1 |
| Ncan Sdc3 | 1.90E-18 | 3.770960995 | 0.111 | 0.008 | 3.20E-15 | 6 | Ncan Sdc3 |
| Dlk1 Notch1 | 2.27E-18 | 2.043670258 | 0.144 | 0.019 | 3.83E-15 | 6 | Dlk1 Notch1 |
| Ntf3 Ntrk2 | 2.39E-18 | 4.648993016 | 0.092 | 0.003 | 4.03E-15 | 6 | Ntf3 Ntrk2 |
| Lrpap1 Sorl1 | 2.57E-18 | 0.221023554 | 0.582 | 0.237 | 4.34E-15 | 6 | Lrpap1 Sorl1 |
| Hbegf Prlr | 2.85E-18 | 3.679918042 | 0.092 | 0.003 | 4.82E-15 | 6 | Hbegf Prlr |
| Qdpr Dysf | 2.86E-18 | 1.408504765 | 0.278 | 0.082 | 4.83E-15 | 6 | Qdpr Dysf |
| Hras Grin2d | 4.98E-18 | 2.047374368 | 0.278 | 0.09 | 8.41E-15 | 6 | Hras Grin2d |
| Pdgfc Flt1 | 6.89E-18 | 2.342699887 | 0.163 | 0.029 | 1.16E-14 | 6 | Pdgfc Flt1 |
| Gdnf Gfra2 | 8.00E-18 | 3.297217721 | 0.124 | 0.013 | 1.35E-14 | 6 | Gdnf Gfra2 |
| Igf1 Igf1r | 8.57E-18 | 1.395550684 | 0.359 | 0.139 | 1.45E-14 | 6 | Igf1 Igf1r |
| Tnc Itga8 | 8.58E-18 | 3.44306985 | 0.121 | 0.012 | 1.45E-14 | 6 | Tnc Itga8 |
| Col4a1 Itgav | 1.18E-17 | 0.716387462 | 0.667 | 0.399 | 1.98E-14 | 6 | Col4a1 Itgav |
| Bdnf Ntrk2 | 1.77E-17 | 4.657373091 | 0.092 | 0.004 | 2.98E-14 | 6 | Bdnf Ntrk2 |

| | | | | | | | |
|---|---|---|---|---|---|---|---|
| Il1rn Il1r1 | 2.03E-17 | 4.049284273 | 0.078 | 0.001 | 3.43E-14 | 6 | Il1rn Il1r1 |
| Wnt5a Fzd7 | 2.41E-17 | 4.54075008 | 0.095 | 0.005 | 4.06E-14 | 6 | Wnt5a Fzd7 |
| Pthlh Pth1r | 2.80E-17 | 1.691631815 | 0.157 | 0.026 | 4.72E-14 | 6 | Pthlh Pth1r |
| Col5a1 Itga1 | 3.43E-17 | 1.414316696 | 0.324 | 0.118 | 5.79E-14 | 6 | Col5a1 Itga1 |
| Gnai2 Lhcgr | 3.93E-17 | 1.390047312 | 0.144 | 0.021 | 6.64E-14 | 6 | Gnai2 Lhcgr |
| Col7a1 Itga2 | 4.21E-17 | 9.621160021 | 0.072 | 0 | 7.11E-14 | 6 | Col7a1 Itga2 |
| Calm2 Kcnq3 | 1.34E-16 | 3.294504659 | 0.092 | 0.005 | 2.26E-13 | 6 | Calm2 Kcnq3 |
| Hgf Sdc2 | 1.44E-16 | 4.157142804 | 0.098 | 0.007 | 2.44E-13 | 6 | Hgf Sdc2 |
| App Lrp11 | 2.04E-16 | 0.721345612 | 0.948 | 0.692 | 3.44E-13 | 6 | App Lrp1 |
| a Mc5r | 2.15E-16 | 3.218011353 | 0.098 | 0.007 | 3.63E-13 | 6 | a Mc5r |
| Gdf10 Acvr1b | 2.31E-16 | 3.564692803 | 0.105 | 0.009 | 3.89E-13 | 6 | Gdf10 Acvr1b |
| Col1a1 Itga5 | 5.16E-16 | 1.287206424 | 0.297 | 0.103 | 8.70E-13 | 6 | Col1a1 Itga5 |
| Dusp18 Itga2 | 6.24E-16 | 4.214704277 | 0.088 | 0.005 | 1.05E-12 | 6 | Dusp18 Itga2 |
| Wnt5a Fzd8 | 6.28E-16 | 5.372079115 | 0.092 | 0.006 | 1.06E-12 | 6 | Wnt5a Fzd8 |
| Slit2 Gpc1 | 6.80E-16 | 2.30905219 | 0.088 | 0.005 | 1.15E-12 | 6 | Slit2 Gpc1 |
| Gstp1 Traf2 | 2.18E-15 | 0.384933106 | 0.235 | 0.066 | 3.69E-12 | 6 | Gstp1 Traf2 |
| Calm1 Pde1c | 3.23E-15 | 0.904303468 | 0.33 | 0.121 | 5.46E-12 | 6 | Calm1 Pde1c |
| Agrn Lrp41 | 4.37E-15 | 0.907970131 | 0.157 | 0.03 | 7.37E-12 | 6 | Agrn Lrp4 |
| Lamc1 Itga1 | 5.43E-15 | 1.420905097 | 0.288 | 0.108 | 9.17E-12 | 6 | Lamc1 Itga1 |
| Psap Lrp11 | 6.40E-15 | 0.700044291 | 0.935 | 0.698 | 1.08E-11 | 6 | Psap Lrp1 |
| Efna2 Epha4 | 9.41E-15 | 3.742835895 | 0.075 | 0.003 | 1.59E-11 | 6 | Efna2 Epha4 |
| Calm1 Kcnq3 | 1.75E-14 | 3.088502872 | 0.082 | 0.005 | 2.96E-11 | 6 | Calm1 Kcnq3 |
| Rims2 Abca1 | 3.04E-14 | 0.419492354 | 0.242 | 0.073 | 5.13E-11 | 6 | Rims2 Abca1 |
| Nlgn3 Nrxn2 | 3.14E-14 | 9.156673852 | 0.059 | 0 | 5.31E-11 | 6 | Nlgn3 Nrxn2 |
| Efna2 Epha3 | 3.14E-14 | 8.996631424 | 0.059 | 0 | 5.31E-11 | 6 | Efna2 Epha3 |
| Nxph1 Nrxn2 | 7.57E-14 | 5.772233221 | 0.062 | 0.001 | 1.28E-10 | 6 | Nxph1 Nrxn2 |
| Bmp7 Bmpr1b | 1.64E-13 | 9.151781631 | 0.056 | 0 | 2.77E-10 | 6 | Bmp7 Bmpr1b |
| Cntn2 Nrcam | 1.64E-13 | 8.797141287 | 0.056 | 0 | 2.77E-10 | 6 | Cntn2 Nrcam |
| Tgfb1 Cav1 | 1.84E-13 | 0.914050226 | 0.402 | 0.192 | 3.11E-10 | 6 | Tgfb1 Cav1 |
| Adam15 Itga5 | 2.07E-13 | 0.457568135 | 0.343 | 0.132 | 3.49E-10 | 6 | Adam15 Itga5 |
| Egf Erbb4 | 2.63E-13 | 2.217737516 | 0.157 | 0.038 | 4.45E-10 | 6 | Egf Erbb4 |
| Timp3 Agtr2 | 2.98E-13 | 2.870126818 | 0.127 | 0.025 | 5.03E-10 | 6 | Timp3 Agtr2 |
| Angpt2 Tek | 3.68E-13 | 1.733357664 | 0.141 | 0.03 | 6.21E-10 | 6 | Angpt2 Tek |
| Ereg Egfr | 5.04E-13 | 2.08063478 | 0.092 | 0.01 | 8.51E-10 | 6 | Ereg Egfr |
| Igfbp4 Fzd8 | 5.77E-13 | 1.646632351 | 0.239 | 0.086 | 9.73E-10 | 6 | Igfbp4 Fzd8 |
| Hspa1a Tlr41 | 6.50E-13 | 0.132794136 | 0.281 | 0.097 | 1.10E-09 | 6 | Hspa1a Tlr4 |
| Il6 F3 | 8.53E-13 | 9.083872518 | 0.052 | 0 | 1.44E-09 | 6 | Il6 F3 |
| Hbegf Cd82 | 9.72E-13 | 1.546991577 | 0.186 | 0.055 | 1.64E-09 | 6 | Hbegf Cd82 |
| Col4a6 Itga1 | 1.32E-12 | 2.083516974 | 0.121 | 0.023 | 2.22E-09 | 6 | Col4a6 Itga1 |
| Rgmb Bmpr1b | 1.36E-12 | 2.052833865 | 0.101 | 0.014 | 2.29E-09 | 6 | Rgmb Bmpr1b |
| Lama2 Itga2 | 1.76E-12 | 3.737271043 | 0.072 | 0.005 | 2.97E-09 | 6 | Lama2 Itga2 |
| App Slc45a3 | 1.79E-12 | 3.375564365 | 0.098 | 0.014 | 3.01E-09 | 6 | App Slc45a3 |
| Ereg Erbb4 | 3.29E-12 | 5.58964879 | 0.059 | 0.002 | 5.55E-09 | 6 | Ereg Erbb4 |
| Nrg2 Erbb2 | 5.01E-12 | 5.228866023 | 0.062 | 0.003 | 8.46E-09 | 6 | Nrg2 Erbb2 |
| Lama4 Itgav | 5.12E-12 | 0.764601486 | 0.608 | 0.385 | 8.64E-09 | 6 | Lama4 Itgav |
| Col4a5 Itga1 | 5.13E-12 | 1.007007929 | 0.19 | 0.058 | 8.65E-09 | 6 | Col4a5 Itga1 |
| Tnc Ptprz1 | 5.83E-12 | 3.659409775 | 0.062 | 0.003 | 9.83E-09 | 6 | Tnc Ptprz1 |
| Wnt5a Fzd5 | 6.25E-12 | 4.133095084 | 0.098 | 0.016 | 1.05E-08 | 6 | Wnt5a Fzd5 |
| Col4a1 Itga2 | 6.31E-12 | 3.041777912 | 0.088 | 0.011 | 1.06E-08 | 6 | Col4a1 Itga2 |
| Col4a5 Itgav | 6.60E-12 | 0.99931575 | 0.324 | 0.147 | 1.11E-08 | 6 | Col4a5 Itgav |
| Fgf2 Sdc1 | 7.21E-12 | 1.787678732 | 0.209 | 0.071 | 1.22E-08 | 6 | Fgf2 Sdc1 |

| | | | | | | | |
|---|---|---|---|---|---|---|---|
| Col4a6 Itgav | 7.89E-12 | 1.376195899 | 0.206 | 0.07 | 1.33E-08 | 6 | Col4a6 Itgav |
| Uba52 Erbb2 | 8.49E-12 | 4.944838319 | 0.092 | 0.013 | 1.43E-08 | 6 | Uba52 Erbb2 |
| Kng1 Gpr135 | 9.85E-12 | 7.583942138 | 0.052 | 0.001 | 1.66E-08 | 6 | Kng1 Gpr135 |
| Wnt11 Fzd7 | 1.05E-11 | 5.022880943 | 0.052 | 0.001 | 1.78E-08 | 6 | Wnt11 Fzd7 |
| Tnfsf12 Tnfrsf12a | 1.07E-11 | 3.06228727 | 0.052 | 0.001 | 1.80E-08 | 6 | Tnfsf12 Tnfrsf12a |
| Efnb1 Epha6 | 1.47E-11 | 0.266287919 | 0.186 | 0.056 | 2.48E-08 | 6 | Efnb1 Epha6 |
| Lgals3bp Vangl1 | 1.55E-11 | 1.954475701 | 0.186 | 0.06 | 2.61E-08 | 6 | Lgals3bp Vangl1 |
| Sorbs1 Insr | 1.90E-11 | 0.447619771 | 0.886 | 0.708 | 3.20E-08 | 6 | Sorbs1 Insr |
| Rspo2 Lgr6 | 2.31E-11 | 9.97450163 | 0.046 | 0 | 3.90E-08 | 6 | Rspo2 Lgr6 |
| Rims1 Slc17a7 | 2.31E-11 | 8.783317472 | 0.046 | 0 | 3.90E-08 | 6 | Rims1 Slc17a7 |
| Rspo2 Lgr4 | 2.95E-11 | 4.70167615 | 0.072 | 0.007 | 4.99E-08 | 6 | Rspo2 Lgr4 |
| Fn1 Tshr | 3.74E-11 | 2.410334591 | 0.078 | 0.009 | 6.31E-08 | 6 | Fn1 Tshr |
| Efemp2 Aqp1 | 5.03E-11 | 1.047034715 | 0.131 | 0.031 | 8.49E-08 | 6 | Efemp2 Aqp1 |
| Adam2 Itga91 | 5.29E-11 | 0.585046874 | 0.637 | 0.384 | 8.92E-08 | 6 | Adam2 Itga9 |
| Il1b Il1r1 | 5.49E-11 | 3.113980233 | 0.049 | 0.001 | 9.27E-08 | 6 | Il1b Il1r1 |
| Tctn1 Tmem671 | 8.00E-11 | 0.414791593 | 0.925 | 0.66 | 1.35E-07 | 6 | Tctn1 Tmem67 |
| Wnt5a Ror2 | 9.02E-11 | 5.313919975 | 0.052 | 0.002 | 1.52E-07 | 6 | Wnt5a Ror2 |
| Cdh1 Cdh2 | 1.20E-10 | 8.56176084 | 0.042 | 0 | 2.03E-07 | 6 | Cdh1 Cdh2 |
| Efna2 Epha1 | 1.20E-10 | 7.987526732 | 0.042 | 0 | 2.03E-07 | 6 | Efna2 Epha1 |
| Gnai2 Agtr2 | 1.40E-10 | 1.871873632 | 0.121 | 0.028 | 2.36E-07 | 6 | Gnai2 Agtr2 |
| Lamb1 Itga7 | 1.45E-10 | 5.794018926 | 0.062 | 0.005 | 2.45E-07 | 6 | Lamb1 Itga7 |
| Gnas Tshr | 1.50E-10 | 1.825976836 | 0.082 | 0.011 | 2.53E-07 | 6 | Gnas Tshr |
| Calr Tshr | 1.52E-10 | 2.24750986 | 0.078 | 0.01 | 2.57E-07 | 6 | Calr Tshr |
| Npnt Itgb1 | 1.74E-10 | 0.592813558 | 0.353 | 0.159 | 2.93E-07 | 6 | Npnt Itgb1 |
| Hmgb1 Thbd | 1.74E-10 | 1.416033903 | 0.353 | 0.192 | 2.93E-07 | 6 | Hmgb1 Thbd |
| Pf4 Sdc2 | 1.77E-10 | 4.390696265 | 0.065 | 0.006 | 2.99E-07 | 6 | Pf4 Sdc2 |
| Fgf5 Fgfr2 | 1.87E-10 | 2.771052169 | 0.072 | 0.008 | 3.16E-07 | 6 | Fgf5 Fgfr2 |
| Ncan Cdh2 | 2.65E-10 | 5.702125909 | 0.046 | 0.001 | 4.48E-07 | 6 | Ncan Cdh2 |
| Anxa1 Egfr | 2.95E-10 | 0.495211659 | 0.408 | 0.194 | 4.98E-07 | 6 | Anxa1 Egfr |
| Igf1 Insr | 3.09E-10 | 0.713185045 | 0.33 | 0.151 | 5.22E-07 | 6 | Igf1 Insr |
| Col3a1 Itga2 | 3.98E-10 | 2.501972633 | 0.082 | 0.012 | 6.72E-07 | 6 | Col3a1 Itga2 |
| Ntng1 Lrrc4c | 4.93E-10 | 3.576392746 | 0.049 | 0.002 | 8.32E-07 | 6 | Ntng1 Lrrc4c |
| Gnai2 Tshr | 6.11E-10 | 1.619826986 | 0.078 | 0.011 | 1.03E-06 | 6 | Gnai2 Tshr |
| Rbp4 Stra6 | 6.24E-10 | 9.251504332 | 0.039 | 0 | 1.05E-06 | 6 | Rbp4 Stra6 |
| Gdnf Gfra1 | 6.24E-10 | 9.057268312 | 0.039 | 0 | 1.05E-06 | 6 | Gdnf Gfra1 |
| Il6 Il6st | 8.86E-10 | 2.211951618 | 0.069 | 0.008 | 1.50E-06 | 6 | Il6 Il6st |
| Bmp4 Bmpr1b | 9.80E-10 | 0.954545677 | 0.088 | 0.016 | 1.65E-06 | 6 | Bmp4 Bmpr1b |
| Bmp6 Bmpr1b | 1.07E-09 | 0.578246672 | 0.088 | 0.016 | 1.80E-06 | 6 | Bmp6 Bmpr1b |
| Pgf Nrp1 | 1.07E-09 | 0.432235327 | 0.199 | 0.069 | 1.81E-06 | 6 | Pgf Nrp1 |
| Ccl25 Ackr4 | 1.14E-09 | 1.535576578 | 0.095 | 0.019 | 1.93E-06 | 6 | Ccl25 Ackr4 |
| Lamc2 Itga2 | 1.45E-09 | 4.175862968 | 0.042 | 0.001 | 2.44E-06 | 6 | Lamc2 Itga2 |
| Tnc Itga5 | 1.64E-09 | 1.035993633 | 0.101 | 0.022 | 2.77E-06 | 6 | Tnc Itga5 |
| Sema6a Plxna2 | 1.64E-09 | 0.228937363 | 0.317 | 0.135 | 2.77E-06 | 6 | Sema6a Plxna2 |
| Tnc Itgav | 1.84E-09 | 1.269762404 | 0.176 | 0.062 | 3.11E-06 | 6 | Tnc Itgav |
| Adam12 Itga9 | 2.61E-09 | 2.286616698 | 0.114 | 0.029 | 4.41E-06 | 6 | Adam12 Itga9 |
| Tgfb1 Acvrl1 | 2.76E-09 | 0.490164609 | 0.395 | 0.206 | 4.65E-06 | 6 | Tgfb1 Acvrl1 |
| Fn1 Tmprss6 | 3.08E-09 | 3.307635828 | 0.062 | 0.007 | 5.20E-06 | 6 | Fn1 Tmprss6 |
| Rspo3 Lgr5 | 3.25E-09 | 10.1740228 | 0.036 | 0 | 5.48E-06 | 6 | Rspo3 Lgr5 |
| Lamb3 Col17a1 | 4.17E-09 | 2.616244334 | 0.052 | 0.004 | 7.04E-06 | 6 | Lamb3 Col17a1 |
| Wnt5a Mcam | 6.94E-09 | 6.86782145 | 0.039 | 0.001 | 1.17E-05 | 6 | Wnt5a Mcam |
| Efna2 Epha7 | 7.24E-09 | 4.882972986 | 0.039 | 0.001 | 1.22E-05 | 6 | Efna2 Epha7 |

| | | | | | | | |
|---|---|---|---|---|---|---|---|
| Lrpap1 Ldlr | 7.98E-09 | 0.429081766 | 0.595 | 0.349 | 1.35E-05 | 6 | Lrpap1 Ldlr |
| Nrg2 Erbb4 | 1.15E-08 | 3.845744542 | 0.042 | 0.002 | 1.95E-05 | 6 | Nrg2 Erbb4 |
| Ncam1 Gfra1 | 1.53E-08 | 2.983255702 | 0.046 | 0.003 | 2.58E-05 | 6 | Ncam1 Gfra1 |
| Tnc Sdc4 | 1.54E-08 | 0.783245088 | 0.144 | 0.047 | 2.59E-05 | 6 | Tnc Sdc4 |
| Sema4f Nrp2 | 2.63E-08 | 1.196155378 | 0.072 | 0.012 | 4.44E-05 | 6 | Sema4f Nrp2 |
| Adam12 Sdc4 | 3.23E-08 | 1.717121252 | 0.095 | 0.023 | 5.45E-05 | 6 | Adam12 Sdc4 |
| Tgfa Egfr1 | 4.19E-08 | 0.452494242 | 0.827 | 0.493 | 7.07E-05 | 6 | Tgfa Egfr |
| Clcf1 Il6st | 4.47E-08 | 1.872335752 | 0.095 | 0.024 | 7.54E-05 | 6 | Clcf1 Il6st |
| Mdk Sdc1 | 5.02E-08 | 1.319292859 | 0.127 | 0.04 | 8.48E-05 | 6 | Mdk Sdc1 |
| Col18a1 Itga5 | 6.88E-08 | 0.929545157 | 0.163 | 0.062 | 0.000116172 | 6 | Col18a1 Itga5 |
| Vtn Plaur | 7.45E-08 | 3.21430293 | 0.042 | 0.003 | 0.000125826 | 6 | Vtn Plaur |
| Farp2 Plxna4 | 9.30E-08 | 1.148453685 | 0.392 | 0.289 | 0.000157005 | 6 | Farp2 Plxna4 |
| Nid1 Itga3 | 9.41E-08 | 1.894539098 | 0.121 | 0.04 | 0.000158786 | 6 | Nid1 Itga3 |
| Col1a1 Itga2 | 9.71E-08 | 2.629493436 | 0.065 | 0.011 | 0.000163972 | 6 | Col1a1 Itga2 |
| Vcan Egfr | 1.67E-07 | 0.799339834 | 0.242 | 0.116 | 0.00028252 | 6 | Vcan Egfr |
| Thbs1 Sdc1 | 1.77E-07 | 0.346666597 | 0.196 | 0.081 | 0.000298647 | 6 | Thbs1 Sdc1 |
| Slit2 Robo1 | 1.89E-07 | 1.439874383 | 0.085 | 0.021 | 0.000319636 | 6 | Slit2 Robo1 |
| Mmp9 Ephb2 | 1.94E-07 | 2.581610378 | 0.033 | 0.001 | 0.000328135 | 6 | Mmp9 Ephb2 |
| Il34 Csf1r | 1.95E-07 | 2.143261576 | 0.069 | 0.013 | 0.000329122 | 6 | Il34 Csf1r |
| Sema3e Plxnd1 | 2.05E-07 | 1.556177966 | 0.069 | 0.013 | 0.000346533 | 6 | Sema3e Plxnd1 |
| Efna4 Epha5 | 2.41E-07 | 2.060248717 | 0.075 | 0.017 | 0.000407584 | 6 | Efna4 Epha5 |
| Fgf5 Fgfr1 | 2.57E-07 | 1.71446724 | 0.069 | 0.013 | 0.000434296 | 6 | Fgf5 Fgfr1 |
| Ptn Ptprz1 | 2.63E-07 | 2.816077595 | 0.052 | 0.007 | 0.000443592 | 6 | Ptn Ptprz1 |
| Sema7a Itga1 | 2.74E-07 | 0.337369045 | 0.069 | 0.013 | 0.000461737 | 6 | Sema7a Itga1 |
| Rspo1 Znrf3 | 2.98E-07 | 1.972556887 | 0.036 | 0.002 | 0.000502719 | 6 | Rspo1 Znrf3 |
| Rims1 Slc18a2 | 3.23E-07 | 4.759662265 | 0.042 | 0.004 | 0.00054503 | 6 | Rims1 Slc18a2 |
| Lrpap1 Lrp8 | 3.25E-07 | 0.801270617 | 0.258 | 0.135 | 0.000548663 | 6 | Lrpap1 Lrp8 |
| Spp1 S1pr1 | 3.26E-07 | 2.329938053 | 0.049 | 0.006 | 0.000550993 | 6 | Spp1 S1pr1 |
| L1cam Ephb2 | 3.37E-07 | 4.054861664 | 0.039 | 0.003 | 0.00056836 | 6 | L1cam Ephb2 |
| Slit2 Robo2 | 3.51E-07 | 3.320127882 | 0.039 | 0.003 | 0.00059175 | 6 | Slit2 Robo2 |
| Hspg2 Chrm3 | 3.76E-07 | 1.034832764 | 0.082 | 0.02 | 0.000635496 | 6 | Hspg2 Chrm3 |
| Il18 Il1rl2 | 4.03E-07 | 2.013438068 | 0.062 | 0.011 | 0.000679928 | 6 | Il18 Il1rl2 |
| Proc Thbd | 4.62E-07 | 0.133214639 | 0.078 | 0.019 | 0.000780152 | 6 | Proc Thbd |
| Ccl19 Ackr2 | 4.63E-07 | 7.815817758 | 0.026 | 0 | 0.000780768 | 6 | Ccl19 Ackr2 |
| Efemp1 Egfr1 | 6.51E-07 | 0.216600624 | 0.346 | 0.176 | 0.001099335 | 6 | Efemp1 Egfr |
| Tgm2 Itga91 | 8.92E-07 | 0.4877407 | 0.673 | 0.45 | 0.001506126 | 6 | Tgm2 Itga9 |
| Nlgn1 Nrxn2 | 9.53E-07 | 5.984575773 | 0.029 | 0.001 | 0.001608607 | 6 | Nlgn1 Nrxn2 |
| Clec11a Kit | 9.59E-07 | 5.453078996 | 0.029 | 0.001 | 0.00161819 | 6 | Clec11a Kit |
| Nppa Npr3 | 9.82E-07 | 4.524951384 | 0.029 | 0.001 | 0.001657072 | 6 | Nppa Npr3 |
| Il4 Il13ra1 | 1.32E-06 | 0.341434849 | 0.078 | 0.02 | 0.002234203 | 6 | Il4 Il13ra1 |
| Gnrh1 Gnrhr | 1.37E-06 | 4.030678625 | 0.033 | 0.002 | 0.002311898 | 6 | Gnrh1 Gnrhr |
| Efnb3 Ephb1 | 1.41E-06 | 3.577644112 | 0.033 | 0.002 | 0.002387308 | 6 | Efnb3 Ephb1 |
| Col4a5 Itga2 | 1.50E-06 | 2.634463233 | 0.042 | 0.005 | 0.00253532 | 6 | Col4a5 Itga2 |
| Col1a2 Itgb11 | 1.58E-06 | 0.363491601 | 0.944 | 0.714 | 0.00266227 | 6 | Col1a2 Itgb1 |
| Calm2 Egfr1 | 2.37E-06 | 0.388032113 | 0.846 | 0.614 | 0.003997962 | 6 | Calm2 Egfr |
| Npy Npy4r | 2.43E-06 | 7.493356115 | 0.023 | 0 | 0.00410621 | 6 | Npy Npy4r |
| Il4 Il13ra2 | 2.43E-06 | 7.433109644 | 0.023 | 0 | 0.00410621 | 6 | Il4 Il13ra2 |
| Lamb1 Itga3 | 4.10E-06 | 2.988029626 | 0.062 | 0.014 | 0.006913418 | 6 | Lamb1 Itga3 |
| Wnt5a Ryk | 4.52E-06 | 2.991978298 | 0.101 | 0.036 | 0.007629019 | 6 | Wnt5a Ryk |
| Uba52 Tgfbr2 | 5.53E-06 | 1.327011009 | 0.095 | 0.031 | 0.00933083 | 6 | Uba52 Tgfbr2 |
| Fgf18 Fgfr3 | 5.73E-06 | 2.048273132 | 0.042 | 0.006 | 0.009665094 | 6 | Fgf18 Fgfr3 |

| | | | | | | | |
|---|---|---|---|---|---|---|---|
| Tnc Itga2 | 6.88E-06 | 2.564371326 | 0.029 | 0.002 | 0.011614206 | 6 | Tnc Itga2 |
| Lama5 Bcam | 6.89E-06 | 0.860820699 | 0.18 | 0.088 | 0.011631277 | 6 | Lama5 Bcam |
| Col1a1 Tmprss6 | 7.24E-06 | 2.779865067 | 0.036 | 0.004 | 0.012225303 | 6 | Col1a1 Tmprss6 |
| Rarres2 Gpr1 | 7.33E-06 | 4.668110168 | 0.033 | 0.003 | 0.01236788 | 6 | Rarres2 Gpr1 |
| Efnb2 Epha3 | 7.59E-06 | 1.266729211 | 0.085 | 0.026 | 0.01281522 | 6 | Efnb2 Epha3 |
| Ngf Sorcs3 | 7.84E-06 | 1.379009419 | 0.033 | 0.003 | 0.013227004 | 6 | Ngf Sorcs3 |
| Hras Cav11 | 8.71E-06 | 0.434461518 | 0.356 | 0.228 | 0.014701447 | 6 | Hras Cav1 |
| Uba52 Tgfbr1 | 1.06E-05 | 1.991030368 | 0.072 | 0.021 | 0.017822611 | 6 | Uba52 Tgfbr1 |
| Lamc1 Itga2 | 1.12E-05 | 1.997768135 | 0.056 | 0.012 | 0.018912908 | 6 | Lamc1 Itga2 |
| Clcf1 Cntfr | 1.29E-05 | 8.087187168 | 0.02 | 0 | 0.021733479 | 6 | Clcf1 Cntfr |
| Oxt Oxtr | 1.29E-05 | 6.732560252 | 0.02 | 0 | 0.021733479 | 6 | Oxt Oxtr |
| Tnc Itga9 | 1.57E-05 | 0.728160835 | 0.167 | 0.078 | 0.02651941 | 6 | Tnc Itga9 |
| Lama5 Itga2 | 1.61E-05 | 1.658064259 | 0.052 | 0.011 | 0.027255219 | 6 | Lama5 Itga2 |
| Fgf16 Fgfr2 | 1.68E-05 | 2.312790837 | 0.042 | 0.007 | 0.028383239 | 6 | Fgf16 Fgfr2 |
| Calm1 Sell | 2.56E-152 | 7.023243571 | 0.673 | 0.023 | 4.32E-149 | 7 | Calm1 Sell |
| Calm2 Sell | 1.36E-148 | 6.613925871 | 0.663 | 0.024 | 2.29E-145 | 7 | Calm2 Sell |
| Calm1 Grm5 | 2.26E-148 | 6.0348417 | 0.743 | 0.067 | 3.81E-145 | 7 | Calm1 Grm5 |
| Calm2 Grm5 | 9.19E-141 | 5.418387623 | 0.733 | 0.076 | 1.55E-137 | 7 | Calm2 Grm5 |
| Efnb2 Grm5 | 3.16E-138 | 7.605066162 | 0.63 | 0.025 | 5.33E-135 | 7 | Efnb2 Grm5 |
| Podxl Sell | 1.84E-137 | 7.354280584 | 0.61 | 0.017 | 3.11E-134 | 7 | Podxl Sell |
| Cd34 Sell | 1.84E-136 | 6.828430694 | 0.617 | 0.022 | 3.10E-133 | 7 | Cd34 Sell |
| Cfh Sell | 1.51E-135 | 6.96772049 | 0.613 | 0.022 | 2.55E-132 | 7 | Cfh Sell |
| Calm1 Scn4a | 6.18E-131 | 7.702735146 | 0.563 | 0.008 | 1.04E-127 | 7 | Calm1 Scn4a |
| Calm2 Scn4a | 8.12E-127 | 6.625009171 | 0.553 | 0.009 | 1.37E-123 | 7 | Calm2 Scn4a |
| Gnai2 S1pr4 | 4.19E-125 | 5.793861127 | 0.597 | 0.029 | 7.08E-122 | 7 | Gnai2 S1pr4 |
| Sema4d Cd72 | 6.61E-120 | 5.289809518 | 0.67 | 0.079 | 1.12E-116 | 7 | Sema4d Cd72 |
| Vcam1 Itgb7 | 2.65E-111 | 5.295771704 | 0.6 | 0.053 | 4.47E-108 | 7 | Vcam1 Itgb7 |
| Icam1 Itgal | 1.22E-103 | 4.451237516 | 0.57 | 0.049 | 2.06E-100 | 7 | Icam1 Itgal |
| Tgfb1 Cxcr4 | 2.15E-99 | 4.360465254 | 0.637 | 0.103 | 3.62E-96 | 7 | Tgfb1 Cxcr4 |
| Calm1 Kcnq5 | 1.41E-95 | 4.097973865 | 0.677 | 0.151 | 2.38E-92 | 7 | Calm1 Kcnq5 |
| Psen1 Notch2 | 1.86E-95 | 1.72222871 | 0.963 | 0.935 | 3.14E-92 | 7 | Psen1 Notch2 |
| App Cd74 | 4.79E-88 | 1.993881699 | 0.933 | 0.633 | 8.09E-85 | 7 | App Cd74 |
| Icam2 Itgal | 1.46E-87 | 4.128237746 | 0.533 | 0.063 | 2.46E-84 | 7 | Icam2 Itgal |
| Tgm2 Itga4 | 3.66E-87 | 2.662900381 | 0.86 | 0.583 | 6.18E-84 | 7 | Tgm2 Itga4 |
| Adam2 Itgb7 | 1.09E-85 | 4.523311358 | 0.523 | 0.06 | 1.84E-82 | 7 | Adam2 Itgb7 |
| Calm2 Kcnq5 | 1.29E-85 | 3.435838666 | 0.667 | 0.166 | 2.17E-82 | 7 | Calm2 Kcnq5 |
| Calm1 Kcnn4 | 1.69E-79 | 3.026957145 | 0.637 | 0.138 | 2.85E-76 | 7 | Calm1 Kcnn4 |
| Icam4 Itgal | 2.66E-79 | 5.218513907 | 0.44 | 0.034 | 4.49E-76 | 7 | Icam4 Itgal |
| Apoe Sorl1 | 9.90E-76 | 4.496483465 | 0.557 | 0.109 | 1.67E-72 | 7 | Apoe Sorl1 |
| Gpi1 Amfr | 3.37E-74 | 1.036759553 | 0.96 | 0.878 | 5.69E-71 | 7 | Gpi1 Amfr |
| Selplg Itgb2 | 8.32E-74 | 6.024532603 | 0.37 | 0.015 | 1.40E-70 | 7 | Selplg Itgb2 |
| Psen1 Cd44 | 7.56E-68 | 2.211292228 | 0.837 | 0.602 | 1.28E-64 | 7 | Psen1 Cd44 |
| Pkm Cd44 | 5.33E-67 | 1.696288491 | 0.84 | 0.6 | 9.00E-64 | 7 | Pkm Cd44 |
| Mmp9 Itgb2 | 1.40E-62 | 9.533516822 | 0.273 | 0.001 | 2.37E-59 | 7 | Mmp9 Itgb2 |
| L1cam Itgav | 2.04E-62 | 3.194143995 | 0.527 | 0.118 | 3.44E-59 | 7 | L1cam Itgav |
| Icam4 Itgb3 | 2.13E-62 | 4.70475768 | 0.367 | 0.03 | 3.59E-59 | 7 | Icam4 Itgb3 |
| Psen1 Ncstn | 3.06E-59 | 1.136666831 | 0.963 | 0.925 | 5.17E-56 | 7 | Psen1 Ncstn |
| Lrpap1 Sorl11 | 1.34E-57 | 2.839218052 | 0.62 | 0.227 | 2.27E-54 | 7 | Lrpap1 Sorl1 |
| Fn1 Cd79a | 3.48E-57 | 5.964411625 | 0.287 | 0.01 | 5.87E-54 | 7 | Fn1 Cd79a |
| Efnb2 Pecam1 | 6.98E-56 | 2.415756399 | 0.637 | 0.271 | 1.18E-52 | 7 | Efnb2 Pecam1 |
| Efnb2 Grm1 | 1.01E-53 | 5.595873098 | 0.33 | 0.031 | 1.71E-50 | 7 | Efnb2 Grm1 |

| | | | | | | | |
|---|---|---|---|---|---|---|---|
| Ntng2 Lrrc4 | 1.83E-53 | 5.151250765 | 0.35 | 0.039 | 3.09E-50 | 7 | Ntng2 Lrrc4 |
| Ebi3 Il6st | 1.98E-53 | 6.219983859 | 0.26 | 0.007 | 3.34E-50 | 7 | Ebi3 Il6st |
| Hsp90b1 Tlr1 | 5.03E-52 | 1.913942924 | 0.79 | 0.615 | 8.50E-49 | 7 | Hsp90b1 Tlr1 |
| Icam4 Itgav | 1.32E-51 | 3.425663758 | 0.513 | 0.15 | 2.23E-48 | 7 | Icam4 Itgav |
| Hsp90b1 Tlr9 | 4.83E-51 | 2.717157811 | 0.593 | 0.227 | 8.15E-48 | 7 | Hsp90b1 Tlr9 |
| Il1b Adrb2 | 5.95E-50 | 6.828718067 | 0.233 | 0.004 | 1.00E-46 | 7 | Il1b Adrb2 |
| Cd14 Itgb1 | 2.40E-49 | 5.881712886 | 0.26 | 0.012 | 4.05E-46 | 7 | Cd14 Itgb1 |
| Gnai2 S1pr11 | 3.35E-49 | 1.090034418 | 0.797 | 0.601 | 5.65E-46 | 7 | Gnai2 S1pr1 |
| Osm Lifr | 7.21E-49 | 8.637098942 | 0.22 | 0.002 | 1.22E-45 | 7 | Osm Lifr |
| Cd14 Itga4 | 7.46E-49 | 7.395453173 | 0.24 | 0.007 | 1.26E-45 | 7 | Cd14 Itga4 |
| Ltb Cd40 | 2.03E-48 | 6.619052868 | 0.223 | 0.003 | 3.42E-45 | 7 | Ltb Cd40 |
| Tgfb1 Itgav1 | 2.95E-47 | 1.906518654 | 0.717 | 0.406 | 4.97E-44 | 7 | Tgfb1 Itgav |
| Gnai2 Oprm1 | 2.51E-46 | 5.453750624 | 0.25 | 0.013 | 4.23E-43 | 7 | Gnai2 Oprm1 |
| Calm1 Oprm1 | 3.01E-46 | 5.379723193 | 0.25 | 0.013 | 5.08E-43 | 7 | Calm1 Oprm1 |
| Vim Cd44 | 3.59E-46 | 1.41179246 | 0.783 | 0.588 | 6.06E-43 | 7 | Vim Cd44 |
| Mfng Notch21 | 3.52E-45 | 1.3837883 | 0.793 | 0.62 | 5.95E-42 | 7 | Mfng Notch2 |
| Hp Itgb2 | 3.87E-44 | 5.328471585 | 0.257 | 0.018 | 6.53E-41 | 7 | Hp Itgb2 |
| Fbln2 Itgb3 | 1.61E-43 | 2.684391057 | 0.417 | 0.091 | 2.72E-40 | 7 | Fbln2 Itgb3 |
| Hras Tlr9 | 2.06E-43 | 2.650650977 | 0.53 | 0.182 | 3.48E-40 | 7 | Hras Tlr9 |
| Tnfsf4 Traf2 | 7.16E-43 | 9.92209358 | 0.19 | 0.001 | 1.21E-39 | 7 | Tnfsf4 Traf2 |
| Mmp9 Cd44 | 2.19E-42 | 7.056242527 | 0.23 | 0.012 | 3.70E-39 | 7 | Mmp9 Cd44 |
| Col4a4 Itgav | 3.11E-42 | 2.114330354 | 0.613 | 0.312 | 5.25E-39 | 7 | Col4a4 Itgav |
| Ltb Tnfrsf1a | 6.91E-42 | 5.633474125 | 0.2 | 0.004 | 1.17E-38 | 7 | Ltb Tnfrsf1a |
| Ltb Ltbr | 3.80E-41 | 5.576891465 | 0.197 | 0.004 | 6.41E-38 | 7 | Ltb Ltbr |
| Gnai2 Adcy71 | 1.81E-40 | 0.831129833 | 0.963 | 0.866 | 3.05E-37 | 7 | Gnai2 Adcy7 |
| Camp P2rx7 | 3.69E-40 | 8.272297555 | 0.217 | 0.011 | 6.23E-37 | 7 | Camp P2rx7 |
| Lrp1b Plaur | 4.71E-40 | 5.579219331 | 0.233 | 0.016 | 7.95E-37 | 7 | Lrp1b Plaur |
| Osm Il6st | 4.98E-40 | 7.307950431 | 0.187 | 0.003 | 8.40E-37 | 7 | Osm Il6st |
| Ebi3 Il27ra | 8.79E-40 | 8.914132845 | 0.177 | 0.001 | 1.48E-36 | 7 | Ebi3 Il27ra |
| Tnfsf14 Ltbr | 3.43E-39 | 5.501409458 | 0.22 | 0.013 | 5.80E-36 | 7 | Tnfsf14 Ltbr |
| Col4a3 Itgav | 2.49E-37 | 2.174567598 | 0.553 | 0.263 | 4.21E-34 | 7 | Col4a3 Itgav |
| Icam4 Itga4 | 1.31E-36 | 3.323667533 | 0.523 | 0.253 | 2.21E-33 | 7 | Icam4 Itga4 |
| Adam9 Itgav | 1.55E-36 | 1.524912085 | 0.717 | 0.532 | 2.62E-33 | 7 | Adam9 Itgav |
| Calm1 Ptpra1 | 1.21E-34 | 0.546472137 | 0.997 | 0.942 | 2.04E-31 | 7 | Calm1 Ptpra |
| Tgfb1 Tgfbr2 | 1.25E-34 | 1.222815889 | 0.76 | 0.63 | 2.10E-31 | 7 | Tgfb1 Tgfbr2 |
| Vcam1 Itga4 | 8.27E-34 | 2.26378523 | 0.66 | 0.544 | 1.40E-30 | 7 | Vcam1 Itga4 |
| a Mgrn1 | 8.63E-34 | 2.112569455 | 0.7 | 0.613 | 1.46E-30 | 7 | a Mgrn1 |
| Sema4a Plxnd1 | 1.48E-33 | 4.729749078 | 0.23 | 0.026 | 2.50E-30 | 7 | Sema4a Plxnd1 |
| Calm1 Trpc5 | 1.91E-33 | 3.528438071 | 0.33 | 0.075 | 3.23E-30 | 7 | Calm1 Trpc5 |
| Proc Itgb2 | 1.03E-32 | 5.210331815 | 0.22 | 0.024 | 1.74E-29 | 7 | Proc Itgb2 |
| Vegfa Itgav | 2.02E-32 | 1.907141768 | 0.643 | 0.455 | 3.41E-29 | 7 | Vegfa Itgav |
| Mmp9 Lrp1 | 3.33E-32 | 4.468822285 | 0.213 | 0.022 | 5.62E-29 | 7 | Mmp9 Lrp1 |
| Adam15 Itgav | 5.47E-32 | 1.799812512 | 0.647 | 0.487 | 9.24E-29 | 7 | Adam15 Itgav |
| Bgn Tlr1 | 1.88E-31 | 1.480577908 | 0.723 | 0.621 | 3.17E-28 | 7 | Bgn Tlr1 |
| Sema7a Plxnc1 | 3.53E-30 | 2.827490135 | 0.317 | 0.074 | 5.96E-27 | 7 | Sema7a Plxnc1 |
| Calm2 Trpc5 | 1.32E-29 | 3.2897824 | 0.323 | 0.082 | 2.23E-26 | 7 | Calm2 Trpc5 |
| Calr Itgav1 | 4.60E-29 | 1.097678099 | 0.757 | 0.606 | 7.77E-26 | 7 | Calr Itgav |
| Ifng Ifngr2 | 1.11E-28 | 11.80349801 | 0.123 | 0 | 1.87E-25 | 7 | Ifng Ifngr2 |
| Alox5ap Alox5 | 3.92E-28 | 3.224026226 | 0.22 | 0.031 | 6.61E-25 | 7 | Alox5ap Alox5 |
| Pdgfb Itgav | 1.07E-27 | 2.213550319 | 0.413 | 0.157 | 1.80E-24 | 7 | Pdgfb Itgav |
| Hdc Hrh1 | 2.78E-27 | 4.143559006 | 0.203 | 0.027 | 4.70E-24 | 7 | Hdc Hrh1 |

| | | | | | | | |
|---|---|---|---|---|---|---|---|
| Hp Asgr2 | 3.19E-27 | 4.497798381 | 0.173 | 0.015 | 5.38E-24 | 7 | Hp Asgr2 |
| Ccl3 Ccr5 | 3.52E-27 | 13.08838746 | 0.117 | 0 | 5.94E-24 | 7 | Ccl3 Ccr5 |
| Dll4 Notch21 | 4.95E-27 | 1.204758201 | 0.68 | 0.594 | 8.35E-24 | 7 | Dll4 Notch2 |
| Ly86 Cd180 | 9.20E-27 | 7.897768833 | 0.137 | 0.005 | 1.55E-23 | 7 | Ly86 Cd180 |
| Fn1 Itgb7 | 1.57E-26 | 2.826986314 | 0.287 | 0.069 | 2.66E-23 | 7 | Fn1 Itgb7 |
| Col4a4 Cd47 | 1.81E-26 | 1.415953424 | 0.653 | 0.594 | 3.05E-23 | 7 | Col4a4 Cd47 |
| Tnf Tnfrsf1b | 3.76E-26 | 5.951212708 | 0.13 | 0.004 | 6.35E-23 | 7 | Tnf Tnfrsf1b |
| Camp Igf1r | 7.48E-26 | 5.2130595 | 0.207 | 0.032 | 1.26E-22 | 7 | Camp Igf1r |
| Ifng Ifngr1 | 1.11E-25 | 12.31472897 | 0.11 | 0 | 1.87E-22 | 7 | Ifng Ifngr1 |
| Il16 Ccr5 | 3.10E-25 | 2.381373178 | 0.32 | 0.089 | 5.23E-22 | 7 | Il16 Ccr5 |
| Tnf Traf2 | 3.28E-25 | 6.677266448 | 0.133 | 0.006 | 5.54E-22 | 7 | Tnf Traf2 |
| Ptdss1 Jmjd6 | 3.71E-25 | 1.278014365 | 0.9 | 0.88 | 6.27E-22 | 7 | Ptdss1 Jmjd6 |
| Il10 Il10ra | 4.70E-25 | 6.42895122 | 0.117 | 0.002 | 7.94E-22 | 7 | Il10 Il10ra |
| Ccl4 Ccr5 | 6.18E-25 | 13.01117811 | 0.107 | 0 | 1.04E-21 | 7 | Ccl4 Ccr5 |
| Adam15 Itgb3 | 7.01E-25 | 2.330093392 | 0.417 | 0.182 | 1.18E-21 | 7 | Adam15 Itgb3 |
| F10 Itgb2 | 1.63E-24 | 4.766117251 | 0.123 | 0.004 | 2.75E-21 | 7 | F10 Itgb2 |
| Dscam Dcc | 3.45E-24 | 11.78416266 | 0.103 | 0 | 5.82E-21 | 7 | Dscam Dcc |
| Il27 Il27ra | 3.94E-24 | 7.11009218 | 0.117 | 0.003 | 6.65E-21 | 7 | Il27 Il27ra |
| Tnfsf14 Tnfrsf14 | 7.37E-24 | 9.810623482 | 0.107 | 0.001 | 1.24E-20 | 7 | Tnfsf14 Tnfrsf14 |
| Jag2 Notch2 | 1.39E-23 | 1.562803038 | 0.527 | 0.317 | 2.35E-20 | 7 | Jag2 Notch2 |
| Il10 Il10rb | 1.73E-23 | 5.635775685 | 0.11 | 0.002 | 2.92E-20 | 7 | Il10 Il10rb |
| Ccl4 Ccr1 | 1.92E-23 | 12.39019716 | 0.1 | 0 | 3.24E-20 | 7 | Ccl4 Ccr1 |
| Il1b Il1rap | 4.38E-23 | 6.284163191 | 0.147 | 0.013 | 7.39E-20 | 7 | Il1b Il1rap |
| Dll1 Notch21 | 1.10E-22 | 1.283781045 | 0.66 | 0.61 | 1.85E-19 | 7 | Dll1 Notch2 |
| Col4a3 Cd47 | 4.83E-22 | 1.55582173 | 0.583 | 0.487 | 8.15E-19 | 7 | Col4a3 Cd47 |
| Ccl3 Ccr1 | 5.90E-22 | 12.33151048 | 0.093 | 0 | 9.96E-19 | 7 | Ccl3 Ccr1 |
| a Atrn | 1.11E-21 | 1.320167861 | 0.7 | 0.61 | 1.88E-18 | 7 | a Atrn |
| Il1a Il1rap | 2.33E-21 | 7.47671734 | 0.1 | 0.002 | 3.93E-18 | 7 | Il1a Il1rap |
| Gzmb Pgrmc1 | 3.26E-21 | 11.05005564 | 0.09 | 0 | 5.51E-18 | 7 | Gzmb Pgrmc1 |
| Tgm2 Itgb31 | 3.72E-21 | 1.845676932 | 0.45 | 0.228 | 6.28E-18 | 7 | Tgm2 Itgb3 |
| App Ncstn1 | 1.76E-20 | 0.394155281 | 0.93 | 0.94 | 2.97E-17 | 7 | App Ncstn |
| Fn1 Plaur | 2.91E-20 | 2.686407231 | 0.3 | 0.105 | 4.90E-17 | 7 | Fn1 Plaur |
| Lta Tnfrsf1b | 3.90E-20 | 8.217843766 | 0.09 | 0.001 | 6.59E-17 | 7 | Lta Tnfrsf1b |
| B2m Kir3dl1 | 7.38E-20 | 4.8035791 | 0.153 | 0.022 | 1.25E-16 | 7 | B2m Kir3dl1 |
| Nucb2 Erap1 | 9.91E-20 | 1.292721844 | 0.667 | 0.744 | 1.67E-16 | 7 | Nucb2 Erap1 |
| Nps Npsr1 | 1.10E-19 | 4.261493842 | 0.17 | 0.029 | 1.86E-16 | 7 | Nps Npsr1 |
| Hgf St14 | 1.16E-19 | 7.548444708 | 0.097 | 0.003 | 1.96E-16 | 7 | Hgf St14 |
| Ccl5 Ccr5 | 1.91E-18 | 3.885271649 | 0.103 | 0.006 | 3.23E-15 | 7 | Ccl5 Ccr5 |
| Il18 Cd48 | 1.97E-18 | 3.731513645 | 0.137 | 0.017 | 3.32E-15 | 7 | Il18 Cd48 |
| Icam4 Rhag | 2.16E-18 | 7.29245136 | 0.087 | 0.002 | 3.64E-15 | 7 | Icam4 Rhag |
| Ccl25 Ccr10 | 6.10E-18 | 4.112990998 | 0.093 | 0.004 | 1.03E-14 | 7 | Ccl25 Ccr10 |
| Efnb2 Epha6 | 1.49E-17 | 3.371730269 | 0.287 | 0.109 | 2.52E-14 | 7 | Efnb2 Epha6 |
| Icam1 Itgb21 | 1.78E-17 | 1.006302926 | 0.573 | 0.425 | 3.01E-14 | 7 | Icam1 Itgb2 |
| Tnf Tnfrsf1a | 4.40E-17 | 4.255147756 | 0.093 | 0.005 | 7.43E-14 | 7 | Tnf Tnfrsf1a |
| Egf Fshr | 6.88E-17 | 4.008857122 | 0.193 | 0.048 | 1.16E-13 | 7 | Egf Fshr |
| Fasl Tnfrsf1a | 8.97E-17 | 10.68643133 | 0.07 | 0 | 1.51E-13 | 7 | Fasl Tnfrsf1a |
| Tnf Tnfrsf21 | 1.38E-16 | 5.544492719 | 0.087 | 0.004 | 2.32E-13 | 7 | Tnf Tnfrsf21 |
| Il15 Il2rg | 1.80E-16 | 5.744784563 | 0.093 | 0.006 | 3.03E-13 | 7 | Il15 Il2rg |
| Ccl5 Ccr1 | 2.74E-16 | 3.324622373 | 0.093 | 0.006 | 4.63E-13 | 7 | Ccl5 Ccr1 |
| Hc C5ar1 | 2.78E-16 | 3.482215419 | 0.127 | 0.017 | 4.69E-13 | 7 | Hc C5ar1 |
| Icam1 Il2rg | 3.52E-16 | 6.082492961 | 0.077 | 0.002 | 5.94E-13 | 7 | Icam1 Il2rg |

| | | | | | | | |
|---|---|---|---|---|---|---|---|
| Selplg Sell | 7.69E-16 | 5.696155737 | 0.083 | 0.004 | 1.30E-12 | 7 | Selplg Sell |
| L1cam Fgfr2 | 1.38E-15 | 2.197332303 | 0.257 | 0.094 | 2.32E-12 | 7 | L1cam Fgfr2 |
| S100a9 Tlr4 | 2.11E-15 | 3.719024797 | 0.24 | 0.086 | 3.57E-12 | 7 | S100a9 Tlr4 |
| Hdc Hrh2 | 2.53E-15 | 5.434941843 | 0.107 | 0.012 | 4.28E-12 | 7 | Hdc Hrh2 |
| Tph1 Htr1f | 2.67E-15 | 11.24572125 | 0.063 | 0 | 4.51E-12 | 7 | Tph1 Htr1f |
| Nmb Nmbr | 6.59E-15 | 3.597682181 | 0.167 | 0.04 | 1.11E-11 | 7 | Nmb Nmbr |
| Apoe Ldlr | 8.24E-15 | 1.645367392 | 0.433 | 0.255 | 1.39E-11 | 7 | Apoe Ldlr |
| B2m Tfrc1 | 1.00E-14 | 0.65589309 | 0.843 | 0.651 | 1.70E-11 | 7 | B2m Tfrc |
| Calm1 Pde1b1 | 1.21E-14 | 0.627431734 | 0.793 | 0.731 | 2.05E-11 | 7 | Calm1 Pde1b |
| Tnfsf4 Tnfrsf4 | 1.45E-14 | 11.47418106 | 0.06 | 0 | 2.45E-11 | 7 | Tnfsf4 Tnfrsf4 |
| Pomc Oprm1 | 2.59E-14 | 4.00106648 | 0.077 | 0.004 | 4.38E-11 | 7 | Pomc Oprm1 |
| Il1rn Il1rl2 | 2.62E-14 | 5.096995271 | 0.083 | 0.006 | 4.42E-11 | 7 | Il1rn Il1rl2 |
| Tnf Ltbr | 3.14E-14 | 3.997975124 | 0.08 | 0.005 | 5.31E-11 | 7 | Tnf Ltbr |
| Fn1 Nt5e | 3.33E-14 | 2.648164912 | 0.163 | 0.04 | 5.63E-11 | 7 | Fn1 Nt5e |
| Jag1 Notch21 | 3.92E-14 | 1.188300177 | 0.693 | 0.884 | 6.62E-11 | 7 | Jag1 Notch2 |
| Nampt Insr1 | 5.30E-14 | 0.691556912 | 0.81 | 0.75 | 8.94E-11 | 7 | Nampt Insr |
| Tnf Slc5a11 | 7.91E-14 | 11.38348489 | 0.057 | 0 | 1.33E-10 | 7 | Tnf Slc5a11 |
| Il15 Il2rb | 1.55E-13 | 4.184834835 | 0.08 | 0.006 | 2.62E-10 | 7 | Il15 Il2rb |
| Ntng2 Lrrc4c | 1.87E-13 | 4.964248786 | 0.1 | 0.013 | 3.15E-10 | 7 | Ntng2 Lrrc4c |
| Rtn4 Rtn4rl1 | 2.61E-13 | 1.569771129 | 0.453 | 0.305 | 4.41E-10 | 7 | Rtn4 Rtn4rl1 |
| Icam5 Itgal | 9.34E-13 | 8.083387653 | 0.057 | 0.001 | 1.58E-09 | 7 | Icam5 Itgal |
| Lta Ltbr | 9.52E-13 | 7.611212312 | 0.057 | 0.001 | 1.61E-09 | 7 | Lta Ltbr |
| Camp Egfr | 1.69E-12 | 4.774131295 | 0.133 | 0.031 | 2.85E-09 | 7 | Camp Egfr |
| Hc C5ar2 | 2.11E-12 | 3.355703705 | 0.103 | 0.016 | 3.55E-09 | 7 | Hc C5ar2 |
| Btla Cd79a | 2.33E-12 | 13.16199243 | 0.05 | 0 | 3.94E-09 | 7 | Btla Cd79a |
| Mmp9 Itgam | 2.33E-12 | 12.06387951 | 0.05 | 0 | 3.94E-09 | 7 | Mmp9 Itgam |
| Plau Itgb2 | 2.57E-12 | 4.543730329 | 0.063 | 0.003 | 4.34E-09 | 7 | Plau Itgb2 |
| S100a8 Tlr4 | 3.95E-12 | 3.642125154 | 0.237 | 0.103 | 6.66E-09 | 7 | S100a8 Tlr4 |
| Selplg Itgam | 5.05E-12 | 7.676486252 | 0.053 | 0.001 | 8.53E-09 | 7 | Selplg Itgam |
| Gnas Adcy71 | 6.14E-12 | 0.407263448 | 0.963 | 0.882 | 1.04E-08 | 7 | Gnas Adcy7 |
| Sema7a Itgb1 | 1.39E-11 | 1.343811845 | 0.253 | 0.104 | 2.35E-08 | 7 | Sema7a Itgb1 |
| Itgb3bp Itgb31 | 2.05E-11 | 1.252526891 | 0.41 | 0.239 | 3.45E-08 | 7 | Itgb3bp Itgb3 |
| Lta Tnfrsf1a | 2.73E-11 | 7.642921947 | 0.05 | 0.001 | 4.61E-08 | 7 | Lta Tnfrsf1a |
| B2m Klrd1 | 2.81E-11 | 4.985610836 | 0.083 | 0.011 | 4.74E-08 | 7 | B2m Klrd1 |
| Icam1 Il2ra | 4.56E-11 | 5.568419022 | 0.053 | 0.002 | 7.69E-08 | 7 | Icam1 Il2ra |
| Icam4 Itgb2 | 7.13E-11 | 1.506671893 | 0.4 | 0.275 | 1.20E-07 | 7 | Icam4 Itgb2 |
| Lipc Lrp1 | 2.56E-10 | 3.373278663 | 0.127 | 0.034 | 4.32E-07 | 7 | Lipc Lrp1 |
| Plau Itgb1 | 3.88E-10 | 3.684712509 | 0.067 | 0.007 | 6.56E-07 | 7 | Plau Itgb1 |
| F8 Ldlr | 3.94E-10 | 1.395967623 | 0.463 | 0.356 | 6.65E-07 | 7 | F8 Ldlr |
| Tnfsf13b Tnfrsf13b | 5.25E-10 | 3.212277454 | 0.08 | 0.012 | 8.87E-07 | 7 | Tnfsf13b Tnfrsf13b |
| Tgfb1 Tgfbr11 | 5.94E-10 | 1.187505436 | 0.477 | 0.402 | 1.00E-06 | 7 | Tgfb1 Tgfbr1 |
| Il4 Cd53 | 6.28E-10 | 3.514373383 | 0.077 | 0.011 | 1.06E-06 | 7 | Il4 Cd53 |
| Ptgs2 Alox5 | 6.39E-10 | 2.42318702 | 0.17 | 0.062 | 1.08E-06 | 7 | Ptgs2 Alox5 |
| Il15 Il2ra | 1.26E-09 | 5.55180495 | 0.047 | 0.002 | 2.13E-06 | 7 | Il15 Il2ra |
| Ltf Lrp1 | 1.29E-09 | 0.614779211 | 0.213 | 0.524 | 2.18E-06 | 7 | Ltf Lrp1 |
| Ccl7 Ccr5 | 2.02E-09 | 11.01222604 | 0.037 | 0 | 3.41E-06 | 7 | Ccl7 Ccr5 |
| Ccl7 Ccr1 | 2.02E-09 | 10.46264497 | 0.037 | 0 | 3.41E-06 | 7 | Ccl7 Ccr1 |
| Pdgfb S1pr11 | 2.89E-09 | 0.880976928 | 0.39 | 0.262 | 4.88E-06 | 7 | Pdgfb S1pr1 |
| Ccl2 Ccr5 | 4.33E-09 | 5.862108357 | 0.04 | 0.001 | 7.31E-06 | 7 | Ccl2 Ccr5 |
| Ccl2 Ccr1 | 4.49E-09 | 5.330581345 | 0.04 | 0.001 | 7.59E-06 | 7 | Ccl2 Ccr1 |
| Hgf Cd44 | 5.11E-09 | 2.413330424 | 0.173 | 0.069 | 8.62E-06 | 7 | Hgf Cd44 |

| | | | | | | | |
|---|---|---|---|---|---|---|---|
| Efna5 Epha6 | 5.17E-09 | 2.860653617 | 0.287 | 0.173 | 8.73E-06 | 7 | Efna5 Epha6 |
| Lrpap1 Ldlr1 | 6.20E-09 | 1.061830307 | 0.483 | 0.384 | 1.05E-05 | 7 | Lrpap1 Ldlr |
| Scgb1a1 Lmbr1l | 1.06E-08 | 1.619809734 | 0.507 | 0.457 | 1.78E-05 | 7 | Scgb1a1 Lmbr1l |
| Il1b Il1r2 | 1.10E-08 | 12.61111361 | 0.033 | 0 | 1.86E-05 | 7 | Il1b Il1r2 |
| Cxcl2 Xcr1 | 1.10E-08 | 12.15533323 | 0.033 | 0 | 1.86E-05 | 7 | Cxcl2 Xcr1 |
| Tph1 Htr7 | 1.10E-08 | 10.4886556 | 0.033 | 0 | 1.86E-05 | 7 | Tph1 Htr7 |
| Ccl5 Sdc1 | 1.10E-08 | 10.35820641 | 0.033 | 0 | 1.86E-05 | 7 | Ccl5 Sdc1 |
| Il23a Il12rb1 | 1.10E-08 | 9.667237742 | 0.033 | 0 | 1.86E-05 | 7 | Il23a Il12rb1 |
| Lta Tnfrsf14 | 2.30E-08 | 7.981658399 | 0.037 | 0.001 | 3.88E-05 | 7 | Lta Tnfrsf14 |
| F12 Cd93 | 2.30E-08 | 7.415241835 | 0.037 | 0.001 | 3.88E-05 | 7 | F12 Cd93 |
| Hp Itgam | 2.30E-08 | 7.132971624 | 0.037 | 0.001 | 3.88E-05 | 7 | Hp Itgam |
| Cxcl12 Cxcr4 | 3.79E-08 | 2.822383161 | 0.107 | 0.031 | 6.40E-05 | 7 | Cxcl12 Cxcr4 |
| Calm2 Pde1b1 | 3.89E-08 | 0.592130918 | 0.783 | 0.744 | 6.56E-05 | 7 | Calm2 Pde1b |
| Ccl5 Sdc4 | 3.91E-08 | 3.113017726 | 0.04 | 0.002 | 6.61E-05 | 7 | Ccl5 Sdc4 |
| Efna1 Epha6 | 3.99E-08 | 2.884709817 | 0.277 | 0.173 | 6.74E-05 | 7 | Efna1 Epha6 |
| Thbs1 Itga4 | 4.14E-08 | 2.263921706 | 0.307 | 0.202 | 6.98E-05 | 7 | Thbs1 Itga4 |
| Btla Tnfrsf14 | 4.26E-08 | 6.373841766 | 0.047 | 0.004 | 7.19E-05 | 7 | Btla Tnfrsf14 |
| Proc Itgam | 5.99E-08 | 11.06470897 | 0.03 | 0 | 0.000101064 | 7 | Proc Itgam |
| Ccl26 Ccr1 | 5.99E-08 | 9.523660733 | 0.03 | 0 | 0.000101064 | 7 | Ccl26 Ccr1 |
| Calm1 Insr1 | 1.45E-07 | 0.284735658 | 0.897 | 0.898 | 0.000244153 | 7 | Calm1 Insr |
| Plau Plaur | 1.78E-07 | 6.199575759 | 0.037 | 0.002 | 0.000300891 | 7 | Plau Plaur |
| Plau Itgav | 2.21E-07 | 4.140999565 | 0.043 | 0.004 | 0.000373297 | 7 | Plau Itgav |
| Ccl2 Ccr2 | 3.27E-07 | 10.40138979 | 0.027 | 0 | 0.000552004 | 7 | Ccl2 Ccr2 |
| Tnfsf13b Tnfrsf13c | 3.27E-07 | 9.859627593 | 0.027 | 0 | 0.000552004 | 7 | Tnfsf13b Tnfrsf13c |
| Plau St14 | 3.27E-07 | 9.084465191 | 0.027 | 0 | 0.000552004 | 7 | Plau St14 |
| Gnai2 Mtnr1b | 5.95E-07 | 3.048762744 | 0.073 | 0.017 | 0.001004551 | 7 | Gnai2 Mtnr1b |
| Il1rn Il1r2 | 6.62E-07 | 8.276941733 | 0.03 | 0.001 | 0.001117494 | 7 | Il1rn Il1r2 |
| Tgm2 Itgb1 | 7.21E-07 | 0.552059548 | 0.63 | 0.836 | 0.001216369 | 7 | Tgm2 Itgb1 |
| B2m Klrc1 | 1.06E-06 | 4.025194192 | 0.037 | 0.003 | 0.001789576 | 7 | B2m Klrc1 |
| Adam28 Itga4 | 1.17E-06 | 4.264185477 | 0.073 | 0.018 | 0.001975261 | 7 | Adam28 Itga4 |
| Ccl7 Ccr2 | 1.79E-06 | 10.28314997 | 0.023 | 0 | 0.003028124 | 7 | Ccl7 Ccr2 |
| Tnfsf8 Tnfrsf8 | 1.79E-06 | 10.15782065 | 0.023 | 0 | 0.003028124 | 7 | Tnfsf8 Tnfrsf8 |
| Il1a Il1r2 | 1.79E-06 | 10.05300271 | 0.023 | 0 | 0.003028124 | 7 | Il1a Il1r2 |
| Gzmb Igf2r | 1.79E-06 | 9.292951009 | 0.023 | 0 | 0.003028124 | 7 | Gzmb Igf2r |
| Ltf Gp91 | 2.26E-06 | 0.11927525 | 0.053 | 0.172 | 0.003820776 | 7 | Ltf Gp9 |
| Tnfsf9 Traf2 | 3.89E-06 | 3.880494866 | 0.07 | 0.018 | 0.006571199 | 7 | Tnfsf9 Traf2 |
| Ntn4 Dcc1 | 4.48E-06 | 0.341175637 | 0.257 | 0.499 | 0.007559189 | 7 | Ntn4 Dcc |
| Lamc1 Itgav1 | 8.77E-06 | 1.000555351 | 0.43 | 0.371 | 0.014804968 | 7 | Lamc1 Itgav |
| Uba52 Fshr | 9.90E-06 | 9.840789878 | 0.02 | 0 | 0.016716956 | 7 | Uba52 Fshr |
| Ccl3 Ccr4 | 9.90E-06 | 9.516269788 | 0.02 | 0 | 0.016716956 | 7 | Ccl3 Ccr4 |
| Fasl Fas | 9.90E-06 | 8.584215347 | 0.02 | 0 | 0.016716956 | 7 | Fasl Fas |
| Ccl25 Ccr9 | 1.38E-05 | 3.220946519 | 0.04 | 0.006 | 0.023330822 | 7 | Ccl25 Ccr9 |
| Liph Lpar2 | 1.47E-05 | 3.508683612 | 0.06 | 0.015 | 0.024771745 | 7 | Liph Lpar2 |
| Mfge8 Itgav1 | 1.80E-05 | 0.448215519 | 0.607 | 0.602 | 0.030359688 | 7 | Mfge8 Itgav |
| Ccl8 Ccr5 | 1.90E-05 | 9.167087184 | 0.023 | 0.001 | 0.032057844 | 7 | Ccl8 Ccr5 |
| Ccl8 Ccr1 | 1.90E-05 | 8.752515885 | 0.023 | 0.001 | 0.032057844 | 7 | Ccl8 Ccr1 |
| Tnfsf9 Tnfrsf9 | 1.90E-05 | 7.618612701 | 0.023 | 0.001 | 0.032057844 | 7 | Tnfsf9 Tnfrsf9 |
| C3 Cd19 | 1.91E-05 | 5.72258377 | 0.023 | 0.001 | 0.032247271 | 7 | C3 Cd19 |
| F10 Itgam | 1.94E-05 | 4.410211285 | 0.023 | 0.001 | 0.03282197 | 7 | F10 Itgam |
| Vegfa Gpc11 | 2.05E-05 | 0.62539551 | 0.01 | 0.079 | 0.03455567 | 7 | Vegfa Gpc1 |
| Calm1 Grm4 | 2.63E-05 | 3.453713767 | 0.043 | 0.008 | 0.044333926 | 7 | Calm1 Grm4 |

**Table S12.** NICHES intercellular communication: unique and common genes

| Nox-Hox unique genes | | Hox-NoxRec unique genes | | | | | | Hox-NoxRecExer unique genes | | | Genes common to all conditions | Genes common to Nox-Hox Hox-NoxRec | | Genes common to Nox-Hox Hox-NoxRecExer | Genes common to Hox-NoxRecExer Hox-Nox |
|---|---|---|---|---|---|---|---|---|---|---|---|---|---|---|---|
| Adcy7 | Robo4 | A2m | Col11a1 | Fzd1 | Kcnj4 | Ogfr | Slc18a2 | Adam28 | Htr7 | Pde1b1 | Cd44 | Abca1 | Plxna4 | Asgr2 | Ccl25 |
| Adcy9 | Rps19 | Ackr2 | Col17a1 | Fzd2 | Kcnq3 | Olfm2 | Slc44a5 | Adcy71 | Icam5 | Pecam1 | Cd47 | Acvr1 | Plxnb2 | B2m | Efna1 |
| Adora2b | Scarb1 | Ackr4 | Col2a1 | Fzd3 | Kidins220 | Oxt | Slc45a3 | Adrb2 | Ifng | Pkm | Egfr | Acvr1b | Plxnd1 | Bgn | Efna5 |
| Adra2a | Scarf1 | Acvr2b1 | Col6a2 | Fzd4 | Kit | Oxtr | Sorbs1 | Alox5 | Ifngr1 | Plau | Fgfr2 | Acvr2a | Psap | C5ar1 | Epha6 |
| Ahsg | Scgb3a1 | Acvrl1 | Col6a3 | Fzd5 | Kng1 | Pdap1 | Sorcs3 | Alox5ap | Ifngr2 | Plxnc1 | Igf1r | Acvr2b | Ptn | Ccr5 | Fas |
| Angpt1 | Selp | Adam12 | Col7a1 | Fzd7 | Lama2 | Pde1a | Sort1 | Amh | Igf2r | Podxl | Itga4 | Adam10 | Rarres2 | Cd34 | Hgf |
| Angptl4 | Sema3a | Adcy1 | Col8a1 | Fzd8 | Lama3 | Pde1c | Spp1 | Atrn | Il10 | Ptpra1 | Itgb1 | Adam17 | Rgmb | Cd93 | Il15 |
| Atp1a3 | Sema3c | Adcy8 | Ctf1 | Gdf10 | Lamb3 | Pdgfa | Sstr2 | Btla | Il10ra | Rhag | Lrp1 | Agrn | S1pr1 | Col4a3 | Il18 |
| Bmp8b | Sema3d | Adcyap1r1 | Dcbld2 | Gdf11 | Lamc2 | Pdgfc | Stra6 | C3 | Il10rb | Rtn4rl1 | Notch2 | Angpt2 | Sdc3 | Col4a4 | Il1b |
| Bst1 | Sema3f | Adipoq | Dchs1 | Gdf9 | Lamc3 | Pdgfd | Tbc1d24 | C5ar2 | Il12rb1 | S100a8 | Sdc1 | Axl | Sema5a | Cxcl12 | Il1rl2 |
| Cd24a | Sema3g | Adipor1 | Dcn | Gdnf | Lamp1 | Pdgfra | Tbxa2r | Camp | Il15 | S100a9 | Tlr4 | Bmp4 | Slit2 | Dcc | Il1rn |
| Cd36 | Sema4c | Adipor2 | Ddr1 | Gfra1 | Lepr | Pdgfrb | Tgfb2 | Ccl2 | Il1a | S1pr11 | Calm1 | Bmp6 | Tctn1 | Dll1 | Il4 |
| Cd81 | Sema6d | Adora1 | Ddr2 | Gfra2 | Lgals3bp | Penk | Tgfb3 | Ccl26 | Il1r2 | S1pr4 | Calm2 | Bmpr1a | Tek | Dll4 | Il6st |
| Cmklr1 | Serping1 | Agrp | Dlk1 | Gnb3 | Lgr4 | Pf4 | Tgfbr1 | Ccl3 | Il1rap | Scgb1a1 | Gnas | Bmpr2 | Tfpi | F8 | Itgav |
| Col4a2 | Sfrp1 | Agtr2 | Dll3 | Gnrh1 | Lgr5 | Pgf | Tgfbr3 | Ccl4 | Il23a | Scn4a | Fn1 | Cav1 | Tgfa | Icam1 | Itgb3 |
| Cp | Sirpa | Alk | Dusp18 | Gnrhr | Lgr6 | Pla2g10 | Thbd | Ccl5 | Il27 | Sell | Gnai2 | Col18a1 | Timp3 | Icam2 | Itgb3bp |
| Csf1 | Slc40a1 | Amelx | Dysf | Gpc1 | Lhcgr | Pla2r1 | Timp1 | Ccl7 | Il27ra | Selplg | Egf | Col1a1 | Tnc | Icam4 | Lamc1 |
| Cthrc1 | Sost | Angptl1 | Ecm1 | Gpc4 | Lif | Plscr4 | Timp2 | Ccl8 | Il2ra | Sema4a | L1cam | Col1a2 | Unc5b | Itgb2 | Ldlr |
| Cx3cl1 | Tfrc | Angptl3 | Eda | Gpr1 | Lingo1 | Plxna2 | Tlr41 | Ccr1 | Il2rb | Slc5a11 | Efnb2 | Col3a1 | Vcan | Jag2 | Lifr |
| Cx3cr1 | Thbs2 | Anxa1 | Eda2r | Gpr135 | Lrp11 | Pon2 | Tmem671 | Ccr10 | Il2rg | Sorl11 | Adam2 | Col4a1 | Vegfb | Ltf | Lrpap1 |
| Dkk2 | Tie1 | Apcdd1 | Edil3 | Grin2d | Lrp41 | Prlr | Tmprss6 | Ccr2 | Ins1 | St14 | Tgm2 | Col4a5 | | Mfng | Lrrc4c |
| Edn1 | Tlr2 | Apod | Ednra | Grm3 | Lrp6 | Pros1 | Tnfrsf11b | Ccr4 | Itgal | Tfrc1 | Adam15 | Col4a6 | | Nampt | Mfge8 |
| Ednrb | Tlr7 | Aqp1 | Efemp1 | Grm7 | Ltbp3 | Pth1r | Tnfrsf12a | Ccr9 | Itgam | Tgfbr11 | Adam9 | Col5a1 | | Nestn | Mmp9 |
| Epo | Tmem67 | Atp6ap2 | Efemp2 | Gstp1 | Magec1 | Pthlh | Tnfsf12 | Cd14 | Itgav1 | Tlr9 | Apoe | Col5a2 | | Ntn4 | Plaur |
| Epor | Tnfrsf10b | Axl1 | Efna2 | Has2 | Mc5r | Ptprf | Tshr | Cd180 | Itgb21 | Tnf | Jag1 | Col6a1 | | Pdgfb | Pomc |
| Fgl1 | Tnfsf10 | BC055324 | Efna4 | Hbegf | Mcam | Ptprm | Tyro3 | Cd19 | Itgb31 | Tnfrsf13b | Psen1 | Csf1r | | Ptdss1 | Proc |
| Fgl2 | Trem2 | Bambi | Efnb1 | Hhipl2 | Mmp2 | Ptprs | Vangl1 | Cd40 | Itgb7 | Tnfrsf13c | Vegfa | Epha4 | | Sema4d | Ptgs2 |
| Flt4 | Tyrobp | Bcam | Efnb3 | Hmgb1 | Mtnr1a | Ptpru | Vldlr | Cd48 | Jmjd6 | Tnfrsf14 | Calr | Ephb1 | | Tlr1 | Rtn4 |
| Glp2r | Unc5a | Bdnf | Egfr1 | Hsp90aa1 | Musk | Ptprz1 | Vtn | Cd53 | Kcnn4 | Tnfrsf1a | Hras | Ephb4 | | Tnfrsf21 | Sdc4 |
| Gp9 | Vegfc | Bmp2 | Eng | Hspa1a | Mylk | Pvr | Wnt11 | Cd72 | Kcnq5 | Tnfrsf1b | Hsp90b1 | Fgf2 | | Vcam1 | Sema7a |
| Gpc3 | Vwf | Bmp5 | Epha1 | Htr2a | Mylk2 | Qdpr | Wnt3a | Cd74 | Kir3dl1 | Tnfrsf4 | App | Fgfr1 | | | Sorl1 |
| Gpihbp1 | | Bmp7 | Epha3 | Htr6 | Myoc | Rbp4 | Wnt4 | Cd79a | Klrc1 | Tnfrsf8 | | Fgfr3 | | | Tgfb1 |
| Hfe | | Bmpr1b | Epha5 | Igf1 | Ncan | Rgma | Wnt5a | Cfh | [illegible] | Tnfrsf9 | | Flt1 | | | Tgfbr2 |
| Itga6 | | C4b | Epha7 | Igfbp4 | Ncp | Rims1 | Znrf3 | Cxcl2 | Lipc | Tnfsf13b | | Fzd6 | | | Thbs1 |
| Itgb4 | | Cachd1 | Ephb2 | Il13ra1 | Nec1 | Rims2 | | Cxcr4 | [illegible] | Tnfsf14 | | Gas6 | | | Traf2 |
| Kdr | | Cacna1c | Ephb3 | Il13ra2 | Ngf | Robo1 | | Dec1 | Lmbr1l | Tnfsf4 | | Hspg2 | | | Uba52 |
| Kel | | Calcr | Ephb6 | Il17rc | Nlgn1 | Robo2 | | Dscam | Lpar2 | Tnfsf8 | | Inha | | | As |
| Kitl | | Cav11 | Erbb2 | Il1r1 | Nlgn3 | Ror1 | | Ebi3 | Lrp1b | Tnfsf9 | | Insr | | | |
| Lama1 | | Ccl19 | Erbb4 | Il1rapl1 | Notch3 | Ror2 | | Erap1 | Lrrc4 | Tph1 | | Itga1 | | | |
| Ly96 | | Cd109 | Ereg | Il34 | Npnt | Rspo1 | | F10 | Lta | Trpc5 | | Itga2b | | | |
| Lyve1 | | Cd151 | F13a1 | Il6 | Nppa | Rspo2 | | F12 | Ltb | Vim | | Itga9 | | | |
| Mag | | Cd40 | F2 | Inhba | Npr1 | Rspo3 | | Fasl | Ltbr | Xcr1 | | Lama4 | | | |
| Marco | | Cd63 | F2r | Inhbb | Npr3 | Rtn4r | | Fbln2 | Ly86 | | | Lama5 | | | |
| Mertk | | Cd82 | F3 | Itga10 | Npy | Ryk | | Fshr | Mgrn1 | | | Lamb1 | | | |
| Met | | Cd9 | Fap | Itga11 | Npy1r | S1pr3 | | Gp91 | Mtnr1b | | | Lin7c | | | |
| Mfap2 | | Cdh1 | Farp2 | Itga2 | Npy4r | Sdc2 | | Gpc11 | Nestn1 | | | Lpl | | | |
| Mfap5 | | Cdh2 | Fat4 | Itga3 | Nrcam | Sema3b | | Gpi1 | Nmb | | | Lrp5 | | | |
| Notch4 | | Cdon | Fbn1 | Itga5 | Nrg1 | Sema3e | | Grm1 | Nmbr | | | Lrp6 | | | |
| Nrxn3 | | Chad | Fgf1 | Itga7 | Nrg2 | Sema4b | | Grm4 | Notch2l | | | Mdk | | | |
| Ntn1 | | Chrm3 | Fgf14 | Itga8 | Nrg3 | Sema4f | | Grm5 | Nps | | | Ncam1 | | | |
| P2ry12 | | Clcf1 | Fgf16 | Itga91 | Nrg4 | Sema4g | | Gzmb | Npsr1 | | | Nid1 | | | |
| Papln | | Clec11a | Fgf18 | Itgbl1 | Nrxn2 | Serpinc1 | | Hc | Nt5e | | | Nlgn2 | | | |
| Pde1b | | Cntf | Fgf22 | Itgb5 | Ntf3 | Serpine1 | | Hdc | Ntng2 | | | Notch1 | | | |
| Pgf | | Cntn2 | Fgf5 | Itgb6 | Ntng1 | Serpine2 | | Hp | Nucb2 | | | Nrp1 | | | |
| Plat | | Cntnap1 | Fgf9 | Itgb8 | Ntrk2 | Sftpc | | Hrh1 | Oprm1 | | | Nrp2 | | | |
| Ptpra | | Cntnap2 | Fgfrl1 | Kcnd1 | Ntrk3 | Shank1 | | Hrh2 | Osm | | | Plxna1 | | | |
| Ptprb | | Fgf11 | Fxyd5 | Kcnj15 | Nxph1 | Slc17a7 | | Htr1f | P2rx7 | | | Plxna3 | | | |

**Table S13**. Key geometric and passive biomechanical metrics for the proximal pulmonary artery in juvenile (3 to 8 weeks of age) and adult (8 to 13 weeks of age) mice exposed to 5 weeks of hypoxia with a subsequent 5 weeks of normoxic recovery without or with voluntary exercise.

| | | | | | | | | | | |
|---|---|---|---|---|---|---|---|---|---|---|
| | Normoxic $FiO_2$ 20% | 5wk FiO2 10% then 5wk $FiO_2$ 21% | | | Normoxic $FiO_2$ 20% | 5wk FiO2 10%, then 5wk $FiO_2$ 21% | | +an additional 5wkFiO2 21% | | Normoxic $FiO_2$ 20% |
| | age-matched control | 5wk FiO2 10%, then 5wk $FiO_2$ 21%+Exercise | | | age-matched control | 5wk FiO2 10%, then 5wk $FiO_2$ 21%+Exercis | | | | age-match control |
| | F8 NOX | F8 HOX | F13R | F13RE | F13 NOX | F13 HOX | F18R | F18RE | F23R | F18 NOX |
| | n = 4 | n = 5 | n=3 | n=5 | n = 6 | n = 5 | n=5 | n=3 | n=5 | n = 4 |
| **Unloaded Dimensions** | | | | | | | | | | |
| Wall Thickness (µm) | 63.72 ± 4.91 | 62.81 ± 0.69 | 66.66 ± 1.33 | 68.89 ± 2.10 | 66.50 ± 3.18 | 74.94 ± 1.52 | 66.50 ± 3.35 | 69.62 ± 4.11 | 71.34 ± 1.98 | 67.71 ± 1.62 |
| Outer Diameter (µm) | 810.83 ± 5.59 | 865.23 ± 6.73 | 826.43 ± 17.26 | 884.97 ± 17.08 | 813.07 ± 28.59 | 906.61 ± 17.42 | 915.55 ± 12.83 | 955.11 ± 18.14 | 953.78 ± 30.52 | 831.52 ± 9.32 |
| Axial Length (mm) | 2.31 ± 0.13 | 3.06 ± 0.20 | 2.54 ± 0.20 | 2.77 ± 0.15 | 2.39 ± 0.14 | 3.01 ± 0.29 | 2.78 ± 0.09 | 2.95 ± 0.52 | 3.25 ± 0.10 | 2.54 ± 0.05 |
| **Loaded Dimensions** | **P = 15.0** | **P = 25.0** | **P = 15.0** | **P = 15.0** | **P = 15.0** | **P = 25.0** | **P = 15.0** | **P = 15.0** | **P = 15.0** | **P = 15.0** |
| Wall Thickness (µm) | 29.20 ± 2.12 | 31.16 ± 0.54 | 36.98 ± 1.32 | 39.80 ± 1.21 | 28.18 ± 1.01 | 34.80 ± 1.30 | 34.85 ± 1.40 | 44.95 ± 4.57 | 37.24 ± 1.81 | 30.31 ± 1.20 |
| Outer Diameter (µm) | 1110.37 ± 22.31 | 1229.56 ± 17.65 | 1040.74 ± 23.19 | 1101.41 ± 36.49 | 1122.96 ± 51.97 | 1314.30 ± 31.25 | 1209.75 ± 14.51 | 1160.82 ± 26.95 | 1232.20 ± 54.73 | 1211.64 ± 60.03 |
| Inner Radius (µm) | 525.99 ± 10.32 | 583.62 ± 8.99 | 483.39 ± 12.41 | 510.91 ± 18.67 | 533.30 ± 26.06 | 622.35 ± 16.27 | 570.03 ± 6.07 | 535.46 ± 15.93 | 578.86 ± 28.94 | 575.51 ± 30.83 |
| *in vivo* Axial Stretch ($\lambda_z^{iv}$) | 1.51 ± 0.03 | 1.35 ± 0.01 | 1.37 ± 0.04 | 1.33 ± 0.02 | 1.60 ± 0.03 | 1.40 ± 0.03 | 1.38 ± 0.02 | 1.38 ± 0.02 | 1.42 ± 0.03 | 1.42 ± 0.04 |
| *in vivo* Circumferential Stress ($\lambda_\theta^{iv}$) | 1.45 ± 0.03 | 1.49 ± 0.02 | 1.32 ± 0.02 | 1.30 ± 0.02 | 1.48 ± 0.04 | 1.54 ± 0.04 | 1.38 ± 0.00 | 1.38 ± 0.00 | 1.35 ± 0.02 | 1.52 ± 0.08 |
| **Cacucy Stresses (kPa)** | | | | | | | | | | |
| Circumferential, $\sigma_\theta$ | 36.49 ± 2.30 | 62.54 ± 1.72 | 26.24 ± 1.44 | 25.81091 ± 1.4127 | 38.12 ± 2.37 | 60.06 ± 3.31 | 32.86 ± 0.96 | 32.86 ± 0.96 | 31.71 ± 3.37 | 38.34 ± 3.42 |
| Axial, $\sigma_z$ | 45.01 ± 3.23 | 48.63 ± 1.76 | 29.18 ± 3.57 | 28.82 ± 1.54 | 50.49 ± 3.02 | 48.28 ± 3.21 | 32.70 ± 1.04 | 32.70 ± 1.04 | 33.35 ± 2.16 | 36.51 ± 3.23 |
| **Linearized Stiffness (MPa)** | | | | | | | | | | |
| Circumferential, $C_{\theta\theta\theta\theta}$ | 0.13 ± 0.01 | 0.39 ± 0.01 | 0.12 ± 0.01 | 0.11 ± 0.01 | 0.13 ± 0.01 | 0.35 ± 0.03 | 0.13 ± 0.00 | 0.11 ± 0.01 | 0.15 ± 0.03 | 0.15 ± 0.02 |
| Axial, $C_{zzzz}$ | 0.22 ± 0.01 | 0.37 ± 0.02 | 0.18 ± 0.04 | 0.16 ± 0.01 | 0.24 ± 0.02 | 0.35 ± 0.02 | 0.18 ± 0.01 | 0.12 ± 0.02 | 0.18 ± 0.03 | 0.17 ± 0.02 |
| **Distensibility ($mmHg^{-1}$)** | 336.66 ± 23.52 | 198.21 ± 11.02 | 210.17 ± 19.79 | 225.67 ± 15.22 | 387.44 ± 36.69 | 217.57 ± 21.24 | 228.71 ± 6.28 | 182.27 ± 11.58 | 211.14 ± 23.80 | 300.62 ± 33.77 |
| **Disetnsibility (1/MPa)** | 0.0449 ± 0.0031 | 0.0264 ± 0.0015 | 0.03 ± 0.00 | 0.03 ± 0.00 | 0.05 ± 0.00 | 0.03 ± 0.00 | 0.03 ± 0.00 | 0.02 ± 0.00 | 0.03 ± 0.00 | 0.0401 ± 0.0045 |
| **PWV (m/s)** | 0.15 ± 0.01 | 0.17 ± 0.01 | 0.17 ± 0.01 | 0.16 ± 0.01 | 0.14 ± 0.01 | 0.17 ± 0.01 | 0.16 ± 0.00 | 0.18 ± 0.01 | 0.17 ± 0.01 | 0.16 ± 0.01 |
| **Stored Energy (kPa)** | 13.15 ± 0.98 | 13.36 ± 0.61 | 6.79 ± 0.94 | 6.40 ± 0.45 | 15.12 ± 1.12 | 13.84 ± 1.28 | 8.90 ± 0.30 | 4.91 ± 1.07 | 8.54 ± 0.74 | 11.80 ± 1.04 |

**Table S14.** Best-fit values of the material parameters in the (pseudo)strain-energy function describing the passive mechanical behavior of the proximal pulmonary artery in juvenile (3 to 8 weeks of age) and adult (8 to 13 weeks of age) mice exposed to 5 weeks of hypoxia with a subsequent 5 weeks of normoxic recovery without or with voluntary exercise.

| | Normoxic $FiO_2$ 20% age-matched control | 5wk FiO2 10% then 5wk $FiO_2$ 21% / 5wk FiO2 10%, then 5wk $FiO_2$ 21%+Exercise | | | Normoxic $FiO_2$ 20% age-matched control | 5wk FiO2 10% then 5wk $FiO_2$ 21% / 5wk FiO2 10%, then 5wk $FiO_2$ 21%+Exercise | | +an additional 5wkFiO2 21% | | Normoxic $FiO_2$ 20% age-match control |
|---|---|---|---|---|---|---|---|---|---|---|
| | F8 NOX | F8 HOX | F13R | F13RE | F13 NOX | F13 HOX | F18R | F18RE | F23R | F18 NOX |
| | n = 4 | n = 5 | n=3 | n=5 | n = 6 | n = 5 | n=5 | n=3 | n=5 | n = 4 |
| **Elastic Fibers** | | | | | | | | | | |
| $c$ (kPa) | 12.913 | 12.659 | 10.513 | 11.815 | 11.335 | 9.919 | 13.061 | 13.853 | 10.871 | 11.986 |
| **Axial Collagen** | | | | | | | | | | |
| $c_1^z$ (kPa) | 1.093 | 0.131 | 0.110 | 0.165 | 0.288 | 0.576 | 0.093 | 0.099 | 0.077 | 0.063 |
| $c_2^z$ | 0.919 | 3.945 | 3.474 | 3.675 | 0.906 | 2.274 | 3.188 | 8.263 | 3.907 | 2.781 |
| **Circumferential CollagenCollagen** | | | | | | | | | | |
| $c_1^θ$ (kPa) | 1.404 | 3.356 | 2.202 | 2.041 | 0.648 | 3.449 | 1.438 | 2.801 | 2.164 | 0.607 |
| $c_2^θ$ | 0.200 | 0.231 | 0.187 | 0.274 | 0.468 | 0.073 | 1.364 | 0.247 | 2.198 | 0.502 |
| **Diagonal Collagen** | | | | | | | | | | |
| $c_1^{3,4}$ (kPa) | 1.932 | 3.447 | 3.163 | 3.759 | 3.306 | 3.559 | 2.305 | 3.880 | 3.103 | 2.431 |
| $c_2^{3,4}$ | 0.565 | 1.329 | 1.157 | 1.113 | 0.363 | 0.960 | 1.128 | 2.252 | 0.873 | 0.645 |
| $\alpha_0$ (deg) | 41.500 | 41.587 | 42.129 | 40.505 | 42.522 | 42.643 | 42.649 | 40.800 | 44.612 | 41.637 |
| **Error (RMSE)** | 0.10 | 0.07 | 0.08 | 0.088 | 0.09 | 0.07 | 0.08 | 0.084 | 0.103 | 0.08 |

**Table S15.** Best-fit values of the material parameters in the (pseudo)strain-energy function describing the passive mechanical behavior of the proximal pulmonary artery in adult (8 to 13 weeks of age) mice exposed to 5 weeks of hypoxia with concurrent drug treatment and/or forced exercise. Also shown are values for adult (8 to 13 weeks of age) *Ccr2*$^{-/-}$ mice with and without 5 weeks of exposure to hypoxia.

| | Normoxic control | 5 weeks Hypoxic Exposure | 5 weeks Hypoxic Exposure+ forced exercise (75% VO2max) | 5 weeks Hypoxic Exposure+ Rapamycin | 5 weeks Hypoxic Exposure+ Metformin (chow) | 5 weeks Hypoxic Exposure + Metformin | Ccr2$^{-/-}$ normoxic control | Ccr2$^{-/-}$ 5 weeks Hypoxic Exposure |
|---|---|---|---|---|---|---|---|---|
| | F13 NOX | F13 HOX | F13 FE | F13 RAPA | F13 METC | F13 METW | FCcr2KO13 NOX | FCcr2KO13 HOX |
| | n = 6 | n = 5 | n = 5 | n = 6 | n=4 | n=4 | n = 4 | n = 5 |
| **Elastic Fibers** | | | | | | | | |
| $c$ (kPa) | 11.335 | 9.919 | 8.052 | 4.981 | 7.47661726 | 5.734685178 | 10.352 | 8.050 |
| **Axial Collagen** | | | | | | | | |
| $c_1^1$ (kPa) | 0.288 | 0.576 | 1.564 | 0.457 | 0.626805426 | 2.717494886 | 3.002 | 0.680 |
| $c_2^1$ | 0.906 | 2.274 | 2.210 | 5.086 | 2.255432268 | 5.387251469 | 2.972 | 3.060 |
| **Circumferential CollagenCollagen** | | | | | | | | |
| $c_1^2$ (kPa) | 0.648 | 3.449 | 4.218 | 6.660 | 5.64400166 | 5.639335156 | 1.436 | 4.890 |
| $c_2^2$ | 0.468 | 0.073 | 0.074 | 0.000 | 0.033130192 | 3.81257E-14 | 0.881 | 0.020 |
| **Diagonal Collagen** | | | | | | | | |
| $c_1^{3,4}$ (kPa) | 3.306 | 3.559 | 4.728 | 8.789 | 3.939857315 | 7.652739446 | 6.160 | 4.350 |
| $c_2^{3,4}$ | 0.363 | 0.960 | 0.875 | 0.890 | 0.980481522 | 1.054010103 | 0.795 | 0.950 |
| $\alpha_0$ (deg) | 42.522 | 42.643 | 36.194 | 32.955 | 37.35372043 | 33.98800661 | 37.990 | 37.550 |
| **Error (RMSE)** | 0.09 | 0.07 | 0.083 | 0.078 | 0.068329767 | 0.070713419 | 0.12 | 0.07 |

**Table S16.** Key geometric and passive biomechanical metrics for the proximal pulmonary artery in adult (8 to 13 weeks of age) mice exposed to 5 weeks of hypoxia with concurrent drug treatment and/or forced exercise. Also shown are values for adult (8 to 13 weeks of age) *Ccr2*$^{-/-}$ mice with and without 5 weeks of exposure to hypoxia.

| | Normoxic control | 5 weeks Hypoxic Exposure | 5 weeks Hypoxic Exposure + force exercise (75% VO2max) | 5 weeks Hypoxic Exposure + Rapamycin | 5 weeks Hypoxic Exposure + Metformin (water) | 5 weeks Hypoxic Exposure + Metformin (chow) | Ccr2$^{-/-}$ normoxic control | Ccr2$^{-/-}$ 5 weeks Hypoxic Exposure |
|---|---|---|---|---|---|---|---|---|
| | **F13 NOX** | **F13 HOX** | **F13 FE** | **F13 RAPA** | **F13 METW** | **F13 METC** | **FCcr2KO13 NOX** | **FCcr2KO13 HOX** |
| | **n = 5** | **n = 5** | **n = 5** | **n = 6** | **n=4** | **n=4** | **n = 4** | **n = 5** |
| **Unloaded Dimensions** | | | | | | | | |
| Wall Thickness (μm) | 66.50 ± 3.18 | 74.94 ± 1.52 | 62.20 ± 2.20 | 61.22 ± 1.15 | 62.54 ± 2.40 | 70.30 ± 1.33 | 58.55 ± 1.60 | 66.99 ± 3.98 |
| Outer Diameter (μm) | 813.07 ± 28.59 | 906.61 ± 17.42 | 723.94 ± 9.94 | 695.52 ± 25.78 | 757.22 ± 48.30 | 750.94 ± 9.80 | 792.11 ± 25.32 | 807.52 ± 15.44 |
| Axial Length (mm) | 2.39 ± 0.14 | 3.01 ± 0.29 | 2.79 ± 0.07 | 2.16 ± 0.14 | 2.59 ± 0.2 | 3.01 ± 0.30 | 3.01 ± 0.13 | 2.75 ± 0.11 |
| **Loaded Dimensions** | **P = 15.0** | **P = 25.0** | **P = 25.0** | **P = 25.0** | **P = 25.0** | **P = 25.0** | **P = 25.0** | **P = 25.0** |
| Wall Thickness (μm) | 28.18 ± 1.01 | 34.80 ± 1.30 | 28.42 ± 1.30 | 30.12 ± 0.80 | 29.57 ± 3.53 | 34.47 ± 0.76 | 23.86 ± 1.20 | 31.47 ± 2.05 |
| Outer Diameter (μm) | 1122.96 ± 51.97 | 1314.30 ± 31.25 | 1084.97 ± 13.16 | 1019.33 ± 48.24 | 1170.55 ± 32.07 | 1043.91 ± 7.19 | 1345.79 ± 69.94 | 1200.30 ± 19.35 |
| Inner Radius (μm) | 533.30 ± 26.06 | 622.35 ± 16.27 | 514.06 ± 7.47 | 479.54 ± 24.32 | 555.70 ± 17.62 | 487.49 ± 2.97 | 649.04 ± 33.77 | 568.68 ± 10.36 |
| *in vivo* Axial Stretch ($\lambda_z^{iv}$) | 1.60 ± 0.03 | 1.40 ± 0.03 | 1.37 ± 0.03 | 1.32 ± 0.05 | 1.30 ± 0.09 | 1.38 ± 0.01 | 1.38 ± 0.10 | 1.35 ± 0.02 |
| *in vivo* Circumferential Stress ($\lambda_\theta^{iv}$) | 1.48 ± 0.04 | 1.54 ± 0.04 | 1.60 ± 0.03 | 1.55 ± 0.05 | 1.66 ± 0.10 | 1.48 ± 0.01 | 1.80 ± 0.08 | 1.58 ± 0.02 |
| **Cacucy Stresses (kPa)** | | | | | | | | |
| Circumferential, $\sigma_\theta$ | 38.12 ± 2.37 | 60.06 ± 3.31 | 60.92 ± 3.52 | 53.35 ± 3.43 | 66.55 ± 11.37 | 47.20 ± 0.84 | 74.90 ± 7.96 | 61.34 ± 4.27 |
| Axial, $\sigma_z$ | 50.49 ± 3.02 | 48.28 ± 3.21 | 52.58 ± 2.05 | 48.75 ± 6.01 | 48.73 ± 11.19 | 42.83 ± 2.39 | 91.59 ± 7.66 | 47.20 ± 3.81 |
| **Linearized Stiffness (MPa)** | | | | | | | | |
| Circumferential, $C_{\theta\theta\theta\theta}$ | 0.13 ± 0.01 | 0.35 ± 0.03 | 0.31 ± 0.02 | 0.27 ± 0.02 | 0.29 ± 0.01 | 0.24 ± 0.01 | 0.58 ± 0.14 | 0.33 ± 0.03 |
| Axial, $C_{zzzz}$ | 0.24 ± 0.02 | 0.35 ± 0.02 | 0.41 ± 0.02 | 0.39 ± 0.04 | 0.29 ± 0.02 | 0.36 ± 0.03 | 0.49 ± 0.03 | 0.37 ± 0.03 |
| **Distensibility (mmHg$^{-1}$)** | 387.44 ± 36.69 | 217.57 ± 21.24 | 258.43 ± 8.16 | 196.93 ± 13.60 | 205.17 ± 18.22 | 204.83 ± 7.33 | 279.07 ± 50.43 | 215.01 ± 13.86 |
| **Disetnsibility (1/MPa)** | 0.0517 ± 0.0049 | 0.0290 ± 0.0028 | 0.0345 ± 0.0011 | 0.0263 ± 0.0018 | 0.0274 ± 0.0024 | 0.0273 ± 0.0010 | 0.0372 ± 0.0100 | 0.0300 ± 0.0000 |
| **PWV (m/s)** | 0.14 ± 0.01 | 0.17 ± 0.01 | 0.15 ± 0.00 | 0.18 ± 0.01 | 0.17 ± 0.01 | 0.17 ± 0.00 | 0.17 ± 0.02 | 0.17 ± 0.01 |
| **Stored Energy (kPa)** | 15.12 ± 1.12 | 13.84 ± 1.28 | 14.53 ± 0.61 | 11.90 ± 1.30 | 14.56 ± 3.68 | 10.74 ± 0.45 | 23.14 ± 2.34 | 13.51 ± 1.04 |
| **Active Contractility** | **n = 5** | **n = 5** | | **n = 4** | | | | |
| Outer Diameter Change KCl (%) | 31 ± 5.04 | 14 ± 4.32 | | 21.2 ± 3.57 | | | | |
| Outer Diameter Change PE (%) | 35 ± 4.86 | 28 ± 3.65 | | 15.4 ± 2.78 | | | | |
